# Rapid emergence of extensively drug-resistant *Shigella sonnei* in France

**DOI:** 10.1101/2022.09.07.506808

**Authors:** Sophie Lefèvre, Elisabeth Njamkepo, Sarah Feldman, Corinne Ruckly, Isabelle Carle, Monique Lejay-Collin, Laëtitia Fabre, Iman Yassine, Lise Frezal, Maria Pardos de la Gandara, Arnaud Fontanet, François-Xavier Weill

## Abstract

*Shigella sonnei*, the main cause of bacillary dysentery in high-income countries, has become increasingly resistant to antibiotics. We monitored the antimicrobial susceptibility of 7,121 *S. sonnei* isolates collected in France between 2005 and 2021. We identified a dramatic increase in the proportion of extensively drug-resistant (XDR) isolates (i.e., simultaneously resistant to ciprofloxacin, third-generation cephalosporins and azithromycin), to 22.3% of all *S. sonnei* isolates in 2021. Our genomic analysis identified 13 different clusters of XDR isolates descended from a ciprofloxacin-resistant sublineage originating from South Asia. The 164 XDR isolates detected were resistant to azithromycin, principally through a pKSR100-like plasmid, and to third-generation cephalosporins through various genes and plasmids. This rapid emergence of XDR *S. sonnei* in different transmission networks, particularly among men who have sex with men, is a matter of concern, and good laboratory-based surveillance of *Shigella* infections will be crucial for informed decision-making and appropriate public health action.

## TEXT

*Shigella* causes invasive intestinal infections in humans, ranging from acute watery diarrhea to dysenteric syndrome. Most people recover spontaneously from shigellosis, but antibiotic therapy is recommended for adults and children with bloody diarrhea, patients at risk of complications or to stop transmission in certain outbreak-prone settings^1–3^. The drugs currently used are ciprofloxacin (CIP), ceftriaxone (a third-generation cephalosporin, 3GC), and azithromycin (AZM). There are four serogroups (formerly species) of *Shigella*, with *S. sonnei* the predominant serogroup circulating in industrialized countries and emerging worldwide, even in countries in which other *Shigella* serogroups have traditionally predominated^4^. In the first decade of this century, the annual number of cases of shigellosis was estimated at almost 500,000 in the United States^5^. *S. sonnei* accounted for 80.5% (10,139/11,779) of culture-confirmed *Shigella* infections in the 2016 US national surveillance report^6^. Many multidrugresistant (MDR) strains of *S. sonnei* have been described in recent years. MDR strains were originally linked to travel to Asia or circulation in Orthodox Jewish communities, but are increasingly being reported in the community of men who have sex with men (MSM) around the world^7–13^. In 2018-2019, so-called “extensively drug-resistant” (XDR) *S. sonnei* isolates were reported among MSM in Australia, but these isolates remained susceptible to 3GCs^14^.

We report here a recent dramatic increase in the number of XDR *S. sonnei* isolates (i.e., simultaneously resistant to 3GCs, CIP and AZM) in France. A review of antimicrobial susceptibility data for *S. sonnei* between 2005 and 2021 (based on 7,121 isolates received and confirmed at the French National Reference Center for *E. coli, Shigella* and *Salmonella*, FNRC-ESS) revealed a sharp increase in the percentage of isolates resistant to 3GCs, CIP and AZM (*P* < 0.001) (Fig. 1). A first isolate resistant to 3GCs was identified as early as 2005 (prevalence of 1/342, 0.3%), whereas the first isolates resistant to CIP were identified in 2008 (prevalence of 2/373, 0.5%). Ten isolates resistant to AZM were detected in 2014, the year in which screening for such resistance began (in April) (10/435, 2.3%). In 2021, 29.5% (131/444) of *S. sonnei* isolates were resistant to 3GCs, 42.3% (188/444) were resistant to CIP and 38.7% (172/444) were resistant to AZM (Fig. 1). The first XDR *S. sonnei* isolate was identified in 2015, and an additional 163 XDR isolates have since been obtained (Fig. 1, Tables S1 and S2). All but one of these 164 XDR isolates were collected from mainland France. The percentage of XDR *S. sonnei* isolates increased over the study period (*P* < 0.001), peaking at 22.3% (99/444) in 2021. All but one of these XDR isolates were also resistant to trimethoprim/sulfamethoxazole. They remained susceptible to carbapenems.

**Figure 1.**
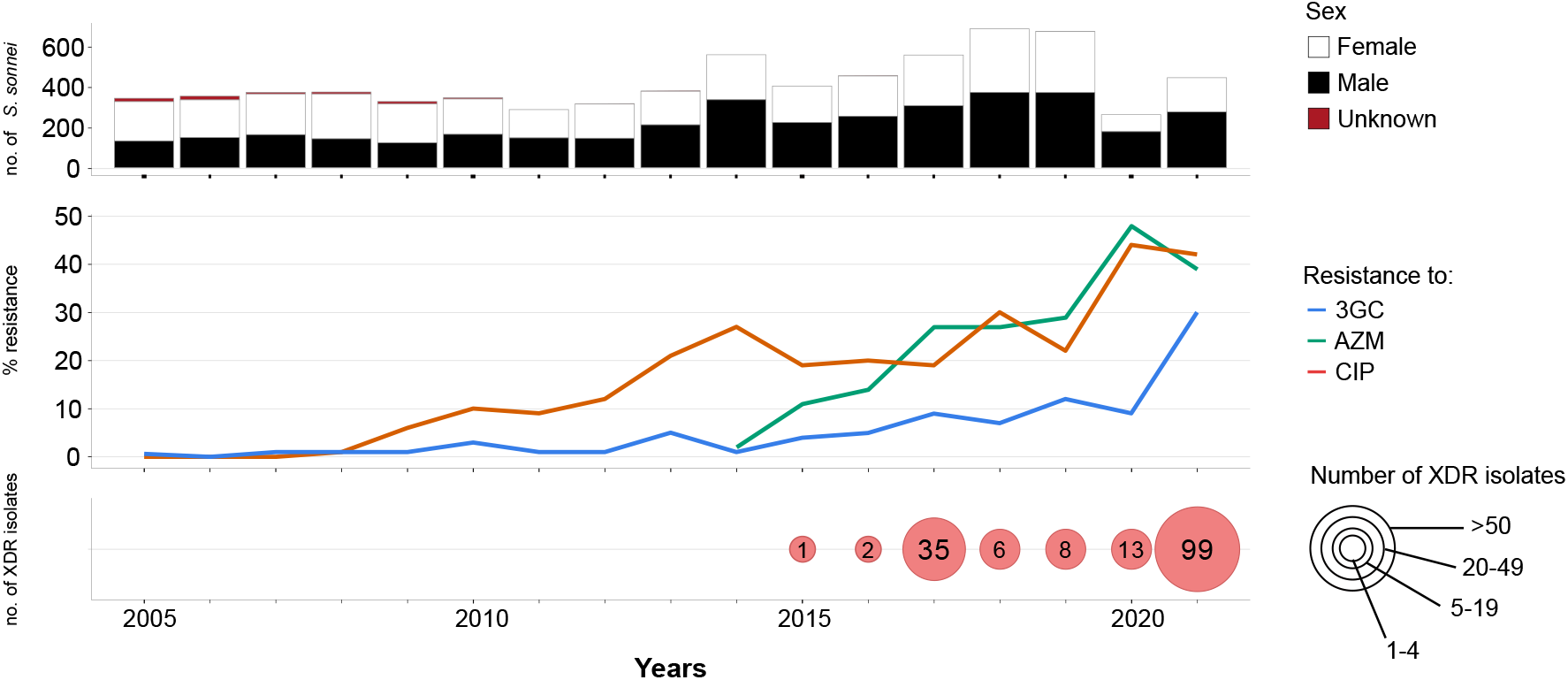
Laboratory surveillance of *Shigella sonnei* infections in France, 2005-2021. The upper panel shows the number of isolates, by patient sex, received and analyzed per year. The middle panel shows the percentage of isolates displaying phenotypic resistance to ciprofloxacin (CIP), azithromycin (AZM) or third-generation cephalosporins (3GCs). The lower panel shows the number of extensively drug-resistant (XDR) *S. sonnei* isolates detected per year. XDR means combined resistance to CIP, AZM and 3GCs.

Previous genomic studies showed that *S. sonnei* forms five different lineages (L1 to L5), one of which, L3 has dominated globally over the last two decades^15,16^. A recently developed genomic tool based on a selection of core-genome single-nucleotide variants (SNVs) can categorize *S. sonnei* into informative phylogenetic genotypes with a universal nomenclature (e.g., genotype 3.6.1.1_CipR, which corresponds to lineage L3, clade 6, subclade 1, subtype 1 with alias name CipR, as used in previous publications for isolates resistant to CIP)^16^. This tool can also identify antimicrobial resistance (AMR) determinants present in *S. sonnei^16^*.

We therefore investigated the phylogenetic relationships and AMR determinants by sequencing the genomes of the 164 French XDR *S. sonnei* isolates. We also included in the phylogenomic analysis 2,976 genomes from isolates and historical strains from the FNRC-ESS collected between 1945 and 2021, to provide a phylogenetic context for the 164 XDR *S. sonnei* isolates (see Methods). A maximum likelihood (ML) phylogenetic tree built from an alignment of 55,099 chromosomal SNVs revealed that the 164 XDR *S. sonnei* isolates were grouped into 13 clusters (X1 to X13) belonging to a single subclade (3.6.1) of lineage L3 (Fig. 2). The genotyping scheme further assigned these isolates to four different, heterogeneously distributed subtypes. Two of these subtypes contain ≤ 5 XDR isolates: 3.6.1_CipR-parent (*n* = 2) and 3.6.1.1_CipR (*n* = 5), and another two each contain more than 30 XDR isolates: 3.6.1.1.1_CipR.SEA (*n* = 36) and 3.6.1.1.2_CipR.MSM5 (*n* = 121) (Fig. 3). For the sake of simplicity, we will use mostly alias names (e.g., CipR-parent, CipR, CipR.SEA, CipR.MSM5) hereafter. The two clusters with the largest numbers of XDR isolates were X4 (*n* = 34) and X10 (*n* = 102) belonging to CipR.SEA and CipR.MSM5, respectively. Some of the XDR *S. sonnei* isolates belonging to CipR-parent, CipR and CipR.SEA were acquired following travel (mostly to South and Southeast Asia) (Fig. 3). For cluster X4 (CipR.SEA), three patients reported travel to Vietnam, including the child who was the index case of an outbreak at an elementary school (91 cases identified) in Southwestern France in March-April 2017, leading to the temporary closure of the school^17^. All but one of the cases contaminated with an isolate from CipR.MSM5 for whom travel information was available reported no travel outside Europe. This widespread genotype was previously reported to be associated with MSM in the UK and Australia^10,16^. Sexual orientation is not recorded on our laboratory-surveillance notification form (see Methods). However, the sex ratio (117 male patients and only four female patients) and age (median age: 34 years; interquartile range 28 – 42 years; age range: 13 to 68 years) of the cases of infection with CipR.MSM5 XDR strains are very suggestive of transmission among MSM (Fig. 3).

**Figure 2.**
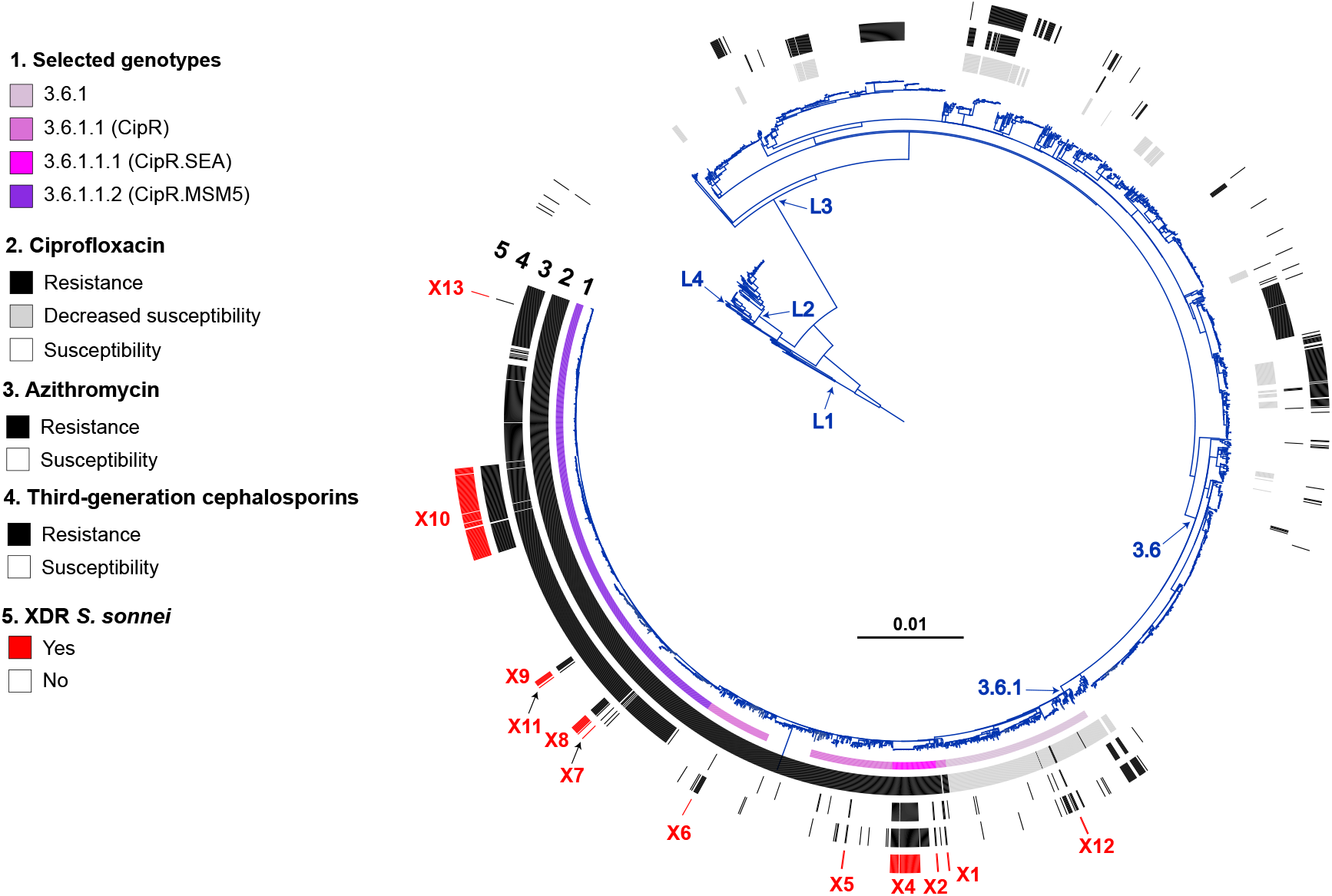
Maximum-likelihood phylogeny of 3,141 *S. sonnei* genomic sequences. The *S. sonnei* reference genome 53G (GenBank accession numbers NC_016822) was also included in addition to the 3,140 genomes from the French National Reference Center for *E. coli, Shigella* and *Salmonella* (Supplementary Table 4). The circular phylogenetic tree (in blue) is rooted on *E. coli* O45:H2 FWSEC0003. *S. sonnei* lineages L1 to L4, clade 3.6 and subclade 3.6.1 are indicated. The rings show the associated information (see key) for each isolate, according to its position in the phylogeny, from the innermost to the outermost, in the following order: (1) selected *S. sonnei* genotypes; (2) antimicrobial susceptibility testing (AST) for ciprofloxacin; (3) AST for azithromycin; (4) AST for third-generation cephalosporins; (5) and the XDR isolates (with the XDR cluster names, X1 to X13, in red). The scale bar indicates the number of substitutions per variable site (SNVs). Detailed information on antimicrobial drug resistance genes and elements are provided in Figs. 4–6. Due to a deliberate strategy to enrich our genomic dataset with azithromycin-resistant, ciprofloxacin-resistant, and third-generation cephalosporin-resistant isolates collected before 2017, the year in which routine genomic surveillance began in France (see Methods), there is an overrepresentation of these resistances in this figure with respect to the global population of 7,121 isolates studied.

**Figure 3.**
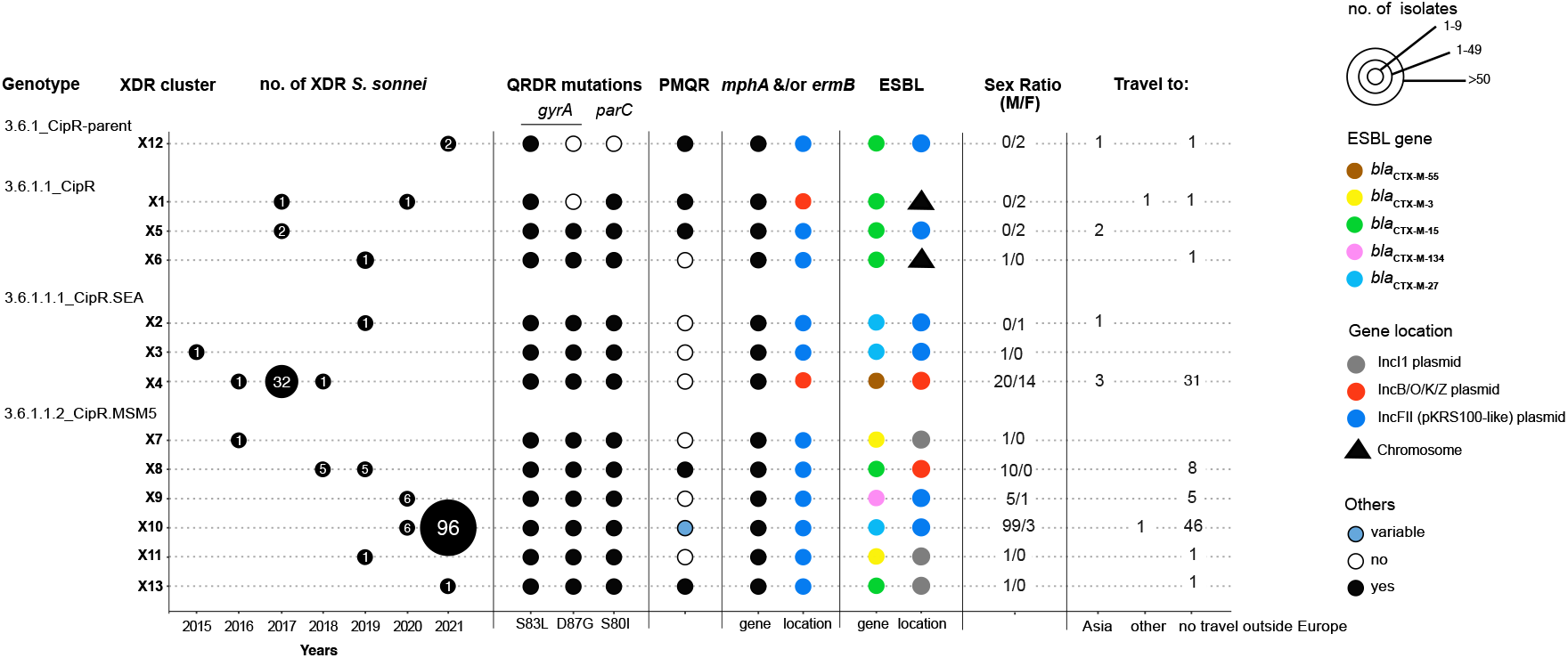
Main characteristics of the 164 XDR *S. sonnei* isolates. The number of XDR isolates, categorized by genotype and XDR genomic cluster, is given by year of isolation. For each XDR genomic cluster, the mechanisms of resistance to CIP (mutation in the quinolone resistance-determining (QRDR) region of *gyrA* and *parC*, presence of a plasmid-mediated quinolone (PMQR) resistance gene), AZM (presence of *mph(A)* and/or *erm(B)* resistance genes) or 3GCs (presence of the extended-spectrum beta-lactamase (ESBL) *bla*_CTX-M_ genes indicated in the key) are indicated. The chromosomal or plasmid location of the AZM and 3GC resistance genes is indicated. For plasmid-borne genes, the type of plasmid is also indicated. Finally, the sex ratio and information about travel (when known) are also indicated for the cases associated with each XDR genomic cluster.

All but four of the XDR *S. sonnei* isolates harbored three mutations in the quinolone resistance-determining regions (QRDR) of the *gyrA* and *parC* genes, conferring a high level of resistance to CIP (MIC > 0.5 mg/L) (Figs. 3 and 4). The other four isolates – from the more ancestral clades, CipR-parent and CipR – harbored only one or two QRDR mutations but also carried a plasmid-borne *qnr* gene conferring a high level of CIP resistance. For better characterization of the mobile (or chromosomally integrated) elements encoding AMR, we also performed long-read sequencing on 16 XDR *S. sonnei* isolates (one to two per XDR cluster). Resistance to AZM was conferred by the *mphA* and/or *ermB* genes, located on two different plasmids. The first was a common plasmid typed as IncFII (PTU-PE) and resembling pKRS100 – previously described in *S. flexneri* 3a, 2a, and *S. sonnei* sublineages associated with the MSM community from Europe and North America – and the second was a less common plasmid typed as IncB/O/K/Z (Figs. 3 and 5, Supplementary Table 3)^10,16,18^. Resistance to 3GCs was conferred by various extended-spectrum beta-lactamase (ESBL) *bla*_CTX-M_ genes (*bla*_CTX-M-3_, *bla*_CTX-M-15_, *bla*_CTX-M-27_, *bla*_CTX-M-55_, and *bla*_CTX-M-134_) located on multiple plasmids of different types (IncFII, including the pKSR100-like plasmid encoding resistance to AZM, IncB/O/K/Z, IncI1) or even integrated into the bacterial chromosome (Figs. 3 and 6, Supplementary Figs. 1-6, Supplementary Table 3, Supplementary Results), suggesting that these antimicrobial drugs have exerted strong selective pressure.

**Figure 4.**
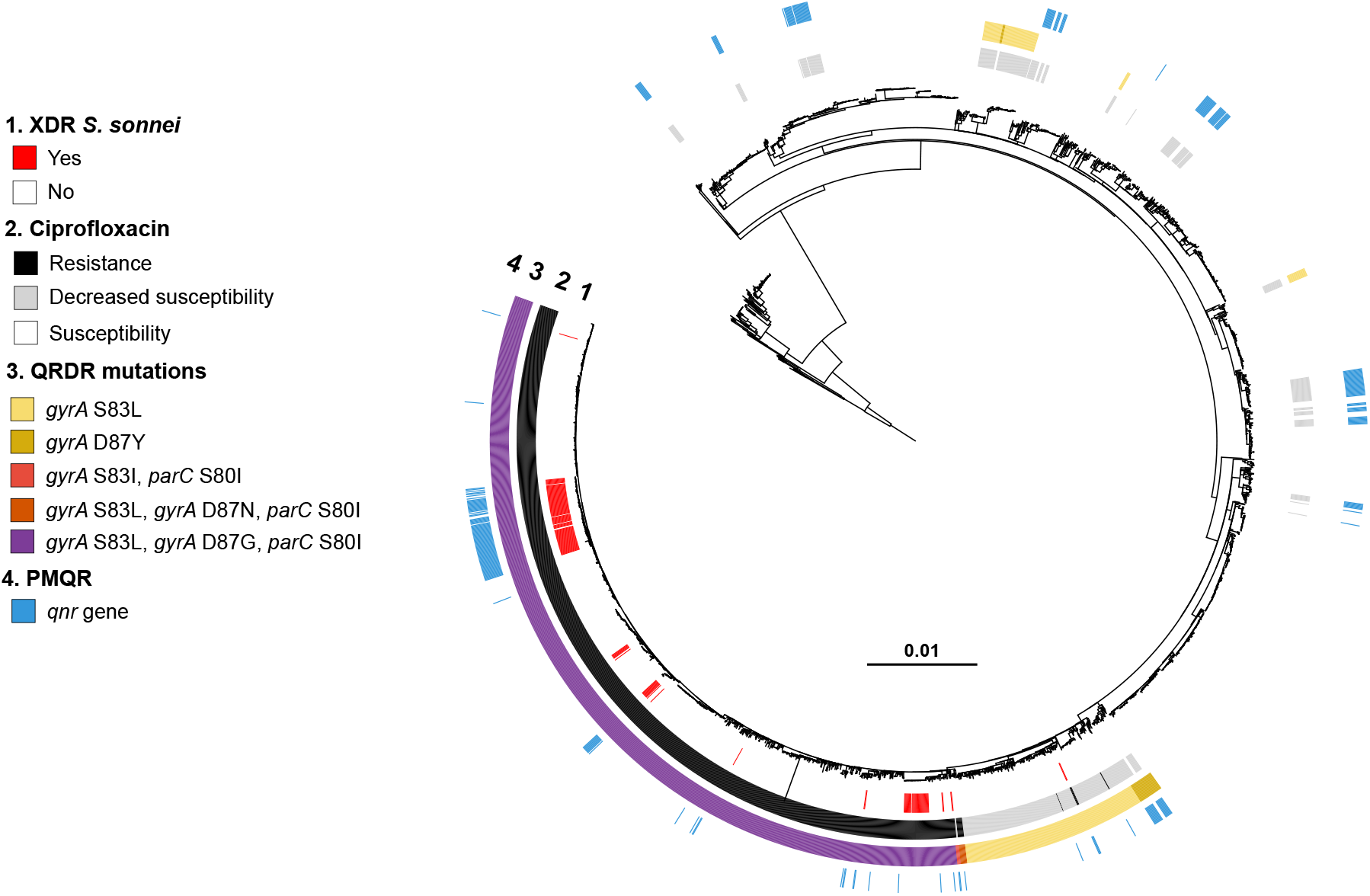
Acquisition of genes encoding resistance to quinolones and fluoroquinolones in our genomic dataset. Maximum-likelihood phylogeny of 3,141 *S. sonnei* genomic sequences as shown in Fig. 2. The rings show the associated information (see key) for each isolate, according to its position in the phylogeny, from the innermost to the outermost, in the following order: (1) the XDR isolates; (2) antimicrobial susceptibility testing for ciprofloxacin (resistance defined as minimum inhibitory concentration [MIC] > 0.5 mg/L; susceptibility as MIC ≤ 0.06 mg/L; decreased susceptibility as MIC between 0.125 and 0.5 mg/L); (3) mutations in the quinolone resistance-determining region (QRDR) of *gyrA* and *parC*; (4) presence of plasmid-mediated quinolone resistance (PMQR) genes of the *qnr* family.

**Figure 5.**
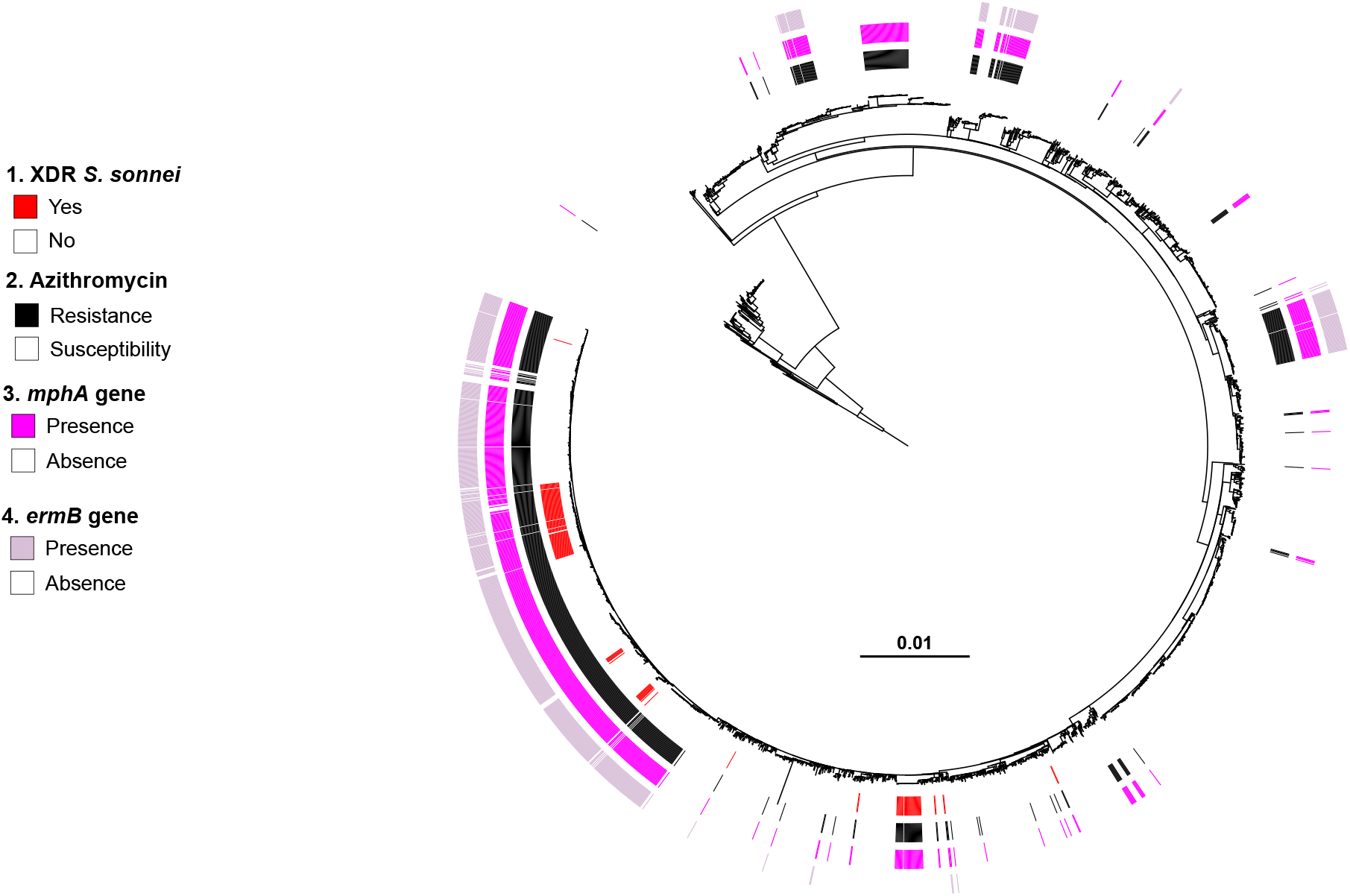
Acquisition of genes encoding resistance to azithromycin in our genomic dataset. Maximum-likelihood phylogeny of 3,141 *S. sonnei* genomic sequences as shown in Fig. 2. The rings show the associated information (see key) for each isolate, according to its position in the phylogeny, from the innermost to the outermost, in the following order: (1) the XDR isolates; (2) antimicrobial susceptibility testing for azithromycin (resistance defined as minimum inhibitory concentration [MIC] ≥ 32 mg/L; susceptibility as MIC ≤ 16 mg/L); (3) presence of the *mph(A)* gene; (4) presence of the *erm(B)* gene.

**Figure 6.**
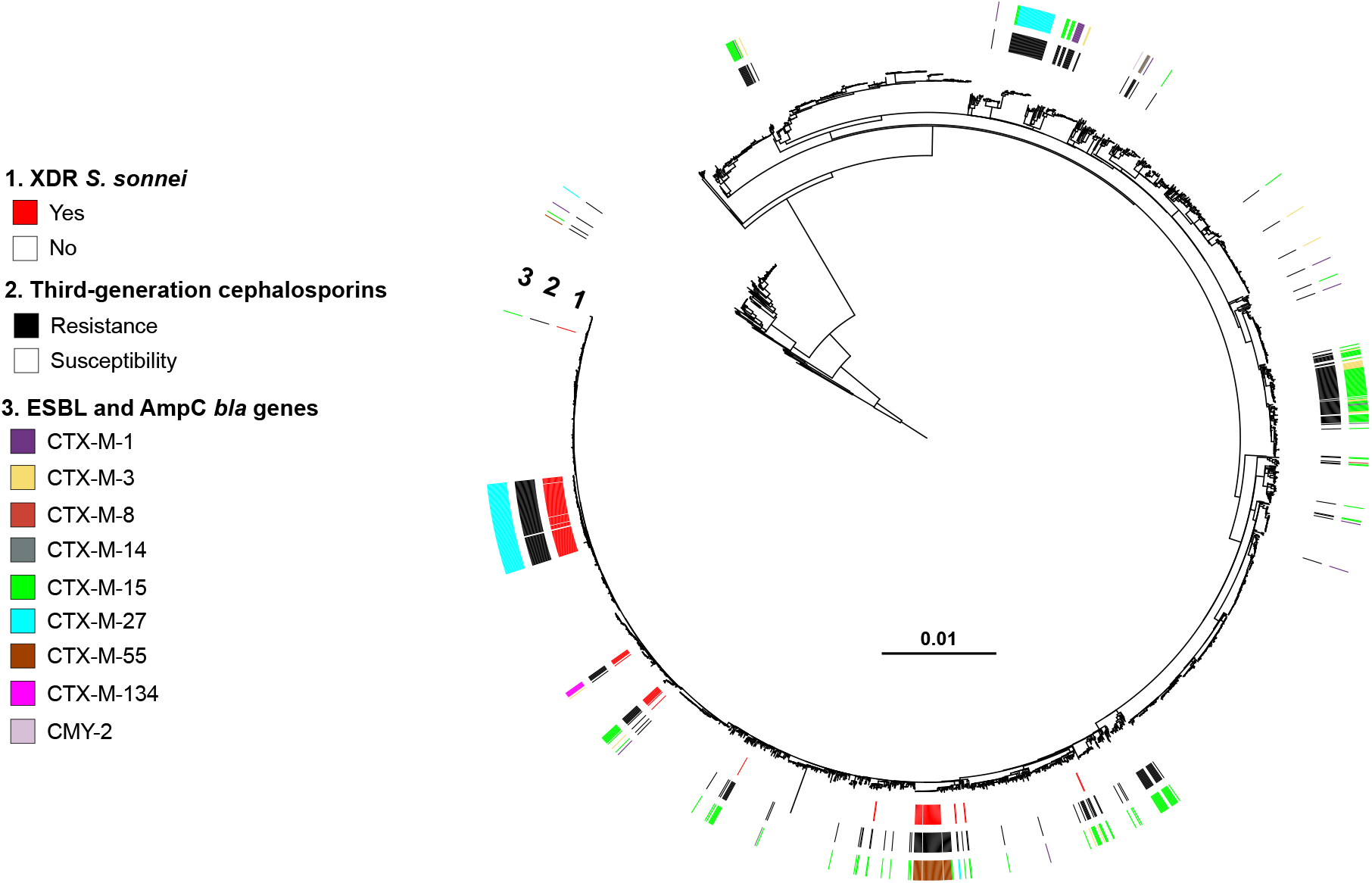
Acquisition of selected genes encoding resistance to third-generation cephalosporins in our genomic dataset. Maximum-likelihood phylogeny of 3,141 *S. sonnei* genomic sequences as shown in Fig. 2. The rings show the associated information (see key) for each isolate, according to its position in the phylogeny, from the innermost to the outermost in the following order: (1) the XDR isolates; (2) antimicrobial susceptibility testing for third-generation cephalosporins (resistance defined as minimum inhibitory concentration [MIC] of ceftriaxone ≥ 4 mg/L; susceptibility as MIC ≤ 1mg/L); (3) presence of *bla* genes encoding extended-spectrum beta-lactamases (ESBLs) or cephamycinases.

Between 2016 and 2019, only 0.8% (36/4,222) of the 4,222 *S. sonnei* genomic sequences obtained in the framework of public health surveillance in England, Australia and the US were inferred to be XDR^16^. These 36 XDR isolates from England (*n* = 20), Australia (*n* = 13) and the US (*n* = 3) also belonged to the CipR, CipR.SEA, and CipR.MSM5 genotypes^16^. Our findings indicate that XDR *S. sonnei* strains are no longer extremely rare, as they accounted for 22.3% (99/444) of all *S. sonnei* isolates in France in 2021. We found these XDR isolates in various epidemiological contexts, initially related to international travel to Asia (South or Southeast Asia), but they have since been identified in various networks, such as a school, or, more frequently, the MSM community. The alarming situation already observed in France is likely to occur in other high-income countries, given the high degree of connectivity between members of the MSM community across the world, and the fact that the CipR.MSM5 genotype is already widespread^10,16,18^. Our largest XDR cluster, X10 – belonging to CipR.MSM5 and containing the ESBL *bla*_CTX-M-27_ gene on a pKSR100-like plasmid – has already been found among MSM in nine other European countries, with a total of 208 isolates as of February 23, 2022 (ref. ^19^). France had the largest number of isolates (*n* = 106) and the earliest isolate, dating from September 2020. A descriptive epidemiological study of this outbreak (72 cases) was recently performed in the UK; 21 of the 27 cases interviewed (78%) were HIV-negative MSM and users of HIV pre-exposure prophylaxis (PrEP)^20^. These individuals also reported having high-risk sex in England or Europe during the incubation period. Due to the few therapeutic options left, the recommended oral antimicrobial therapy – when needed – was either pivmecillinam or fosfomycin (chloramphenicol not being readily accessible in the UK)^20^. For hospitalized or severe cases, carbapenems (colistin in case of allergy to beta-lactams) were used for three to five days, followed by an oral step-down treatment^20^.

In summary, our analysis of antimicrobial susceptibility testing data at national level over a long time period (from 2005 to 2021 with over 7,000 *S. sonnei* isolates) identified the first XDR *S. sonnei* isolate in France in 2015. The number of XDR isolates has since increased considerably. Our genomic analysis showed that all our 164 XDR isolates were descended from a successful sublineage (3.6.1_CipR) known to have acquired full resistance to CIP in South Asia by around 2007 (refs. ^7,16^). However, the XDR phenotype was not the result of a single clonal expansion of this sublineage, but arose independently on multiple occasions, through different plasmids and resistance genes. An effective national laboratory-based surveillance of *Shigella* infections, including antimicrobial susceptibility data, is therefore essential for informed decision-making and appropriate public health action to tackle the spread of these XDR *S. sonnei* strains circulating in different transmission networks. Genomic surveillance with this new genotyping scheme with a common nomenclature will facilitate the detection and tracking at global scale of these alarming XDR *S. sonnei* strains.

## METHODS

### Bacterial isolates

The French national surveillance program for *Shigella* infections is based on a voluntary laboratory-based network consisting of approximately 1,000 clinical laboratories located in mainland France and its overseas territories in South America (French Guiana), the Caribbean (Martinique, Guadeloupe) and the Indian Ocean (La Réunion, Mayotte), which send about 1,000-1,200 *Shigella* spp. isolates to the FNRC-ESS each year (only 600 in 2020, due to the COVID pandemic). The *Shigella* isolates are sent to the FNRC-ESS with a notification form that includes basic data: patient name, date of birth, sex, postcode, stools or other types of sample, isolation date, clinical symptoms and their onset, illness or asymptomatic carriage, (if illness, types of symptoms), information about international travel (if yes, date and country), sporadic or outbreak isolates (if clustered, hospital, school, household, nursery, etc.).

We studied all 7,121 *S. sonnei* isolates (one per patient) received by the French National Reference Center for *E. coli, Shigella* and *Salmonella* (FNRC-ESS) between 2005 and 2021, in the framework of the French national surveillance program for *Shigella* infections. It has been estimated that this surveillance system detects 50-60% of laboratory-confirmed *Shigella* infections in France^21^.The 7,121 *S. sonnei* human isolates studied consisted of 6,700 (94.1%) from mainland France and 421 (5.9%) from French overseas territories. From January 2005 to September 2021 (when phenotypic typing was definitively replaced by genomic surveillance), all these isolates were thoroughly characterized with a panel of biochemical tests and serotyped with slide agglutination assays according to standard protocols, as previously described^22^.

### Antimicrobial drug susceptibility testing

Antimicrobial drug susceptibility testing was performed on all *S. sonnei* isolates, at the time of reception. Isolates were first tested with the disk diffusion (DD) method on Mueller-Hinton agar (Bio-Rad, Marnes-la-Coquette, France) according to the 2005-2015 guidelines of the antibiogram committee of the French Society for Microbiology (CA-SFM), in accordance with the recommendations of the European Committee on Antimicrobial Susceptibility Testing (EUCAST) (https://www.sfm-microbiologie.org/casfm/). The following disks (Bio-Rad, Marnes-La-Coquette, France) were used for the DD method: amoxicillin (AMX, 10 μg) or ampicillin (AMP, 10 μg), cefotaxime (CTX, 5 μg), ceftazidime (CAZ, 30 μg before 2015, 10 μg from 2015), ertapenem (ETP, 10 μg), chloramphenicol (CHL, 30 μg), sulfonamides (SMX, 200 μg), trimethoprim (TMP, 5 μg), trimethoprim-sulfamethoxazole (SXT, 1.25 μg/23.75 μg), streptomycin (STR, 10 μg), amikacin (AKN, 30 μg), gentamicin (GEN, 10 μg), tetracycline (TET, 30 μg), nalidixic acid (NAL, 30 μg), ofloxacin (OFX, 5 μg) or pefloxacin (PEF, 5 μg), ciprofloxacin (CIP, 5 μg), and azithromycin (AZM, 15 μg). Susceptibility to AZM was tested systematically only from April 2014. Resistance to 3GCs was defined as resistance to ceftazidime, cefotaxime or ceftriaxone. For isolates resistant to NAL, CIP, AZM or 3GCs, we confirmed the DD data by determining the minimum inhibitory concentrations (MICs) of these drugs with Etest strips (AB Biodisk, Solna, Sweden; bioMérieux, Marcy L’Etoile, France). The Clinical and Laboratory Standards Institute (CLSI) criteria were then used for the final interpretation^23^. As a means of distinguishing *Shigella* isolates susceptible to ciprofloxacin (minimum inhibitory concentration [MIC] ≤ 0.25 mg/L) that are wild-type (WT) from those that are non-WT, we defined two categories based on the epidemiological cutoffs used by the CLSI for *Salmonella* spp.: decreased susceptibility to ciprofloxacin (MIC between 0.125 and 0.5 mg/L) and true susceptibility to ciprofloxacin (MIC ≤ 0.06 mg/L)^23^.

### Whole-genome sequencing

In total, 3,109 *S. sonnei* isolates from the 7,121 collected between 2005 and 2021 were sequenced, including all 164 XDR isolates. Between 2017 (the year in which genomic surveillance began in France) and 2021, 2,618 clinical isolates were received and sequenced at the FNRC-ESS. We included in this study the 2,558 genomes (97.7%) that passed the EnteroBase quality control criteria (https://enterobase.warwick.ac.uk/species/index/ecoli). We also sequenced a selection of 551/4,503 (12.2%) of the 2005-2016 isolates. This selection contained, in particular, 96.7% (89/92) of all isolates resistant to 3GCs, 56.3% (294/522) of all isolates resistant to CIP, and 96.7% (116/120) of the isolates resistant to AZM detected from April 2014 to December 2016. Total DNA was extracted with the MagNA Pure DNA isolation kit (Roche Molecular Systems, Indianapolis, IN, USA), in accordance with the manufacturer’s recommendations. Whole-genome sequencing was performed as part of routine procedures at the FNRC-ESS, and at the Mutualized Platform for Microbiology (P2M) at Institut Pasteur, Paris. The libraries were prepared with the Nextera XT kit (Illumina, San Diego, CA, USA) and sequencing was performed with the NextSeq 500 system (Illumina) generating 150 bp paired-end reads. All reads were filtered with FqCleanER version 21.06 (https://gitlab.pasteur.fr/GIPhy/fqCleanER) with options -q 15 -l 50 to eliminate adaptor sequences and discard low-quality reads with phred scores below 15 and a length of less than 50 bp^24^.

Assemblies were generated with SPAdes version 3.15.2 (ref. ^25^).

Short-read sequence data were submitted to EnteroBase (https://enterobase.warwick.ac.uk/) and the European Nucleotide Archive (ENA, https://www.ebi.ac.uk/ena/) under study number PRJEB44801 (Supplementary Table S2).

### Genotyping

All the genomes studied were genotyped with the hierarchical SNV-based genotyping scheme for *S. sonnei* described by Hawkey *et al*.^16^ and implemented in Mykrobe software version 0.9.0 (https://github.com/katholt/sonneityping)^26^.

### Phylogenomic analysis

We included in the phylogenomic analysis 3,140 *S. sonnei* genomic sequences from isolates and historical strains of the FNRC-ESS (Supplementary Table S4) originating from 3,109 *S. sonnei* isolates collected between 2005 and 2021 and 31 isolated before 2005, to enrich the dataset with rare lineages of *S. sonnei* (L1, L2 and L4)^8,15^.

The paired-end reads were mapped onto the reference genome of *S. sonnei* 53G (GenBank accession numbers NC_016822)^15^ with Snippy version 4.6.0/BWA-MEM version 0.7.17 (https://github.com/tseemann/snippy). SNVs were called with Snippy version 4.6.0/Freebayes version 1.3.2 (https://github.com/tseemann/snippy) under the following constraints: mapping quality of 60, a minimum base quality of 13, a minimum read coverage of 4, and a 75% read concordance at a locus for a variant to be reported. An alignment of core genome SNVs was produced in Snippy version 4.6.0 for phylogenetic inference.

Repetitive regions (i.e., insertion sequences, tRNAs) in the alignment were masked (https://doi.org/10.26180/5f1a443b19b2f)^16^. Putative recombinogenic regions were detected and masked with Gubbins version 3.2.0 (ref. ^27^). A maximum likelihood (ML) phylogenetic tree was built from an alignment of 55,099 chromosomal SNVs, with RAxML version 8.2.12, under the GTR model, with 200 bootstrap values^28^. The final tree was rooted on the *E. coli* O45:H2 FWSEC0003 genome (GenBank accession no. CP031916) genome and visualized with iTOL version 6 (https://itol.embl.de)^29^.

### Resistance gene analysis

The presence and type of acquired antibiotic resistance genes (ARGs) were determined with ResFinder version 4.0.1 (https://cge.cbs.dtu.dk/services/ResFinder/)^30^, Sonneityping/Mykrobe version 0.9.0 (https://github.com/katholt/sonneityping)^16,26^. The presence of mutations in genes encoding resistance to quinolones (*gyrA, parC*) was also investigated by analyzing the sequences assembled *de novo* with BLAST version 2.2.26.

### Plasmid sequencing

Sixteen XDR *S. sonnei* isolates (one to two per XDR cluster) were selected (Supplementary Table 3) and sequenced with a Nanopore MinION sequencer (Oxford Nanopore Technologies). Genomic DNA was prepared as follows: the isolates were cultured overnight at 37 °C in alkaline nutrient agar (20 g casein meat peptone E2 from Organotechnie; 5 g sodium chloride from Sigma; 15 g Bacto agar from Difco; distilled water to 1 L; adjusted to pH 8.4; autoclaved at 121°C for 15 min). A few isolated colonies from the overnight culture were used to inoculate 20 mL of brain-heart infusion (BHI) broth, and were cultured until a final OD_600_ of 0.8 was reached at 37°C with shaking (200 rpm —Thermo Fisher Scientific MaxQ 6800). The bacterial cells were harvested by centrifugation and DNA was extracted with one of the two following methods. The first method corresponded to the protocol described by von Mentzer *et al*.^31^, except that MaXtract High-Density columns (Qiagen) were used (instead of phase-lock tubes) and the DNA was resuspended in molecular biology-grade water (instead of 10 mM Tris pH 8.0). In the second method, we used Genomic-tip 100/G columns (Qiagen) according to the manufacturer’s protocol. The library was prepared according to the instructions of the “Native barcoding genomic DNA (with EXP-NBD104, EXP-NBD114, and SQK-LSK109)” procedure provided by Oxford Nanopore Technology. Sequencing was then performed on a MinION Mk1C apparatus (Oxford Nanopore Technologies). The genomic sequences of the isolates were assembled from long and short reads, with a hybrid approach and UniCycler version 0.4.8 (ref. ^32^). A polishing step was performed with Pilon version 1.23 (ref. ^33^) to generate a high-quality sequence composed of chromosomal and plasmid sequences. The plasmids were then annotated with Prokka version 1.14.5 (https://github.com/tseemann/prokka)^34^ and corrected manually. Plasmids were aligned and visualized with BRIG version 0.95 (http://sourceforge.net/projects/brig)^35^.

### Plasmid typing

The plasmids were typed with PlasmidFinder version 2.1.1. (https://cge.cbs.dtu.dk/services/PlasmidFinder/)^36^, pMLST version 1.2 (https://cge.cbs.dtu.dk/services/pMLST/)^36^, and COPLA version 1.0 (https://castillo.dicom.unican.es/copla/)^37^ on SPAdes assemblies.

## Supporting information

Supplementary Results, Supplementary Figs. 1-7, Supplementary Tables 1-4, Supplementary References

## Statistical analysis

Chi-squared tests for trends were used to analyze the proportion of bacterial strains resistant to antimicrobial drugs by year.

## Data collection

The data were entered into an Excel (Microsoft) version 15.41 spreadsheet.

## Data availability statement

Short-read sequence data were submitted to EnteroBase (https://enterobase.warwick.ac.uk/) and to the European Nucleotide Archive (ENA, https://www.ebi.ac.uk/ena/) under study number PRJEB44801. All the accession numbers of the genomes produced and used in this study are listed in Supplementary Tables 2 and 4). The plasmid sequences obtained were deposited in GenBank under accession numbers OP038267-OP038301 and OP038303 (Supplementary Table 3).

## Code availability statement

No custom computer code nor custom algorithm was used in this study.

## ACKNOWLEDGMENTS

This study was supported by the *Institut Pasteur, Santé publique France*, and the French government’s *Investissement d’Avenir* program, *Laboratoire d’Excellence* ‘Integrative Biology of Emerging Infectious Diseases’ (grant number ANR-10-LABX-62-IBEID). We also thank all the corresponding laboratories of the French National Reference Center for

*Escherichia coli*, *Shigella*, and *Salmonella*.

The funders had no role in study design, data collection and analysis, decision to publish, or preparation of the manuscript.

## AUTHOR CONTRIBUTIONS STATEMENT

S.L. and F.-X.W. conceptualized and designed the study. S.L., E.N., I.Y. and F.-X.W. did the genomic analyses. S.L., E.N., I.Y., L.F., and F.-X.W. contributed to data interpretation and visualization. C.R., I.C., M.L.C. and E.N. performed the laboratory experiments. S.L. and S.F. were involved in sample collection and metadata curation. S. F. and A.F. performed statistical analyses. F.-X.W. supervised the project. F.-X.W. and A.F. were responsible for funding acquisition. F.-X.W. drafted the article. S.L., E.N., S.F., L.F., I.Y., L.F., M.P.G. and A.F. critically reviewed the draft. All the authors read and approved the final manuscript. S.L., E.N. and F.-X.W. accessed and verified the underlying data.

## COMPETING INTERESTS STATEMENT

The authors have no competing financial interests to declare.

Correspondence and requests for materials should be addressed to F.-X.W.

