## Supplementary Results, Supplementary Figs. 1-7, Supplementary Tables 1-4, Supplementary References for "Rapid emergence of extensively drug-resistant *Shigella sonnei* in France"

### Plasmids encoding antimicrobial drug resistance

Plasmid analysis was performed on the 16 *S. sonnei* (one to two isolates per XDR genomic cluster) isolates sequenced with an Oxford Nanopore MinION sequencer. The XDR *S. sonnei* isolates contained one to three plasmids carrying antimicrobial drug resistance genes (Supplementary Table 3). Plasmid sizes ranged from 8,379 kb to 106,936 kb. A small ~ 8 kb (either non-typable or typed as PTU-E63) plasmid encoding resistance to streptomycin (*strA* and *strB* genes), sulfonamides (*sul2*) and tetracyclines (*tet(A)*) was found in 12 of 16 XDR isolates from all four genotypes. This plasmid was highly similar to pMHMC-012 (GenBank accession no. CP053763) (Supplementary Fig. 7) from a *S. sonnei* isolate collected from an MSM patient in Boston, USA in 2017 (ref. [1]). Most of the other AMR plasmids belonged to IncFII (PTU-FE) and were related to pKSR100, a plasmid initially from a *S. flexneri* 3a isolate collected in Canada in 2013 but subsequently found in Europe, North America and Australasia in MSM-associated *S. sonnei* and *S. flexneri* sublineages[2-4].

Highly similar ESBL plasmids (98-99% nucleotide identity) were found in different XDR clusters. For example, an IncI1 (PTU-I1) plasmid carrying $bla_{\text{CTX-M-3}}$ was identified in clusters X7 and X11 (same genotype, 3.6.1.1.2_CipR.MSM5) and an IncFII (PTU-FE) plasmid carrying $bla_{\text{CTX-M-27}}$ in clusters X2, X3 and X10 (two different genotypes, 3.6.1.1.1_CipR.SEA and 3.6.1.1.2_CipR.MSM5) (Extended Data Fig. 1, Supplementary Table 2, Supplementary Figs. 4 and 6).

Some of our ESBL plasmids were also similar to certain plasmids described in previous studies. Hence, the $bla_{\text{CTX-M-27}}$-carrying p202008564-6 plasmid (cluster X10, genotype 3.6.1.1.2_CipR.MSM5) displayed 99.6% nucleotide identity to p893916, from a *S. sonnei* isolate collected in London, UK, in 2020 (Supplementary Fig. 4)[5]. The second form of the ESBL plasmid found in cluster X10 isolates – and represented by p202008118-4 – probably resulted from an insertion sequence (IS)-driven deletion or acquisition of the azithromycin-resistance gene, *erm(B)* (Supplementary Fig. 4). The *bla*$_{CTX-M-134}$-carrying p202000562-4 plasmid (cluster X9, genotype 3.6.1.1.2_CipR.MSM5) displayed 90.3% nucleotide identity to p3123885, from a *S. sonnei* isolate acquired in Israel in 2019 (Supplementary Fig. 5)[6]. Finally, the *bla*$_{CTX-M-3}$-carrying p201908234-4 plasmid (cluster X11, genotype 3.6.1.1.2_CipR.MSM5) displayed 99.7% nucleotide identity to p711-69, from a *S. sonnei* isolate acquired in Turkey in 2019 (Supplementary Fig. 6)[6].

In two *S. sonnei* isolates, 201701093 (cluster X1, genotype 3.6.1.1_CipR) and 201908033 (cluster X6, also 3.6.1.1_CipR), the *bla*$_{CTX-M-15}$ ESBL gene was not located on a plasmid but on the bacterial chromosome (Extended Data Fig. 1, Supplementary Table 3). In isolate 201908033, the ESBL gene was part of a ~ 42 kb genomic island described in Supplementary Fig. 1, whereas in isolate 201701093, the ESBL gene was integrated into a ~ 57 kb prophage sequence absent from other XDR genomes (Supplementary Fig. 2).

In the XDR genomic cluster X10, two isolates, 202110142 (accession no. ERR9940949) and 202111148 (ERR9941000) were phenotypically similar to other X10 isolates, except that they were susceptible to 3GCs (and were therefore not classified as XDR). However, they had the same antimicrobial drug resistance gene content (including *bla*$_{CTX-M-27}$) as the other X10 XDR isolates. A careful inspection of the assemblies identified an IS (IS*26*) in both isolates, integrated at the same position of the *bla*$_{CTX-M-27}$ promotor and probably leading to a non-functional ESBL gene.

**Supplementary Figure 1.** Representation of the genomic island containing the *bla*$_{CTX-M-15}$ gene in the XDR *S. sonnei* isolate 201908033

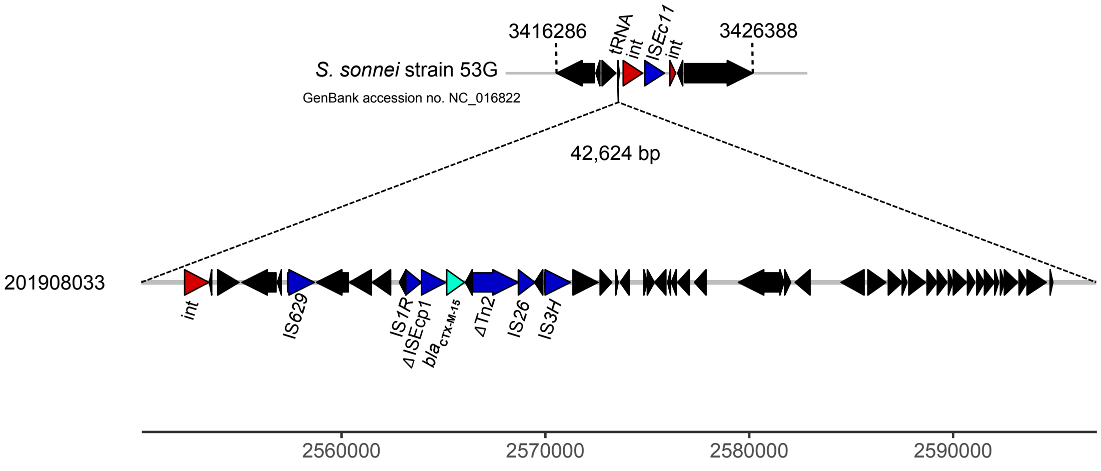

**Supplementary Figure 2.** Representation of the prophage containing the *bla*$_{CTX-M-15}$ gene in the XDR *S. sonnei* isolate 201701093

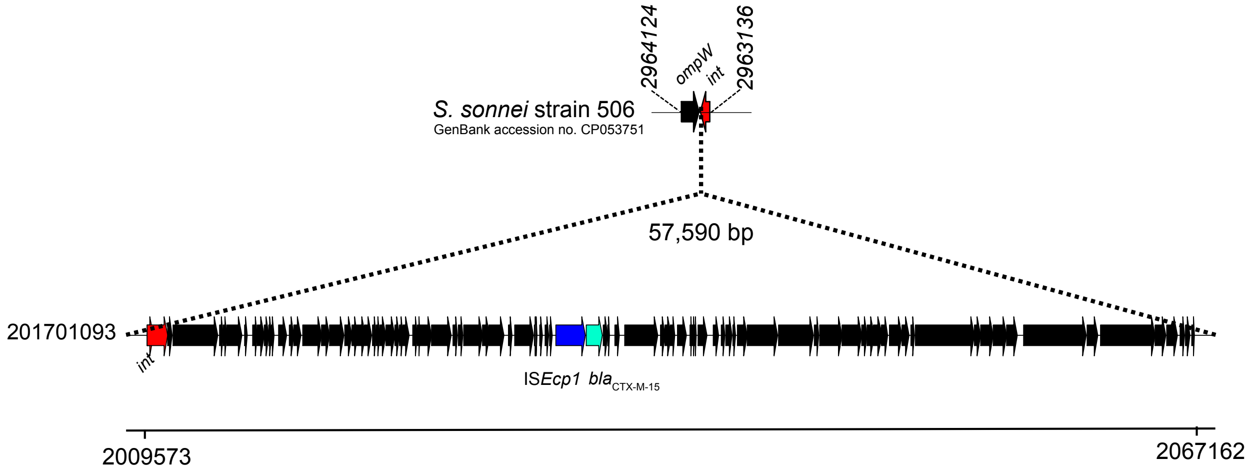

**Supplementary Figure 3.** Circular map and comparative analysis of IncF plasmids carrying the ESBL *bla*$_{CTX-M-15}$ gene

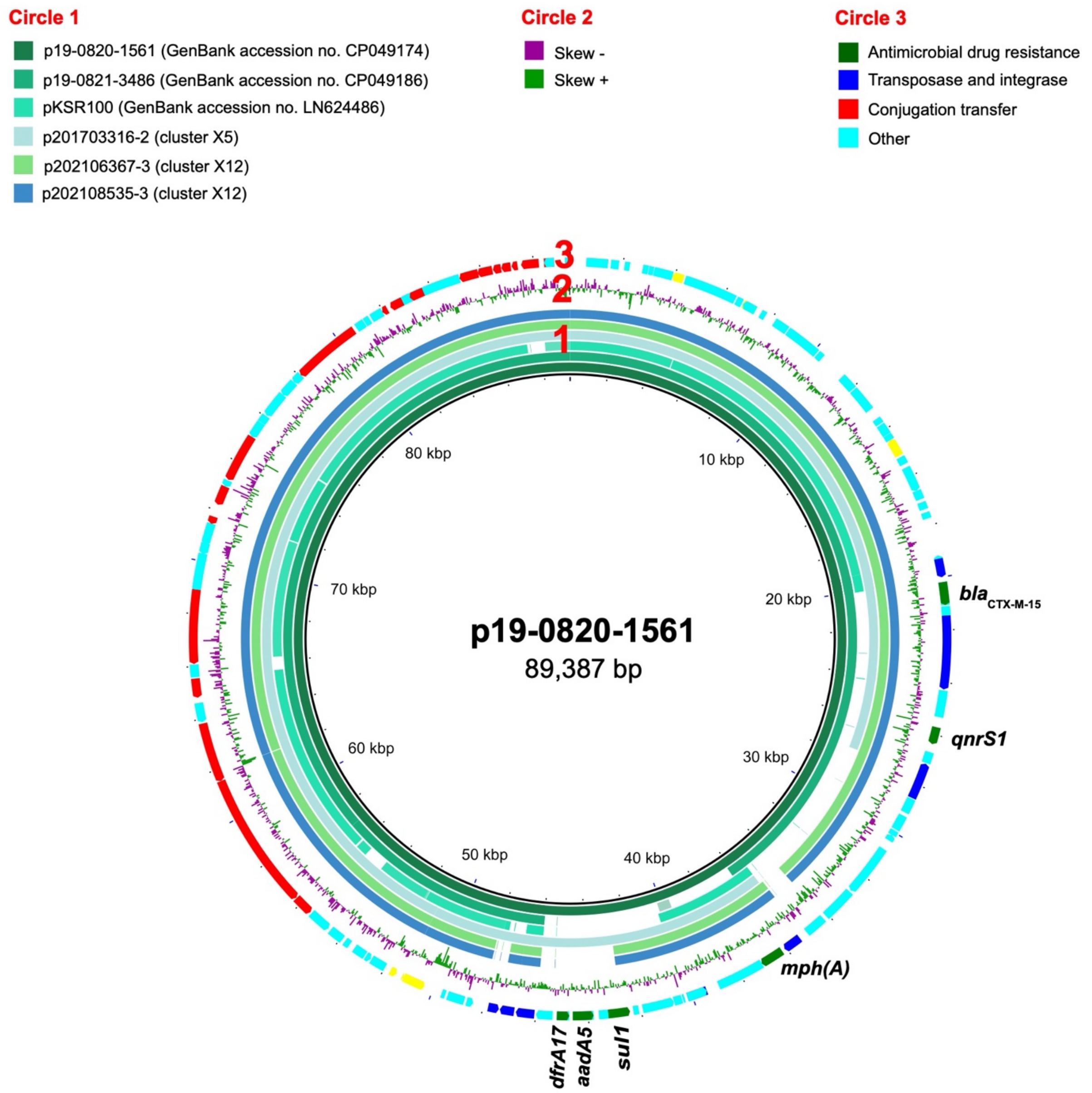


Circles from inside to outside indicate (1) the nucleotide position of p19-0820-1561, a plasmid from a *S. sonnei* isolate acquired in Nepal[6], and regions of p19-0820-1561 displaying high levels of sequence identity to plasmids p19-0821-3486 (from a *S. sonnei* isolate acquired in Egypt)[6], pKSR100 (from a *S. flexneri* 3a isolate, SF7955, collected in Canada, in 2013)[3], p201703316-2 (XDR genomic cluster X5, our study), p202106367-3 (cluster X12, our study), and p202108535-3 (cluster X12, our study), (2) a G+C content map of p19-0820-1561, and (3) coding sequences (CDS) colored according to their functions. The antimicrobial drug resistance genes are indicated.

**Supplementary Figure 4.** Circular map and comparative analysis of IncF plasmids carrying the ESBL *bla*$_{CTX-M-27}$ gene

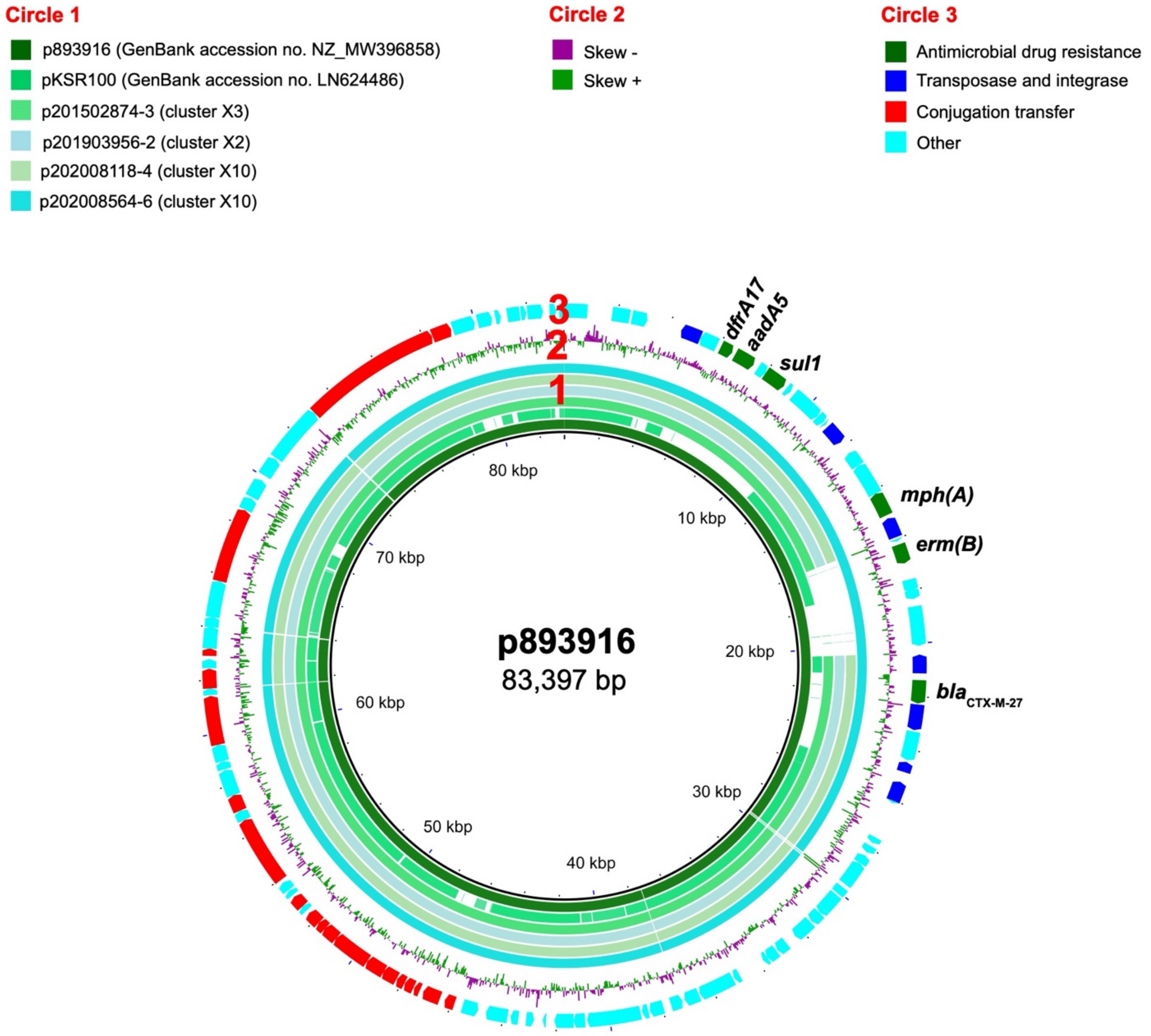


Circles from inside to outside indicate (1) the nucleotide position of p893916, a plasmid from a *S. sonnei* isolate collected in the UK in 2020 (ref. [5]), and regions of p893916 that displaying high levels of sequence identity to plasmids pKSR100 (from a *S. flexneri* 3a isolate, SF7955, collected in Canada in 2013)[3], p201502874-3 (XDR genomic cluster X3, our study), p201903956-2 (cluster X2, our study), p202008118-4 (cluster X10, our study), and p202008564-6 (cluster X10, our study), (2) a G+C content map of p893916, and (3) coding sequences (CDS) colored according to their functions. The antimicrobial drug resistance genes are indicated.

**Supplementary Figure 5.** Circular map and comparative analysis of IncF plasmids carrying the ESBL *bla*$_{CTX-M-134}$ gene

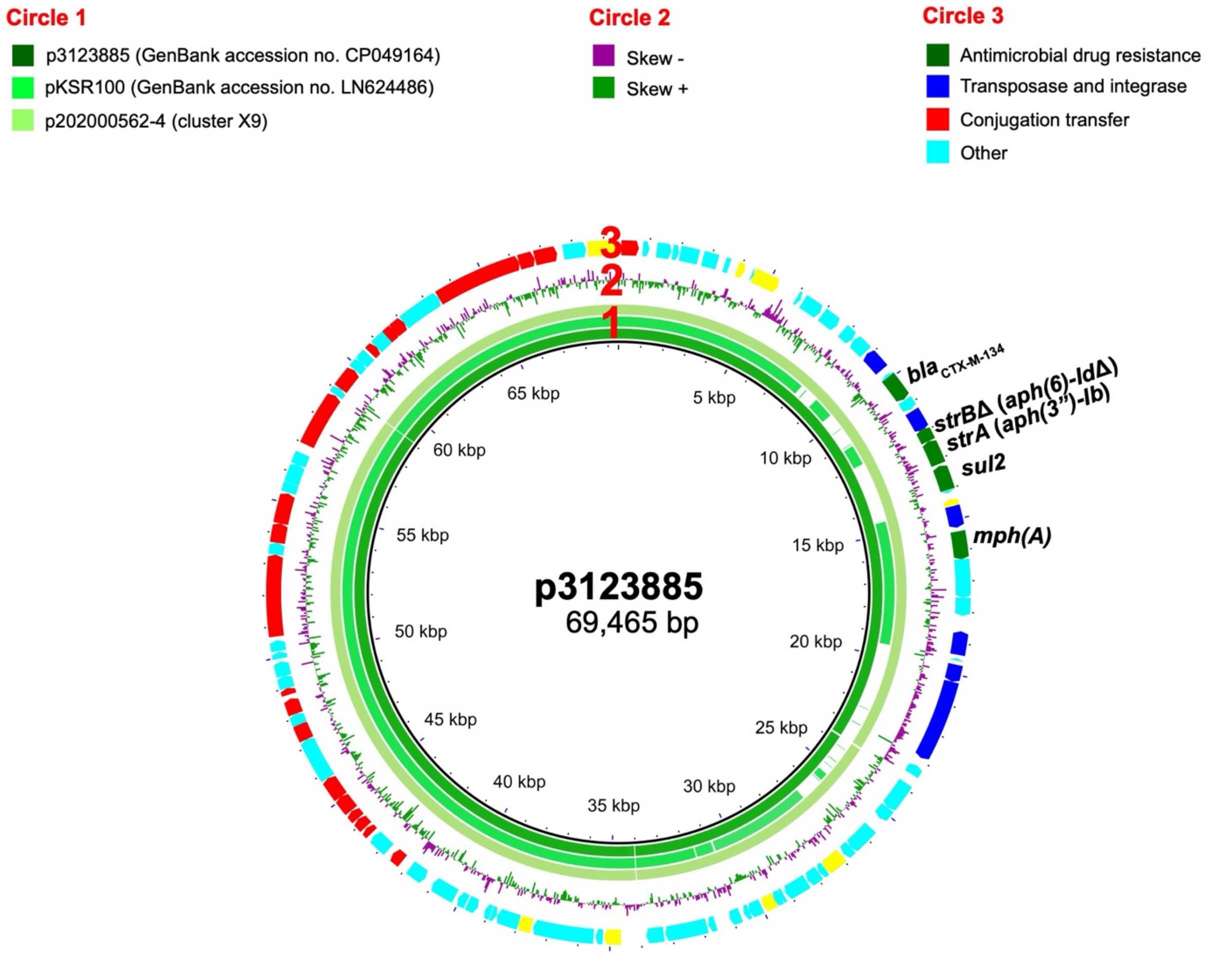


Circles from inside to outside indicate (1) the nucleotide position of p3123885, a plasmid from a *S. sonnei* isolate acquired in Israel in 2019 (ref. [6]), and regions of p3123885 displaying high levels of sequence identity to pKSR100 (from a *S. flexneri* 3a isolate, SF7955, collected in Canada in 2013)[3], and p202000562-4 (XDR genomic cluster X9, our study), (2) a G+C content map of p3123885, and (3) coding sequences (CDS) colored according to their functions. The antimicrobial drug resistance genes are indicated (most of the *strB* gene is deleted).

**Supplementary Figure 6.** Circular map and comparative analysis of IncI1 plasmids carrying the ESBL *bla*$_{CTX\text{-}M\text{-}3}$ gene

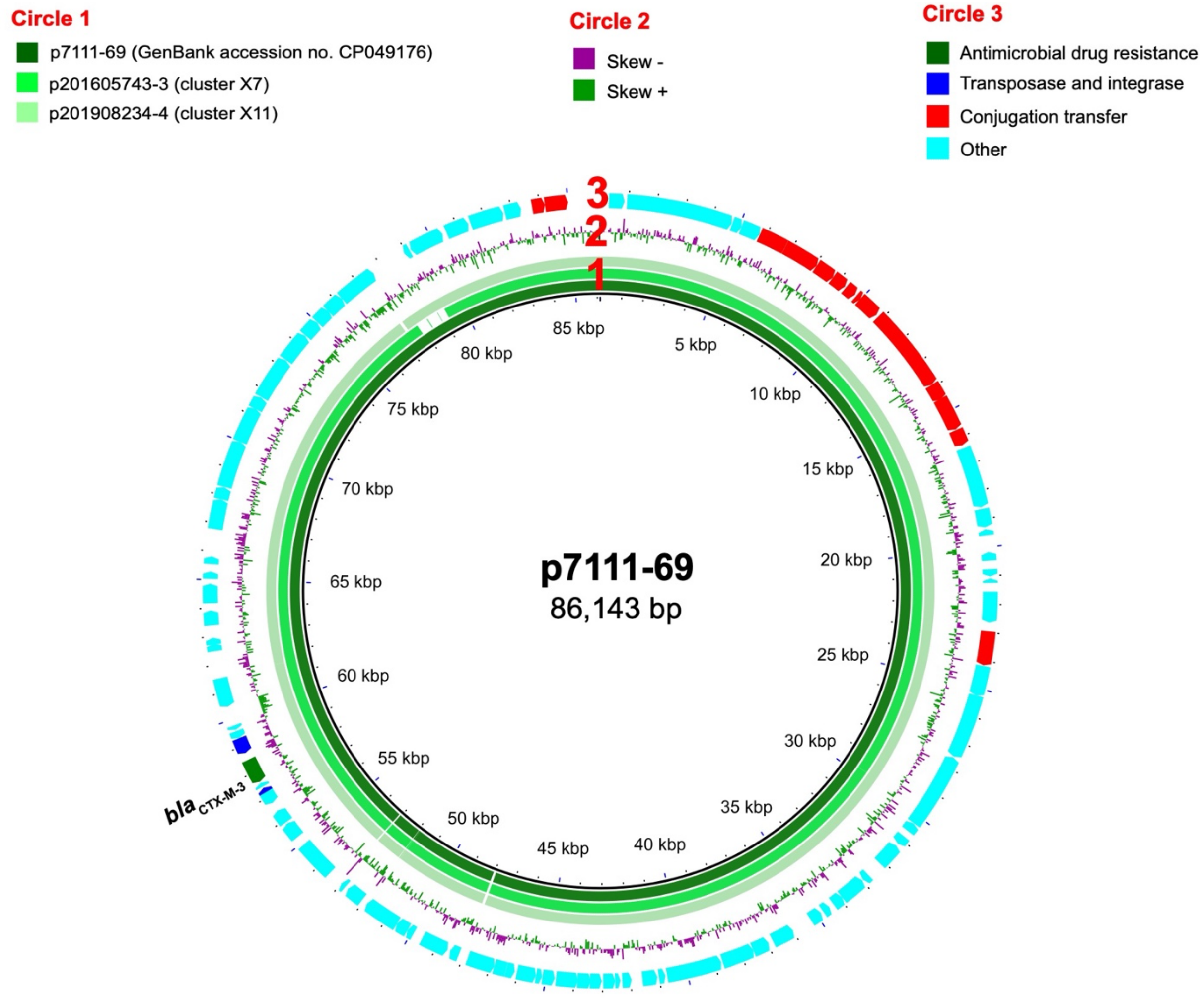


Circles from inside to outside indicate (1) the nucleotide position of p711-69, a plasmid from a *S. sonnei* isolate acquired in Turkey in 2019 (ref. [6]), and regions of p7111-69 displaying high levels of sequence identity to pKSR100 (from a *S. flexneri* 3a isolate, SF7955, collected in Canada in 2013)[3], p201605743-3 (XDR genomic cluster X7, our study), and p201908234-4 (cluster X11, our study), (2) a G+C content map of p711-69, and (3) coding sequences (CDS) colored according to their functions. The antimicrobial drug resistance gene is indicated.

**Supplementary Figure 7.** Circular map and comparative analysis of the ~ 8 kb MDR plasmids present in our XDR *S. sonnei* isolates

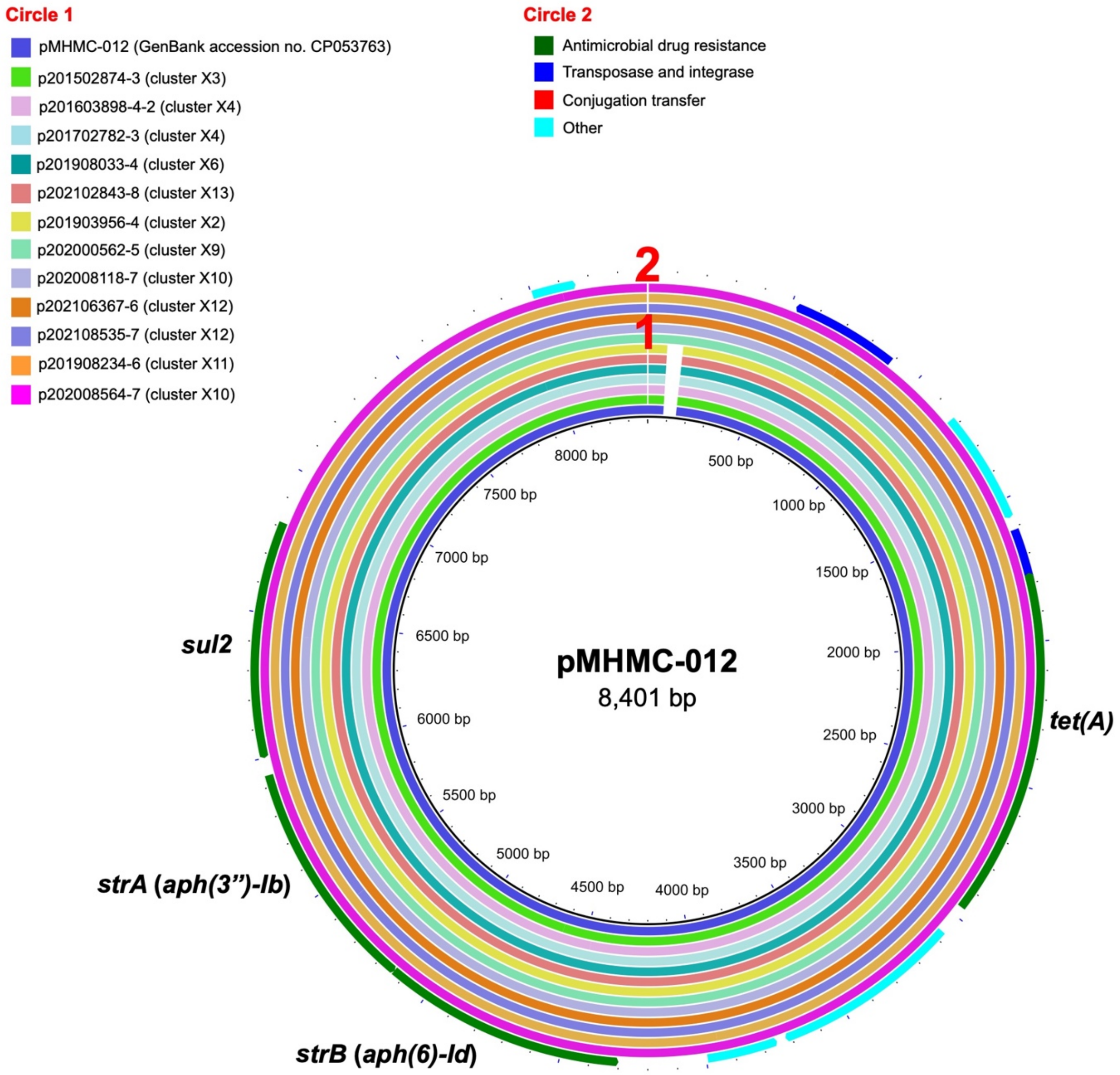


Circles from inside to outside indicate (1) the nucleotide position of pMHMC-012, a plasmid from a *S. sonnei* isolate acquired in Boston, USA in 2017 (ref. [1]), and regions of pMHMC-012 displaying high levels of identity to the sequences of 12 plasmids from XDR *S. sonnei* isolates in our study. The names of these plasmids and the XDR genomic clusters of their host *S. sonnei* isolates are shown in the legend, and (2) coding sequences (CDS) colored according to their functions. The antimicrobial drug resistance genes are indicated.

**Supplementary Table 1.** Phenotypic characteristics and metadata for the 164 XDR *S. sonnei* studied.

| Isolate | Year | Age group (year) | Sex | French region | Travel | 2017 school outbreak | Phenotypic antimicrobial drug resistance profile | MIC (mg/L) | | |
|---|---|---|---|---|---|---|---|---|---|---|
| | | | | | | | | CIP | AZM | CRO |
| 201502874 | 2015 | >15 | M | Metropole | Unknown | No | AMP CRO STR SMX TMP SXT TET NAL CIP AZM | 4 | 96 | 24 |
| 201603898 | 2016 | >15 | M | Metropole | Vietnam | No | AMP CRO STR GEN SMX TMP SXT TET NAL CIP AZM | 2 | 48 | 32 |
| 201605743 | 2016 | >15 | M | Metropole | Unknown | No | AMP CRO STR SMX TMP SXT NAL CIP AZM | 3 | ≥256 | 64 |
| 201701093 | 2017 | >15 | F | Metropole | None | No | AMP CRO SMX TMP SXT NAL CIP AZM | 8 | ≥256 | 128 |
| 201702421 | 2017 | 0-15 | F | Metropole | None | Yes | AMP CRO STR GEN SMX TMP SXT TET NAL CIP AZM | 3 | 64 | 96 |
| 201702422 | 2017 | 0-15 | F | Metropole | None | Yes | AMP CRO STR GEN SMX TMP SXT TET NAL CIP AZM | 4 | 64 | 64 |
| 201702423 | 2017 | 0-15 | M | Metropole | None | Yes | AMP CRO STR GEN SMX TMP SXT TET NAL CIP AZM | 4 | 48 | 192 |
| 201702424 | 2017 | 0-15 | F | Metropole | None | Yes | AMP CRO STR GEN SMX TMP SXT TET NAL CIP AZM | 4 | 128 | 96 |
| 201702425 | 2017 | 0-15 | M | Metropole | None | Yes | AMP CRO STR GEN SMX TMP SXT TET NAL CIP AZM | 4 | 64 | ≥256 |
| 201702483 | 2017 | 0-15 | F | Metropole | None | Yes | AMP CRO STR GEN SMX TMP SXT TET NAL CIP AZM | 3 | 64 | 64 |
| 201702484 | 2017 | 0-15 | M | Metropole | None | Yes | AMP CRO STR GEN SMX TMP SXT TET NAL CIP AZM | 3 | 64 | 64 |
| 201702485 | 2017 | 0-15 | M | Metropole | None | Yes | AMP CRO STR GEN SMX TMP SXT TET NAL CIP AZM | 3 | 96 | 192 |
| 201702486 | 2017 | 0-15 | M | Metropole | None | Yes | AMP CRO STR GEN SMX TMP SXT TET NAL CIP AZM | 3 | 64 | 96 |
| 201702487 | 2017 | 0-15 | M | Metropole | None | Yes | AMP CRO STR GEN SMX TMP SXT TET NAL CIP AZM | 2 | 64 | 64 |
| 201702488 | 2017 | 0-15 | F | Metropole | None | Yes | AMP CRO STR GEN SMX TMP SXT TET NAL CIP AZM | 3 | 64 | 64 |
| 201702489 | 2017 | 0-15 | M | Metropole | None | Yes | AMP CRO STR GEN SMX TMP SXT TET NAL CIP AZM | 3 | 64 | 96 |
| 201702490 | 2017 | 0-15 | F | Metropole | None | Yes | AMP CRO STR GEN SMX TMP SXT TET NAL CIP AZM | 2 | 64 | 48 |
| 201702491 | 2017 | 0-15 | M | Metropole | None | Yes | AMP CRO STR GEN SMX TMP SXT TET NAL CIP AZM | 3 | 64 | 96 |
| 201702544 | 2017 | 0-15 | F | Metropole | None | Yes | AMP CRO STR GEN SMX TMP SXT TET NAL CIP AZM | 3 | 96 | ≥256 |
| 201702545 | 2017 | 0-15 | M | Metropole | None | Yes | AMP CRO STR GEN SMX TMP SXT TET NAL CIP AZM | 2 | 64 | 96 |
| 201702562 | 2017 | 0-15 | M | Metropole | None | Yes | AMP CRO STR GEN SMX TMP SXT TET NAL CIP AZM | 3 | 64 | 64 |
| 201702563 | 2017 | 0-15 | F | Metropole | None | Yes | AMP CRO STR GEN SMX TMP SXT TET NAL CIP AZM | 3 | 96 | 32 |
| 201702620 | 2017 | >15 | M | Metropole | None | Yes | AMP CRO STR GEN SMX TMP SXT TET NAL CIP AZM | 3 | 128 | ≥256 |
| 201702781 | 2017 | 0-15 | M | Metropole | Vietnam | Yes | AMP CRO STR GEN SMX TMP SXT TET NAL CIP AZM | 3 | 96 | ≥256 |
| 201702782 | 2017 | 0-15 | F | Metropole | None | Yes | AMP CRO STR GEN SMX TMP SXT TET NAL CIP AZM | 4 | 128 | 64 |
| 201702783 | 2017 | 0-15 | M | Metropole | None | Yes | AMP CRO STR GEN SMX TMP SXT TET NAL CIP AZM | 3 | 96 | 96 |
| 201702784 | 2017 | 0-15 | M | Metropole | None | Yes | AMP CRO STR GEN SMX TMP SXT TET NAL CIP AZM | 3 | 96 | 64 |
| 201702786 | 2017 | 0-15 | M | Metropole | None | Yes | AMP CRO STR GEN SMX TMP SXT TET NAL CIP AZM | 3 | 96 | 192 |
| 201702787 | 2017 | >15 | M | Metropole | None | Yes | AMP CRO STR GEN SMX TMP SXT TET NAL CIP AZM | 3 | 96 | 128 |
| 201702788 | 2017 | >15 | F | Metropole | None | Yes | AMP CRO STR GEN SMX TMP SXT TET NAL CIP AZM | 3 | 48 | 128 |
| 201702789 | 2017 | 0-15 | F | Metropole | None | Yes | AMP CRO STR GEN SMX TMP SXT TET NAL CIP AZM | 3 | 128 | 96 |
| 201702790 | 2017 | 0-15 | M | Metropole | None | Yes | AMP CRO STR GEN SMX TMP SXT TET NAL CIP AZM | 3 | 96 | 96 |
| 201702791 | 2017 | 0-15 | M | Metropole | None | Yes | AMP CRO STR GEN SMX TMP SXT TET NAL CIP AZM | 3 | 128 | 64 |
| 201702847 | 2017 | >15 | F | Metropole | None | Yes | AMP CRO STR GEN SMX TMP SXT TET NAL CIP AZM | 3 | 96 | 96 |
| 201703089 | 2017 | >15 | F | Metropole | None | Yes | AMP CRO STR GEN SMX TMP SXT TET NAL CIP AZM | >32 | 96 | 48 |
| 201703090 | 2017 | 0-15 | M | Metropole | None | Yes | AMP CRO STR GEN SMX TMP SXT TET NAL CIP AZM | 4 | 128 | ≥256 |
| 201703128 | 2017 | 0-15 | F | Metropole | India | No | AMP CRO STR SMX TMP SXT NAL CIP AZM | 6 | 128 | 12 |
| 201703316 | 2017 | >15 | F | Metropole | India | No | AMP CRO STR SMX TMP SXT NAL CIP AZM | 6 | 192 | 16 |
| 201802246 | 2018 | 0-15 | F | Metropole | Vietnam | No | AMP CRO STR SMX TMP SXT TET NAL CIP AZM | 4 | 128 | 128 |
| 201807563 | 2018 | >15 | M | Metropole | None | No | AMP CRO STR SMX TMP SXT TET NAL CIP AZM | >32 | ≥256 | 128 |
| 201808736 | 2018 | >15 | M | Metropole | Unknown | No | AMP CRO STR SMX TMP SXT TET NAL CIP AZM | 24 | ≥256 | 24 |
| 201809101 | 2018 | >15 | M | Metropole | None | No | AMP CRO STR SMX TMP SXT TET NAL CIP AZM | 24 | ≥256 | 24 |

| | | | | | | | | | | |
|---|---|---|---|---|---|---|---|---|---|---|
| 201810383 | 2018 | >15 | M | Metropole | None | No | AMP CRO STR SMX TMP SXT TET NAL CIP AZM | >32 | ≥256 | 12 |
| 201810803 | 2018 | >15 | M | Metropole | Spain | No | AMP CRO STR SMX TMP SXT TET NAL CIP AZM | 24 | ≥256 | 32 |
| 201900053 | 2019 | >15 | M | Metropole | Unknown | No | AMP CRO STR SMX TMP SXT TET NAL CIP AZM | 24 | ≥256 | 32 |
| 201903956 | 2019 | >15 | F | Metropole | Cambodia | No | AMP CRO STR SMX TMP SXT TET NAL CIP AZM | 6 | 192 | 64 |
| 201904981 | 2019 | >15 | M | Metropole | None | No | AMP CRO STR SMX TMP SXT TET NAL CIP AZM | >32 | ≥256 | 24 |
| 201908033 | 2019 | 0-15 | M | Metropole | None | No | AMP CRO STR SMX TMP SXT TET NAL CIP AZM | 4 | ≥256 | ≥256 |
| 201908234 | 2019 | >15 | M | Metropole | None | No | AMP CRO STR SMX TMP SXT TET NAL CIP AZM | 4 | ≥256 | 32 |
| 201909147 | 2019 | >15 | M | Metropole | None | No | AMP CRO STR SMX TMP SXT TET NAL CIP AZM | 24 | ≥256 | 24 |
| 201910769 | 2019 | >15 | M | Metropole | Spain | No | AMP CRO STR SMX TMP SXT TET NAL CIP AZM | >32 | ≥256 | 128 |
| 201910803 | 2019 | >15 | M | Metropole | None | No | AMP CRO STR SMX TMP SXT TET NAL CIP AZM | >32 | ≥256 | 48 |
| 202000267 | 2020 | >15 | M | Metropole | None | No | AMP CRO STR SMX TMP SXT TET NAL CIP AZM | 4 | 192 | 24 |
| 202000322 | 2020 | >15 | M | Metropole | Unknown | No | AMP CRO STR SMX TMP SXT TET NAL CIP AZM | 4 | 192 | 48 |
| 202000562 | 2020 | >15 | M | Metropole | None | No | AMP CRO STR SMX TMP SXT TET NAL CIP AZM | 3 | 192 | 48 |
| 202001062 | 2020 | >15 | M | Metropole | Switzerland | No | AMP CRO STR SMX TMP SXT TET NAL CIP AZM | 4 | 192 | 96 |
| 202001063 | 2020 | >15 | F | Metropole | Spain | No | AMP CRO STR SMX TMP SXT TET NAL CIP AZM | 4 | 192 | 96 |
| 202001623 | 2020 | >15 | M | Metropole | None | No | AMP CRO STR SMX TMP SXT TET NAL CIP AZM | 4 | 192 | 128 |
| 202005154 | 2020 | >15 | F | La Réunion | None | No | AMP CRO SMX TMP SXT NAL CIP AZM | 6 | ≥256 | 16 |
| 202006637 | 2020 | >15 | M | Metropole | None | No | AMP CRO STR SMX TMP SXT TET NAL CIP AZM | 12 | ≥256 | 18 |
| 202007856 | 2020 | >15 | M | Metropole | None | No | AMP CRO STR SMX TMP SXT TET NAL CIP AZM | 12 | ≥256 | 48 |
| 202008118 | 2020 | >15 | M | Metropole | None | No | AMP CRO STR SMX TMP SXT TET NAL CIP AZM | 8 | 192 | 32 |
| 202008158 | 2020 | >15 | M | Metropole | None | No | AMP CRO STR SMX TMP SXT TET NAL CIP AZM | 12 | ≥256 | 16 |
| 202008564 | 2020 | >15 | M | Metropole | None | No | AMP CRO STR SMX TMP SXT TET NAL CIP AZM | 12 | ≥256 | 32 |
| 202008707 | 2020 | >15 | M | Metropole | None | No | AMP CRO STR SMX TMP SXT TET NAL CIP AZM | 8 | 128 | 16 |
| 202100373 | 2021 | >15 | M | Metropole | Unknown | No | AMP CRO STR SMX TMP SXT TET NAL CIP AZM | 8 | 192 | 24 |
| 202100420 | 2021 | >15 | M | Metropole | Unknown | No | AMP CRO STR SMX TMP SXT TET NAL CIP AZM | 8 | 192 | 128 |
| 202100759 | 2021 | >15 | M | Metropole | Unknown | No | AMP CRO STR SMX TMP SXT TET NAL CIP AZM | 12 | ≥256 | 16 |
| 202101412 | 2021 | >15 | M | Metropole | Unknown | No | AMP CRO STR SMX TMP SXT TET NAL CIP AZM | 12 | ≥256 | 32 |
| 202101541 | 2021 | >15 | M | Metropole | None | No | AMP CRO STR SMX TMP SXT NAL CIP AZM | 8 | ≥256 | 48 |
| 202101624 | 2021 | >15 | M | Metropole | Unknown | No | AMP CRO STR SMX TMP SXT TET NAL CIP AZM | 12 | ≥256 | 96 |
| 202101651 | 2021 | >15 | M | Metropole | None | No | AMP CRO STR SMX TMP SXT TET NAL CIP AZM | 32 | ≥256 | 64 |
| 202101727 | 2021 | >15 | M | Metropole | None | No | AMP CRO STR SMX TMP SXT TET NAL CIP AZM | 12 | ≥256 | 24 |
| 202101879 | 2021 | >15 | M | Metropole | None | No | AMP CRO STR SMX TMP SXT TET NAL CIP AZM | 6 | ≥256 | 32 |
| 202101885 | 2021 | >15 | M | Metropole | Unknown | No | AMP CRO STR SMX TMP SXT TET NAL CIP AZM | 8 | 192 | 32 |
| 202101930 | 2021 | >15 | M | Metropole | Unknown | No | AMP CRO STR SMX TMP SXT TET NAL CIP AZM | 12 | ≥256 | 24 |
| 202102119 | 2021 | >15 | M | Metropole | Unknown | No | AMP CRO STR SMX TMP SXT TET NAL CIP AZM | 8 | ≥256 | 48 |
| 202102200 | 2021 | >15 | M | Metropole | Unknown | No | AMP CRO STR SMX TMP SXT TET NAL CIP AZM | 8 | ≥256 | 32 |
| 202102776 | 2021 | >15 | M | Metropole | None | No | AMP CRO STR SMX TMP SXT TET NAL CIP AZM | 8 | ≥256 | 48 |
| 202102820 | 2021 | >15 | M | Metropole | None | No | AMP CRO STR SMX TMP SXT TET NAL CIP AZM | 8 | ≥256 | 24 |
| 202102843 | 2021 | >15 | M | Metropole | None | No | AMP CRO STR SMX TMP SXT TET NAL CIP AZM | 6 | ≥256 | 12 |
| 202102980 | 2021 | >15 | M | Metropole | None | No | AMP CRO STR SMX TMP SXT TET NAL CIP AZM | 6 | 128 | 24 |
| 202102993 | 2021 | >15 | M | Metropole | Unknown | No | AMP CRO STR SMX TMP SXT TET NAL CIP AZM | 8 | ≥256 | 64 |
| 202103005 | 2021 | >15 | M | Metropole | None | No | AMP CRO STR SMX TMP SXT TET NAL CIP AZM | 6 | 96 | 24 |
| 202103026 | 2021 | >15 | M | Metropole | None | No | AMP CRO STR SMX TMP SXT TET NAL CIP AZM | 8 | ≥256 | 32 |
| 202103166 | 2021 | >15 | M | Metropole | None | No | AMP CRO STR SMX TMP SXT TET NAL CIP AZM | 6 | ≥256 | 24 |
| 202103248 | 2021 | >15 | M | Metropole | None | No | AMP CRO STR SMX TMP SXT TET NAL CIP AZM | 6 | ≥256 | 24 |
| 202103290 | 2021 | >15 | M | Metropole | None | No | AMP CRO STR SMX TMP SXT TET NAL CIP AZM | 6 | 32 | 24 |
| 202103342 | 2021 | >15 | M | Metropole | None | No | AMP CRO STR SMX TMP SXT TET NAL CIP AZM | 6 | 96 | 24 |
| 202103419 | 2021 | >15 | M | Metropole | None | No | AMP CRO STR SMX TMP SXT TET NAL CIP AZM | 6 | 96 | 24 |
| 202103424 | 2021 | >15 | M | Metropole | Cameroon | No | AMP CRO STR SMX TMP SXT TET NAL CIP AZM | 6 | 32 | 48 |
| 202103567 | 2021 | >15 | M | Metropole | Unknown | No | AMP CRO STR SMX TMP SXT TET NAL CIP AZM | 8 | ≥256 | 24 |
| 202103754 | 2021 | >15 | M | Metropole | Unknown | No | AMP CRO STR SMX TMP SXT TET NAL CIP AZM | 6 | ≥256 | 32 |

| | | | | | | | | | | |
|---|---|---|---|---|---|---|---|---|---|---|
| 202103844 | 2021 | >15 | M | Metropole | Unknown | No | AMP CRO STR SMX TMP SXT TET NAL CIP AZM | 6 | ≥256 | 48 |
| 202103880 | 2021 | >15 | M | Metropole | None | No | AMP CRO STR SMX TMP SXT TET NAL CIP AZM | 8 | 48 | 32 |
| 202103930 | 2021 | >15 | M | Metropole | None | No | AMP CRO STR SMX TMP SXT TET NAL CIP AZM | 6 | ≥256 | 32 |
| 202104060 | 2021 | >15 | F | Metropole | Unknown | No | AMP CRO STR SMX TMP SXT TET NAL CIP AZM | 4 | ≥256 | 32 |
| 202104078 | 2021 | >15 | M | Metropole | None | No | AMP CRO STR SMX TMP SXT TET NAL CIP AZM | 8 | 128 | 48 |
| 202104261 | 2021 | >15 | M | Metropole | None | No | AMP CRO STR SMX TMP SXT TET NAL CIP AZM | 6 | ≥256 | 16 |
| 202104281 | 2021 | >15 | M | Metropole | None | No | AMP CRO STR SMX TMP SXT TET NAL CIP AZM | 24 | ≥256 | 96 |
| 202104371 | 2021 | >15 | M | Metropole | Unknown | No | AMP CRO STR SMX TMP SXT TET NAL CIP AZM | 6 | ≥256 | 24 |
| 202104519 | 2021 | >15 | M | Metropole | None | No | AMP CRO STR SMX TMP SXT TET NAL CIP AZM | 6 | ≥256 | 16 |
| 202104685 | 2021 | >15 | M | Metropole | Unknown | No | AMP CRO STR SMX TMP SXT TET NAL CIP AZM | 6 | ≥256 | 16 |
| 202104831 | 2021 | >15 | M | Metropole | None | No | AMP CRO STR SMX TMP SXT TET NAL CIP AZM | 8 | ≥256 | 48 |
| 202105132 | 2021 | >15 | M | Metropole | Unknown | No | AMP CRO STR SMX TMP SXT TET NAL CIP AZM | 8 | 192 | 32 |
| 202105166 | 2021 | >15 | M | Metropole | Unknown | No | AMP CRO STR SMX TMP SXT TET NAL CIP AZM | 6 | ≥256 | 32 |
| 202105284 | 2021 | >15 | M | Metropole | None | No | AMP CRO STR SMX TMP SXT TET NAL CIP AZM | 8 | ≥256 | 192 |
| 202105285 | 2021 | >15 | M | Metropole | None | No | AMP CRO STR SMX TMP SXT TET NAL CIP AZM | 12 | 96 | 48 |
| 202105373 | 2021 | >15 | M | Metropole | Unknown | No | AMP CRO STR SMX TMP SXT TET NAL CIP AZM | 16 | ≥256 | 24 |
| 202105408 | 2021 | >15 | M | Metropole | Unknown | No | AMP CRO STR SMX TMP SXT TET NAL CIP AZM | 12 | ≥256 | 24 |
| 202105440 | 2021 | >15 | M | Metropole | Unknown | No | AMP CRO STR SMX TMP SXT TET NAL CIP AZM | 8 | ≥256 | 192 |
| 202105446 | 2021 | >15 | M | Metropole | Unknown | No | AMP CRO TMP NAL CIP AZM | 12 | 48 | 48 |
| 202105523 | 2021 | >15 | M | Metropole | None | No | AMP CRO STR SMX TMP SXT TET NAL CIP AZM | 4 | ≥256 | 64 |
| 202105574 | 2021 | >15 | M | Metropole | None | No | AMP CRO STR SMX TMP SXT TET NAL CIP AZM | 12 | ≥256 | 64 |
| 202105953 | 2021 | >15 | F | Metropole | Unknown | No | AMP CRO STR SMX TMP SXT TET NAL CIP AZM | 12 | ≥256 | 32 |
| 202106029 | 2021 | >15 | M | Metropole | Unknown | No | AMP CRO STR SMX TMP SXT TET NAL CIP AZM | 8 | ≥256 | 32 |
| 202106219 | 2021 | >15 | M | Metropole | None | No | AMP CRO STR SMX TMP SXT TET NAL CIP AZM | 8 | ≥256 | 24 |
| 202106247 | 2021 | 0-15 | M | Metropole | Unknown | No | AMP CRO STR SMX TMP SXT TET NAL CIP AZM | 12 | ≥256 | 16 |
| 202106367 | 2021 | >15 | F | Metropole | Lebanon | No | AMP CRO STR SMX TMP SXT TET NAL CIP AZM | 1 | 96 | 6 |
| 202106455 | 2021 | >15 | M | Metropole | None | No | AMP CRO STR SMX TMP SXT TET NAL CIP AZM | 4 | ≥256 | 24 |
| 202106629 | 2021 | >15 | M | Metropole | Unknown | No | AMP CRO STR SMX TMP SXT TET NAL CIP AZM | 32 | ≥256 | 24 |
| 202106664 | 2021 | >15 | M | Metropole | None | No | AMP CRO STR SMX TMP SXT NAL CIP AZM | 16 | ≥256 | 24 |
| 202106716 | 2021 | >15 | M | Metropole | Spain | No | AMP CRO STR SMX TMP SXT TET NAL CIP AZM | 12 | ≥256 | 24 |
| 202106961 | 2021 | >15 | M | Metropole | Unknown | No | AMP CRO STR SMX TMP SXT TET NAL CIP AZM | 32 | ≥256 | 12 |
| 202107022 | 2021 | >15 | M | Metropole | None | No | AMP CRO STR SMX TMP SXT TET NAL CIP AZM | 12 | ≥256 | 32 |
| 202107051 | 2021 | >15 | M | Metropole | None | No | AMP CRO STR SMX TMP SXT TET NAL CIP AZM | 16 | ≥256 | 48 |
| 202107052 | 2021 | >15 | M | Metropole | None | No | AMP CRO STR SMX TMP SXT TET NAL CIP AZM | 16 | ≥256 | 32 |
| 202107188 | 2021 | >15 | M | Metropole | Unknown | No | AMP CRO STR SMX TMP SXT NAL CIP AZM | 16 | ≥256 | 32 |
| 202107323 | 2021 | >15 | M | Metropole | Unknown | No | AMP CRO STR SMX TMP SXT TET NAL CIP AZM | 16 | ≥256 | 32 |
| 202107395 | 2021 | >15 | M | Metropole | Unknown | No | AMP CRO STR SMX TMP SXT TET NAL CIP AZM | 6 | ≥256 | 48 |
| 202107415 | 2021 | >15 | M | Metropole | Unknown | No | AMP CRO STR SMX TMP SXT TET NAL CIP AZM | 6 | ≥256 | 48 |
| 202107446 | 2021 | >15 | M | Metropole | None | No | AMP CRO STR SMX TMP SXT TET NAL CIP AZM | 12 | ≥256 | 48 |
| 202107466 | 2021 | >15 | M | Metropole | None | No | AMP CRO STR SMX TMP SXT TET NAL CIP AZM | 6 | ≥256 | 192 |
| 202107818 | 2021 | >15 | M | Metropole | Unknown | No | AMP CRO STR SMX TMP SXT TET NAL CIP AZM | 6 | ≥256 | 32 |
| 202108294 | 2021 | >15 | M | Metropole | Unknown | No | AMP CRO STR SMX TMP SXT TET NAL CIP AZM | 12 | ≥256 | 24 |
| 202108535 | 2021 | >15 | F | Metropole | None | No | AMP CRO STR SMX TMP SXT TET NAL CIP AZM | 1 | 96 | 16 |
| 202108795 | 2021 | >15 | M | Metropole | Unknown | No | AMP CRO STR SMX TMP SXT TET NAL CIP AZM | 6 | ≥256 | 24 |
| 202108919 | 2021 | >15 | M | Metropole | None | No | AMP CRO STR SMX TMP SXT TET NAL CIP AZM | 4 | ≥256 | 24 |
| 202108953 | 2021 | >15 | M | Metropole | Unknown | No | AMP CRO STR SMX TMP SXT TET NAL CIP AZM | 8 | ≥256 | 48 |
| 202109588 | 2021 | >15 | M | Metropole | Unknown | No | AMP CRO STR SMX TMP SXT TET NAL CIP AZM | 6 | ≥256 | 24 |
| 202109651 | 2021 | >15 | M | Metropole | None | No | AMP CRO STR SMX TMP SXT TET NAL CIP AZM | 4 | ≥256 | 24 |
| 202109656 | 2021 | >15 | M | Metropole | Unknown | No | AMP CRO STR SMX TMP SXT TET NAL CIP AZM | 16 | ≥256 | 24 |
| 202109849 | 2021 | >15 | M | Metropole | Unknown | No | AMP CRO STR SMX TMP SXT NAL CIP AZM | 12 | ≥256 | 24 |
| 202109909 | 2021 | >15 | M | Metropole | Unknown | No | AMP CRO STR SMX TMP SXT TET NAL CIP AZM | 4 | ≥256 | 96 |

| | | | | | | | | | | |
|---|---|---|---|---|---|---|---|---|---|---|
| 202109960 | 2021 | >15 | M | Metropole | Unknown | No | AMP CRO STR SMX TMP SXT TET NAL CIP AZM | 6 | ≥256 | 48 |
| 202110146 | 2021 | >15 | M | Metropole | Unknown | No | AMP CRO STR SMX TMP SXT NAL CIP AZM | 8 | ≥256 | 32 |
| 202110251 | 2021 | >15 | M | Metropole | Unknown | No | AMP CRO STR SMX TMP SXT NAL CIP AZM | 8 | ≥256 | 24 |
| 202110403 | 2021 | >15 | M | Metropole | Unknown | No | AMP CRO STR SMX TMP SXT TET NAL CIP AZM | 12 | ≥256 | 32 |
| 202110415 | 2021 | >15 | M | Metropole | None | No | AMP CRO STR SMX TMP SXT NAL CIP AZM | 8 | ≥256 | 32 |
| 202110472 | 2021 | >15 | M | Metropole | None | No | AMP CRO STR SMX TMP SXT TET NAL CIP AZM | 8 | ≥256 | 32 |
| 202110584 | 2021 | >15 | M | Metropole | Unknown | No | AMP CRO STR SMX TMP SXT NAL CIP AZM | 8 | ≥256 | 32 |
| 202110641 | 2021 | >15 | M | Metropole | Unknown | No | AMP CRO STR SMX TMP SXT NAL CIP AZM | 8 | ≥256 | 32 |
| 202110674 | 2021 | >15 | M | Metropole | Unknown | No | AMP CRO STR SMX TMP SXT NAL CIP AZM | 8 | ≥256 | 24 |
| 202110841 | 2021 | >15 | M | Metropole | Spain | No | AMP CRO STR SMX TMP SXT NAL CIP AZM | 8 | ≥256 | 16 |
| 202110869 | 2021 | >15 | F | Metropole | Unknown | No | AMP CRO STR SMX TMP SXT NAL CIP AZM | 16 | ≥256 | 48 |
| 202110877 | 2021 | >15 | M | Metropole | Unknown | No | AMP CRO STR SMX TMP SXT NAL CIP AZM | 12 | ≥256 | 48 |
| 202110960 | 2021 | >15 | M | Metropole | Unknown | No | AMP CRO STR SMX TMP SXT TET NAL CIP AZM | 16 | ≥256 | 24 |
| 202110961 | 2021 | >15 | M | Metropole | Unknown | No | AMP CRO STR SMX TMP SXT TET NAL CIP AZM | 12 | ≥256 | 32 |
| 202111030 | 2021 | >15 | M | Metropole | Unknown | No | AMP CRO STR SMX TMP SXT TET NAL CIP AZM | 12 | ≥256 | 24 |
| 202111082 | 2021 | >15 | M | Metropole | Unknown | No | AMP CRO STR SMX TMP SXT TET NAL CIP AZM | 12 | ≥256 | 32 |
| 202111134 | 2021 | >15 | M | Metropole | Unknown | No | AMP CRO STR SMX TMP SXT TET NAL CIP AZM | 4 | ≥256 | 24 |
| 202111144 | 2021 | >15 | M | Metropole | Unknown | No | AMP CRO STR SMX TMP SXT TET NAL CIP AZM | 8 | 192 | 48 |
| 202111184 | 2021 | >15 | M | Metropole | None | No | AMP CRO STR SMX TMP SXT NAL CIP AZM | 12 | ≥256 | 24 |
| 202200143 | 2021 | >15 | M | Metropole | Unknown | No | AMP CRO STR SMX TMP SXT NAL CIP AZM | 8 | ≥256 | 48 |

M, male; F, female; MIC; minimum inhibitory concentration; AMP, ampicillin; CRO, ceftriaxone; STR, streptomycin; SMX, sulfamethoxazole; TMP, trimethoprim; TET, tetracycline; NAL, nalidixic acid; CIP, ciprofloxacin; AZM, azithromycin.

**Supplementary Table 2.** Genomic characteristics for the 164 XDR *S. sonnei* isolates under study.

| Isolate | Cluster | Genotype | Genotype alias | Acquired antimicrobial drug resistance genes | QRDR mutation *gyrA* | QRDR mutation *parC* | Accession no. |
|---|---|---|---|---|---|---|---|
| 201701093 | X1 | 3.6.1.1 | CipR | *bla*$_{\text{CTX-M-15}}$, *sul1, dfrA1, dfrA5, qnrS13, mph(A), erm(B)* | S83L | S80I | ERS6492764 |
| 202005154 | X1 | 3.6.1.1 | CipR | *bla*$_{\text{CTX-M-15}}$, *sul1, dfrA1, dfrA5, qnrS13, mph(A), erm(B)* | S83L | S80I | ERS6576287 |
| 201903956 | X2 | 3.6.1.1.1 | CipR.SEA | *bla*$_{\text{CTX-M-27}}$, *strA, strB, aadA5, sul1, sul2, dfrA1, dfrA17, tet(A), mph(A)* | S83L, D87G | S80I | ERS6495067 |
| 201502874 | X3 | 3.6.1.1.1 | CipR.SEA | *bla*$_{\text{CTX-M-27}}$, *strA, strB, aadA5, sul1, sul2, dfrA1, dfrA17, tet(A), mph(A)* | S83L, D87G | S80I | ERS12446082 |
| 201603898 | X4 | 3.6.1.1.1 | CipR.SEA | *bla*$_{\text{CTX-M-55}}$, *strA, strB, aac(3)-IIa, sul2, dfrA1, tet(A), mph(A)* | S83L, D87G | S80I | ERS12446174 |
| 201702421 | X4 | 3.6.1.1.1 | CipR.SEA | *bla*$_{\text{CTX-M-55}}$, *strA, strB, aac(3)-IIa, sul2, dfrA1, tet(A), mph(A)* | S83L, D87G | S80I | ERS6495734 |
| 201702422 | X4 | 3.6.1.1.1 | CipR.SEA | *bla*$_{\text{CTX-M-55}}$, *strA, strB, aac(3)-IIa, sul2, dfrA1, tet(A), mph(A)* | S83L, D87G | S80I | ERS6495735 |
| 201702423 | X4 | 3.6.1.1.1 | CipR.SEA | *bla*$_{\text{CTX-M-55}}$, *strA, strB, aac(3)-IIa, sul2, dfrA1, tet(A), mph(A)* | S83L, D87G | S80I | ERS6495736 |
| 201702424 | X4 | 3.6.1.1.1 | CipR.SEA | *bla*$_{\text{CTX-M-55}}$, *strA, strB, aac(3)-IIa, sul2, dfrA1, tet(A), mph(A)* | S83L, D87G | S80I | ERS6495737 |
| 201702425 | X4 | 3.6.1.1.1 | CipR.SEA | *bla*$_{\text{CTX-M-55}}$, *strA, strB, aac(3)-IIa, sul2, dfrA1, tet(A), mph(A)* | S83L, D87G | S80I | ERS6495738 |
| 201702483 | X4 | 3.6.1.1.1 | CipR.SEA | *bla*$_{\text{CTX-M-55}}$, *strA, strB, aac(3)-IIa, sul2, dfrA1, tet(A), mph(A)* | S83L, D87G | S80I | ERS6495742 |
| 201702484 | X4 | 3.6.1.1.1 | CipR.SEA | *bla*$_{\text{CTX-M-55}}$, *strA, strB, aac(3)-IIa, sul2, dfrA1, tet(A), mph(A)* | S83L, D87G | S80I | ERS6495743 |
| 201702485 | X4 | 3.6.1.1.1 | CipR.SEA | *bla*$_{\text{CTX-M-55}}$, *strA, strB, aac(3)-IIa, sul2, dfrA1, tet(A), mph(A)* | S83L, D87G | S80I | ERS6495744 |
| 201702486 | X4 | 3.6.1.1.1 | CipR.SEA | *bla*$_{\text{CTX-M-55}}$, *strA, strB, aac(3)-IIa, sul2, dfrA1, tet(A), mph(A)* | S83L, D87G | S80I | ERS6495745 |
| 201702487 | X4 | 3.6.1.1.1 | CipR.SEA | *bla*$_{\text{CTX-M-55}}$, *strA, strB, aac(3)-IIa, sul2, dfrA1, tet(A), mph(A)* | S83L, D87G | S80I | ERS6495746 |
| 201702488 | X4 | 3.6.1.1.1 | CipR.SEA | *bla*$_{\text{CTX-M-55}}$, *strA, strB, aac(3)-IIa, sul2, dfrA1, tet(A), mph(A)* | S83L, D87G | S80I | ERS6495747 |
| 201702489 | X4 | 3.6.1.1.1 | CipR.SEA | *bla*$_{\text{CTX-M-55}}$, *strA, strB, aac(3)-IIa, sul2, dfrA1, tet(A), mph(A)* | S83L, D87G | S80I | ERS6495748 |
| 201702490 | X4 | 3.6.1.1.1 | CipR.SEA | *bla*$_{\text{CTX-M-55}}$, *strA, strB, aac(3)-IIa, sul2, dfrA1, tet(A), mph(A)* | S83L, D87G | S80I | ERS6495749 |
| 201702491 | X4 | 3.6.1.1.1 | CipR.SEA | *bla*$_{\text{CTX-M-55}}$, *strA, strB, aac(3)-IIa, sul2, dfrA1, tet(A), mph(A)* | S83L, D87G | S80I | ERS6495750 |
| 201702544 | X4 | 3.6.1.1.1 | CipR.SEA | *bla*$_{\text{CTX-M-55}}$, *strA, strB, aac(3)-IIa, sul2, dfrA1, tet(A), mph(A)* | S83L, D87G | S80I | ERS6495756 |
| 201702545 | X4 | 3.6.1.1.1 | CipR.SEA | *bla*$_{\text{CTX-M-55}}$, *strA, strB, aac(3)-IIa, sul2, dfrA1, tet(A), mph(A)* | S83L, D87G | S80I | ERS6495757 |
| 201702562 | X4 | 3.6.1.1.1 | CipR.SEA | *bla*$_{\text{CTX-M-55}}$, *strA, strB, aac(3)-IIa, sul2, dfrA1, tet(A), mph(A)* | S83L, D87G | S80I | ERS6495758 |
| 201702563 | X4 | 3.6.1.1.1 | CipR.SEA | *bla*$_{\text{CTX-M-55}}$, *strA, strB, aac(3)-IIa, sul2, dfrA1, tet(A), mph(A)* | S83L, D87G | S80I | ERS6495759 |
| 201702620 | X4 | 3.6.1.1.1 | CipR.SEA | *bla*$_{\text{CTX-M-55}}$, *strA, strB, aac(3)-IIa, sul2, dfrA1, tet(A), mph(A)* | S83L, D87G | S80I | ERS6495761 |
| 201702781 | X4 | 3.6.1.1.1 | CipR.SEA | *bla*$_{\text{CTX-M-55}}$, *strA, strB, aac(3)-IIa, sul2, dfrA1, tet(A), mph(A)* | S83L, D87G | S80I | ERS6495763 |
| 201702782 | X4 | 3.6.1.1.1 | CipR.SEA | *bla*$_{\text{CTX-M-55}}$, *strA, strB, aac(3)-IIa, sul2, dfrA1, tet(A), mph(A)* | S83L, D87G | S80I | ERS6495764 |
| 201702783 | X4 | 3.6.1.1.1 | CipR.SEA | *bla*$_{\text{CTX-M-55}}$, *strA, strB, aac(3)-IIa, sul2, dfrA1, tet(A), mph(A)* | S83L, D87G | S80I | ERS6495765 |
| 201702784 | X4 | 3.6.1.1.1 | CipR.SEA | *bla*$_{\text{CTX-M-55}}$, *strA, strB, aac(3)-IIa, sul2, dfrA1, tet(A), mph(A)* | S83L, D87G | S80I | ERS6495766 |
| 201702786 | X4 | 3.6.1.1.1 | CipR.SEA | *bla*$_{\text{CTX-M-55}}$, *strA, strB, aac(3)-IIa, sul2, dfrA1, tet(A), mph(A)* | S83L, D87G | S80I | ERS6495768 |
| 201702787 | X4 | 3.6.1.1.1 | CipR.SEA | *bla*$_{\text{CTX-M-55}}$, *strA, strB, aac(3)-IIa, sul2, dfrA1, tet(A), mph(A)* | S83L, D87G | S80I | ERS6575823 |
| 201702788 | X4 | 3.6.1.1.1 | CipR.SEA | *bla*$_{\text{CTX-M-55}}$, *strA, strB, aac(3)-IIa, sul2, dfrA1, tet(A), mph(A)* | S83L, D87G | S80I | ERS6575824 |
| 201702789 | X4 | 3.6.1.1.1 | CipR.SEA | *bla*$_{\text{CTX-M-55}}$, *strA, strB, aac(3)-IIa, sul2, dfrA1, tet(A), mph(A)* | S83L, D87G | S80I | ERS6575825 |
| 201702790 | X4 | 3.6.1.1.1 | CipR.SEA | *bla*$_{\text{CTX-M-55}}$, *strA, strB, aac(3)-IIa, sul2, dfrA1, tet(A), mph(A)* | S83L, D87G | S80I | ERS6575826 |
| 201702791 | X4 | 3.6.1.1.1 | CipR.SEA | *bla*$_{\text{CTX-M-55}}$, *strA, strB, aac(3)-IIa, sul2, dfrA1, tet(A), mph(A)* | S83L, D87G | S80I | ERS6575827 |
| 201702847 | X4 | 3.6.1.1.1 | CipR.SEA | *bla*$_{\text{CTX-M-55}}$, *strA, strB, aac(3)-IIa, sul2, dfrA1, tet(A), mph(A)* | S83L, D87G | S80I | ERS6575830 |
| 201703089 | X4 | 3.6.1.1.1 | CipR.SEA | *bla*$_{\text{CTX-M-55}}$, *strA, strB, aac(3)-IIa, sul2, dfrA1, tet(A), mph(A)* | S83L, D87G | S80I | ERS6575842 |
| 201703090 | X4 | 3.6.1.1.1 | CipR.SEA | *bla*$_{\text{CTX-M-55}}$, *strA, strB, aac(3)-IIa, sul2, dfrA1, tet(A), mph(A)* | S83L, D87G | S80I | ERS6575843 |

| | | | | | | | |
|---|---|---|---|---|---|---|---|
| 201802246 | X4 | 3.6.1.1.1 | CipR.SEA | *bla*$_{\text{CTX-M-55}}$, *strA, strB, sul2, dfrA1, tet(A), mph(A)* | S83L, D87G | S80I | ERS6575238 |
| 201703128 | X5 | 3.6.1.1 | CipR | *bla*$_{\text{CTX-M-15}}$, *aadA5, sul1, dfrA1, dfrA17, qnrS1, mph(A)* | S83L, D87G | S80I | ERS6575844 |
| 201703316 | X5 | 3.6.1.1 | CipR | *bla*$_{\text{CTX-M-15}}$, *aadA5, sul1, dfrA1, dfrA17, qnrS1, mph(A)* | S83L, D87G | S80I | ERS6575847 |
| 201908033 | X6 | 3.6.1.1 | CipR | *bla*$_{\text{CTX-M-15}}$, *strA, strB, sul2, dfrA1, tet(A), mph(A), erm(B)* | S83L, D87G | S80I | ERS6495503 |
| 201605743 | X7 | 3.6.1.1.2 | CipR.MSM5 | *bla*$_{\text{TEM-1B}}$, *bla*$_{\text{CTX-M-3}}$, *aadA5, sul1, dfrA1, dfrA17, mph(A), erm(B)* | S83L, D87G | S80I | ERS12446187 |
| 201807563 | X8 | 3.6.1.1.2 | CipR.MSM5 | *bla*$_{\text{TEM-1B}}$, *bla*$_{\text{CTX-M-15}}$, *aadA1, aadA5, sul1, sul2, dfrA1, dfrA17, tet(B), qnrS1, mph(A), erm(B)* | S83L, D87G | S80I | ERS6575531 |
| 201808736 | X8 | 3.6.1.1.2 | CipR.MSM5 | *bla*$_{\text{TEM-1B}}$, *bla*$_{\text{CTX-M-15}}$, *aadA1, aadA5, sul1, sul2, dfrA1, dfrA17, tet(B), qnrS1, mph(A), erm(B)* | S83L, D87G | S80I | ERS6492807 |
| 201809101 | X8 | 3.6.1.1.2 | CipR.MSM5 | *bla*$_{\text{TEM-1B}}$, *bla*$_{\text{CTX-M-15}}$, *aadA1, aadA5, sul1, sul2, dfrA1, dfrA17, tet(B), qnrS1, mph(A), erm(B)* | S83L, D87G | S80I | ERS6492833 |
| 201810383 | X8 | 3.6.1.1.2 | CipR.MSM5 | *bla*$_{\text{TEM-1B}}$, *bla*$_{\text{CTX-M-15}}$, *aadA1, aadA5, sul1, sul2, dfrA1, dfrA17, tet(B), qnrS1, mph(A), erm(B)* | S83L, D87G | S80I | ERS6494748 |
| 201810803 | X8 | 3.6.1.1.2 | CipR.MSM5 | *bla*$_{\text{TEM-1B}}$, *bla*$_{\text{CTX-M-15}}$, *aadA1, aadA5, sul1, sul2, dfrA1, dfrA17, tet(B), qnrS1, mph(A), erm(B)* | S83L, D87G | S80I | ERS6494788 |
| 201900053 | X8 | 3.6.1.1.2 | CipR.MSM5 | *bla*$_{\text{TEM-1B}}$, *bla*$_{\text{CTX-M-15}}$, *aadA1, aadA5, sul1, sul2, dfrA1, dfrA17, tet(B), qnrS1, mph(A), erm(B)* | S83L, D87G | S80I | ERS6575939 |
| 201904981 | X8 | 3.6.1.1.2 | CipR.MSM5 | *bla*$_{\text{TEM-1B}}$, *bla*$_{\text{CTX-M-15}}$, *aadA1, aadA5, sul1, sul2, dfrA1, dfrA17, tet(B), qnrS1, mph(A), erm(B)* | S83L, D87G | S80I | ERS6495091 |
| 201909147 | X8 | 3.6.1.1.2 | CipR.MSM5 | *bla*$_{\text{TEM-1B}}$, *bla*$_{\text{CTX-M-15}}$, *aadA1, aadA5, sul1, sul2, dfrA1, dfrA17, tet(B), qnrS1, mph(A), erm(B)* | S83L, D87G | S80I | ERS6495579 |
| 201910769 | X8 | 3.6.1.1.2 | CipR.MSM5 | *bla*$_{\text{TEM-1B}}$, *bla*$_{\text{CTX-M-15}}$, *aadA1, aadA5, sul1, sul2, dfrA1, dfrA17, tet(B), qnrS1, mph(A), erm(B)* | S83L, D87G | S80I | ERS6575039 |
| 201910803 | X8 | 3.6.1.1.2 | CipR.MSM5 | *bla*$_{\text{TEM-1B}}$, *bla*$_{\text{CTX-M-15}}$, *aadA1, aadA5, sul1, sul2, dfrA1, dfrA17, tet(B), qnrS1, mph(A), erm(B)* | S83L, D87G | S80I | ERS6575045 |
| 202000267 | X9 | 3.6.1.1.2 | CipR.MSM5 | *bla*$_{\text{CTX-M-134}}$, *strA, strB, sul2, dfrA1, tet(A), mph(A)* | S83L, D87G | S80I | ERS6495836 |
| 202000322 | X9 | 3.6.1.1.2 | CipR.MSM5 | *bla*$_{\text{CTX-M-134}}$, *strA, strB, sul2, dfrA1, tet(A), mph(A)* | S83L, D87G | S80I | ERS6495841 |
| 202000562 | X9 | 3.6.1.1.2 | CipR.MSM5 | *bla*$_{\text{CTX-M-134}}$, *strA, strB, sul2, dfrA1, tet(A), mph(A)* | S83L, D87G | S80I | ERS6575389 |
| 202001062 | X9 | 3.6.1.1.2 | CipR.MSM5 | *bla*$_{\text{CTX-M-134}}$, *strA, strB, sul2, dfrA1, tet(A), mph(A)* | S83L, D87G | S80I | ERS6576215 |
| 202001063 | X9 | 3.6.1.1.2 | CipR.MSM5 | *bla*$_{\text{CTX-M-134}}$, *strA, strB, sul2, dfrA1, tet(A), mph(A)* | S83L, D87G | S80I | ERS6576216 |
| 202001623 | X9 | 3.6.1.1.2 | CipR.MSM5 | *bla*$_{\text{CTX-M-134}}$, *strA, strB, sul2, dfrA1, tet(A), mph(A)* | S83L, D87G | S80I | ERS6576007 |
| 202006637 | X10 | 3.6.1.1.2 | CipR.MSM5 | *bla*$_{\text{CTX-M-27}}$, *strA, strB, aadA5, sul1, sul2, dfrA1, dfrA17, tet(A), qnrB19, mph(A), erm(B)* | S83L, D87G | S80I | ERS6576310 |
| 202007856 | X10 | 3.6.1.1.2 | CipR.MSM5 | *bla*$_{\text{CTX-M-27}}$, *strA, strB, aadA5, sul1, sul2, dfrA1, dfrA17, tet(A), qnrB19, mph(A), erm(B)* | S83L, D87G | S80I | ERS6576333 |
| 202008118 | X10 | 3.6.1.1.2 | CipR.MSM5 | *bla*$_{\text{CTX-M-27}}$, *strA, strB, aadA5, sul1, sul2, dfrA1, dfrA17, tet(A), qnrB19, mph(A)* | S83L, D87G | S80I | ERS12446247 |
| 202008158 | X10 | 3.6.1.1.2 | CipR.MSM5 | *bla*$_{\text{CTX-M-27}}$, *strA, strB, aadA5, sul1, sul2, dfrA1, dfrA17, tet(A), qnrB19, mph(A), erm(B)* | S83L, D87G | S80I | ERS6576346 |
| 202008564 | X10 | 3.6.1.1.2 | CipR.MSM5 | *bla*$_{\text{CTX-M-27}}$, *strA, strB, aadA5, sul1, sul2, dfrA1, dfrA17, tet(A), qnrB19, mph(A), erm(B)* | S83L, D87G | S80I | ERS6576362 |
| 202008707 | X10 | 3.6.1.1.2 | CipR.MSM5 | *bla*$_{\text{CTX-M-27}}$, *strA, strB, aadA5, sul1, sul2, dfrA1, dfrA17, tet(A), qnrB19, mph(A)* | S83L, D87G | S80I | ERS6578764 |
| 202100373 | X10 | 3.6.1.1.2 | CipR.MSM5 | *bla*$_{\text{CTX-M-27}}$, *strA, strB, aadA5, sul1, sul2, dfrA1, dfrA17, tet(A), qnrB19, mph(A)* | S83L, D87G | S80I | ERS12156949 |
| 202100420 | X10 | 3.6.1.1.2 | CipR.MSM5 | *bla*$_{\text{CTX-M-27}}$, *strA, strB, aadA5, sul1, sul2, dfrA1, dfrA17, tet(A), qnrB19, mph(A)* | S83L, D87G | S80I | ERS12156952 |
| 202100759 | X10 | 3.6.1.1.2 | CipR.MSM5 | *bla*$_{\text{CTX-M-27}}$, *strA, strB, aadA5, sul1, sul2, dfrA1, dfrA17, tet(A), qnrB19, mph(A), erm(B)* | S83L, D87G | S80I | ERS12156964 |
| 202101412 | X10 | 3.6.1.1.2 | CipR.MSM5 | *bla*$_{\text{CTX-M-27}}$, *strA, strB, aadA5, sul1, sul2, dfrA1, dfrA17, tet(A), qnrB19, mph(A), erm(B)* | S83L, D87G | S80I | ERS12156980 |
| 202101541 | X10 | 3.6.1.1.2 | CipR.MSM5 | *bla*$_{\text{CTX-M-27}}$, *aadA5, sul1, dfrA1, dfrA17, qnrB19, mph(A), erm(B)* | S83L, D87G | S80I | ERS12156987 |
| 202101624 | X10 | 3.6.1.1.2 | CipR.MSM5 | *bla*$_{\text{CTX-M-27}}$, *strA, strB, aadA5, sul1, sul2, dfrA1, dfrA17, tet(A), qnrB19, mph(A), erm(B)* | S83L, D87G | S80I | ERS12156996 |
| 202101651 | X10 | 3.6.1.1.2 | CipR.MSM5 | *bla*$_{\text{CTX-M-27}}$, *strA, strB, aadA5, sul1, sul2, dfrA1, dfrA17, tet(A), qnrB19, mph(A), erm(B)* | S83L, D87G | S80I | ERS12156997 |
| 202101727 | X10 | 3.6.1.1.2 | CipR.MSM5 | *bla*$_{\text{CTX-M-27}}$, *strA, strB, aadA5, sul1, sul2, dfrA1, dfrA17, tet(A), qnrB19, mph(A), erm(B)* | S83L, D87G | S80I | ERS12157001 |
| 202101879 | X10 | 3.6.1.1.2 | CipR.MSM5 | *bla*$_{\text{CTX-M-27}}$, *strA, strB, aadA5, sul1, sul2, dfrA1, dfrA17, tet(A), qnrB19, mph(A), erm(B)* | S83L, D87G | S80I | ERS12157007 |
| 202101885 | X10 | 3.6.1.1.2 | CipR.MSM5 | *bla*$_{\text{CTX-M-27}}$, *strA, strB, aadA5, sul1, sul2, dfrA1, dfrA17, tet(A), qnrB19, mph(A)* | S83L, D87G | S80I | ERS12157008 |
| 202101930 | X10 | 3.6.1.1.2 | CipR.MSM5 | *bla*$_{\text{CTX-M-27}}$, *strA, strB, aadA5, sul1, sul2, dfrA1, dfrA17, tet(A), qnrB19, mph(A), erm(B)* | S83L, D87G | S80I | ERS12157011 |
| 202102119 | X10 | 3.6.1.1.2 | CipR.MSM5 | *bla*$_{\text{CTX-M-27}}$, *strA, strB, aadA5, sul1, sul2, dfrA1, dfrA17, tet(A), qnrB19, mph(A), erm(B)* | S83L, D87G | S80I | ERS12157014 |
| 202102200 | X10 | 3.6.1.1.2 | CipR.MSM5 | *bla*$_{\text{CTX-M-27}}$, *strA, strB, aadA5, sul1, sul2, dfrA1, dfrA17, tet(A), qnrB19, mph(A), erm(B)* | S83L, D87G | S80I | ERS12446248 |
| 202102776 | X10 | 3.6.1.1.2 | CipR.MSM5 | *bla*$_{\text{CTX-M-27}}$, *strA, strB, aadA5, sul1, sul2, dfrA1, dfrA17, tet(A), qnrB19, mph(A), erm(B)* | S83L, D87G | S80I | ERS12157026 |
| 202102820 | X10 | 3.6.1.1.2 | CipR.MSM5 | *bla*$_{\text{CTX-M-27}}$, *strA, strB, aadA5, sul1, sul2, dfrA1, dfrA17, tet(A), qnrB19, mph(A), erm(B)* | S83L, D87G | S80I | ERS12157027 |
| 202102980 | X10 | 3.6.1.1.2 | CipR.MSM5 | *bla*$_{\text{CTX-M-27}}$, *strA, strB, aadA5, sul1, sul2, dfrA1, dfrA17, tet(A), qnrB19, mph(A)* | S83L, D87G | S80I | ERS12157031 |

| | | | | | | | |
|---|---|---|---|---|---|---|---|
| 202102993 | X10 | 3.6.1.1.2 | CipR.MSM5 | *bla*$_{CTX-M-27}$, *strA, strB, aadA5, sul1, sul2, dfrA1, dfrA17, tet(A), qnrB19, mph(A), erm(B)* | S83L, D87G | S80I | ERS12157032 |
| 202103005 | X10 | 3.6.1.1.2 | CipR.MSM5 | *bla*$_{CTX-M-27}$, *strA, strB, sul2, dfrA1, tet(A), qnrB19, erm(B)* | S83L, D87G | S80I | ERS12157033 |
| 202103026 | X10 | 3.6.1.1.2 | CipR.MSM5 | *bla*$_{CTX-M-27}$, *strA, strB, aadA5, sul1, sul2, dfrA1, dfrA17, tet(A), qnrB19, mph(A), erm(B)* | S83L, D87G | S80I | ERS12157034 |
| 202103166 | X10 | 3.6.1.1.2 | CipR.MSM5 | *bla*$_{CTX-M-27}$, *strA, strB, aadA5, sul1, sul2, dfrA1, dfrA17, tet(A), qnrB19, mph(A), erm(B)* | S83L, D87G | S80I | ERS12157040 |
| 202103248 | X10 | 3.6.1.1.2 | CipR.MSM5 | *bla*$_{CTX-M-27}$, *strA, strB, aadA5, sul1, sul2, dfrA1, dfrA17, tet(A), qnrB19, mph(A), erm(B)* | S83L, D87G | S80I | ERS12157044 |
| 202103290 | X10 | 3.6.1.1.2 | CipR.MSM5 | *bla*$_{CTX-M-27}$, *strA, strB, sul2, dfrA1, tet(A), qnrB19, erm(B)* | S83L, D87G | S80I | ERS12157048 |
| 202103342 | X10 | 3.6.1.1.2 | CipR.MSM5 | *bla*$_{CTX-M-27}$, *strA, strB, aadA5, sul1, sul2, dfrA1, dfrA17, tet(A), qnrB19, mph(A)* | S83L, D87G | S80I | ERS12157051 |
| 202103419 | X10 | 3.6.1.1.2 | CipR.MSM5 | *bla*$_{CTX-M-27}$, *strA, strB, sul2, dfrA1, tet(A), qnrB19, erm(B)* | S83L, D87G | S80I | ERS12157053 |
| 202103424 | X10 | 3.6.1.1.2 | CipR.MSM5 | *bla*$_{CTX-M-27}$, *strA, strB, sul2, dfrA1, tet(A), qnrB19, erm(B)* | S83L, D87G | S80I | ERS12157055 |
| 202103567 | X10 | 3.6.1.1.2 | CipR.MSM5 | *bla*$_{CTX-M-27}$, *strA, strB, aadA5, sul1, sul2, dfrA1, dfrA17, tet(A), qnrB19, mph(A), erm(B)* | S83L, D87G | S80I | ERS12157058 |
| 202103754 | X10 | 3.6.1.1.2 | CipR.MSM5 | *bla*$_{CTX-M-27}$, *strA, strB, aadA5, sul1, sul2, dfrA1, dfrA17, tet(A), qnrB19, mph(A), erm(B)* | S83L, D87G | S80I | ERS12157060 |
| 202103844 | X10 | 3.6.1.1.2 | CipR.MSM5 | *bla*$_{CTX-M-27}$, *strA, strB, aadA5, sul1, sul2, dfrA1, dfrA17, tet(A), qnrB19, mph(A), erm(B)* | S83L, D87G | S80I | ERS12157061 |
| 202103880 | X10 | 3.6.1.1.2 | CipR.MSM5 | *bla*$_{CTX-M-27}$, *strA, strB, sul2, dfrA1, tet(A), qnrB19, erm(B)* | S83L, D87G | S80I | ERS12157062 |
| 202103930 | X10 | 3.6.1.1.2 | CipR.MSM5 | *bla*$_{CTX-M-27}$, *strA, strB, aadA5, sul1, sul2, dfrA1, dfrA17, tet(A), qnrB19, mph(A), erm(B)* | S83L, D87G | S80I | ERS12157064 |
| 202104060 | X10 | 3.6.1.1.2 | CipR.MSM5 | *bla*$_{CTX-M-27}$, *strA, strB, aadA5, sul1, sul2, dfrA1, dfrA17, tet(A), qnrB19, mph(A), erm(B)* | S83L, D87G | S80I | ERS12157066 |
| 202104078 | X10 | 3.6.1.1.2 | CipR.MSM5 | *bla*$_{CTX-M-27}$, *strA, strB, sul2, dfrA1, tet(A), qnrB19, erm(B)* | S83L, D87G | S80I | ERS12157067 |
| 202104261 | X10 | 3.6.1.1.2 | CipR.MSM5 | *bla*$_{CTX-M-27}$, *strA, strB, aadA5, sul1, sul2, dfrA1, dfrA17, tet(A), qnrB19, mph(A), erm(B)* | S83L, D87G | S80I | ERS12157068 |
| 202104281 | X10 | 3.6.1.1.2 | CipR.MSM5 | *bla*$_{CTX-M-27}$, *strA, strB, aadA5, sul1, sul2, dfrA1, dfrA17, tet(A), qnrB19, mph(A), erm(B)* | S83L, D87G | S80I | ERS12157069 |
| 202104371 | X10 | 3.6.1.1.2 | CipR.MSM5 | *bla*$_{CTX-M-27}$, *strA, strB, aadA5, sul1, sul2, dfrA1, dfrA17, tet(A), qnrB19, mph(A), erm(B)* | S83L, D87G | S80I | ERS12157071 |
| 202104519 | X10 | 3.6.1.1.2 | CipR.MSM5 | *bla*$_{CTX-M-27}$, *strA, strB, aadA5, sul1, sul2, dfrA1, dfrA17, tet(A), qnrB19, mph(A), erm(B)* | S83L, D87G | S80I | ERS12157072 |
| 202104685 | X10 | 3.6.1.1.2 | CipR.MSM5 | *bla*$_{CTX-M-27}$, *strA, strB, aadA5, sul1, sul2, dfrA1, dfrA17, tet(A), qnrB19, mph(A), erm(B)* | S83L, D87G | S80I | ERS12157074 |
| 202104831 | X10 | 3.6.1.1.2 | CipR.MSM5 | *bla*$_{CTX-M-27}$, *strA, strB, sul1, sul2, dfrA1, dfrA5, tet(A), qnrB19, qnrS1, mph(A), erm(B)* | S83L, D87G | S80I | ERS12157076 |
| 202105132 | X10 | 3.6.1.1.2 | CipR.MSM5 | *bla*$_{CTX-M-27}$, *strA, strB, aadA5, sul1, sul2, dfrA1, dfrA17, tet(A), qnrB19, mph(A)* | S83L, D87G | S80I | ERS12157079 |
| 202105166 | X10 | 3.6.1.1.2 | CipR.MSM5 | *bla*$_{CTX-M-27}$, *strA, strB, aadA5, sul1, sul2, dfrA1, dfrA17, tet(A), qnrB19, mph(A), erm(B)* | S83L, D87G | S80I | ERS12157080 |
| 202105284 | X10 | 3.6.1.1.2 | CipR.MSM5 | *bla*$_{CTX-M-27}$, *strA, strB, sul2, dfrA1, tet(A), qnrB19, erm(B)* | S83L, D87G | S80I | ERS12157083 |
| 202105285 | X10 | 3.6.1.1.2 | CipR.MSM5 | *bla*$_{CTX-M-27}$, *strA, strB, sul2, dfrA1, tet(A), qnrB19, erm(B)* | S83L, D87G | S80I | ERS12157084 |
| 202105373 | X10 | 3.6.1.1.2 | CipR.MSM5 | *bla*$_{CTX-M-27}$, *strA, strB, aadA5, sul1, sul2, dfrA1, dfrA17, tet(A), qnrB19, mph(A), erm(B)* | S83L, D87G | S80I | ERS12157087 |
| 202105408 | X10 | 3.6.1.1.2 | CipR.MSM5 | *bla*$_{CTX-M-27}$, *strA, strB, aadA5, sul1, sul2, dfrA1, dfrA17, tet(A), qnrB19, mph(A), erm(B)* | S83L, D87G | S80I | ERS12157089 |
| 202105440 | X10 | 3.6.1.1.2 | CipR.MSM5 | *bla*$_{CTX-M-27}$, *strA, strB, aadA5, sul1, sul2, dfrA1, dfrA17, tet(A), qnrB19, mph(A), erm(B)* | S83L, D87G | S80I | ERS12157092 |
| 202105446 | X10 | 3.6.1.1.2 | CipR.MSM5 | *bla*$_{CTX-M-27}$, *dfrA1, qnrB19, erm(B)* | S83L, D87G | S80I | ERS12157093 |
| 202105523 | X10 | 3.6.1.1.2 | CipR.MSM5 | *bla*$_{CTX-M-27}$, *strA, strB, aadA5, sul1, sul2, dfrA1, dfrA17, tet(A), mph(A), erm(B)* | S83L, D87G | S80I | ERS12157094 |
| 202105574 | X10 | 3.6.1.1.2 | CipR.MSM5 | *bla*$_{CTX-M-27}$, *strA, strB, aadA5, sul1, sul2, dfrA1, dfrA17, tet(A), qnrB19, mph(A), erm(B)* | S83L, D87G | S80I | ERS12157095 |
| 202105953 | X10 | 3.6.1.1.2 | CipR.MSM5 | *bla*$_{CTX-M-27}$, *strA, strB, aadA5, sul1, sul2, dfrA1, dfrA17, tet(A), qnrB19, mph(A), erm(B)* | S83L, D87G | S80I | ERS12157110 |
| 202106029 | X10 | 3.6.1.1.2 | CipR.MSM5 | *bla*$_{CTX-M-27}$, *strA, strB, aadA5, sul1, sul2, dfrA1, dfrA17, tet(A), qnrB19, mph(A), erm(B)* | S83L, D87G | S80I | ERS12157114 |
| 202106219 | X10 | 3.6.1.1.2 | CipR.MSM5 | *bla*$_{CTX-M-27}$, *strA, strB, aadA5, sul1, sul2, dfrA1, dfrA17, tet(A), mph(A), erm(B)* | S83L, D87G | S80I | ERS12157121 |
| 202106247 | X10 | 3.6.1.1.2 | CipR.MSM5 | *bla*$_{CTX-M-27}$, *strA, strB, aadA5, sul1, sul2, dfrA1, dfrA17, tet(A), qnrB19, mph(A), erm(B)* | S83L, D87G | S80I | ERS12157122 |
| 202106455 | X10 | 3.6.1.1.2 | CipR.MSM5 | *bla*$_{CTX-M-27}$, *strA, strB, aadA5, sul1, sul2, dfrA1, dfrA17, tet(A), mph(A), erm(B)* | S83L, D87G | S80I | ERS12157133 |
| 202106629 | X10 | 3.6.1.1.2 | CipR.MSM5 | *bla*$_{CTX-M-27}$, *strA, strB, aadA5, sul1, sul2, dfrA1, dfrA17, tet(A), qnrB19, mph(A), erm(B)* | S83L, D87G | S80I | ERS12157142 |
| 202106664 | X10 | 3.6.1.1.2 | CipR.MSM5 | *bla*$_{CTX-M-27}$, *aadA5, sul1, dfrA1, dfrA17, qnrB19, mph(A), erm(B)* | S83L, D87G | S80I | ERS12157145 |
| 202106716 | X10 | 3.6.1.1.2 | CipR.MSM5 | *bla*$_{CTX-M-27}$, *strA, strB, aadA5, sul1, sul2, dfrA1, dfrA17, tet(A), qnrB19, mph(A), erm(B)* | S83L, D87G | S80I | ERS12157148 |
| 202106961 | X10 | 3.6.1.1.2 | CipR.MSM5 | *bla*$_{CTX-M-27}$, *strA, strB, aadA5, sul1, sul2, dfrA1, dfrA17, tet(A), qnrB19, mph(A), erm(B)* | S83L, D87G | S80I | ERS12157163 |
| 202107022 | X10 | 3.6.1.1.2 | CipR.MSM5 | *bla*$_{CTX-M-27}$, *strA, strB, aadA5, sul1, sul2, dfrA1, dfrA17, tet(A), qnrB19, mph(A), erm(B)* | S83L, D87G | S80I | ERS12157165 |
| 202107051 | X10 | 3.6.1.1.2 | CipR.MSM5 | *bla*$_{CTX-M-27}$, *strA, strB, aadA5, sul1, sul2, dfrA1, dfrA17, tet(A), qnrB19, mph(A), erm(B)* | S83L, D87G | S80I | ERS12157167 |

| | | | | | | | |
|---|---|---|---|---|---|---|---|
| 202107052 | X10 | 3.6.1.1.2 | CipR.MSM5 | $bla_{CTX-M-27}$, *strA, strB, aadA5, sul1, sul2, dfrA1, dfrA17, tet(A), qnrB19, mph(A), erm(B)* | S83L, D87G | S80I | ERS12157168 |
| 202107188 | X10 | 3.6.1.1.2 | CipR.MSM5 | $bla_{CTX-M-27}$, *aadA5, sul1, dfrA1, dfrA17, qnrB19, mph(A), erm(B)* | S83L, D87G | S80I | ERS12157174 |
| 202107323 | X10 | 3.6.1.1.2 | CipR.MSM5 | $bla_{CTX-M-27}$, *strA, strB, aadA5, sul1, sul2, dfrA1, dfrA17, tet(A), qnrB19, mph(A), erm(B)* | S83L, D87G | S80I | ERS12157179 |
| 202107395 | X10 | 3.6.1.1.2 | CipR.MSM5 | $bla_{CTX-M-27}$, *strA, strB, aadA5, sul1, sul2, dfrA1, dfrA17, tet(A), mph(A)* | S83L, D87G | S80I | ERS12157182 |
| 202107415 | X10 | 3.6.1.1.2 | CipR.MSM5 | $bla_{CTX-M-27}$, *strA, strB, aadA5, sul1, sul2, dfrA1, dfrA17, tet(A), mph(A), erm(B)* | S83L, D87G | S80I | ERS12157183 |
| 202107446 | X10 | 3.6.1.1.2 | CipR.MSM5 | $bla_{CTX-M-27}$, *strA, strB, aadA5, sul1, sul2, dfrA1, dfrA17, tet(A), qnrB19, mph(A), erm(B)* | S83L, D87G | S80I | ERS12157184 |
| 202107466 | X10 | 3.6.1.1.2 | CipR.MSM5 | $bla_{CTX-M-27}$, *strA, strB, aadA5, sul1, sul2, dfrA1, dfrA17, tet(A), mph(A), erm(B)* | S83L, D87G | S80I | ERS12157186 |
| 202107818 | X10 | 3.6.1.1.2 | CipR.MSM5 | $bla_{CTX-M-27}$, *strA, strB, aadA5, sul1, sul2, dfrA1, dfrA17, tet(A), mph(A), erm(B)* | S83L, D87G | S80I | ERS12157209 |
| 202108294 | X10 | 3.6.1.1.2 | CipR.MSM5 | $bla_{CTX-M-27}$, *strA, strB, aadA5, sul1, sul2, dfrA1, dfrA17, tet(A), qnrB19, mph(A), erm(B)* | S83L, D87G | S80I | ERS12446254 |
| 202108795 | X10 | 3.6.1.1.2 | CipR.MSM5 | $bla_{CTX-M-27}$, *strA, strB, aadA5, sul1, sul2, dfrA1, dfrA17, tet(A), mph(A), erm(B)* | S83L, D87G | S80I | ERS12446289 |
| 202108919 | X10 | 3.6.1.1.2 | CipR.MSM5 | $bla_{CTX-M-27}$, *strA, strB, aadA5, sul1, sul2, dfrA1, dfrA17, tet(A), mph(A), erm(B)* | S83L, D87G | S80I | ERS12446294 |
| 202108953 | X10 | 3.6.1.1.2 | CipR.MSM5 | $bla_{CTX-M-27}$, *strA, strB, aadA5, sul1, sul2, dfrA1, dfrA17, tet(A), qnrB19, mph(A), erm(B)* | S83L, D87G | S80I | ERS12446298 |
| 202109588 | X10 | 3.6.1.1.2 | CipR.MSM5 | $bla_{CTX-M-27}$, *strA, strB, aadA5, sul1, sul2, dfrA1, dfrA17, tet(A), mph(A), erm(B)* | S83L, D87G | S80I | ERS12446318 |
| 202109651 | X10 | 3.6.1.1.2 | CipR.MSM5 | $bla_{CTX-M-27}$, *strA, strB, aadA5, sul1, sul2, dfrA1, dfrA17, tet(A), qnrB19, mph(A), erm(B)* | S83L, D87G | S80I | ERS12446319 |
| 202109656 | X10 | 3.6.1.1.2 | CipR.MSM5 | $bla_{CTX-M-27}$, *strA, strB, aadA5, sul1, sul2, dfrA1, dfrA17, tet(A), qnrB19, mph(A), erm(B)* | S83L, D87G | S80I | ERS12446320 |
| 202109849 | X10 | 3.6.1.1.2 | CipR.MSM5 | $bla_{CTX-M-27}$, *aadA5, sul1, dfrA1, dfrA17, qnrB19, mph(A), erm(B)* | S83L, D87G | S80I | ERS12446324 |
| 202109909 | X10 | 3.6.1.1.2 | CipR.MSM5 | $bla_{CTX-M-27}$, *strA, strB, aadA5, sul1, sul2, dfrA1, dfrA17, tet(A), mph(A), erm(B)* | S83L, D87G | S80I | ERS12446330 |
| 202109960 | X10 | 3.6.1.1.2 | CipR.MSM5 | $bla_{CTX-M-27}$, *strA, strB, aadA5, sul1, sul2, dfrA1, dfrA17, tet(A), mph(A), erm(B)* | S83L, D87G | S80I | ERS12446335 |
| 202110146 | X10 | 3.6.1.1.2 | CipR.MSM5 | $bla_{CTX-M-27}$, *aadA5, sul1, dfrA1, dfrA17, qnrB19, mph(A), erm(B)* | S83L, D87G | S80I | ERS12446347 |
| 202110251 | X10 | 3.6.1.1.2 | CipR.MSM5 | $bla_{CTX-M-27}$, *aadA5, sul1, dfrA1, dfrA17, qnrB19, mph(A), erm(B)* | S83L, D87G | S80I | ERS12446356 |
| 202110403 | X10 | 3.6.1.1.2 | CipR.MSM5 | $bla_{CTX-M-27}$, *strA, strB, aadA5, sul1, sul2, dfrA1, dfrA17, tet(A), qnrB19, mph(A), erm(B)* | S83L, D87G | S80I | ERS12446360 |
| 202110415 | X10 | 3.6.1.1.2 | CipR.MSM5 | $bla_{CTX-M-27}$, *aadA5, sul1, dfrA1, dfrA17, qnrB19, mph(A), erm(B)* | S83L, D87G | S80I | ERS12446361 |
| 202110472 | X10 | 3.6.1.1.2 | CipR.MSM5 | $bla_{CTX-M-27}$, *strA, strB, aadA5, sul1, sul2, dfrA1, dfrA17, tet(A), qnrB19, mph(A), erm(B)* | S83L, D87G | S80I | ERS12446362 |
| 202110584 | X10 | 3.6.1.1.2 | CipR.MSM5 | $bla_{CTX-M-27}$, *aadA5, sul1, dfrA1, dfrA17, qnrB19, mph(A), erm(B)* | S83L, D87G | S80I | ERS12446365 |
| 202110641 | X10 | 3.6.1.1.2 | CipR.MSM5 | $bla_{CTX-M-27}$, *aadA5, sul1, dfrA1, dfrA17, qnrB19, mph(A), erm(B)* | S83L, D87G | S80I | ERS12446370 |
| 202110674 | X10 | 3.6.1.1.2 | CipR.MSM5 | $bla_{CTX-M-27}$, *aadA5, sul1, dfrA1, dfrA17, qnrB19, mph(A), erm(B)* | S83L, D87G | S80I | ERS12446373 |
| 202110841 | X10 | 3.6.1.1.2 | CipR.MSM5 | $bla_{CTX-M-27}$, *aadA5, sul1, dfrA1, dfrA17, qnrB19, mph(A), erm(B)* | S83L, D87G | S80I | ERS12446377 |
| 202110869 | X10 | 3.6.1.1.2 | CipR.MSM5 | $bla_{CTX-M-27}$, *aadA5, sul1, dfrA1, dfrA17, qnrB19, mph(A), erm(B)* | S83L, D87G | S80I | ERS12446379 |
| 202110877 | X10 | 3.6.1.1.2 | CipR.MSM5 | $bla_{CTX-M-27}$, *aadA5, sul1, dfrA1, dfrA17, qnrB19, mph(A), erm(B)* | S83L, D87G | S80I | ERS12446381 |
| 202110960 | X10 | 3.6.1.1.2 | CipR.MSM5 | $bla_{CTX-M-27}$, *strA, strB, aadA5, sul1, sul2, dfrA1, dfrA17, tet(A), qnrB19, mph(A), erm(B)* | S83L, D87G | S80I | ERS12446386 |
| 202110961 | X10 | 3.6.1.1.2 | CipR.MSM5 | $bla_{CTX-M-27}$, *strA, strB, aadA5, sul1, sul2, dfrA1, dfrA17, tet(A), qnrB19, mph(A), erm(B)* | S83L, D87G | S80I | ERS12446387 |
| 202111030 | X10 | 3.6.1.1.2 | CipR.MSM5 | $bla_{CTX-M-27}$, *strA, strB, aadA5, sul1, sul2, dfrA1, dfrA17, tet(A), qnrB19, mph(A), erm(B)* | S83L, D87G | S80I | ERS12446392 |
| 202111082 | X10 | 3.6.1.1.2 | CipR.MSM5 | $bla_{CTX-M-27}$, *strA, strB, aadA5, sul1, sul2, dfrA1, dfrA17, tet(A), qnrB19, mph(A), erm(B)* | S83L, D87G | S80I | ERS12446394 |
| 202111134 | X10 | 3.6.1.1.2 | CipR.MSM5 | $bla_{CTX-M-27}$, *strA, strB, aadA5, sul1, sul2, dfrA1, dfrA17, tet(A), mph(A), erm(B)* | S83L, D87G | S80I | ERS12446395 |
| 202111144 | X10 | 3.6.1.1.2 | CipR.MSM5 | $bla_{CTX-M-27}$, *strA, strB, sul2, dfrA1, tet(A), qnrB19, erm(B)* | S83L, D87G | S80I | ERS12446396 |
| 202111184 | X10 | 3.6.1.1.2 | CipR.MSM5 | $bla_{CTX-M-27}$, *aadA5, sul1, dfrA1, dfrA17, qnrB19, mph(A), erm(B)* | S83L, D87G | S80I | ERS12446399 |
| 202200143 | X10 | 3.6.1.1.2 | CipR.MSM5 | $bla_{CTX-M-27}$, *aadA5, sul1, dfrA1, dfrA17, qnrB19, mph(A), erm(B)* | S83L, D87G | S80I | ERS12446406 |
| 201908234 | X11 | 3.6.1.1.2 | CipR.MSM5 | $bla_{TEM-1B}$, $bla_{CTX-M-3}$, *strA, strB, aadA5, sul1, sul2, dfrA1, dfrA17, tet(A), mph(A), erm(B)* | S83L, D87G | S80I | ERS12446240 |
| 202106367 | X12 | 3.6.1 | CipR-Parent | $bla_{CTX-M-15}$, *strA, strB, sul2, dfrA1, tet(A), qnrS1, mph(A)* | S83L | - | ERS12157130 |
| 202108535 | X12 | 3.6.1 | CipR-Parent | $bla_{CTX-M-15}$, *strA, strB, sul2, dfrA1, tet(A), qnrB19, qnrS1, mph(A)* | S83L | - | ERS12446269 |
| 202102843 | X13 | 3.6.1.1.2 | CipR.MSM5 | $bla_{TEM-1B}$, $bla_{CTX-M-15}$, *strA, strB, aadA5, sul1, sul2, dfrA1, dfrA17, tet(A), qnrS1, mph(A), erm(B)* | S83L, D87G | S80I | ERS12157029 |

QRDR, quinolone resistance-determining region

**Supplementary Table 3.** Description of the AMR elements identified after long-read sequencing

| | | | QRDR mutations | | Chromosomal AMR genes[#] | AMR plasmids | | | | | |
|---|---|---|---|---|---|---|---|---|---|---|---|
| Genotype | XDR cluster | Isolate | *gyrA* | *parC* | | Name | Size in kb | Inc type[£] | PTU type[£] | AMR genes | GenBank accession nos. |
| **3.6.1_CipR-parent** | | | | | | | | | | | |
| | X12 | 202106367 | S83L | | *dfrA1* | p202106367-3 | 87,521 | FII | FE | $bla_{CTX-M-15}$, *mph(A)*, *qnrS1* | OP038295 |
| | | | | | | p202106367-6 | 8,379 | NT | NT | *strA*, *strB*, *sul2*, *tet(A)* | OP038296 |
| | | 202108535 | S83L | | *dfrA1* | p202108535-3 | 87,540 | FII | FE | $bla_{CTX-M-15}$, *mph(A)*, *qnrS1* | OP038298 |
| | | | | | | p202108535-7 | 8,379 | NT | NT | *strA*, *strB*, *sul2*, *tet(A)* | OP038299 |
| | | | | | | p202108535-13 | 2,699 | Col(pH) | E62 | *qnrB19* | OP038297 |
| **3.6.1.1_CipR** | | | | | | | | | | | |
| | X1 | 201701093 | S83L | S80I | $bla_{CTX-M-15}$, *dfrA1* | p201701093-2 | 102,387 | B/O/K/Z | B/O/K/Z | *sul1*, *dfrA5*, *mph(A)*, *erm(B)*, *qnrS13* | OP038271 |
| | X5 | 201703316 | S83L, D87G | S80I | *dfrA1* | p201703316-2 | 82,941 | FII | FE | $bla_{CTX-M-15}$, *aadA5*, *sul1*, *dfrA17*, *mph(A)*, *qnrS1* | OP038274 |
| | X6 | 201908033 | S83L, D87G | S80I | $bla_{CTX-M-15}$, *dfrA1* | p201908033-3 | 73,493 | FII | FE | *mph(A)*, *erm(B)* | OP038279 |
| | | | | | | p201908033-4 | 8,401 | NT | E63 | *strA*, *strB*, *sul2*, *tet(A)* | OP038280 |
| **3.6.1.1.1_CipR.SEA** | | | | | | | | | | | |
| | X2 | 201903956 | S83L, D87G | S80I | *dfrA1* | p201903956-2 | 78,123 | FII | FE | $bla_{CTX-M-27}$, *aadA5*, *sul1*, *dfrA17*, *mph(A)* | OP038278 |
| | | | | | | p201903956-4 | 8,401 | NT | E63 | *strA*, *strB*, *sul2*, *tet(A)* | OP038303 |
| | X3 | 201502874 | S83L, D87G | S80I | *dfrA1* | p201502874-3 | 79,375 | FII | FE | $bla_{CTX-M-27}$, *aadA5*, *sul1*, *dfrA17*, *mph(A)* | OP038267 |
| | | | | | | p201502874-5 | 8,401 | NT | NT | *strA*, *strB*, *sul2*, *tet(A)* | OP038268 |
| | X4 | 201603898 | S83L, D87G | S80I | *dfrA1* | p201603898-2 | 102,642 | B/O/K/Z | B/O/K/Z | $bla_{CTX-M-55}$, *aac(3)-IIa*, *mph(A)* | OP038269 |
| | | | | | | p201603898-4 | 8,401 | NT | NT | *strA*, *strB*, *sul2*, *tet(A)* | OP038270 |
| | | 201702782 | S83L, D87G | S80I | *dfrA1* | p201702782-2 | 97,029 | B/O/K/Z | B/O/K/Z | $bla_{CTX-M-55}$, *aac(3)-IIa*, *mph(A)* | OP038272 |
| | | | | | | p201702782-3 | 8,401 | NT | E63 | *strA*, *strB*, *sul2*, *tet(A)* | OP038273 |
| **3.6.1.1.2_CipR.MSM5** | | | | | | | | | | | |
| | X7 | 201605743 | S83L, D87G | S80I | *dfrA1* | p201605743-3 | 84,796 | I1 | I1 | $bla_{CTX-M-3}$ | OP038300 |
| | | | | | | p201605743-4 | 80,127 | FII | FE | $bla_{TEM-1B}$, *aadA5*, *sul1*, *dfrA17*, *mph(A)*, *erm(B)* | OP038301 |
| | X8 | 201809101 | S83L, D87G | S80I | *dfrA1* | p201809101-4 | 106,936 | I1 | I1 | $bla_{CTX-M-15}$, *aadA1*, *sul2*, *dfrA1*, *qnrS1* | OP038275 |
| | | | | | | p201809101-5* | 88,949 | B/O/K/Z | B/O/K/Z | *sul2*, *tet(B)* | OP038276 |
| | | | | | | p201809101-6 | 80,134 | FII | FE | $bla_{TEM-1B}$, *aadA5*, *sul1*, *dfrA17*, *mph(A)*, *erm(B)* | OP038277 |
| | X9 | 202000562 | S83L, D87G | S80I | *dfrA1* | p202000562-4 | 76,702 | FII | FE | $bla_{CTX-M-134}$, *strA*, *sul2*, *mph(A)* | OP038284 |
| | | | | | | p202000562-5 | 8,379 | NT | NT | *strA*, *strB*, *sul2*, *tet(A)* | OP038285 |
| | X10 | 202008118 | S83L, D87G | S80I | *dfrA1* | p202008118-4 | 78,102 | FII | FE | $bla_{CTX-M-27}$, *aadA5*, *sul1*, *dfrA17*, *mph(A)* | OP038287 |
| | | | | | | p202008118-7 | 8,390 | NT | NT | *strA*, *strB*, *sul2*, *tet(A)* | OP038288 |
| | | | | | | p202008118-16 | 2,579 | Col(pH) | E76 | *qnrB19* | OP038286 |

| | | | | | | | | | | |
|---|---|---|---|---|---|---|---|---|---|---|
| | 202008564 | S83L, D87G | S80I | *dfrA1* | p202008564-6 | 83,397 | FII | FE | *bla*$_{\text{CTX-M-27}}$, *aadA5*, *sul1*, *dfrA17*, *mph(A)*, *erm(B)* | OP038290 |
| | | | | | p202008564-7 | 8,390 | NT | NT | *strA*, *strB*, *sul2*, *tet(A)* | OP038291 |
| | | | | | p202008564-21 | 2,579 | Col(pH) | E76 | *qnrB19* | OP038289 |
| X11 | 201908234 | S83L, D87G | S80I | *dfrA1* | p201908234-4 | 86,101 | I1 | I1 | *bla*$_{\text{CTX-M-3}}$ | OP038281 |
| | | | | | p201908234-5 | 82,851 | FII | FE | *bla*$_{\text{TEM-1B}}$, *aadA5*, *sul1*, *dfrA17*, *mph(A)*, *erm(B)* | OP038282 |
| | | | | | p201908234-6 | 8,379 | NT | E63 | *strA*, *strB*, *sul2*, *tet(A)* | OP038283 |
| X13 | 202102843 | S83L, D87G | S80I | *dfrA1* | p202102843-3 | 89,779 | I1 | I1 | *bla*$_{\text{CTX-M-15}}$, *qnrS1* | OP038292 |
| | | | | | p202102843-4 | 80,145 | FII | FE | *bla*$_{\text{TEM-1B}}$, *aadA5*, *sul1*, *dfrA17*, *mph(A)*, *erm(B)* | OP038293 |
| | | | | | p202102843-8 | 8,401 | NT | E63 | *strA*, *strB*, *sul2*, *tet(A)* | OP038294 |

[#], The *dfrA1* gene is a class 2 integron gene cassette present in a chromosomally inserted Tn*7* (downstream from the bacterial gene *glmS*); *, non-circularized; [£], NT, non typable, Col(pH) means Col(pHAD28).

**Supplementary Table 4.** List of the 3,140 *S. sonnei* genomic sequences from isolates and historical strains of the FNRC-ESS included in the phylogenetic tree

| Isolate | Alternative name | Year | Country of isolation | Metropole/Overseas | Travel | Accession no. |
|---|---|---|---|---|---|---|
| 54210 | | 1943 | Sweden | Not Applicable | Unknown | ERR025732 |
| 54228 | | 1947 | Sweden | Not Applicable | Unknown | ERR025724 |
| 6-58 | | 1958 | France | Metropole | Unknown | ERR025734 |
| 2-59 | | 1959 | France | Metropole | Unknown | ERR025737 |
| 11-66 | | 1966 | France | Metropole | Unknown | ERR025743 |
| 11-73 | | 1973 | France | Metropole | Unknown | ERR025746 |
| 20-73 | | 1973 | France | Metropole | Unknown | ERR025749 |
| 2-73 | | 1973 | Senegal | Not Applicable | Unknown | ERR025750 |
| 25-74 | | 1974 | France | Metropole | Unknown | ERR025753 |
| 43-74 | | 1974 | France | Metropole | Unknown | ERR025754 |
| 44-74 | | 1974 | France | Metropole | Unknown | ERR025755 |
| 4-76 | | 1976 | France | Metropole | Unknown | ERR025756 |
| 88-83 | | 1983 | France | Metropole | Unknown | ERR025747 |
| 96-1420 | sh96-1420 | 1996 | France | Metropole | Unknown | ERR042788 |
| 96-2938 | sh96-2938 | 1996 | France | Metropole | Israel | ERR042789 |
| 96-5282 | sh96-5282 | 1996 | France | Metropole | Unknown | ERR042790 |
| 96-6494 | sh96-6494 | 1996 | France | Metropole | Unknown | ERR042791 |
| 98-10267 | | 1998 | France | Metropole | Madagascar | ERR025762 |
| 98-8743 | | 1998 | France | Metropole | French Guiana | ERR025758 |
| 98-9560 | | 1998 | France | Metropole | Madagascar | ERR025761 |
| 200005827 | sh00-5827 | 2000 | France | Metropole | Madagascar | ERR025765 |
| 200207265 | sh02-7265 | 2002 | France | Metropole | Unknown | ERR042775 |
| 200208933 | sh02-8933 | 2002 | France | Metropole | Unknown | ERR042776 |
| 200209633 | sh02-9633 | 2002 | France | Metropole | Unknown | ERR042777 |
| 200300995 | sh03-0995 | 2003 | France | Metropole | Unknown | ERR042778 |
| 200301223 | sh03-1223 | 2003 | France | Metropole | Unknown | ERR042779 |
| 200301382 | sh03-1382 | 2003 | France | Metropole | Israel | ERR025767 |
| 200301382 | sh03-1382 | 2003 | France | Metropole | Israel | ERR042794 |
| 200302222 | sh03-2222 | 2003 | France | Metropole | Cuba | ERR025768 |
| 200303567 | sh03-3567 | 2003 | France | Metropole | Unknown | ERR042780 |
| 200401191 | sh04-1191 | 2004 | France | Metropole | Tanzania | ERR024604 |
| 200503579 | | 2005 | France | Metropole | India | ERR1290040 |
| 200504771 | | 2005 | France | Metropole | India | ERR1290041 |
| 200506395 | | 2005 | France | Metropole | India | ERR1290042 |
| 200506437 | | 2005 | France | Metropole | India | ERR1290043 |
| 200506859 | | 2005 | France | Metropole | India | ERR1290044 |
| 200507122 | | 2005 | France | Metropole | India | ERR1290045 |
| 200507941 | | 2005 | France | Metropole | India | ERR1290046 |
| 200507958 | | 2005 | France | Metropole | None | ERS12446407 |
| 200508237 | | 2005 | France | Metropole | Unknown | ERS12445939 |
| 200505623 | sh05-5623 | 2005 | France | Metropole | Morocco | ERR024607 |
| 200600108 | sh06-0108 | 2005 | France | French Guiana | None | ERR024608 |
| 200600063 | | 2006 | France | Metropole | India | ERR1290047 |
| 200602347 | | 2006 | France | Metropole | Nepal | ERR1290048 |
| 200602456 | | 2006 | France | Metropole | Nepal | ERR1290049 |
| 200602762 | | 2006 | France | Metropole | Pakistan | ERR1290050 |
| 200605380 | | 2006 | France | Metropole | China | ERS12446408 |
| 200607712 | | 2006 | France | Metropole | India | ERR1290051 |
| 200609049 | | 2006 | France | Metropole | Pakistan | ERR1290052 |
| 200602542 | sh06-2542 | 2006 | France | Metropole | Burkina Faso | ERR024609 |
| 200605179 | sh06-5179 | 2006 | France | Metropole | Senegal | ERR024610 |
| 200605387 | sh06-5387 | 2006 | France | Metropole | Morocco | ERR024611 |
| 200605623 | sh06-5623 | 2006 | France | Metropole | Morocco | ERR024612 |
| 200606396 | sh06-6396 | 2006 | France | Metropole | Unknown | ERR024614 |
| 200606470 | sh06-6470 | 2006 | France | Metropole | Haiti | ERR024605 |
| 200700106 | | 2007 | France | Metropole | India | ERR1290104 |
| 200700426 | sh07-0426 | 2007 | France | Metropole | Unknown | ERR042782 |
| 200701018 | sh07-1018 | 2007 | France | Metropole | Unknown | ERR042783 |
| 200701244 | | 2007 | France | Metropole | Egypt | ERS12446409 |
| 200701608 | sh07-1608 | 2007 | France | Metropole | Unknown | ERR042784 |
| 200701748 | | 2007 | France | Metropole | India | ERR1290105 |
| 200701910 | sh07-1910 | 2007 | France | Metropole | Unknown | ERR042785 |
| 200702004 | sh07-2004 | 2007 | France | Metropole | Unknown | ERR042786 |
| 200702224 | sh07-2224 | 2007 | France | Metropole | Unknown | ERR042787 |
| 200702523 | | 2007 | France | Metropole | India | ERR1290106 |
| 200703781 | | 2007 | France | Metropole | India | ERS12445940 |
| 200705022 | | 2007 | France | Metropole | Pakistan | ERR1290107 |
| 200705156 | | 2007 | France | Metropole | Nepal | ERR1290108 |
| 200705535 | | 2007 | France | Metropole | Nepal | ERS12446410 |
| 200705552 | | 2007 | France | Metropole | Nepal | ERR1290109 |
| 200705956 | | 2007 | France | Metropole | None | ERS12446411 |
| 200706723 | | 2007 | France | Metropole | India | ERR1290110 |
| 200704369 | Sh07-4369 | 2007 | France | Metropole | Unknown | ERR024606 |
| 200801815 | | 2008 | France | Metropole | India | ERR1290111 |
| 200802680 | | 2008 | France | Metropole | India | ERR1290112 |
| 200806701 | | 2008 | France | Metropole | India | ERR1290113 |
| 200807227 | | 2008 | France | Metropole | India | ERS12446412 |
| 200807710 | | 2008 | France | Metropole | India | ERR1290114 |
| 200807785 | | 2008 | France | Metropole | Switzerland | ERS12446413 |
| 200807891 | | 2008 | France | Metropole | None | ERS12446414 |
| 200808768 | | 2008 | France | Metropole | India | ERR1290115 |
| 200810423 | | 2008 | France | Metropole | India | ERR1290116 |
| 200901360 | | 2009 | France | Metropole | Canary Islands | ERS12446415 |
| 200901709 | | 2009 | France | Metropole | Tunisia | ERS12446416 |
| 200902265 | | 2009 | France | Metropole | India | ERS12445941 |
| 200902471 | | 2009 | France | Metropole | India | ERR1290072 |
| 200902613 | | 2009 | France | Metropole | India | ERR1290073 |
| 200903618 | | 2009 | France | Metropole | India | ERR1290074 |
| 200904563 | | 2009 | France | Metropole | India | ERR1290075 |
| 200905271 | | 2009 | France | Metropole | India | ERR1290076 |
| 200905450 | | 2009 | France | Metropole | India | ERR1290077 |
| 200905584 | | 2009 | France | Metropole | India | ERR1290078 |

| | | | | | |
|---|---|---|---|---|---|
| 200905610 | 2009 | France | Metropole | India | ERR1290079 |
| 200905611 | 2009 | France | Metropole | India | ERS12445942 |
| 200905799 | 2009 | France | Metropole | India | ERR1290080 |
| 200905819 | 2009 | France | Metropole | India | ERR1290081 |
| 200906003 | 2009 | France | Metropole | India | ERR1290082 |
| 200906032 | 2009 | France | Metropole | India | ERS12445943 |
| 200906336 | 2009 | France | Metropole | India | ERR1290083 |
| 200906354 | 2009 | France | Metropole | Unknown | ERS12445944 |
| 200906870 | 2009 | France | Metropole | India | ERS12445945 |
| 200907471 | 2009 | France | Metropole | None | ERS12445946 |
| 200907727 | 2009 | France | Metropole | India | ERR1290084 |
| 200908210 | 2009 | France | Metropole | India | ERR1290085 |
| 200908973 | 2009 | France | Metropole | None | ERR1516064 |
| 200909097 | 2009 | France | Metropole | Unknown | ERS12445947 |
| 201000148 | 2010 | France | Metropole | India | ERR1290086 |
| 201000294 | 2010 | France | Metropole | India | ERR1290087 |
| 201000844 | 2010 | France | Metropole | Nepal | ERS12445948 |
| 201001217 | 2010 | France | Metropole | None | ERR1516065 |
| 201001454 | 2010 | France | Metropole | India | ERR1290088 |
| 201001899 | 2010 | France | Metropole | Unknown | ERS12445949 |
| 201002095 | 2010 | France | Metropole | India | ERR1290089 |
| 201002215 | 2010 | France | Metropole | None | ERR1516066 |
| 201002275 | 2010 | France | Metropole | Ecuador | ERR1516067 |
| 201002412 | 2010 | France | Metropole | India | ERR1290090 |
| 201002549 | 2010 | France | Metropole | Unknown | ERS12446417 |
| 201002692 | 2010 | France | Metropole | Unknown | ERS12445950 |
| 201002795 | 2010 | France | Metropole | India | ERR1290091 |
| 201002971 | 2010 | France | Metropole | Maldives | ERS12446418 |
| 201003014 | 2010 | France | Metropole | India | ERS12445951 |
| 201003098 | 2010 | France | Metropole | India | ERR1290092 |
| 201003858 | 2010 | France | Metropole | Unknown | ERS12445952 |
| 201003859 | 2010 | France | Metropole | None | ERR1516068 |
| 201004217 | 2010 | France | Metropole | Unknown | ERS12446419 |
| 201004269 | 2010 | France | Metropole | Unknown | ERS12445953 |
| 201004369 | 2010 | France | Metropole | Unknown | ERS12445954 |
| 201004450 | 2010 | France | Metropole | India | ERR1290093 |
| 201004619 | 2010 | France | Metropole | India | ERS12445955 |
| 201004745 | 2010 | France | Metropole | Unknown | ERS12446420 |
| 201004752 | 2010 | France | Metropole | India | ERR1516069 |
| 201004767 | 2010 | France | Metropole | None | ERR1516070 |
| 201004778 | 2010 | France | Metropole | Unknown | ERS12445956 |
| 201005110 | 2010 | France | Metropole | India | ERR1290095 |
| 201005130 | 2010 | France | Metropole | India | ERR1290096 |
| 201005165 | 2010 | France | Metropole | India | ERR1290097 |
| 201005309 | 2010 | France | Metropole | None | ERR1516071 |
| 201005369 | 2010 | France | Metropole | Egypt | ERR1290098 |
| 201005497 | 2010 | France | Metropole | Unknown | ERS12445957 |
| 201005526 | 2010 | France | Metropole | India | ERR1290099 |
| 201005586 | 2010 | France | Metropole | Algeria | ERR1516072 |
| 201005808 | 2010 | France | Metropole | None | ERR1516073 |
| 201005926 | 2010 | France | Metropole | None | ERR1516074 |
| 201005979 | 2010 | France | Metropole | Egypt | ERR1516075 |
| 201006110 | 2010 | France | Metropole | Nepal | ERR1290100 |
| 201006730 | 2010 | France | Metropole | Pakistan | ERS12446421 |
| 201006903 | 2010 | France | Metropole | India | ERS12445958 |
| 201007497 | 2010 | France | Metropole | Vietnam | ERS12446422 |
| 201007778 | 2010 | France | Metropole | Unknown | ERS12445959 |
| 201007817 | 2010 | France | Metropole | India | ERR1290101 |
| 201007882 | 2010 | France | Metropole | None | ERR1516076 |
| 201007942 | 2010 | France | Metropole | Egypt | ERS12446423 |
| 201008355 | 2010 | France | Metropole | Jordan | ERS12446424 |
| 201008713 | 2010 | France | Metropole | Unknown | ERS12445960 |
| 201008890 | 2010 | France | Metropole | None | ERR1516077 |
| 201008898 | 2010 | France | Metropole | Mexico | ERR1516078 |
| 201008902 | 2010 | France | Metropole | India | ERR1290102 |
| 201009047 | 2010 | France | Metropole | Egypt | ERS12446425 |
| 201009092 | 2010 | France | Metropole | Nepal | ERR1290103 |
| 201009281 | 2010 | France | Metropole | None | ERR1516079 |
| 201009353 | 2010 | France | Metropole | Unknown | ERS12445961 |
| 201009356 | 2010 | France | Metropole | India | ERR1290053 |
| 201009404 | 2010 | France | Metropole | None | ERR1516080 |
| 201009462 | 2010 | France | Metropole | India | ERS12446426 |
| 201009559 | 2010 | France | Metropole | Italy | ERS12445962 |
| 201009727 | 2010 | France | Metropole | None | ERR1516081 |
| 201100184 | 2011 | France | Metropole | India | ERR1290054 |
| 201100668 | 2011 | France | Metropole | None | ERR1516082 |
| 201100743 | 2011 | France | Metropole | None | ERR1516083 |
| 201101448 | 2011 | France | Metropole | India | ERS12445963 |
| 201101579 | 2011 | France | Metropole | Egypt | ERS12445964 |
| 201101951 | 2011 | France | Metropole | None | ERR1516084 |
| 201102248 | 2011 | France | Metropole | None | ERR1516085 |
| 201102489 | 2011 | France | Metropole | India | ERR1290055 |
| 201103739 | 2011 | France | Metropole | Cameroon | ERR1290056 |
| 201104124 | 2011 | France | Metropole | Unknown | ERS12445965 |
| 201104990 | 2011 | France | Metropole | Senegal | ERS12445966 |
| 201105547 | 2011 | France | Metropole | Unknown | ERS12445967 |
| 201106135 | 2011 | France | Metropole | Unknown | ERS12445968 |
| 201106390 | 2011 | France | Metropole | Unknown | ERS12445969 |
| 201107359 | 2011 | France | Metropole | India | ERR1290057 |
| 201107439 | 2011 | France | Metropole | None | ERR1516086 |
| 201107604 | 2011 | France | Metropole | Unknown | ERS12445970 |
| 201107709 | 2011 | France | Metropole | Unknown | ERS12445971 |
| 201108308 | 2011 | France | Metropole | India | ERR1290058 |
| 201108320 | 2011 | France | Metropole | Unknown | ERS12445972 |
| 201109322 | 2011 | France | Metropole | Unknown | ERS12445973 |
| 201110226 | 2011 | France | Metropole | Unknown | ERS12445974 |
| 201110264 | 2011 | France | Metropole | None | ERR1516087 |
| 201110473 | 2011 | France | Metropole | India | ERR1290059 |
| 201110495 | 2011 | France | Metropole | Spain | ERS12445975 |
| 201110496 | 2011 | France | Metropole | Spain | ERS12445976 |
| 201110602 | 2011 | France | Metropole | Unknown | ERS12445977 |
| 201110800 | 2011 | France | Metropole | None | ERR1516088 |
| 201110914 | 2011 | France | Metropole | None | ERR1541572 |

| | | | | | |
|---|---|---|---|---|---|
| 201111387 | 2011 | France | Metropole | None | ERR1541573 |
| 201111388 | 2011 | France | Metropole | None | ERR1541574 |
| 201111634 | 2011 | France | Metropole | India | ERR1290060 |
| 201111854 | 2011 | France | Metropole | Unknown | ERS12445978 |
| 201111941 | 2011 | France | Metropole | Unknown | ERS12445979 |
| 201112145 | 2011 | France | Metropole | None | ERR1541575 |
| 201200003 | 2011 | France | Metropole | Unknown | ERS12445980 |
| 201200407 | 2012 | France | Metropole | Unknown | ERS12445981 |
| 201201002 | 2012 | France | Metropole | Unknown | ERS12445982 |
| 201201272 | 2012 | France | Metropole | Unknown | ERS12445983 |
| 201201921 | 2012 | France | Metropole | Unknown | ERS12445984 |
| 201202008 | 2012 | France | Metropole | Unknown | ERS12445985 |
| 201202272 | 2012 | France | Metropole | India | ERR1290061 |
| 201202578 | 2012 | France | Metropole | Nepal | ERS12445986 |
| 201202584 | 2012 | France | Metropole | Unknown | ERS12445987 |
| 201202864 | 2012 | France | Metropole | Mexico | ERS12445988 |
| 201202874 | 2012 | France | Metropole | None | ERR1541576 |
| 201202950 | 2012 | France | Metropole | India | ERR1290062 |
| 201203160 | 2012 | France | Metropole | India | ERR1290063 |
| 201203380 | 2012 | France | Metropole | Unknown | ERS12445989 |
| 201203410 | 2012 | France | Metropole | India | ERR1290064 |
| 201204522 | 2012 | France | Metropole | Myanmar | ERR1290065 |
| 201204835 | 2012 | France | Metropole | India | ERR1290066 |
| 201204980 | 2012 | France | Metropole | Unknown | ERS12445990 |
| 201205359 | 2012 | France | Metropole | None | ERR1541577 |
| 201206128 | 2012 | France | Metropole | Unknown | ERS12445991 |
| 201206137 | 2012 | France | Metropole | Unknown | ERS12445992 |
| 201207010 | 2012 | France | Metropole | India | ERR1290067 |
| 201207078 | 2012 | France | Metropole | None | ERR1541578 |
| 201207116 | 2012 | France | Metropole | Unknown | ERS12445993 |
| 201207175 | 2012 | France | Metropole | Portugal | ERS12445994 |
| 201207290 | 2012 | France | Metropole | Unknown | ERS12445995 |
| 201207316 | 2012 | France | Metropole | India | ERR1290068 |
| 201208025 | 2012 | France | Metropole | Unknown | ERS12445996 |
| 201208633 | 2012 | France | Metropole | None | ERR1541580 |
| 201209000 | 2012 | France | Metropole | None | ERR1541581 |
| 201209109 | 2012 | France | Metropole | None | ERR1541582 |
| 201209330 | 2012 | France | Metropole | Unknown | ERS12445997 |
| 201209805 | 2012 | France | Metropole | Morocco | ERR1541583 |
| 201210011 | 2012 | France | Metropole | Unknown | ERS12445998 |
| 201210189 | 2012 | France | Metropole | India | ERS12445999 |
| 201210535 | 2012 | France | Metropole | Madagascar | ERR1541585 |
| 201210795 | 2012 | France | Metropole | Unknown | ERS12446000 |
| 201210818 | 2012 | France | Metropole | Turkey | ERS12446001 |
| 201211112 | 2012 | France | Metropole | Unknown | ERS12446002 |
| 201211243 | 2012 | France | Metropole | None | ERR1541586 |
| 201211330 | 2012 | France | Metropole | None | ERR1541587 |
| 201211333 | 2012 | France | Metropole | Unknown | ERS12446003 |
| 201211533 | 2012 | France | Metropole | Unknown | ERS12446004 |
| 201211535 | 2012 | France | Metropole | None | ERR1541588 |
| 201300694 | 2013 | France | Metropole | Unknown | ERS12446005 |
| 201301070 | 2013 | France | Metropole | Unknown | ERS12446006 |
| 201301550 | 2013 | France | Metropole | Unknown | ERS12446007 |
| 201302179 | 2013 | France | Metropole | Unknown | ERS12446008 |
| 201303008 | 2013 | France | Metropole | None | ERR1541590 |
| 201303126 | 2013 | France | Metropole | Unknown | ERS12446009 |
| 201303256 | 2013 | France | Metropole | India | ERS12446010 |
| 201303423 | 2013 | France | Metropole | Unknown | ERS12446011 |
| 201303702 | 2013 | France | Metropole | Unknown | ERS12446012 |
| 201303975 | 2013 | France | Metropole | Unknown | ERS12446013 |
| 201305624 | 2013 | France | Metropole | Unknown | ERS12446014 |
| 201305634 | 2013 | France | Metropole | India | ERS12446015 |
| 201305930 | 2013 | France | Metropole | Egypt | ERS12446016 |
| 201305931 | 2013 | France | Metropole | None | ERS12446017 |
| 201306244 | 2013 | France | Metropole | Unknown | ERS12446018 |
| 201306316 | 2013 | France | Metropole | None | ERS12446019 |
| 201306421 | 2013 | France | Metropole | Algeria | ERR1290070 |
| 201306478 | 2013 | France | Metropole | Unknown | ERS12446020 |
| 201306864 | 2013 | France | Metropole | None | ERR1541595 |
| 201307032 | 2013 | France | Metropole | Unknown | ERS12446021 |
| 201307033 | 2013 | France | Metropole | Unknown | ERS12446022 |
| 201307043 | 2013 | France | Metropole | Unknown | ERS12446023 |
| 201307374 | 2013 | France | Metropole | USA | ERR1290071 |
| 201307390 | 2013 | France | Metropole | Unknown | ERS12446024 |
| 201307631 | 2013 | France | Metropole | None | ERR1541596 |
| 201307632 | 2013 | France | Metropole | Spain | ERS12446025 |
| 201307843 | 2013 | France | Metropole | Unknown | ERS12446026 |
| 201307912 | 2013 | France | Metropole | India | ERR1290117 |
| 201307998 | 2013 | France | Metropole | India | ERR1290118 |
| 201308172 | 2013 | France | Metropole | None | ERR1541597 |
| 201308207 | 2013 | France | Metropole | Maghreb | ERS12446027 |
| 201308616 | 2013 | France | Metropole | Unknown | ERS12446028 |
| 201308995 | 2013 | France | Metropole | Unknown | ERS12446029 |
| 201309044 | 2013 | France | Metropole | Unknown | ERS12446030 |
| 201309047 | 2013 | France | Metropole | Unknown | ERS12446031 |
| 201309930 | 2013 | France | Metropole | India | ERR1290119 |
| 201310047 | 2013 | France | Metropole | Unknown | ERS12446032 |
| 201310566 | 2013 | France | Metropole | Tunisia | ERS12446033 |
| 201310767 | 2013 | France | Metropole | Germany | ERS12446034 |
| 201311321 | 2013 | France | Metropole | None | ERR1541598 |
| 201311425 | 2013 | France | Metropole | India | ERR1290120 |
| 201311547 | 2013 | France | Metropole | India | ERR1290121 |
| 201311626 | 2013 | France | Metropole | Unknown | ERR861643 |
| 201311838 | 2013 | France | Metropole | Unknown | ERR861644 |
| 201311953 | 2013 | France | Metropole | Unknown | ERR861645 |
| 201312273 | 2013 | France | Metropole | Unknown | ERR861646 |
| 201312374 | 2013 | France | Metropole | Unknown | ERR861647 |
| 201312454 | 2013 | France | Metropole | None | ERR1541599 |
| 201312520 | 2013 | France | Metropole | None | ERR1541600 |
| 201312542 | 2013 | France | Metropole | None | ERR861649 |
| 201312549 | 2013 | France | Metropole | Unknown | ERR861650 |
| 201400003 | 2013 | France | Metropole | Unknown | ERR861651 |
| 201400029 | 2013 | France | Metropole | Unknown | ERR861652 |

| | | | | | |
|---|---|---|---|---|---|
| 201400046 | 2013 | France | Metropole | Unknown | ERR861653 |
| 201400049 | 2013 | France | Metropole | Unknown | ERR861654 |
| 201400063 | 2013 | France | Metropole | Unknown | ERR861655 |
| 201400526 | 2014 | France | Metropole | Unknown | ERR861656 |
| 201400633 | 2014 | France | Metropole | Unknown | ERR861657 |
| 201400675 | 2014 | France | Metropole | None | ERR1541602 |
| 201400696 | 2014 | France | Metropole | Dominican Republic | ERR861658 |
| 201400752 | 2014 | France | Metropole | Unknown | ERR861659 |
| 201400855 | 2014 | France | Metropole | Unknown | ERR861660 |
| 201400893 | 2014 | France | Metropole | Unknown | ERR861661 |
| 201401034 | 2014 | France | Metropole | India | ERR1290122 |
| 201401054 | 2014 | France | Metropole | Unknown | ERR861662 |
| 201401096 | 2014 | France | Metropole | Unknown | ERR861663 |
| 201401124 | 2014 | France | Metropole | Unknown | ERR861664 |
| 201401237 | 2014 | France | Metropole | Unknown | ERR861665 |
| 201401254 | 2014 | France | Metropole | Dominican Republic | ERR1541603 |
| 201401278 | 2014 | France | Metropole | Unknown | ERR861666 |
| 201401461 | 2014 | France | Metropole | Cambodia | ERR1290123 |
| 201401463 | 2014 | France | Metropole | Unknown | ERR861667 |
| 201401464 | 2014 | France | Metropole | Unknown | ERR861668 |
| 201401573 | 2014 | France | Metropole | Unknown | ERR861669 |
| 201401659 | 2014 | France | Metropole | India | ERR1290124 |
| 201401695 | 2014 | France | Metropole | India | ERR1290125 |
| 201401742 | 2014 | France | Metropole | Unknown | ERR861670 |
| 201401743 | 2014 | France | Metropole | Unknown | ERR861671 |
| 201401787 | 2014 | France | Metropole | Unknown | ERS12446035 |
| 201401809 | 2014 | France | Metropole | Unknown | ERS12446036 |
| 201401842 | 2014 | France | Metropole | Unknown | ERR861672 |
| 201401879 | 2014 | France | Metropole | Unknown | ERR861673 |
| 201401929 | 2014 | France | Metropole | India | ERR1290126 |
| 201401955 | 2014 | France | Metropole | Unknown | ERS12446037 |
| 201402218 | 2014 | France | Metropole | Unknown | ERS12446038 |
| 201402395 | 2014 | France | Metropole | None | ERS12446039 |
| 201402408 | 2014 | France | Metropole | Unknown | ERR861674 |
| 201402425 | 2014 | France | Metropole | Unknown | ERR861675 |
| 201402444 | 2014 | France | Metropole | Unknown | ERS12446040 |
| 201402642 | 2014 | France | Metropole | French Polynesia | ERS12446041 |
| 201402814 | 2014 | France | Metropole | Unknown | ERS12446042 |
| 201402995 | 2014 | France | Metropole | Israel | ERR861676 |
| 201403274 | 2014 | France | Metropole | None | ERS12446043 |
| 201403597 | 2014 | France | Metropole | None | ERR1541604 |
| 201403691 | 2014 | France | Metropole | None | ERS12446044 |
| 201403715 | 2014 | France | Metropole | Thailand | ERR1290127 |
| 201403802 | 2014 | France | Metropole | Unknown | ERS12446045 |
| 201403834 | 2014 | France | Metropole | Canada | ERR861677 |
| 201403888 | 2014 | France | Metropole | None | ERS12446046 |
| 201403906 | 2014 | France | Metropole | Unknown | ERR861678 |
| 201403955 | 2014 | France | Metropole | Unknown | ERR861679 |
| 201403956 | 2014 | France | Metropole | Israel | ERR861680 |
| 201403963 | 2014 | France | Metropole | Israel | ERR861681 |
| 201404255 | 2014 | France | Metropole | None | ERS12446047 |
| 201404284 | 2014 | France | Metropole | None | ERS12446048 |
| 201404332 | 2014 | France | Metropole | Unknown | ERR861682 |
| 201404890 | 2014 | France | Metropole | Unknown | ERS12446049 |
| 201404910 | 2014 | France | Metropole | None | ERR1541605 |
| 201405003 | 2014 | France | Metropole | None | ERS12446050 |
| 201405020 | 2014 | France | Metropole | None | ERR1541606 |
| 201405079 | 2014 | France | Metropole | None | ERR861683 |
| 201405193 | 2014 | France | Metropole | None | ERR861684 |
| 201405267 | 2014 | France | Metropole | Unknown | ERR861685 |
| 201405269 | 2014 | France | Metropole | Unknown | ERS12446051 |
| 201405302 | 2014 | France | Metropole | Unknown | ERR861686 |
| 201405327 | 2014 | France | Metropole | Unknown | ERR861687 |
| 201405357 | 2014 | France | Metropole | Unknown | ERR861688 |
| 201405358 | 2014 | France | Metropole | Unknown | ERR861689 |
| 201405414 | 2014 | France | Metropole | None | ERS12446052 |
| 201405577 | 2014 | France | Metropole | None | ERR1541607 |
| 201405578 | 2014 | France | Metropole | None | ERS12446053 |
| 201405579 | 2014 | France | Metropole | Morocco | ERR1290128 |
| 201406445 | 2014 | France | Metropole | India | ERR1290129 |
| 201406456 | 2014 | France | Metropole | Unknown | ERS12446054 |
| 201406550 | 2014 | France | Metropole | India | ERR1290130 |
| 201406733 | 2014 | France | Metropole | None | ERS12446055 |
| 201407018 | 2014 | France | Metropole | None | ERR1541608 |
| 201407047 | 2014 | France | Metropole | Unknown | ERS12445938 |
| 201407049 | 2014 | France | Metropole | None | ERR1541609 |
| 201407070 | 2014 | France | Metropole | Unknown | ERS12446056 |
| 201407081 | 2014 | France | Metropole | India | ERR1290131 |
| 201407301 | 2014 | France | Metropole | Greece | ERS12446057 |
| 201407537 | 2014 | France | Metropole | India | ERR1290132 |
| 201407682 | 2014 | France | Metropole | Unknown | ERS12446058 |
| 201407778 | 2014 | France | Metropole | None | ERR1541610 |
| 201408083 | 2014 | France | Metropole | None | ERR1541611 |
| 201408085 | 2014 | France | Metropole | None | ERR1541612 |
| 201408517 | 2014 | France | Metropole | Unknown | ERS12446059 |
| 201408685 | 2014 | France | Metropole | None | ERR1541613 |
| 201408687 | 2014 | France | Metropole | Unknown | ERS12446060 |
| 201409042 | 2014 | France | Metropole | None | ERR1541614 |
| 201409146 | 2014 | France | Metropole | Unknown | ERS12446061 |
| 201409153 | 2014 | France | Metropole | Unknown | ERS12446062 |
| 201409154 | 2014 | France | Metropole | Algeria | ERS12446063 |
| 201409413 | 2014 | France | Metropole | None | ERR1541615 |
| 201409655 | 2014 | France | Metropole | Nepal | ERR1290133 |
| 201409824 | 2014 | France | Metropole | Unknown | ERS12446064 |
| 201409985 | 2014 | France | Metropole | None | ERR1541616 |
| 201410017 | 2014 | France | Metropole | Unknown | ERS12446065 |
| 201410020 | 2014 | France | Metropole | Italy | ERS12446066 |
| 201410165 | 2014 | France | Metropole | Unknown | ERS12446067 |
| 201410234 | 2014 | France | Metropole | Unknown | ERS12446068 |
| 201410939 | 2014 | France | Metropole | India | ERR1290134 |
| 201411698 | 2014 | France | Metropole | Cambodia | ERR1290135 |
| 201411716 | 2014 | France | Metropole | Morocco | ERR1541618 |
| 201411769 | 2014 | France | Metropole | None | ERS12446069 |
| 201412007 | 2014 | France | Metropole | India | ERS12446070 |

| | | | | | |
|---|---|---|---|---|---|
| 201412038 | 2014 | France | Metropole | Myanmar | ERR1541601 |
| 201412082 | 2014 | France | Metropole | None | ERS12446071 |
| 201412145 | 2014 | France | Metropole | None | ERR1541619 |
| 201412248 | 2014 | France | Metropole | None | ERS12446072 |
| 201412812 | 2014 | France | Metropole | Sudan | ERR1541620 |
| 201500606 | 2015 | France | Metropole | Cambodia | ERS12446073 |
| 201500996 | 2015 | France | Metropole | Cambodia | ERS12446074 |
| 201501147 | 2015 | France | Metropole | Cambodia | ERS12446075 |
| 201501197 | 2015 | France | Metropole | India | ERS12446076 |
| 201501493 | 2015 | France | Metropole | None | ERS12446077 |
| 201501582 | 2015 | France | Metropole | Unknown | ERS12446078 |
| 201502330 | 2015 | France | Metropole | Unknown | ERS12446079 |
| 201502475 | 2015 | France | Metropole | Unknown | ERS12446080 |
| 201502596 | 2015 | France | Metropole | Unknown | ERS12446081 |
| 201502874 | 2015 | France | Metropole | Unknown | ERS12446082 |
| 201502894 | 2015 | France | Metropole | Cambodia | ERS12446083 |
| 201502962 | 2015 | France | Metropole | Unknown | ERS12446084 |
| 201503165 | 2015 | France | Metropole | Unknown | ERS12446085 |
| 201503442 | 2015 | France | Metropole | Unknown | ERS12446086 |
| 201503669 | 2015 | France | Metropole | India | ERS12446087 |
| 201503778 | 2015 | France | Metropole | Unknown | ERS12446088 |
| 201503779 | 2015 | France | Metropole | Unknown | ERS12446089 |
| 201503780 | 2015 | France | Metropole | India | ERS12446090 |
| 201503782 | 2015 | France | Metropole | None | ERS12446091 |
| 201503792 | 2015 | France | Metropole | Unknown | ERS12446092 |
| 201503825 | 2015 | France | Metropole | None | ERS12446093 |
| 201504267 | 2015 | France | Metropole | Morocco | ERS12446094 |
| 201504297 | 2015 | France | Metropole | Unknown | ERS12446095 |
| 201504751 | 2015 | France | Metropole | None | ERS12446096 |
| 201504752 | 2015 | France | Metropole | Unknown | ERS12446097 |
| 201504754 | 2015 | France | Metropole | Unknown | ERS12446098 |
| 201505381 | 2015 | France | Metropole | Morocco | ERS12446099 |
| 201505383 | 2015 | France | Metropole | Unknown | ERS12446100 |
| 201505587 | 2015 | France | Metropole | Unknown | ERS12446101 |
| 201505659 | 2015 | France | Metropole | None | ERS12446102 |
| 201505943 | 2015 | France | Metropole | Unknown | ERS12446103 |
| 201505949 | 2015 | France | Metropole | Unknown | ERS12446104 |
| 201506202 | 2015 | France | Metropole | Germany | ERS12446105 |
| 201506485 | 2015 | France | Metropole | Asia | ERS12446106 |
| 201506814 | 2015 | France | Metropole | Thailand | ERS12446107 |
| 201506834 | 2015 | France | Metropole | Unknown | ERS12446108 |
| 201507184 | 2015 | France | Metropole | Unknown | ERS12446109 |
| 201508501 | 2015 | France | Metropole | Unknown | ERS12446110 |
| 201509157 | 2015 | France | Metropole | None | ERS12446111 |
| 201509163 | 2015 | France | Metropole | Costa Rica | ERS12446112 |
| 201509508 | 2015 | France | Metropole | Unknown | ERS12446113 |
| 201509509 | 2015 | France | Metropole | Unknown | ERS12446114 |
| 201509788 | 2015 | France | Metropole | Unknown | ERS12446115 |
| 201510021 | 2015 | France | Metropole | Unknown | ERS12446116 |
| 201510074 | 2015 | France | Metropole | Unknown | ERS12446117 |
| 201510152 | 2015 | France | Metropole | Unknown | ERS12446118 |
| 201510242 | 2015 | France | Metropole | Unknown | ERS12446119 |
| 201510390 | 2015 | France | Metropole | Unknown | ERS12446120 |
| 201510391 | 2015 | France | Metropole | Unknown | ERS12446121 |
| 201510465 | 2015 | France | Metropole | India | ERS12446122 |
| 201510619 | 2015 | France | Metropole | Unknown | ERS12446123 |
| 201510681 | 2015 | France | Metropole | Unknown | ERS12446124 |
| 201511284 | 2015 | France | Metropole | Unknown | ERS12446125 |
| 201511286 | 2015 | France | Metropole | Unknown | ERS12446126 |
| 201511415 | 2015 | France | Metropole | Unknown | ERS12446127 |
| 201511545 | 2015 | France | Metropole | Unknown | ERS12446128 |
| 201511600 | 2015 | France | Metropole | None | ERS12446129 |
| 201511668 | 2015 | France | Metropole | Sri Lanka | ERS12446130 |
| 201511681 | 2015 | France | Metropole | None | ERS12446131 |
| 201511775 | 2015 | France | Metropole | India | ERS12446132 |
| 201512005 | 2015 | France | Metropole | Unknown | ERS12446133 |
| 201512345 | 2015 | France | Metropole | Guinea | ERS12446134 |
| 201512528 | 2015 | France | Metropole | Unknown | ERS12446135 |
| 201512604 | 2015 | France | Metropole | India | ERS12446136 |
| 201512633 | 2015 | France | Metropole | Unknown | ERS12446137 |
| 201512634 | 2015 | France | Metropole | Unknown | ERS12446138 |
| 201512731 | 2015 | France | Metropole | Unknown | ERS12446139 |
| 201512907 | 2015 | France | Metropole | Unknown | ERS12446140 |
| 201512940 | 2015 | France | Metropole | Mauritius | ERS12446141 |
| 201513097 | 2015 | France | Metropole | Unknown | ERS12446142 |
| 201600158 | 2015 | France | Metropole | Unknown | ERS12446144 |
| 201600162 | 2015 | France | Metropole | Unknown | ERS12446145 |
| 201600117 | 2016 | France | Metropole | Unknown | ERS12446143 |
| 201600647 | 2016 | France | Metropole | India | ERS12446146 |
| 201600651 | 2016 | France | Metropole | Unknown | ERS12446147 |
| 201600653 | 2016 | France | Metropole | Unknown | ERS12446148 |
| 201600884 | 2016 | France | Metropole | Unknown | ERS12446149 |
| 201600966 | 2016 | France | Metropole | Unknown | ERS12446150 |
| 201601117 | 2016 | France | Metropole | Unknown | ERS12446151 |
| 201601157 | 2016 | France | Metropole | Unknown | ERS12446152 |
| 201601246 | 2016 | France | Metropole | Cambodia | ERS12446153 |
| 201601367 | 2016 | France | Metropole | China | ERS12446154 |
| 201601509 | 2016 | France | Metropole | Unknown | ERS12446155 |
| 201601512 | 2016 | France | Metropole | Unknown | ERS12446156 |
| 201601652 | 2016 | France | Metropole | Cambodia | ERS12446157 |
| 201601748 | 2016 | France | Metropole | Unknown | ERS12446158 |
| 201602115 | 2016 | France | Metropole | Unknown | ERS12446159 |
| 201602347 | 2016 | France | Metropole | Unknown | ERS12446160 |
| 201602350 | 2016 | France | Metropole | Brazil | ERS12446161 |
| 201602860 | 2016 | France | Metropole | Unknown | ERS12446162 |
| 201602861 | 2016 | France | Metropole | Unknown | ERS12446163 |
| 201602863 | 2016 | France | Metropole | Unknown | ERS12446164 |
| 201602864 | 2016 | France | Metropole | Unknown | ERS12446165 |
| 201602867 | 2016 | France | Metropole | Unknown | ERS12446166 |
| 201603117 | 2016 | France | Metropole | Unknown | ERS12446167 |
| 201603190 | 2016 | France | Metropole | Morocco | ERS12446168 |
| 201603252 | 2016 | France | Metropole | India | ERS12446169 |
| 201603634 | 2016 | France | Metropole | Unknown | ERS12446170 |

| | | | | | |
|---|---|---|---|---|---|
| 201603739 | 2016 | France | Metropole | Unknown | ERS12446171 |
| 201603742 | 2016 | France | Metropole | Unknown | ERS12446172 |
| 201603813 | 2016 | France | Metropole | None | ERS12446173 |
| 201603898 | 2016 | France | Metropole | Unknown | ERS12446174 |
| 201604144 | 2016 | France | Metropole | Unknown | ERS12446175 |
| 201604283 | 2016 | France | Metropole | Spain | ERS12446176 |
| 201604332 | 2016 | France | Metropole | None | ERS12446177 |
| 201604478 | 2016 | France | Metropole | None | ERS12446178 |
| 201604588 | 2016 | France | Metropole | None | ERS12446179 |
| 201604682 | 2016 | France | Metropole | Turkey | ERS12446180 |
| 201604687 | 2016 | France | Metropole | Unknown | ERS12446181 |
| 201604761 | 2016 | France | Metropole | Italy | ERS12446182 |
| 201605018 | 2016 | France | Metropole | Unknown | ERS12446183 |
| 201605089 | 2016 | France | Metropole | Unknown | ERS12446184 |
| 201605090 | 2016 | France | Metropole | Unknown | ERS12446185 |
| 201605735 | 2016 | France | Metropole | None | ERS12446186 |
| 201605743 | 2016 | France | Metropole | Unknown | ERS12446187 |
| 201605745 | 2016 | France | Metropole | Unknown | ERS12446188 |
| 201605809 | 2016 | France | Metropole | Unknown | ERS12446189 |
| 201605946 | 2016 | France | Metropole | None | ERS12446190 |
| 201605965 | 2016 | France | Metropole | Unknown | ERS12446191 |
| 201606031 | 2016 | France | Metropole | Sudan | ERS12446192 |
| 201606036 | 2016 | France | Metropole | Unknown | ERS12446193 |
| 201606447 | 2016 | France | Metropole | India | ERS12446194 |
| 201606592 | 2016 | France | Metropole | USA | ERS12446195 |
| 201606749 | 2016 | France | Metropole | Unknown | ERS12446196 |
| 201606817 | 2016 | France | Metropole | Unknown | ERS12446197 |
| 201606870 | 2016 | France | Metropole | Unknown | ERS12446198 |
| 201606993 | 2016 | France | Metropole | Unknown | ERS12446199 |
| 201607131 | 2016 | France | Metropole | Unknown | ERS12446200 |
| 201607242 | 2016 | France | Metropole | Unknown | ERS12446201 |
| 201607279 | 2016 | France | Metropole | None | ERS12446202 |
| 201607467 | 2016 | France | Metropole | Unknown | ERS12446203 |
| 201607759 | 2016 | France | Metropole | Unknown | ERS12446204 |
| 201607786 | 2016 | France | Metropole | Peru | ERS12446205 |
| 201607913 | 2016 | France | Metropole | None | ERS12446206 |
| 201608087 | 2016 | France | Metropole | Unknown | ERS12446207 |
| 201608145 | 2016 | France | Metropole | Unknown | ERS12446208 |
| 201608572 | 2016 | France | Metropole | Unknown | ERS12446209 |
| 201608836 | 2016 | France | Metropole | Cuba | ERS12446210 |
| 201608858 | 2016 | France | Metropole | Unknown | ERS12446211 |
| 201608950 | 2016 | France | Metropole | Unknown | ERS12446212 |
| 201609048 | 2016 | France | Metropole | Unknown | ERS12446213 |
| 201609067 | 2016 | France | Metropole | Unknown | ERS12446214 |
| 201609069 | 2016 | France | Metropole | Unknown | ERS12446215 |
| 201609123 | 2016 | France | Metropole | Unknown | ERS12446216 |
| 201609226 | 2016 | France | Metropole | Unknown | ERS12446217 |
| 201609525 | 2016 | France | Metropole | Unknown | ERS12446218 |
| 201609526 | 2016 | France | Metropole | Unknown | ERS12446219 |
| 201609802 | 2016 | France | Metropole | Unknown | ERS12446220 |
| 201609836 | 2016 | France | Metropole | Unknown | ERS12446221 |
| 201609975 | 2016 | France | Metropole | Unknown | ERS12446222 |
| 201610028 | 2016 | France | Metropole | Unknown | ERS12446223 |
| 201610080 | 2016 | France | Metropole | Unknown | ERS12446224 |
| 201610090 | 2016 | France | Metropole | Unknown | ERS12446225 |
| 201610216 | 2016 | France | Metropole | None | ERS12446226 |
| 201610335 | 2016 | France | Metropole | None | ERS12446227 |
| 201610461 | 2016 | France | Metropole | Unknown | ERS12446228 |
| 201610631 | 2016 | France | Metropole | None | ERS12446229 |
| 201610666 | 2016 | France | Metropole | None | ERS12446230 |
| 201610996 | 2016 | France | Metropole | Nepal | ERS12446231 |
| 201611169 | 2016 | France | Metropole | Unknown | ERS12446232 |
| 201611351 | 2016 | France | Metropole | Unknown | ERS12446233 |
| 201611359 | 2016 | France | Metropole | Unknown | ERS12446234 |
| 201611488 | 2016 | France | Metropole | Unknown | ERS12446235 |
| 201611650 | 2016 | France | Metropole | None | ERS12446236 |
| 201611705 | 2016 | France | Metropole | None | ERS12446237 |
| 201611756 | 2016 | France | Metropole | Thailand | ERS12446238 |
| 201700025 | 2016 | France | Metropole | Unknown | ERS6495648 |
| 201700037 | 2016 | France | Metropole | Africa | ERS6495649 |
| 201700298 | 2016 | France | Metropole | Unknown | ERS6495651 |
| 201700454 | 2016 | France | French Guiana | Unknown | ERS6495652 |
| 201700456 | 2016 | France | French Guiana | Unknown | ERS6495653 |
| 201700726 | 2016 | France | Metropole | None | ERS6495661 |
| 201700895 | 2016 | France | French Guiana | Unknown | ERS6495668 |
| 201700897 | 2016 | France | French Guiana | Unknown | ERS6495669 |
| 201702142 | 2016 | France | French Guiana | Unknown | ERS6495725 |
| 201700244 | 2017 | France | Metropole | Unknown | ERS6495650 |
| 201700471 | 2017 | France | Metropole | India | ERS6495654 |
| 201700472 | 2017 | France | Metropole | Unknown | ERS6495655 |
| 201700474 | 2017 | France | Metropole | Unknown | ERS6495656 |
| 201700585 | 2017 | France | Metropole | None | ERS6495657 |
| 201700623 | 2017 | France | Metropole | Mali | ERS6495658 |
| 201700723 | 2017 | France | Metropole | Unknown | ERS6495659 |
| 201700724 | 2017 | France | Metropole | Unknown | ERS6495660 |
| 201700752 | 2017 | France | Metropole | Unknown | ERS6495662 |
| 201700773 | 2017 | France | Metropole | Saudi Arabia | ERS6495663 |
| 201700801 | 2017 | France | Metropole | Unknown | ERS6495664 |
| 201700823 | 2017 | France | French Guiana | Unknown | ERS6495665 |
| 201700824 | 2017 | France | Metropole | Unknown | ERS6495666 |
| 201700852 | 2017 | France | Metropole | Unknown | ERS6495667 |
| 201700905 | 2017 | France | Metropole | Unknown | ERS6495670 |
| 201700936 | 2017 | France | French Guiana | Unknown | ERS6495671 |
| 201700937 | 2017 | France | Metropole | Unknown | ERS6495672 |
| 201700965 | 2017 | France | Metropole | Unknown | ERS6495673 |
| 201701025 | 2017 | France | Metropole | Unknown | ERS6495674 |
| 201701049 | 2017 | France | Metropole | Colombia | ERS6495675 |
| 201701093 | 2017 | France | Metropole | Unknown | ERS6492764 |
| 201701125 | 2017 | France | Metropole | India | ERS6495676 |
| 201701239 | 2017 | France | Metropole | Unknown | ERS6495677 |
| 201701293 | 2017 | France | Metropole | Unknown | ERS6495678 |
| 201701294 | 2017 | France | Metropole | Unknown | ERS6495679 |
| 201701295 | 2017 | France | Metropole | Unknown | ERS6495680 |

| | | | | | |
|---|---|---|---|---|---|
| 201701296 | 2017 | France | Metropole | Unknown | ERS6495681 |
| 201701297 | 2017 | France | Metropole | Unknown | ERS6495682 |
| 201701298 | 2017 | France | Metropole | Unknown | ERS6495683 |
| 201701299 | 2017 | France | Metropole | Unknown | ERS6495684 |
| 201701300 | 2017 | France | Metropole | Unknown | ERS6495685 |
| 201701337 | 2017 | France | Metropole | Reunion | ERS6575812 |
| 201701367 | 2017 | France | Metropole | None | ERS6495686 |
| 201701372 | 2017 | France | Metropole | Unknown | ERS6495687 |
| 201701373 | 2017 | France | Metropole | Unknown | ERS6495688 |
| 201701407 | 2017 | France | Metropole | Costa Rica | ERS6495689 |
| 201701433 | 2017 | France | Metropole | Unknown | ERS6495690 |
| 201701437 | 2017 | France | Metropole | Unknown | ERS6495691 |
| 201701504 | 2017 | France | Mayotte | Unknown | ERS6495692 |
| 201701512 | 2017 | France | Metropole | Unknown | ERS6495693 |
| 201701529 | 2017 | France | Metropole | Unknown | ERS6495694 |
| 201701530 | 2017 | France | Metropole | Unknown | ERS6495695 |
| 201701531 | 2017 | France | Metropole | Unknown | ERS6495696 |
| 201701532 | 2017 | France | Metropole | Unknown | ERS6495697 |
| 201701533 | 2017 | France | Metropole | Unknown | ERS6495698 |
| 201701534 | 2017 | France | Metropole | Unknown | ERS6495699 |
| 201701535 | 2017 | France | Metropole | Unknown | ERS6495700 |
| 201701545 | 2017 | France | Metropole | Unknown | ERS6495701 |
| 201701548 | 2017 | France | Metropole | Unknown | ERS6495702 |
| 201701554 | 2017 | France | Metropole | Myanmar | ERS6495703 |
| 201701555 | 2017 | France | Metropole | Unknown | ERS6495704 |
| 201701576 | 2017 | France | Metropole | Unknown | ERS6574674 |
| 201701584 | 2017 | France | Metropole | None | ERS6495705 |
| 201701701 | 2017 | France | Metropole | None | ERS6495706 |
| 201701714 | 2017 | France | Metropole | Unknown | ERS6495707 |
| 201701715 | 2017 | France | Metropole | Unknown | ERS6495708 |
| 201701716 | 2017 | France | Metropole | Unknown | ERS6495709 |
| 201701800 | 2017 | France | Metropole | Guinea | ERS6495710 |
| 201701807 | 2017 | France | Metropole | Unknown | ERS6495713 |
| 201701808 | 2017 | France | Metropole | Unknown | ERS6495714 |
| 201701845 | 2017 | France | Metropole | Thailand | ERS6495715 |
| 201701847 | 2017 | France | Metropole | Unknown | ERS6495716 |
| 201701849 | 2017 | France | Metropole | Nicaragua | ERS6495717 |
| 201701853 | 2017 | France | Metropole | Unknown | ERS6495718 |
| 201701870 | 2017 | France | Metropole | Côte d'Ivoire | ERS6495719 |
| 201701912 | 2017 | France | Metropole | Cuba | ERS6495720 |
| 201701936 | 2017 | France | Metropole | Unknown | ERS6575813 |
| 201701937 | 2017 | France | Metropole | Haiti | ERS6495721 |
| 201702075 | 2017 | France | Metropole | Unknown | ERS6575814 |
| 201702096 | 2017 | France | Metropole | Dominican Republic | ERS6495722 |
| 201702105 | 2017 | France | Metropole | Unknown | ERS6495723 |
| 201702106 | 2017 | France | Metropole | None | ERS6495724 |
| 201702179 | 2017 | France | Metropole | Unknown | ERS6495726 |
| 201702180 | 2017 | France | Metropole | Unknown | ERS6495727 |
| 201702219 | 2017 | France | Metropole | India | ERS6495728 |
| 201702296 | 2017 | France | Metropole | Unknown | ERS6495729 |
| 201702322 | 2017 | France | Metropole | Unknown | ERS6495730 |
| 201702347 | 2017 | France | Metropole | Unknown | ERS6495731 |
| 201702350 | 2017 | France | Metropole | Unknown | ERS6495732 |
| 201702375 | 2017 | France | Metropole | Unknown | ERS6495733 |
| 201702421 | 2017 | France | Metropole | Unknown | ERS6495734 |
| 201702422 | 2017 | France | Metropole | Unknown | ERS6495735 |
| 201702423 | 2017 | France | Metropole | Unknown | ERS6495736 |
| 201702424 | 2017 | France | Metropole | Unknown | ERS6495737 |
| 201702425 | 2017 | France | Metropole | Unknown | ERS6495738 |
| 201702469 | 2017 | France | Metropole | Unknown | ERS6495739 |
| 201702470 | 2017 | France | Metropole | Unknown | ERS6495740 |
| 201702483 | 2017 | France | Metropole | Unknown | ERS6495742 |
| 201702484 | 2017 | France | Metropole | Unknown | ERS6495743 |
| 201702485 | 2017 | France | Metropole | Unknown | ERS6495744 |
| 201702486 | 2017 | France | Metropole | Unknown | ERS6495745 |
| 201702487 | 2017 | France | Metropole | Unknown | ERS6495746 |
| 201702488 | 2017 | France | Metropole | Unknown | ERS6495747 |
| 201702489 | 2017 | France | Metropole | Unknown | ERS6495748 |
| 201702490 | 2017 | France | Metropole | Unknown | ERS6495749 |
| 201702491 | 2017 | France | Metropole | None | ERS6495750 |
| 201702492 | 2017 | France | Metropole | Dominican Republic | ERS6495751 |
| 201702539 | 2017 | France | Metropole | Unknown | ERS6495754 |
| 201702540 | 2017 | France | Metropole | None | ERS6495755 |
| 201702544 | 2017 | France | Metropole | Unknown | ERS6495756 |
| 201702545 | 2017 | France | Metropole | Unknown | ERS6495757 |
| 201702562 | 2017 | France | Metropole | Unknown | ERS6495758 |
| 201702563 | 2017 | France | Metropole | Unknown | ERS6495759 |
| 201702564 | 2017 | France | Metropole | Unknown | ERS6495760 |
| 201702620 | 2017 | France | Metropole | Unknown | ERS6495761 |
| 201702631 | 2017 | France | Metropole | None | ERS6495762 |
| 201702695 | 2017 | France | Metropole | Unknown | ERS6575816 |
| 201702701 | 2017 | France | Mayotte | Unknown | ERS6575817 |
| 201702702 | 2017 | France | Mayotte | Unknown | ERS6575818 |
| 201702704 | 2017 | France | Mayotte | Unknown | ERS6575819 |
| 201702706 | 2017 | France | Mayotte | Unknown | ERS6575820 |
| 201702742 | 2017 | France | Metropole | Unknown | ERS6575821 |
| 201702746 | 2017 | France | Metropole | None | ERS6575822 |
| 201702781 | 2017 | France | Metropole | Unknown | ERS6495763 |
| 201702782 | 2017 | France | Metropole | Unknown | ERS6495764 |
| 201702783 | 2017 | France | Metropole | Unknown | ERS6495765 |
| 201702784 | 2017 | France | Metropole | Unknown | ERS6495766 |
| 201702786 | 2017 | France | Metropole | Unknown | ERS6495768 |
| 201702787 | 2017 | France | Metropole | Unknown | ERS6575823 |
| 201702788 | 2017 | France | Metropole | Unknown | ERS6575824 |
| 201702789 | 2017 | France | Metropole | Unknown | ERS6575825 |
| 201702790 | 2017 | France | Metropole | Unknown | ERS6575826 |
| 201702791 | 2017 | France | Metropole | Unknown | ERS6575827 |
| 201702809 | 2017 | France | Metropole | Unknown | ERS6575828 |
| 201702846 | 2017 | France | Metropole | Unknown | ERS6575829 |
| 201702847 | 2017 | France | Metropole | Unknown | ERS6575830 |
| 201702850 | 2017 | France | Metropole | Unknown | ERS6575831 |
| 201702901 | 2017 | France | Metropole | Brazil | ERS6575832 |
| 201702902 | 2017 | France | Metropole | Unknown | ERS6575833 |

| | | | | | |
|---|---|---|---|---|---|
| 201702903 | 2017 | France | Metropole | Unknown | ERS6575834 |
| 201702904 | 2017 | France | Metropole | Unknown | ERS6575835 |
| 201702905 | 2017 | France | Metropole | Unknown | ERS6575836 |
| 201702906 | 2017 | France | Metropole | Unknown | ERS6575837 |
| 201702936 | 2017 | France | Metropole | None | ERS6575838 |
| 201702962 | 2017 | France | Metropole | Unknown | ERS6575839 |
| 201703031 | 2017 | France | Metropole | Unknown | ERS6575840 |
| 201703068 | 2017 | France | Metropole | Unknown | ERS6575841 |
| 201703089 | 2017 | France | Metropole | Unknown | ERS6575842 |
| 201703090 | 2017 | France | Metropole | Unknown | ERS6575843 |
| 201703128 | 2017 | France | Metropole | India | ERS6575844 |
| 201703205 | 2017 | France | Metropole | None | ERS6575845 |
| 201703224 | 2017 | France | Metropole | None | ERS6575846 |
| 201703316 | 2017 | France | Metropole | India | ERS6575847 |
| 201703318 | 2017 | France | Metropole | Thailand | ERS6575848 |
| 201703369 | 2017 | France | Metropole | Unknown | ERS6575849 |
| 201703389 | 2017 | France | Metropole | Unknown | ERS6575850 |
| 201703436 | 2017 | France | Metropole | Thailand | ERS6575851 |
| 201703437 | 2017 | France | Metropole | Unknown | ERS6575852 |
| 201703446 | 2017 | France | French Guiana | Unknown | ERS6575853 |
| 201703512 | 2017 | France | Metropole | Unknown | ERS6575854 |
| 201703663 | 2017 | France | Metropole | India | ERS6575855 |
| 201703727 | 2017 | France | Metropole | None | ERS6575856 |
| 201703779 | 2017 | France | Metropole | Unknown | ERS6575857 |
| 201703781 | 2017 | France | Metropole | Unknown | ERS6575858 |
| 201703843 | 2017 | France | Metropole | Unknown | ERS6575859 |
| 201703900 | 2017 | France | Metropole | Lebanon | ERS6575860 |
| 201703902 | 2017 | France | Metropole | Unknown | ERS6575861 |
| 201703904 | 2017 | France | Metropole | Unknown | ERS6575416 |
| 201703907 | 2017 | France | Metropole | Unknown | ERS6575862 |
| 201703908 | 2017 | France | Metropole | Unknown | ERS6575863 |
| 201703912 | 2017 | France | Metropole | Unknown | ERS6575864 |
| 201703915 | 2017 | France | Metropole | Unknown | ERS6575865 |
| 201703925 | 2017 | France | Metropole | Unknown | ERS6575866 |
| 201703952 | 2017 | France | Metropole | Unknown | ERS6575867 |
| 201703966 | 2017 | France | Metropole | None | ERS6575868 |
| 201704018 | 2017 | France | Metropole | Indonesia | ERS6575869 |
| 201704049 | 2017 | France | Metropole | Unknown | ERS6575870 |
| 201704062 | 2017 | France | Metropole | Unknown | ERS6575871 |
| 201704090 | 2017 | France | Metropole | Unknown | ERS6575872 |
| 201704091 | 2017 | France | Metropole | Unknown | ERS6575873 |
| 201704112 | 2017 | France | Metropole | Unknown | ERS6575874 |
| 201704120 | 2017 | France | Metropole | Unknown | ERS6575875 |
| 201704134 | 2017 | France | Metropole | Unknown | ERS6575876 |
| 201704135 | 2017 | France | Metropole | Unknown | ERS6575877 |
| 201704136 | 2017 | France | Metropole | Unknown | ERS6575878 |
| 201704137 | 2017 | France | Metropole | Unknown | ERS6575879 |
| 201704138 | 2017 | France | Metropole | Unknown | ERS6575880 |
| 201704139 | 2017 | France | Metropole | Unknown | ERS6575881 |
| 201704140 | 2017 | France | Metropole | Unknown | ERS6575882 |
| 201704259 | 2017 | France | Metropole | Unknown | ERS6575888 |
| 201704289 | 2017 | France | Metropole | Martinique | ERS6575889 |
| 201704301 | 2017 | France | Metropole | Unknown | ERS6575890 |
| 201704332 | 2017 | France | Metropole | None | ERS6575891 |
| 201704333 | 2017 | France | Metropole | Unknown | ERS6575892 |
| 201704336 | 2017 | France | Metropole | Unknown | ERS6575894 |
| 201704354 | 2017 | France | Metropole | Unknown | ERS6575895 |
| 201704371 | 2017 | France | Metropole | Unknown | ERS6575896 |
| 201704480 | 2017 | France | Metropole | Unknown | ERS6575897 |
| 201704483 | 2017 | France | Metropole | Madagascar | ERS6575898 |
| 201704554 | 2017 | France | Metropole | None | ERS6495769 |
| 201704580 | 2017 | France | Metropole | Unknown | ERS6495770 |
| 201704581 | 2017 | France | Metropole | Unknown | ERS6495771 |
| 201704591 | 2017 | France | Metropole | Unknown | ERS6495772 |
| 201704599 | 2017 | France | Metropole | None | ERS6495773 |
| 201704604 | 2017 | France | Metropole | Egypt | ERS6495774 |
| 201704605 | 2017 | France | Metropole | Morocco | ERS6495775 |
| 201704626 | 2017 | France | Metropole | None | ERS6495776 |
| 201704644 | 2017 | France | Metropole | Unknown | ERS6495777 |
| 201704655 | 2017 | France | French Guiana | Unknown | ERS6495778 |
| 201704656 | 2017 | France | French Guiana | Unknown | ERS6495779 |
| 201704700 | 2017 | France | Metropole | None | ERS6495780 |
| 201704709 | 2017 | France | Metropole | Unknown | ERS6495781 |
| 201704740 | 2017 | France | Metropole | None | ERS6495782 |
| 201704741 | 2017 | France | Metropole | Unknown | ERS6495783 |
| 201704742 | 2017 | France | Metropole | Unknown | ERS6495784 |
| 201704743 | 2017 | France | Metropole | Unknown | ERS6495785 |
| 201704744 | 2017 | France | Metropole | Unknown | ERS6495786 |
| 201704747 | 2017 | France | Metropole | Togo | ERS6495787 |
| 201704748 | 2017 | France | Metropole | None | ERS6495788 |
| 201704781 | 2017 | France | Metropole | Unknown | ERS6495789 |
| 201704799 | 2017 | France | Metropole | Unknown | ERS6495790 |
| 201704802 | 2017 | France | Metropole | Unknown | ERS6495791 |
| 201704803 | 2017 | France | Metropole | Unknown | ERS6495792 |
| 201704804 | 2017 | France | Metropole | Unknown | ERS6495793 |
| 201704853 | 2017 | France | Metropole | Israel | ERS6495794 |
| 201704868 | 2017 | France | Metropole | Lebanon | ERS6495795 |
| 201704871 | 2017 | France | Metropole | None | ERS6575899 |
| 201704900 | 2017 | France | Metropole | Mali | ERS6495796 |
| 201704904 | 2017 | France | Metropole | Unknown | ERS6495797 |
| 201704908 | 2017 | France | Metropole | Unknown | ERS6495798 |
| 201704911 | 2017 | France | Metropole | Unknown | ERS6495799 |
| 201704942 | 2017 | France | Metropole | Israel | ERS6495800 |
| 201704993 | 2017 | France | Metropole | Unknown | ERS6495801 |
| 201705046 | 2017 | France | Metropole | Unknown | ERS6495802 |
| 201705062 | 2017 | France | Metropole | Unknown | ERS6495803 |
| 201705116 | 2017 | France | Metropole | None | ERS6495804 |
| 201705119 | 2017 | France | Metropole | Unknown | ERS6495805 |
| 201705123 | 2017 | France | French Guiana | Unknown | ERS6495806 |
| 201705133 | 2017 | France | Metropole | Unknown | ERS6495808 |
| 201705150 | 2017 | France | Metropole | Unknown | ERS6495809 |
| 201705214 | 2017 | France | Metropole | None | ERS6495810 |
| 201705240 | 2017 | France | Metropole | None | ERS6574675 |

| | | | | | |
|---|---|---|---|---|---|
| 201705248 | 2017 | France | Metropole | Unknown | ERS6574676 |
| 201705252 | 2017 | France | Metropole | Unknown | ERS6574677 |
| 201705280 | 2017 | France | Metropole | India | ERS6574678 |
| 201705306 | 2017 | France | Metropole | Unknown | ERS6574679 |
| 201705308 | 2017 | France | Metropole | Unknown | ERS6574680 |
| 201705310 | 2017 | France | Metropole | Unknown | ERS6574681 |
| 201705311 | 2017 | France | Metropole | Unknown | ERS6574682 |
| 201705312 | 2017 | France | Metropole | Unknown | ERS6574683 |
| 201705313 | 2017 | France | Metropole | Unknown | ERS6574684 |
| 201705314 | 2017 | France | Metropole | Unknown | ERS6574685 |
| 201705315 | 2017 | France | Metropole | Unknown | ERS6574686 |
| 201705334 | 2017 | France | Metropole | Unknown | ERS6574687 |
| 201705337 | 2017 | France | Metropole | Unknown | ERS6574688 |
| 201705361 | 2017 | France | Metropole | Unknown | ERS6574689 |
| 201705377 | 2017 | France | Metropole | None | ERS6574690 |
| 201705413 | 2017 | France | Metropole | Unknown | ERS6574691 |
| 201705415 | 2017 | France | Metropole | Unknown | ERS6574692 |
| 201705418 | 2017 | France | Metropole | Unknown | ERS6574693 |
| 201705437 | 2017 | France | Metropole | Unknown | ERS6574694 |
| 201705464 | 2017 | France | Metropole | Unknown | ERS6574695 |
| 201705478 | 2017 | France | Metropole | Unknown | ERS6574696 |
| 201705484 | 2017 | France | Metropole | None | ERS6574697 |
| 201705487 | 2017 | France | Metropole | Unknown | ERS6574698 |
| 201705569 | 2017 | France | Metropole | Unknown | ERS6574699 |
| 201705583 | 2017 | France | Metropole | Unknown | ERS6574700 |
| 201705586 | 2017 | France | Metropole | Unknown | ERS6575900 |
| 201705612 | 2017 | France | Metropole | None | ERS6574701 |
| 201705650 | 2017 | France | Metropole | Unknown | ERS6574702 |
| 201705654 | 2017 | France | Metropole | Unknown | ERS6574703 |
| 201705694 | 2017 | France | Metropole | None | ERS6574704 |
| 201705719 | 2017 | France | Metropole | Unknown | ERS6574705 |
| 201705767 | 2017 | France | Metropole | None | ERS6574706 |
| 201705836 | 2017 | France | Metropole | Unknown | ERS6574707 |
| 201705978 | 2017 | France | Metropole | Unknown | ERS6574708 |
| 201705982 | 2017 | France | Metropole | Morocco | ERS6574709 |
| 201706028 | 2017 | France | Metropole | Unknown | ERS6574710 |
| 201706048 | 2017 | France | Metropole | Unknown | ERS6574711 |
| 201706069 | 2017 | France | Metropole | Unknown | ERS6574712 |
| 201706072 | 2017 | France | Metropole | Morocco | ERS6574714 |
| 201706073 | 2017 | France | Metropole | Unknown | ERS6574715 |
| 201706074 | 2017 | France | Metropole | Unknown | ERS6574716 |
| 201706076 | 2017 | France | Metropole | Unknown | ERS6574717 |
| 201706095 | 2017 | France | Metropole | Unknown | ERS6574718 |
| 201706104 | 2017 | France | Metropole | Unknown | ERS6574719 |
| 201706135 | 2017 | France | Metropole | Unknown | ERS6574720 |
| 201706146 | 2017 | France | Metropole | Unknown | ERS6574721 |
| 201706147 | 2017 | France | Metropole | Unknown | ERS6574722 |
| 201706198 | 2017 | France | Metropole | Unknown | ERS6574723 |
| 201706282 | 2017 | France | Metropole | Unknown | ERS6575417 |
| 201706285 | 2017 | France | Metropole | Unknown | ERS6575901 |
| 201706375 | 2017 | France | Metropole | Unknown | ERS6575902 |
| 201706406 | 2017 | France | Metropole | Cuba | ERS6575903 |
| 201706426 | 2017 | France | Metropole | Unknown | ERS6575904 |
| 201706517 | 2017 | France | Metropole | Unknown | ERS6575905 |
| 201706518 | 2017 | France | Metropole | Gambia | ERS6575906 |
| 201706522 | 2017 | France | Metropole | Unknown | ERS6575907 |
| 201706539 | 2017 | France | Metropole | Unknown | ERS6575908 |
| 201706573 | 2017 | France | Metropole | Italy | ERS6575909 |
| 201706608 | 2017 | France | Metropole | Unknown | ERS6575910 |
| 201706648 | 2017 | France | French Guiana | Unknown | ERS6575911 |
| 201706711 | 2017 | France | Metropole | Peru | ERS6575912 |
| 201706740 | 2017 | France | Metropole | Unknown | ERS6575913 |
| 201706741 | 2017 | France | Metropole | Unknown | ERS6575914 |
| 201706763 | 2017 | France | Metropole | Unknown | ERS6575915 |
| 201706767 | 2017 | France | Metropole | Unknown | ERS6575916 |
| 201706820 | 2017 | France | Metropole | None | ERS6575917 |
| 201706899 | 2017 | France | Metropole | Mexico | ERS6575918 |
| 201706913 | 2017 | France | Metropole | Senegal | ERS6575919 |
| 201706929 | 2017 | France | Metropole | Unknown | ERS6575920 |
| 201706933 | 2017 | France | Metropole | Mali | ERS6575921 |
| 201706983 | 2017 | France | Metropole | Unknown | ERS6575922 |
| 201706989 | 2017 | France | Metropole | Cambodia | ERS6574724 |
| 201707008 | 2017 | France | Metropole | Algeria | ERS6574725 |
| 201707022 | 2017 | France | Metropole | None | ERS6576210 |
| 201707079 | 2017 | France | Metropole | Unknown | ERS6574726 |
| 201707086 | 2017 | France | Metropole | Unknown | ERS6574727 |
| 201707195 | 2017 | France | Metropole | Unknown | ERS6574728 |
| 201707284 | 2017 | France | Metropole | Indonesia | ERS6574730 |
| 201707285 | 2017 | France | Metropole | Spain | ERS6574731 |
| 201707286 | 2017 | France | Metropole | Unknown | ERS6574732 |
| 201707302 | 2017 | France | Metropole | Unknown | ERS6574733 |
| 201707308 | 2017 | France | Metropole | Unknown | ERS6574734 |
| 201707311 | 2017 | France | Metropole | Algeria | ERS6574735 |
| 201707335 | 2017 | France | Metropole | Morocco | ERS6574736 |
| 201707343 | 2017 | France | Metropole | Unknown | ERS6574737 |
| 201707380 | 2017 | France | Metropole | Morocco | ERS6574738 |
| 201707399 | 2017 | France | Metropole | Thailand | ERS6574739 |
| 201707422 | 2017 | France | Metropole | Unknown | ERS6574740 |
| 201707473 | 2017 | France | Metropole | Unknown | ERS6574742 |
| 201707474 | 2017 | France | Metropole | Mexico | ERS6574743 |
| 201707483 | 2017 | France | Metropole | Unknown | ERS6574744 |
| 201707500 | 2017 | France | Metropole | Morocco | ERS6574745 |
| 201707514 | 2017 | France | French Guiana | Unknown | ERS6574746 |
| 201707523 | 2017 | France | Metropole | Unknown | ERS6574747 |
| 201707615 | 2017 | France | Metropole | Algeria | ERS6574749 |
| 201707640 | 2017 | France | Metropole | Unknown | ERS6574751 |
| 201707659 | 2017 | France | Metropole | None | ERS6574752 |
| 201707660 | 2017 | France | Metropole | Morocco | ERS6574753 |
| 201707837 | 2017 | France | Metropole | Unknown | ERS6574754 |
| 201707853 | 2017 | France | Metropole | Morocco | ERS6574755 |
| 201707862 | 2017 | France | Metropole | Vietnam | ERS6574756 |
| 201707913 | 2017 | France | Mayotte | Unknown | ERS6574757 |
| 201707956 | 2017 | France | Metropole | Lebanon | ERS6574758 |

| | | | | | |
|---|---|---|---|---|---|
| 201707971 | 2017 | France | Metropole | Unknown | ERS6574759 |
| 201707976 | 2017 | France | Mayotte | Unknown | ERS6574760 |
| 201707978 | 2017 | France | Mayotte | Unknown | ERS6574761 |
| 201707999 | 2017 | France | Metropole | Unknown | ERS6574762 |
| 201708001 | 2017 | France | Metropole | Morocco | ERS6574763 |
| 201708002 | 2017 | France | Metropole | Vietnam | ERS6574764 |
| 201708016 | 2017 | France | Metropole | Egypt | ERS6574765 |
| 201708020 | 2017 | France | Metropole | Morocco | ERS6574766 |
| 201708044 | 2017 | France | Metropole | Turkey | ERS6574767 |
| 201708050 | 2017 | France | Metropole | Unknown | ERS6574768 |
| 201708055 | 2017 | France | Metropole | Unknown | ERS6574769 |
| 201708112 | 2017 | France | Metropole | Unknown | ERS6574770 |
| 201708115 | 2017 | France | Metropole | India | ERS6574771 |
| 201708157 | 2017 | France | Metropole | Unknown | ERS6574772 |
| 201708158 | 2017 | France | Metropole | Morocco | ERS6574773 |
| 201708165 | 2017 | France | Metropole | Algeria | ERS6574774 |
| 201708167 | 2017 | France | Metropole | None | ERS6574775 |
| 201708171 | 2017 | France | Metropole | None | ERS6574776 |
| 201708198 | 2017 | France | Metropole | Unknown | ERS6574778 |
| 201708278 | 2017 | France | Metropole | Unknown | ERS6574779 |
| 201708289 | 2017 | France | Metropole | Unknown | ERS6574780 |
| 201708297 | 2017 | France | Metropole | None | ERS6574781 |
| 201708298 | 2017 | France | Metropole | Morocco | ERS6574782 |
| 201708332 | 2017 | France | Metropole | Morocco | ERS6574783 |
| 201708393 | 2017 | France | Metropole | Croatia | ERS6574784 |
| 201708474 | 2017 | France | Metropole | Algeria | ERS6574785 |
| 201708475 | 2017 | France | Metropole | Algeria | ERS6574786 |
| 201708477 | 2017 | France | Metropole | Unknown | ERS6574787 |
| 201708491 | 2017 | France | Metropole | Unknown | ERS6574788 |
| 201708494 | 2017 | France | Metropole | Spain | ERS6574789 |
| 201708495 | 2017 | France | Metropole | Morocco | ERS6574790 |
| 201708496 | 2017 | France | French Guiana | Unknown | ERS6574791 |
| 201708625 | 2017 | France | Metropole | Egypt | ERS6574792 |
| 201708646 | 2017 | France | Metropole | Tunisia | ERS6574793 |
| 201708709 | 2017 | France | Metropole | Unknown | ERS6574794 |
| 201708710 | 2017 | France | Metropole | Unknown | ERS6574921 |
| 201708712 | 2017 | France | Metropole | Unknown | ERS6575923 |
| 201708713 | 2017 | France | Metropole | Unknown | ERS6574922 |
| 201708716 | 2017 | France | Metropole | None | ERS6574923 |
| 201708788 | 2017 | France | Metropole | Unknown | ERS6574924 |
| 201708829 | 2017 | France | Metropole | Haiti | ERS6574925 |
| 201708971 | 2017 | France | Metropole | Algeria | ERS6574926 |
| 201708972 | 2017 | France | Metropole | Unknown | ERS6574927 |
| 201708974 | 2017 | France | Metropole | Pakistan | ERS6574928 |
| 201708975 | 2017 | France | Metropole | Unknown | ERS6574929 |
| 201708977 | 2017 | France | Metropole | Portugal | ERS6574930 |
| 201708982 | 2017 | France | Metropole | Gambia | ERS6574931 |
| 201708983 | 2017 | France | Metropole | Unknown | ERS6574932 |
| 201708984 | 2017 | France | Metropole | Unknown | ERS6574933 |
| 201708985 | 2017 | France | Metropole | Morocco | ERS6574934 |
| 201708986 | 2017 | France | Metropole | Lebanon | ERS6574935 |
| 201709008 | 2017 | France | Metropole | None | ERS6575924 |
| 201709009 | 2017 | France | Metropole | None | ERS6574936 |
| 201709052 | 2017 | France | Metropole | Unknown | ERS6574937 |
| 201709054 | 2017 | France | Metropole | Algeria | ERS6574938 |
| 201709059 | 2017 | France | Metropole | Tunisia | ERS6574939 |
| 201709145 | 2017 | France | Metropole | Algeria | ERS6574940 |
| 201709146 | 2017 | France | Metropole | Unknown | ERS6574941 |
| 201709151 | 2017 | France | Metropole | None | ERS6574942 |
| 201709155 | 2017 | France | Metropole | Unknown | ERS6574795 |
| 201709392 | 2017 | France | Metropole | Unknown | ERS6574796 |
| 201709404 | 2017 | France | Metropole | Spain | ERS6574943 |
| 201709405 | 2017 | France | Metropole | Unknown | ERS6574944 |
| 201709420 | 2017 | France | Metropole | None | ERS6575925 |
| 201709424 | 2017 | France | Metropole | Unknown | ERS6574945 |
| 201709456 | 2017 | France | Metropole | Unknown | ERS6574946 |
| 201709461 | 2017 | France | Metropole | None | ERS6574947 |
| 201709496 | 2017 | France | Metropole | Unknown | ERS6574948 |
| 201709497 | 2017 | France | Metropole | Unknown | ERS6574949 |
| 201709513 | 2017 | France | Metropole | Unknown | ERS6574950 |
| 201709514 | 2017 | France | Metropole | Unknown | ERS6574951 |
| 201709516 | 2017 | France | Metropole | Unknown | ERS6574952 |
| 201709517 | 2017 | France | Metropole | Morocco | ERS6574953 |
| 201709518 | 2017 | France | Metropole | Saudi Arabia | ERS6574954 |
| 201709519 | 2017 | France | Metropole | Unknown | ERS6574955 |
| 201709550 | 2017 | France | Metropole | Africa | ERS6574956 |
| 201709582 | 2017 | France | Metropole | Turkey | ERS6574957 |
| 201709587 | 2017 | France | Metropole | Unknown | ERS6574958 |
| 201709588 | 2017 | France | Metropole | Unknown | ERS6574959 |
| 201709636 | 2017 | France | Metropole | Unknown | ERS6574797 |
| 201709638 | 2017 | France | Metropole | Morocco | ERS6574798 |
| 201709655 | 2017 | France | Metropole | Algeria | ERS6574799 |
| 201709678 | 2017 | France | Metropole | Greece | ERS6574800 |
| 201709693 | 2017 | France | Metropole | Unknown | ERS6574801 |
| 201709694 | 2017 | France | Metropole | Unknown | ERS6574802 |
| 201709695 | 2017 | France | Metropole | Unknown | ERS6574803 |
| 201709696 | 2017 | France | Metropole | Unknown | ERS6574804 |
| 201709697 | 2017 | France | Metropole | Unknown | ERS6574805 |
| 201709723 | 2017 | France | Metropole | Unknown | ERS6574806 |
| 201709771 | 2017 | France | Metropole | Algeria | ERS6574807 |
| 201709775 | 2017 | France | Metropole | Unknown | ERS6574808 |
| 201709776 | 2017 | France | Metropole | Unknown | ERS6574809 |
| 201709779 | 2017 | France | Metropole | Romania | ERS6574810 |
| 201709780 | 2017 | France | Metropole | Unknown | ERS6574811 |
| 201709798 | 2017 | France | Metropole | Unknown | ERS6574812 |
| 201709817 | 2017 | France | Metropole | Unknown | ERS6574813 |
| 201709837 | 2017 | France | Metropole | Unknown | ERS6574814 |
| 201709838 | 2017 | France | Metropole | Unknown | ERS6574815 |
| 201709947 | 2017 | France | Metropole | Unknown | ERS6574817 |
| 201709970 | 2017 | France | Metropole | Unknown | ERS6574818 |
| 201709974 | 2017 | France | Metropole | Senegal | ERS6574819 |
| 201709975 | 2017 | France | Metropole | Unknown | ERS6574820 |
| 201709976 | 2017 | France | Metropole | Unknown | ERS6574821 |

| | | | | | |
|---|---|---|---|---|---|
| 201709977 | 2017 | France | Metropole | Unknown | ERS6574822 |
| 201710011 | 2017 | France | Metropole | Morocco | ERS6574823 |
| 201710068 | 2017 | France | Metropole | Morocco | ERS6574824 |
| 201710098 | 2017 | France | Metropole | Spain | ERS6574825 |
| 201710119 | 2017 | France | Metropole | Unknown | ERS6574826 |
| 201710120 | 2017 | France | Metropole | Morocco | ERS6574827 |
| 201710121 | 2017 | France | Metropole | None | ERS6574828 |
| 201710165 | 2017 | France | Metropole | Unknown | ERS6574829 |
| 201710197 | 2017 | France | Metropole | Unknown | ERS6574830 |
| 201710214 | 2017 | France | Metropole | Unknown | ERS6574831 |
| 201710232 | 2017 | France | Metropole | Unknown | ERS6574832 |
| 201710248 | 2017 | France | Metropole | Unknown | ERS6574833 |
| 201710257 | 2017 | France | Metropole | Cape Verde | ERS6574834 |
| 201710294 | 2017 | France | Metropole | Unknown | ERS6575926 |
| 201710303 | 2017 | France | Metropole | Unknown | ERS6574835 |
| 201710326 | 2017 | France | Metropole | Tunisia | ERS6574836 |
| 201710327 | 2017 | France | Metropole | Unknown | ERS6574837 |
| 201710354 | 2017 | France | Metropole | Morocco | ERS6574838 |
| 201710380 | 2017 | France | Metropole | Unknown | ERS6574839 |
| 201710385 | 2017 | France | Metropole | None | ERS6574840 |
| 201710399 | 2017 | France | Metropole | Unknown | ERS6574841 |
| 201710406 | 2017 | France | Metropole | Unknown | ERS6574842 |
| 201710497 | 2017 | France | Metropole | Morocco | ERS6574843 |
| 201710499 | 2017 | France | Metropole | None | ERS6574844 |
| 201710501 | 2017 | France | Metropole | Unknown | ERS6574845 |
| 201710524 | 2017 | France | Metropole | Uzbekistan | ERS6574846 |
| 201710529 | 2017 | France | Metropole | Unknown | ERS6574847 |
| 201710544 | 2017 | France | Metropole | Spain | ERS6574848 |
| 201710655 | 2017 | France | Metropole | Ukraine | ERS6574960 |
| 201710658 | 2017 | France | Metropole | Unknown | ERS6574961 |
| 201710661 | 2017 | France | Metropole | Unknown | ERS6574962 |
| 201710722 | 2017 | France | Metropole | Unknown | ERS6574963 |
| 201710723 | 2017 | France | Metropole | Morocco | ERS6574964 |
| 201710792 | 2017 | France | Metropole | Unknown | ERS6574965 |
| 201710832 | 2017 | France | Metropole | Unknown | ERS6574966 |
| 201710833 | 2017 | France | Metropole | Unknown | ERS6574967 |
| 201710834 | 2017 | France | Metropole | Unknown | ERS6574968 |
| 201710852 | 2017 | France | Metropole | None | ERS6574969 |
| 201710853 | 2017 | France | Metropole | Unknown | ERS6574970 |
| 201710873 | 2017 | France | Metropole | Mexico | ERS6574971 |
| 201710888 | 2017 | France | Metropole | Unknown | ERS6574972 |
| 201710982 | 2017 | France | Metropole | Unknown | ERS6574973 |
| 201711012 | 2017 | France | Metropole | Unknown | ERS6574974 |
| 201711035 | 2017 | France | Metropole | None | ERS6574975 |
| 201711072 | 2017 | France | Metropole | Cape Verde | ERS6574976 |
| 201711083 | 2017 | France | Metropole | India | ERS6574977 |
| 201711084 | 2017 | France | Metropole | Unknown | ERS6574978 |
| 201711085 | 2017 | France | Metropole | Unknown | ERS6574979 |
| 201711086 | 2017 | France | Metropole | Cuba | ERS6574980 |
| 201711087 | 2017 | France | Metropole | Indonesia | ERS6574981 |
| 201711088 | 2017 | France | Metropole | Morocco | ERS6574982 |
| 201711089 | 2017 | France | Metropole | Unknown | ERS6574983 |
| 201711114 | 2017 | France | Metropole | Unknown | ERS6574984 |
| 201711143 | 2017 | France | Metropole | Unknown | ERS6574985 |
| 201711175 | 2017 | France | Metropole | Morocco | ERS6574986 |
| 201711202 | 2017 | France | Metropole | Morocco | ERS6574987 |
| 201711219 | 2017 | France | Metropole | Morocco | ERS6575927 |
| 201711220 | 2017 | France | Metropole | Morocco | ERS6574988 |
| 201711262 | 2017 | France | Metropole | None | ERS6575928 |
| 201711285 | 2017 | France | French Guiana | Unknown | ERS6574989 |
| 201711286 | 2017 | France | French Guiana | Unknown | ERS6575192 |
| 201711287 | 2017 | France | French Guiana | Unknown | ERS6574990 |
| 201711291 | 2017 | France | Metropole | Tunisia | ERS6575193 |
| 201711299 | 2017 | France | Metropole | Dominican Republic | ERS6575194 |
| 201711318 | 2017 | France | Metropole | None | ERS6574991 |
| 201711319 | 2017 | France | Metropole | India | ERS6574992 |
| 201711321 | 2017 | France | Metropole | Unknown | ERS6575195 |
| 201711329 | 2017 | France | Metropole | Haiti | ERS6574993 |
| 201711355 | 2017 | France | Metropole | Dominican Republic | ERS6575196 |
| 201711356 | 2017 | France | Metropole | Morocco | ERS6574994 |
| 201711388 | 2017 | France | Metropole | Unknown | ERS6575197 |
| 201711389 | 2017 | France | Metropole | Dominican Republic | ERS6575198 |
| 201711394 | 2017 | France | Metropole | Unknown | ERS6574995 |
| 201711402 | 2017 | France | Metropole | Egypt | ERS6575199 |
| 201711427 | 2017 | France | Metropole | Algeria | ERS6575200 |
| 201711521 | 2017 | France | Metropole | Unknown | ERS6574997 |
| 201711530 | 2017 | France | Metropole | Egypt | ERS6574998 |
| 201711542 | 2017 | France | Metropole | Unknown | ERS6574999 |
| 201711555 | 2017 | France | Metropole | Asia | ERS6575000 |
| 201711556 | 2017 | France | Metropole | Unknown | ERS6575001 |
| 201711574 | 2017 | France | Metropole | Morocco | ERS6575002 |
| 201711612 | 2017 | France | Metropole | Unknown | ERS6575929 |
| 201711621 | 2017 | France | Metropole | India | ERS6575003 |
| 201711622 | 2017 | France | Metropole | Bolivia | ERS6575004 |
| 201711623 | 2017 | France | Metropole | Latin America | ERS6575005 |
| 201711656 | 2017 | France | Metropole | Unknown | ERS6575007 |
| 201711701 | 2017 | France | Mayotte | Unknown | ERS6575008 |
| 201711705 | 2017 | France | Mayotte | Unknown | ERS6575009 |
| 201711706 | 2017 | France | Mayotte | Unknown | ERS6575010 |
| 201711721 | 2017 | France | Metropole | Tunisia | ERS6575011 |
| 201711742 | 2017 | France | Metropole | None | ERS6575012 |
| 201711745 | 2017 | France | French Guiana | Unknown | ERS6575013 |
| 201711771 | 2017 | France | Metropole | None | ERS6575014 |
| 201711793 | 2017 | France | Metropole | Morocco | ERS6575015 |
| 201711794 | 2017 | France | Metropole | Cape Verde | ERS6575016 |
| 201711854 | 2017 | France | Metropole | Cuba | ERS6575017 |
| 201711886 | 2017 | France | Metropole | Unknown | ERS6575018 |
| 201711980 | 2017 | France | Metropole | Unknown | ERS6575019 |
| 201711997 | 2017 | France | Metropole | Unknown | ERS6575020 |
| 201711998 | 2017 | France | Metropole | Unknown | ERS6575021 |
| 201712011 | 2017 | France | French Guiana | Unknown | ERS6575022 |
| 201712082 | 2017 | France | Metropole | None | ERS6575023 |
| 201712098 | 2017 | France | Metropole | Unknown | ERS6575024 |

| | | | | | |
|---|---|---|---|---|---|
| 201712146 | 2017 | France | Metropole | Unknown | ERS6575025 |
| 201800113 | 2017 | France | Metropole | Greece | ERS6575026 |
| 201800199 | 2017 | France | Metropole | Unknown | ERS6575029 |
| 201800136 | 2018 | France | Metropole | Unknown | ERS6575027 |
| 201800191 | 2018 | France | Metropole | None | ERS6575028 |
| 201800290 | 2018 | France | Metropole | Thailand | ERS6575030 |
| 201800302 | 2018 | France | Metropole | Morocco | ERS6575031 |
| 201800327 | 2018 | France | Metropole | Unknown | ERS6575032 |
| 201800332 | 2018 | France | French Guiana | Unknown | ERS6575033 |
| 201800345 | 2018 | France | Metropole | Unknown | ERS6575034 |
| 201800426 | 2018 | France | Metropole | Unknown | ERS6575201 |
| 201800428 | 2018 | France | Metropole | None | ERS6575202 |
| 201800443 | 2018 | France | Metropole | None | ERS6575203 |
| 201800445 | 2018 | France | Metropole | None | ERS6575204 |
| 201800446 | 2018 | France | Metropole | Unknown | ERS6575205 |
| 201800449 | 2018 | France | Metropole | Unknown | ERS6575206 |
| 201800646 | 2018 | France | Metropole | Unknown | ERS6575207 |
| 201800662 | 2018 | France | Metropole | None | ERS6575208 |
| 201800665 | 2018 | France | Metropole | None | ERS6575209 |
| 201800729 | 2018 | France | Metropole | Unknown | ERS6575210 |
| 201800760 | 2018 | France | Metropole | Unknown | ERS12446239 |
| 201800794 | 2018 | France | Metropole | Unknown | ERS6575211 |
| 201800795 | 2018 | France | Metropole | Indonesia | ERS6492765 |
| 201800796 | 2018 | France | Metropole | Cuba | ERS6575930 |
| 201800834 | 2018 | France | Metropole | Egypt | ERS6575212 |
| 201800835 | 2018 | France | Metropole | None | ERS6575213 |
| 201800864 | 2018 | France | Metropole | Unknown | ERS6575214 |
| 201800879 | 2018 | France | Metropole | Unknown | ERS6575215 |
| 201800881 | 2018 | France | Metropole | Senegal | ERS6575216 |
| 201800926 | 2018 | France | Metropole | Unknown | ERS6575217 |
| 201800948 | 2018 | France | Metropole | Togo | ERS6575218 |
| 201801077 | 2018 | France | Metropole | Unknown | ERS6575219 |
| 201801137 | 2018 | France | Metropole | Unknown | ERS6575220 |
| 201801162 | 2018 | France | Metropole | Unknown | ERS6492766 |
| 201801191 | 2018 | France | Metropole | Unknown | ERS6575221 |
| 201801276 | 2018 | France | Metropole | Unknown | ERS6575223 |
| 201801294 | 2018 | France | Metropole | Unknown | ERS6575224 |
| 201801462 | 2018 | France | Metropole | Unknown | ERS6575225 |
| 201801463 | 2018 | France | Metropole | None | ERS6575226 |
| 201801473 | 2018 | France | Metropole | Mexico | ERS6575227 |
| 201801603 | 2018 | France | Metropole | Dominican Republic | ERS6575228 |
| 201801620 | 2018 | France | Metropole | Unknown | ERS6575229 |
| 201801648 | 2018 | France | Metropole | India | ERS6575230 |
| 201801672 | 2018 | France | Metropole | Unknown | ERS6492767 |
| 201801676 | 2018 | France | Metropole | India | ERS6575231 |
| 201801678 | 2018 | France | Metropole | Bolivia | ERS6575232 |
| 201801683 | 2018 | France | Metropole | India | ERS6575233 |
| 201801897 | 2018 | France | Metropole | India | ERS6575234 |
| 201801912 | 2018 | France | Metropole | India | ERS6575235 |
| 201801924 | 2018 | France | Metropole | Saudi Arabia | ERS6492768 |
| 201801951 | 2018 | France | Metropole | Egypt | ERS6575931 |
| 201801985 | 2018 | France | Metropole | Unknown | ERS6575932 |
| 201801998 | 2018 | France | Metropole | India | ERS6575933 |
| 201802079 | 2018 | France | Metropole | Netherlands | ERS6575934 |
| 201802081 | 2018 | France | Metropole | Unknown | ERS6575935 |
| 201802184 | 2018 | France | Metropole | Unknown | ERS6575936 |
| 201802242 | 2018 | France | Metropole | Sri Lanka | ERS6575236 |
| 201802243 | 2018 | France | Metropole | Unknown | ERS6575237 |
| 201802245 | 2018 | France | Metropole | Unknown | ERS6492769 |
| 201802246 | 2018 | France | Metropole | Vietnam | ERS6575238 |
| 201802247 | 2018 | France | Metropole | Sri Lanka | ERS6575239 |
| 201802293 | 2018 | France | Metropole | Unknown | ERS6575240 |
| 201802294 | 2018 | France | Metropole | Unknown | ERS6575241 |
| 201802339 | 2018 | France | Metropole | Unknown | ERS6575242 |
| 201802398 | 2018 | France | Metropole | India | ERS6575243 |
| 201802430 | 2018 | France | Metropole | Unknown | ERS6575244 |
| 201802433 | 2018 | France | Metropole | Unknown | ERS6575245 |
| 201802494 | 2018 | France | Metropole | Unknown | ERS6575246 |
| 201802495 | 2018 | France | Metropole | Unknown | ERS6575247 |
| 201802557 | 2018 | France | Metropole | Unknown | ERS6575248 |
| 201802590 | 2018 | France | Metropole | Unknown | ERS6575249 |
| 201802644 | 2018 | France | Metropole | Unknown | ERS6575250 |
| 201802686 | 2018 | France | Metropole | Unknown | ERS6575251 |
| 201802687 | 2018 | France | Metropole | Unknown | ERS6575252 |
| 201802820 | 2018 | France | Metropole | Bolivia | ERS6575253 |
| 201802851 | 2018 | France | Metropole | None | ERS6575254 |
| 201802864 | 2018 | France | Metropole | Unknown | ERS6575255 |
| 201802871 | 2018 | France | Metropole | Colombia | ERS6575256 |
| 201802905 | 2018 | France | Metropole | Pakistan | ERS6575257 |
| 201802937 | 2018 | France | Metropole | Unknown | ERS6575258 |
| 201802944 | 2018 | France | Metropole | Peru | ERS6575259 |
| 201802973 | 2018 | France | Metropole | Unknown | ERS6575260 |
| 201802978 | 2018 | France | Metropole | Unknown | ERS6575261 |
| 201803032 | 2018 | France | French Guiana | Unknown | ERS6575262 |
| 201803033 | 2018 | France | French Guiana | Unknown | ERS6575263 |
| 201803102 | 2018 | France | Metropole | Unknown | ERS6492770 |
| 201803216 | 2018 | France | Metropole | Unknown | ERS6575264 |
| 201803257 | 2018 | France | Metropole | Unknown | ERS6575265 |
| 201803260 | 2018 | France | Metropole | Peru | ERS6575266 |
| 201803306 | 2018 | France | Metropole | Unknown | ERS6575267 |
| 201803368 | 2018 | France | Metropole | India | ERS6575268 |
| 201803370 | 2018 | France | Metropole | Unknown | ERS6575269 |
| 201803384 | 2018 | France | Metropole | Asia | ERS6575270 |
| 201803414 | 2018 | France | Metropole | Unknown | ERS6575271 |
| 201803416 | 2018 | France | Metropole | Indonesia | ERS6575272 |
| 201803437 | 2018 | France | French Guiana | Unknown | ERS6575273 |
| 201803440 | 2018 | France | French Guiana | Unknown | ERS6575274 |
| 201803442 | 2018 | France | French Guiana | Unknown | ERS6575275 |
| 201803443 | 2018 | France | French Guiana | Unknown | ERS6575276 |
| 201803466 | 2018 | France | Metropole | India | ERS6575277 |
| 201803467 | 2018 | France | Metropole | India | ERS6575278 |
| 201803502 | 2018 | France | Metropole | Unknown | ERS6575279 |
| 201803510 | 2018 | France | Metropole | Unknown | ERS6575280 |

| | | | | | |
|---|---|---|---|---|---|
| 201803511 | 2018 | France | Metropole | Unknown | ERS6575281 |
| 201803536 | 2018 | France | Metropole | India | ERS6575282 |
| 201803633 | 2018 | France | Metropole | Unknown | ERS6575284 |
| 201803654 | 2018 | France | Metropole | Unknown | ERS6575285 |
| 201803687 | 2018 | France | Metropole | Morocco | ERS6575286 |
| 201803734 | 2018 | France | Metropole | Morocco | ERS6575287 |
| 201803773 | 2018 | France | Metropole | None | ERS6575288 |
| 201803790 | 2018 | France | French Guiana | Unknown | ERS6575289 |
| 201803886 | 2018 | France | Metropole | Unknown | ERS6575290 |
| 201803887 | 2018 | France | Metropole | None | ERS6492771 |
| 201803920 | 2018 | France | Metropole | India | ERS6575291 |
| 201803921 | 2018 | France | Metropole | Chad | ERS6575292 |
| 201803955 | 2018 | France | Metropole | Egypt | ERS6575293 |
| 201803980 | 2018 | France | Metropole | Unknown | ERS6575294 |
| 201803991 | 2018 | France | Metropole | Unknown | ERS6575295 |
| 201804007 | 2018 | France | Metropole | Unknown | ERS6575296 |
| 201804146 | 2018 | France | Metropole | None | ERS6575297 |
| 201804166 | 2018 | France | Metropole | Morocco | ERS6575298 |
| 201804168 | 2018 | France | Metropole | America | ERS6575299 |
| 201804196 | 2018 | France | Metropole | None | ERS6575300 |
| 201804244 | 2018 | France | Mayotte | Unknown | ERS6575301 |
| 201804247 | 2018 | France | Mayotte | Unknown | ERS6575302 |
| 201804248 | 2018 | France | Mayotte | Unknown | ERS6575303 |
| 201804249 | 2018 | France | Mayotte | Unknown | ERS6575304 |
| 201804251 | 2018 | France | Mayotte | Unknown | ERS6575305 |
| 201804252 | 2018 | France | Mayotte | Unknown | ERS6575306 |
| 201804254 | 2018 | France | Mayotte | Unknown | ERS6575307 |
| 201804274 | 2018 | France | Metropole | Dominican Republic | ERS6575308 |
| 201804275 | 2018 | France | Metropole | Unknown | ERS6575309 |
| 201804277 | 2018 | France | Metropole | Portugal | ERS6575310 |
| 201804488 | 2018 | France | Metropole | Côte d’Ivoire | ERS6575311 |
| 201804490 | 2018 | France | Metropole | Unknown | ERS6575312 |
| 201804491 | 2018 | France | Metropole | Colombia | ERS6575313 |
| 201804492 | 2018 | France | Metropole | Unknown | ERS6575314 |
| 201804494 | 2018 | France | Metropole | India | ERS6575315 |
| 201804496 | 2018 | France | Metropole | Unknown | ERS6575316 |
| 201804499 | 2018 | France | Metropole | Spain | ERS6575318 |
| 201804500 | 2018 | France | Metropole | Brazil | ERS6575319 |
| 201804501 | 2018 | France | Metropole | Unknown | ERS6575320 |
| 201804530 | 2018 | France | Metropole | Unknown | ERS6575321 |
| 201804667 | 2018 | France | Metropole | None | ERS6575322 |
| 201804701 | 2018 | France | Metropole | Unknown | ERS6492772 |
| 201804702 | 2018 | France | Metropole | Unknown | ERS6575323 |
| 201804737 | 2018 | France | Metropole | None | ERS6492773 |
| 201804778 | 2018 | France | Metropole | India | ERS6575324 |
| 201804820 | 2018 | France | Metropole | Morocco | ERS6575325 |
| 201804829 | 2018 | France | Metropole | Bulgaria | ERS6575326 |
| 201804938 | 2018 | France | Metropole | Norway | ERS6575327 |
| 201804965 | 2018 | France | Metropole | Unknown | ERS6575328 |
| 201804969 | 2018 | France | Metropole | None | ERS6492774 |
| 201804970 | 2018 | France | Metropole | None | ERS6492775 |
| 201804971 | 2018 | France | Metropole | None | ERS6575329 |
| 201804989 | 2018 | France | Metropole | None | ERS6575330 |
| 201804993 | 2018 | France | Metropole | Unknown | ERS6575331 |
| 201804994 | 2018 | France | Metropole | Unknown | ERS6575332 |
| 201805012 | 2018 | France | Metropole | Unknown | ERS6575333 |
| 201805013 | 2018 | France | Metropole | Unknown | ERS6575334 |
| 201805028 | 2018 | France | Metropole | Unknown | ERS6575335 |
| 201805051 | 2018 | France | Metropole | Unknown | ERS6575336 |
| 201805144 | 2018 | France | Metropole | Unknown | ERS6575337 |
| 201805166 | 2018 | France | French Guiana | Unknown | ERS6575338 |
| 201805174 | 2018 | France | Metropole | Unknown | ERS6575339 |
| 201805223 | 2018 | France | French Guiana | Unknown | ERS6575340 |
| 201805224 | 2018 | France | French Guiana | Unknown | ERS6575341 |
| 201805226 | 2018 | France | French Guiana | Unknown | ERS6575342 |
| 201805227 | 2018 | France | French Guiana | Unknown | ERS6575343 |
| 201805242 | 2018 | France | Metropole | None | ERS6575344 |
| 201805312 | 2018 | France | Metropole | None | ERS6492776 |
| 201805313 | 2018 | France | Metropole | None | ERS6492777 |
| 201805338 | 2018 | France | Metropole | Unknown | ERS6575345 |
| 201805366 | 2018 | France | Metropole | Unknown | ERS6575346 |
| 201805373 | 2018 | France | Metropole | Unknown | ERS6575347 |
| 201805386 | 2018 | France | Metropole | None | ERS6575348 |
| 201805409 | 2018 | France | Metropole | Unknown | ERS6575349 |
| 201805411 | 2018 | France | Guadeloupe | Unknown | ERS6575350 |
| 201805434 | 2018 | France | Metropole | Unknown | ERS6575351 |
| 201805436 | 2018 | France | Metropole | Unknown | ERS6575352 |
| 201805463 | 2018 | France | Metropole | Egypt | ERS6575353 |
| 201805486 | 2018 | France | Metropole | Argentina | ERS6575354 |
| 201805506 | 2018 | France | Metropole | Russia | ERS6575355 |
| 201805522 | 2018 | France | Metropole | None | ERS6492778 |
| 201805523 | 2018 | France | Metropole | None | ERS6492779 |
| 201805524 | 2018 | France | Metropole | None | ERS6492780 |
| 201805526 | 2018 | France | Metropole | None | ERS6492781 |
| 201805529 | 2018 | France | Metropole | Egypt | ERS6575356 |
| 201805547 | 2018 | France | Metropole | Egypt | ERS6575357 |
| 201805549 | 2018 | France | Metropole | Peru | ERS6575358 |
| 201805559 | 2018 | France | Metropole | Unknown | ERS6575359 |
| 201805599 | 2018 | France | Metropole | Colombia | ERS6575360 |
| 201805633 | 2018 | France | Metropole | None | ERS6575361 |
| 201805655 | 2018 | France | Metropole | None | ERS6492782 |
| 201805674 | 2018 | France | Metropole | None | ERS6575362 |
| 201805675 | 2018 | France | Metropole | None | ERS6575363 |
| 201805687 | 2018 | France | Metropole | Unknown | ERS6575364 |
| 201805694 | 2018 | France | Metropole | None | ERS6575365 |
| 201805695 | 2018 | France | Metropole | Unknown | ERS6575366 |
| 201805729 | 2018 | France | Metropole | Togo | ERS6575367 |
| 201805730 | 2018 | France | Metropole | Cuba | ERS6575368 |
| 201805739 | 2018 | France | Metropole | Unknown | ERS6575369 |
| 201805759 | 2018 | France | Metropole | Unknown | ERS6575370 |
| 201805762 | 2018 | France | Metropole | Spain | ERS6575371 |
| 201805829 | 2018 | France | Metropole | Unknown | ERS6575372 |
| 201805830 | 2018 | France | Metropole | Unknown | ERS6575373 |

| | | | | | |
|---|---|---|---|---|---|
| 201805843 | 2018 | France | Metropole | Cambodia | ERS6575374 |
| 201805875 | 2018 | France | Metropole | Portugal | ERS6575375 |
| 201805894 | 2018 | France | Metropole | Unknown | ERS6575376 |
| 201805917 | 2018 | France | Metropole | None | ERS6492783 |
| 201805918 | 2018 | France | Metropole | None | ERS6492784 |
| 201805919 | 2018 | France | Metropole | None | ERS6492785 |
| 201805920 | 2018 | France | Metropole | None | ERS6492786 |
| 201805921 | 2018 | France | Metropole | None | ERS6492787 |
| 201805928 | 2018 | France | Metropole | Israel | ERS6575418 |
| 201805935 | 2018 | France | Metropole | Unknown | ERS6575419 |
| 201805936 | 2018 | France | Metropole | Unknown | ERS6575420 |
| 201805953 | 2018 | France | Metropole | India | ERS6575421 |
| 201805969 | 2018 | France | Metropole | Côte d'Ivoire | ERS6575422 |
| 201805980 | 2018 | France | Metropole | None | ERS6575423 |
| 201805998 | 2018 | France | Metropole | Unknown | ERS6575424 |
| 201806009 | 2018 | France | Metropole | Carribean | ERS6575425 |
| 201806015 | 2018 | France | French Guiana | Unknown | ERS6575426 |
| 201806027 | 2018 | France | Metropole | Spain | ERS6575427 |
| 201806034 | 2018 | France | Metropole | Unknown | ERS6575428 |
| 201806048 | 2018 | France | Metropole | Unknown | ERS6575429 |
| 201806079 | 2018 | France | Metropole | Unknown | ERS6575430 |
| 201806085 | 2018 | France | Metropole | None | ERS6575431 |
| 201806123 | 2018 | France | Metropole | None | ERS6492788 |
| 201806124 | 2018 | France | Metropole | None | ERS6492789 |
| 201806125 | 2018 | France | Metropole | Unknown | ERS6575432 |
| 201806146 | 2018 | France | Metropole | Senegal | ERS6575433 |
| 201806148 | 2018 | France | Metropole | Unknown | ERS6575434 |
| 201806152 | 2018 | France | Metropole | None | ERS6575435 |
| 201806153 | 2018 | France | Metropole | Unknown | ERS6575436 |
| 201806154 | 2018 | France | Metropole | Unknown | ERS6575437 |
| 201806155 | 2018 | France | Metropole | None | ERS6575438 |
| 201806198 | 2018 | France | Metropole | Indonesia | ERS6575439 |
| 201806200 | 2018 | France | Metropole | Morocco | ERS6575440 |
| 201806203 | 2018 | France | Metropole | Unknown | ERS6575441 |
| 201806204 | 2018 | France | Metropole | Spain | ERS6575442 |
| 201806253 | 2018 | France | Metropole | None | ERS6575443 |
| 201806254 | 2018 | France | Metropole | Tanzania | ERS6575444 |
| 201806256 | 2018 | France | Metropole | None | ERS6575445 |
| 201806264 | 2018 | France | Metropole | Unknown | ERS6575446 |
| 201806284 | 2018 | France | Metropole | None | ERS6575447 |
| 201806287 | 2018 | France | Metropole | Unknown | ERS6575448 |
| 201806295 | 2018 | France | Metropole | Egypt | ERS6575449 |
| 201806296 | 2018 | France | Metropole | Unknown | ERS6575450 |
| 201806298 | 2018 | France | Metropole | None | ERS6575451 |
| 201806300 | 2018 | France | Metropole | Unknown | ERS6575452 |
| 201806306 | 2018 | France | Metropole | Unknown | ERS6575453 |
| 201806318 | 2018 | France | Metropole | Unknown | ERS6575454 |
| 201806319 | 2018 | France | Metropole | Colombia | ERS6575455 |
| 201806322 | 2018 | France | Metropole | Unknown | ERS6575456 |
| 201806411 | 2018 | France | Metropole | None | ERS6575457 |
| 201806415 | 2018 | France | Metropole | Unknown | ERS6575458 |
| 201806421 | 2018 | France | Metropole | Cuba | ERS6575459 |
| 201806433 | 2018 | France | Metropole | None | ERS6575460 |
| 201806467 | 2018 | France | Metropole | Togo | ERS6575461 |
| 201806524 | 2018 | France | Metropole | Unknown | ERS6575462 |
| 201806529 | 2018 | France | Metropole | None | ERS6575463 |
| 201806556 | 2018 | France | Metropole | None | ERS6492790 |
| 201806560 | 2018 | France | Metropole | Unknown | ERS6575464 |
| 201806597 | 2018 | France | Metropole | Unknown | ERS6575465 |
| 201806612 | 2018 | France | Metropole | Côte d'Ivoire | ERS6575466 |
| 201806629 | 2018 | France | Metropole | Morocco | ERS6575467 |
| 201806658 | 2018 | France | Metropole | Unknown | ERS6575468 |
| 201806666 | 2018 | France | Metropole | India | ERS6575469 |
| 201806668 | 2018 | France | Metropole | India | ERS6575470 |
| 201806673 | 2018 | France | Metropole | None | ERS6492791 |
| 201806686 | 2018 | France | Metropole | Unknown | ERS6575471 |
| 201806688 | 2018 | France | Metropole | None | ERS6575472 |
| 201806721 | 2018 | France | Metropole | Morocco | ERS6575473 |
| 201806745 | 2018 | France | Metropole | Unknown | ERS6575474 |
| 201806775 | 2018 | France | Metropole | Unknown | ERS6575475 |
| 201806776 | 2018 | France | Metropole | Unknown | ERS6575476 |
| 201806812 | 2018 | France | Metropole | India | ERS6575477 |
| 201806824 | 2018 | France | Metropole | None | ERS6492792 |
| 201806825 | 2018 | France | Metropole | Sri Lanka | ERS6492793 |
| 201806827 | 2018 | France | Metropole | None | ERS6492794 |
| 201806828 | 2018 | France | Metropole | Morocco | ERS6575478 |
| 201806836 | 2018 | France | Metropole | Unknown | ERS6575479 |
| 201806851 | 2018 | France | Metropole | Haiti | ERS6575480 |
| 201806890 | 2018 | France | Metropole | Madagascar | ERS6575481 |
| 201806891 | 2018 | France | Metropole | Unknown | ERS6575482 |
| 201806894 | 2018 | France | Metropole | Unknown | ERS6575483 |
| 201806907 | 2018 | France | Metropole | Africa | ERS6575484 |
| 201806908 | 2018 | France | Metropole | Unknown | ERS6575485 |
| 201806929 | 2018 | France | Metropole | Morocco | ERS6575486 |
| 201806954 | 2018 | France | Metropole | Dominican Republic | ERS6575487 |
| 201806959 | 2018 | France | Metropole | None | ERS6575488 |
| 201807014 | 2018 | France | Metropole | Morocco | ERS6575489 |
| 201807017 | 2018 | France | Metropole | Netherlands | ERS6575490 |
| 201807021 | 2018 | France | Metropole | Morocco | ERS6575492 |
| 201807026 | 2018 | France | Metropole | Unknown | ERS6575493 |
| 201807044 | 2018 | France | Metropole | None | ERS6575494 |
| 201807083 | 2018 | France | Metropole | Cape Verde | ERS6575495 |
| 201807085 | 2018 | France | Metropole | Unknown | ERS6575496 |
| 201807086 | 2018 | France | Metropole | Unknown | ERS6575497 |
| 201807117 | 2018 | France | Metropole | India | ERS6575498 |
| 201807119 | 2018 | France | Metropole | Mexico | ERS6575499 |
| 201807120 | 2018 | France | Metropole | Algeria | ERS6575500 |
| 201807133 | 2018 | France | Metropole | Unknown | ERS6575501 |
| 201807173 | 2018 | France | Metropole | Unknown | ERS6575502 |
| 201807176 | 2018 | France | Metropole | Unknown | ERS6575503 |
| 201807186 | 2018 | France | Metropole | None | ERS6575504 |
| 201807190 | 2018 | France | Metropole | Algeria | ERS6575505 |
| 201807205 | 2018 | France | Metropole | None | ERS6575506 |

| | | | | | |
|---|---|---|---|---|---|
| 201807206 | 2018 | France | Metropole | None | ERS6575507 |
| 201807208 | 2018 | France | Metropole | None | ERS6575508 |
| 201807210 | 2018 | France | Metropole | Unknown | ERS6575509 |
| 201807278 | 2018 | France | Metropole | Egypt | ERS6575510 |
| 201807279 | 2018 | France | Metropole | Unknown | ERS6575511 |
| 201807280 | 2018 | France | Metropole | Burkina Faso | ERS6575512 |
| 201807350 | 2018 | France | Metropole | Egypt | ERS6575513 |
| 201807397 | 2018 | France | Metropole | Unknown | ERS6575514 |
| 201807398 | 2018 | France | Metropole | Unknown | ERS6575515 |
| 201807409 | 2018 | France | Metropole | Unknown | ERS6575516 |
| 201807410 | 2018 | France | Metropole | Unknown | ERS6575517 |
| 201807411 | 2018 | France | Metropole | Brazil | ERS6575518 |
| 201807432 | 2018 | France | Metropole | Unknown | ERS6575519 |
| 201807437 | 2018 | France | Metropole | Bolivia | ERS6575520 |
| 201807458 | 2018 | France | Metropole | Unknown | ERS6575521 |
| 201807502 | 2018 | France | Metropole | Morocco | ERS6575522 |
| 201807503 | 2018 | France | Metropole | Morocco | ERS6575523 |
| 201807505 | 2018 | France | Metropole | Morocco | ERS6575524 |
| 201807507 | 2018 | France | Metropole | Africa | ERS6575525 |
| 201807524 | 2018 | France | Metropole | Cape Verde | ERS6575526 |
| 201807525 | 2018 | France | Metropole | Senegal | ERS6575527 |
| 201807556 | 2018 | France | Metropole | Morocco | ERS6575528 |
| 201807557 | 2018 | France | Metropole | Thailand | ERS6492795 |
| 201807560 | 2018 | France | Metropole | Unknown | ERS6575529 |
| 201807561 | 2018 | France | Metropole | Spain | ERS6575530 |
| 201807563 | 2018 | France | Metropole | None | ERS6575531 |
| 201807566 | 2018 | France | Metropole | Unknown | ERS6575532 |
| 201807567 | 2018 | France | Metropole | Unknown | ERS6575533 |
| 201807578 | 2018 | France | Metropole | Unknown | ERS6575534 |
| 201807593 | 2018 | France | Metropole | Unknown | ERS6575535 |
| 201807594 | 2018 | France | Metropole | Unknown | ERS6575536 |
| 201807603 | 2018 | France | Metropole | Unknown | ERS6575537 |
| 201807604 | 2018 | France | Metropole | Unknown | ERS6575538 |
| 201807622 | 2018 | France | French Guiana | Unknown | ERS6575539 |
| 201807624 | 2018 | France | Metropole | Unknown | ERS6575540 |
| 201807626 | 2018 | France | Metropole | Unknown | ERS6575541 |
| 201807629 | 2018 | France | Metropole | Morocco | ERS6575542 |
| 201807630 | 2018 | France | Metropole | None | ERS6575543 |
| 201807662 | 2018 | France | Metropole | Unknown | ERS6575544 |
| 201807678 | 2018 | France | Metropole | Netherlands | ERS6575545 |
| 201807679 | 2018 | France | Metropole | Israel | ERS6575546 |
| 201807681 | 2018 | France | Metropole | Unknown | ERS6575547 |
| 201807682 | 2018 | France | Metropole | Unknown | ERS6575548 |
| 201807685 | 2018 | France | Metropole | Unknown | ERS6575549 |
| 201807688 | 2018 | France | Metropole | None | ERS6575550 |
| 201807733 | 2018 | France | Metropole | Africa | ERS6575551 |
| 201807738 | 2018 | France | Metropole | Africa | ERS6575552 |
| 201807750 | 2018 | France | Metropole | Unknown | ERS6575553 |
| 201807769 | 2018 | France | Metropole | Senegal | ERS6575554 |
| 201807774 | 2018 | France | Metropole | India | ERS6575555 |
| 201807789 | 2018 | France | Metropole | Unknown | ERS6575556 |
| 201807826 | 2018 | France | Metropole | Morocco | ERS6575557 |
| 201807828 | 2018 | France | Metropole | None | ERS6575558 |
| 201807829 | 2018 | France | Metropole | Tunisia | ERS6575559 |
| 201807830 | 2018 | France | Metropole | Morocco | ERS6575560 |
| 201807832 | 2018 | France | Metropole | Algeria | ERS6575561 |
| 201807834 | 2018 | France | Metropole | None | ERS6575562 |
| 201807835 | 2018 | France | Metropole | Unknown | ERS6575563 |
| 201807836 | 2018 | France | Metropole | Unknown | ERS6575564 |
| 201807837 | 2018 | France | Metropole | Unknown | ERS6575565 |
| 201807845 | 2018 | France | Metropole | Morocco | ERS6575566 |
| 201807846 | 2018 | France | Metropole | Morocco | ERS6575567 |
| 201807854 | 2018 | France | Metropole | Morocco | ERS6575568 |
| 201807863 | 2018 | France | Metropole | Unknown | ERS6575569 |
| 201807874 | 2018 | France | Metropole | Unknown | ERS6575570 |
| 201807880 | 2018 | France | Metropole | Unknown | ERS6575571 |
| 201807881 | 2018 | France | Metropole | Tunisia | ERS6575572 |
| 201807886 | 2018 | France | Metropole | Unknown | ERS6575573 |
| 201807903 | 2018 | France | Metropole | Unknown | ERS6575574 |
| 201807938 | 2018 | France | Metropole | Unknown | ERS6575575 |
| 201807953 | 2018 | France | Metropole | Benin | ERS6575576 |
| 201807955 | 2018 | France | Metropole | Unknown | ERS6575577 |
| 201807963 | 2018 | France | Metropole | Unknown | ERS6492796 |
| 201807972 | 2018 | France | Metropole | Unknown | ERS6575578 |
| 201808025 | 2018 | France | Metropole | Morocco | ERS6575579 |
| 201808038 | 2018 | France | Metropole | Morocco | ERS6575580 |
| 201808057 | 2018 | France | Metropole | Egypt | ERS6575581 |
| 201808082 | 2018 | France | Metropole | Unknown | ERS6575582 |
| 201808084 | 2018 | France | Metropole | Unknown | ERS6575583 |
| 201808090 | 2018 | France | Metropole | Unknown | ERS6575584 |
| 201808097 | 2018 | France | Metropole | Unknown | ERS6492797 |
| 201808098 | 2018 | France | Metropole | Algeria | ERS6492798 |
| 201808105 | 2018 | France | Metropole | Algeria | ERS6575585 |
| 201808114 | 2018 | France | Metropole | Morocco | ERS6575586 |
| 201808116 | 2018 | France | Metropole | Algeria | ERS6575587 |
| 201808185 | 2018 | France | Metropole | Algeria | ERS6492799 |
| 201808189 | 2018 | France | Metropole | Unknown | ERS6492800 |
| 201808235 | 2018 | France | Metropole | Unknown | ERS6575588 |
| 201808243 | 2018 | France | Metropole | None | ERS6575589 |
| 201808253 | 2018 | France | Metropole | Unknown | ERS6575590 |
| 201808254 | 2018 | France | Metropole | Unknown | ERS6492801 |
| 201808259 | 2018 | France | Metropole | None | ERS6492802 |
| 201808266 | 2018 | France | Metropole | Egypt | ERS6492803 |
| 201808282 | 2018 | France | Metropole | Unknown | ERS6575591 |
| 201808283 | 2018 | France | Metropole | Unknown | ERS6575592 |
| 201808316 | 2018 | France | Metropole | None | ERS6575593 |
| 201808359 | 2018 | France | Metropole | Italy | ERS6492804 |
| 201808398 | 2018 | France | Metropole | Unknown | ERS6575594 |
| 201808410 | 2018 | France | Metropole | Unknown | ERS6575595 |
| 201808411 | 2018 | France | Metropole | Unknown | ERS6575596 |
| 201808420 | 2018 | France | Metropole | Mexico | ERS6575597 |
| 201808429 | 2018 | France | Metropole | Unknown | ERS6575598 |
| 201808451 | 2018 | France | Metropole | Unknown | ERS6575599 |

| | | | | | |
|---|---|---|---|---|---|
| 201808457 | 2018 | France | Metropole | Unknown | ERS6575600 |
| 201808458 | 2018 | France | Metropole | India | ERS6575601 |
| 201808470 | 2018 | France | Metropole | Unknown | ERS6575602 |
| 201808486 | 2018 | France | Metropole | Unknown | ERS6575603 |
| 201808491 | 2018 | France | Metropole | Unknown | ERS6575604 |
| 201808525 | 2018 | France | Metropole | Unknown | ERS6575605 |
| 201808526 | 2018 | France | Metropole | Cameroon | ERS6575606 |
| 201808573 | 2018 | France | Metropole | Unknown | ERS6575607 |
| 201808574 | 2018 | France | Metropole | Unknown | ERS6575608 |
| 201808590 | 2018 | France | Metropole | Unknown | ERS6575609 |
| 201808595 | 2018 | France | Metropole | None | ERS6575610 |
| 201808600 | 2018 | France | Metropole | Morocco | ERS6575611 |
| 201808610 | 2018 | France | Metropole | Unknown | ERS6575612 |
| 201808611 | 2018 | France | Metropole | Unknown | ERS6575613 |
| 201808638 | 2018 | France | Metropole | Tunisia | ERS6575614 |
| 201808663 | 2018 | France | Metropole | Unknown | ERS6492805 |
| 201808684 | 2018 | France | Metropole | Unknown | ERS6575615 |
| 201808692 | 2018 | France | Metropole | Turkey | ERS6492806 |
| 201808736 | 2018 | France | Metropole | Unknown | ERS6492807 |
| 201808737 | 2018 | France | Metropole | Morocco | ERS6492808 |
| 201808757 | 2018 | France | Metropole | Unknown | ERS6492809 |
| 201808758 | 2018 | France | Metropole | Unknown | ERS6492810 |
| 201808759 | 2018 | France | Metropole | United Kingdom | ERS6492811 |
| 201808770 | 2018 | France | Metropole | Unknown | ERS6492812 |
| 201808793 | 2018 | France | Metropole | Unknown | ERS6492813 |
| 201808797 | 2018 | France | Metropole | None | ERS6492814 |
| 201808798 | 2018 | France | Metropole | None | ERS6492815 |
| 201808831 | 2018 | France | Metropole | Madagascar | ERS6492816 |
| 201808832 | 2018 | France | Metropole | Morocco | ERS6492817 |
| 201808833 | 2018 | France | Metropole | Morocco | ERS6492818 |
| 201808840 | 2018 | France | Metropole | Unknown | ERS6492819 |
| 201808845 | 2018 | France | Metropole | Unknown | ERS6492820 |
| 201808859 | 2018 | France | Metropole | Morocco | ERS6492821 |
| 201808860 | 2018 | France | Metropole | Unknown | ERS6492822 |
| 201808905 | 2018 | France | Metropole | Unknown | ERS6492823 |
| 201808974 | 2018 | France | Metropole | Unknown | ERS6492824 |
| 201808998 | 2018 | France | Metropole | None | ERS6492825 |
| 201809053 | 2018 | France | Metropole | Unknown | ERS6492826 |
| 201809061 | 2018 | France | Metropole | Unknown | ERS6492827 |
| 201809062 | 2018 | France | Metropole | Unknown | ERS6492828 |
| 201809068 | 2018 | France | Metropole | India | ERS6492829 |
| 201809069 | 2018 | France | Metropole | Unknown | ERS6492830 |
| 201809074 | 2018 | France | Metropole | Togo | ERS6492831 |
| 201809098 | 2018 | France | Metropole | Algeria | ERS6492832 |
| 201809101 | 2018 | France | Metropole | None | ERS6492833 |
| 201809106 | 2018 | France | Metropole | Unknown | ERS6492834 |
| 201809160 | 2018 | France | Metropole | Unknown | ERS6492835 |
| 201809177 | 2018 | France | Metropole | Unknown | ERS6492836 |
| 201809213 | 2018 | France | Metropole | None | ERS6492837 |
| 201809214 | 2018 | France | Metropole | Unknown | ERS6492838 |
| 201809245 | 2018 | France | Metropole | Unknown | ERS6492839 |
| 201809261 | 2018 | France | Metropole | Unknown | ERS6492840 |
| 201809293 | 2018 | France | Metropole | None | ERS6492841 |
| 201809302 | 2018 | France | Metropole | Unknown | ERS6492842 |
| 201809307 | 2018 | France | Metropole | Madagascar | ERS6492843 |
| 201809320 | 2018 | France | Metropole | Unknown | ERS6492844 |
| 201809321 | 2018 | France | Metropole | Unknown | ERS6492845 |
| 201809330 | 2018 | France | Metropole | Unknown | ERS6492846 |
| 201809339 | 2018 | France | Metropole | Unknown | ERS6492847 |
| 201809344 | 2018 | France | Metropole | Unknown | ERS6492848 |
| 201809355 | 2018 | France | Metropole | None | ERS6492849 |
| 201809451 | 2018 | France | Metropole | Unknown | ERS6492850 |
| 201809502 | 2018 | France | Metropole | India | ERS6492851 |
| 201809507 | 2018 | France | Metropole | Unknown | ERS6492852 |
| 201809509 | 2018 | France | Metropole | India | ERS6492853 |
| 201809511 | 2018 | France | Metropole | Cape Verde | ERS6492854 |
| 201809512 | 2018 | France | Metropole | Unknown | ERS6492855 |
| 201809513 | 2018 | France | Metropole | India | ERS6492856 |
| 201809514 | 2018 | France | Metropole | Cape Verde | ERS6492857 |
| 201809529 | 2018 | France | Metropole | Canada | ERS6492858 |
| 201809532 | 2018 | France | Metropole | Unknown | ERS6492859 |
| 201809538 | 2018 | France | Metropole | None | ERS6492860 |
| 201809539 | 2018 | France | Metropole | None | ERS6492861 |
| 201809585 | 2018 | France | Metropole | Unknown | ERS6492862 |
| 201809588 | 2018 | France | Metropole | Unknown | ERS6492863 |
| 201809604 | 2018 | France | Metropole | India | ERS6492864 |
| 201809616 | 2018 | France | Metropole | Unknown | ERS6492865 |
| 201809631 | 2018 | France | Metropole | Unknown | ERS6492866 |
| 201809642 | 2018 | France | Metropole | Unknown | ERS6492867 |
| 201809661 | 2018 | France | Metropole | Unknown | ERS6492868 |
| 201809682 | 2018 | France | Metropole | Unknown | ERS6492869 |
| 201809693 | 2018 | France | Metropole | Senegal | ERS6492871 |
| 201809713 | 2018 | France | Metropole | Unknown | ERS6492872 |
| 201809753 | 2018 | France | Metropole | Morocco | ERS6492873 |
| 201809756 | 2018 | France | Metropole | Morocco | ERS6492874 |
| 201809766 | 2018 | France | Metropole | Unknown | ERS6492875 |
| 201809774 | 2018 | France | Metropole | None | ERS6492876 |
| 201809783 | 2018 | France | Metropole | Unknown | ERS6492877 |
| 201809795 | 2018 | France | Metropole | Peru | ERS6492878 |
| 201809800 | 2018 | France | Metropole | None | ERS6492879 |
| 201809801 | 2018 | France | Metropole | None | ERS6492880 |
| 201809803 | 2018 | France | Metropole | Unknown | ERS6492881 |
| 201809830 | 2018 | France | Metropole | Unknown | ERS6492882 |
| 201809870 | 2018 | France | Metropole | Algeria | ERS6492883 |
| 201809880 | 2018 | France | Metropole | Morocco | ERS6492884 |
| 201809882 | 2018 | France | Metropole | None | ERS6492885 |
| 201809893 | 2018 | France | Metropole | Unknown | ERS6492886 |
| 201809899 | 2018 | France | Metropole | Unknown | ERS6492887 |
| 201809960 | 2018 | France | Metropole | Unknown | ERS6492888 |
| 201809979 | 2018 | France | Metropole | Egypt | ERS6492889 |
| 201809992 | 2018 | France | Metropole | Algeria | ERS6492890 |
| 201809995 | 2018 | France | Metropole | Morocco | ERS6492891 |
| 201809996 | 2018 | France | Metropole | Unknown | ERS6492892 |

| | | | | | |
|---|---|---|---|---|---|
| 201809997 | 2018 | France | Metropole | Unknown | ERS6492893 |
| 201809998 | 2018 | France | Metropole | Morocco | ERS6492894 |
| 201810021 | 2018 | France | Metropole | Unknown | ERS6492895 |
| 201810034 | 2018 | France | Metropole | Unknown | ERS6492896 |
| 201810035 | 2018 | France | Metropole | Madagascar | ERS6492897 |
| 201810036 | 2018 | France | Metropole | None | ERS6492898 |
| 201810043 | 2018 | France | Metropole | Unknown | ERS6575937 |
| 201810051 | 2018 | France | Metropole | Morocco | ERS6492899 |
| 201810111 | 2018 | France | Metropole | Tunisia | ERS6492901 |
| 201810125 | 2018 | France | Metropole | None | ERS6492902 |
| 201810126 | 2018 | France | Metropole | Unknown | ERS6492903 |
| 201810127 | 2018 | France | Metropole | Algeria | ERS6492904 |
| 201810136 | 2018 | France | Metropole | Unknown | ERS6492905 |
| 201810149 | 2018 | France | Metropole | Benin | ERS6492906 |
| 201810157 | 2018 | France | Metropole | Cameroon | ERS6492907 |
| 201810227 | 2018 | France | Mayotte | Unknown | ERS6492908 |
| 201810231 | 2018 | France | Metropole | Mayotte | ERS6492909 |
| 201810233 | 2018 | France | Mayotte | Unknown | ERS6494743 |
| 201810234 | 2018 | France | Mayotte | Unknown | ERS6494744 |
| 201810238 | 2018 | France | Metropole | Unknown | ERS6492910 |
| 201810240 | 2018 | France | Metropole | Mexico | ERS6492911 |
| 201810271 | 2018 | France | Metropole | Unknown | ERS6492912 |
| 201810274 | 2018 | France | Metropole | Unknown | ERS6492913 |
| 201810312 | 2018 | France | Metropole | None | ERS6492914 |
| 201810319 | 2018 | France | Metropole | Togo | ERS6492915 |
| 201810349 | 2018 | France | Metropole | Unknown | ERS6494745 |
| 201810367 | 2018 | France | Metropole | Dominican Republic | ERS6494746 |
| 201810382 | 2018 | France | Metropole | Cape Verde | ERS6494747 |
| 201810383 | 2018 | France | Metropole | None | ERS6494748 |
| 201810429 | 2018 | France | Metropole | Morocco | ERS6494749 |
| 201810436 | 2018 | France | Metropole | Vietnam | ERS6494750 |
| 201810440 | 2018 | France | Metropole | None | ERS6494751 |
| 201810452 | 2018 | France | Metropole | Unknown | ERS6494752 |
| 201810457 | 2018 | France | Metropole | Unknown | ERS6494753 |
| 201810461 | 2018 | France | Metropole | India | ERS6492916 |
| 201810480 | 2018 | France | Metropole | Unknown | ERS6494754 |
| 201810486 | 2018 | France | Metropole | Mexico | ERS6494755 |
| 201810540 | 2018 | France | Metropole | Unknown | ERS6494756 |
| 201810607 | 2018 | France | Metropole | Unknown | ERS6494757 |
| 201810613 | 2018 | France | Metropole | None | ERS6494758 |
| 201810616 | 2018 | France | Metropole | Unknown | ERS6494759 |
| 201810617 | 2018 | France | Metropole | Unknown | ERS6494760 |
| 201810618 | 2018 | France | Metropole | Unknown | ERS6494761 |
| 201810619 | 2018 | France | Metropole | Unknown | ERS6494762 |
| 201810620 | 2018 | France | Metropole | Unknown | ERS6494763 |
| 201810622 | 2018 | France | Metropole | Unknown | ERS6494764 |
| 201810623 | 2018 | France | Metropole | Unknown | ERS6494765 |
| 201810624 | 2018 | France | Metropole | Unknown | ERS6494766 |
| 201810625 | 2018 | France | Metropole | Unknown | ERS6494767 |
| 201810626 | 2018 | France | Metropole | Unknown | ERS6494768 |
| 201810627 | 2018 | France | Metropole | Unknown | ERS6492917 |
| 201810628 | 2018 | France | Metropole | Unknown | ERS6494769 |
| 201810629 | 2018 | France | Metropole | Unknown | ERS6494770 |
| 201810630 | 2018 | France | Metropole | Unknown | ERS6494771 |
| 201810631 | 2018 | France | Metropole | Unknown | ERS6494772 |
| 201810632 | 2018 | France | Metropole | Unknown | ERS6494773 |
| 201810635 | 2018 | France | Metropole | Unknown | ERS6494774 |
| 201810678 | 2018 | France | Metropole | Algeria | ERS6494775 |
| 201810690 | 2018 | France | Metropole | Unknown | ERS6494776 |
| 201810691 | 2018 | France | Metropole | None | ERS6494777 |
| 201810707 | 2018 | France | Metropole | Unknown | ERS6494778 |
| 201810713 | 2018 | France | Metropole | Algeria | ERS6494779 |
| 201810729 | 2018 | France | Metropole | Unknown | ERS6494780 |
| 201810730 | 2018 | France | Metropole | Unknown | ERS6494781 |
| 201810748 | 2018 | France | Metropole | India | ERS6494782 |
| 201810749 | 2018 | France | Metropole | Albania | ERS6494783 |
| 201810750 | 2018 | France | Metropole | Egypt | ERS6494784 |
| 201810751 | 2018 | France | Metropole | Senegal | ERS6494785 |
| 201810752 | 2018 | France | Metropole | Unknown | ERS6492918 |
| 201810761 | 2018 | France | Metropole | Indonesia | ERS6494786 |
| 201810775 | 2018 | France | Metropole | Unknown | ERS6494787 |
| 201810803 | 2018 | France | Metropole | Unknown | ERS6494788 |
| 201810807 | 2018 | France | Metropole | Unknown | ERS6494789 |
| 201810830 | 2018 | France | Metropole | Unknown | ERS6494790 |
| 201810839 | 2018 | France | Metropole | Unknown | ERS6494791 |
| 201810840 | 2018 | France | Metropole | Unknown | ERS6494792 |
| 201810841 | 2018 | France | Metropole | Unknown | ERS6494793 |
| 201810842 | 2018 | France | Metropole | Unknown | ERS6494794 |
| 201810843 | 2018 | France | Metropole | Unknown | ERS6494795 |
| 201810844 | 2018 | France | Metropole | Unknown | ERS6494796 |
| 201810847 | 2018 | France | Metropole | Unknown | ERS6494799 |
| 201810875 | 2018 | France | Metropole | Unknown | ERS6494800 |
| 201810877 | 2018 | France | Metropole | Unknown | ERS6494801 |
| 201810898 | 2018 | France | Metropole | Unknown | ERS6494802 |
| 201810920 | 2018 | France | Metropole | None | ERS6494803 |
| 201810933 | 2018 | France | Metropole | Unknown | ERS6494804 |
| 201810934 | 2018 | France | Metropole | Unknown | ERS6494805 |
| 201810935 | 2018 | France | Metropole | None | ERS6492919 |
| 201810952 | 2018 | France | Metropole | Unknown | ERS6494806 |
| 201810955 | 2018 | France | Metropole | Unknown | ERS6575938 |
| 201810975 | 2018 | France | Metropole | Unknown | ERS6494807 |
| 201811026 | 2018 | France | Metropole | Unknown | ERS6494808 |
| 201811036 | 2018 | France | Metropole | Unknown | ERS6494809 |
| 201811048 | 2018 | France | Metropole | None | ERS6494810 |
| 201811064 | 2018 | France | Metropole | Colombia | ERS6494811 |
| 201811106 | 2018 | France | Metropole | Jordan | ERS6494812 |
| 201811114 | 2018 | France | Metropole | Spain | ERS6494813 |
| 201811116 | 2018 | France | Metropole | India | ERS6494814 |
| 201811118 | 2018 | France | Metropole | None | ERS6492920 |
| 201811119 | 2018 | France | Metropole | Myanmar | ERS6494815 |
| 201811120 | 2018 | France | Metropole | Cape Verde | ERS6494816 |
| 201811123 | 2018 | France | Metropole | None | ERS6494817 |
| 201811157 | 2018 | France | Metropole | Unknown | ERS6494818 |

| | | | | | |
|---|---|---|---|---|---|
| 201811197 | 2018 | France | Metropole | Unknown | ERS6576212 |
| 201811207 | 2018 | France | Metropole | Unknown | ERS6494819 |
| 201811228 | 2018 | France | Metropole | Algeria | ERS6494820 |
| 201811250 | 2018 | France | Metropole | Unknown | ERS6494821 |
| 201811257 | 2018 | France | Metropole | Unknown | ERS6494822 |
| 201811303 | 2018 | France | Metropole | India | ERS6494823 |
| 201811341 | 2018 | France | French Guiana | Unknown | ERS6494824 |
| 201811367 | 2018 | France | Metropole | Unknown | ERS6494825 |
| 201811368 | 2018 | France | Metropole | None | ERS6494826 |
| 201811405 | 2018 | France | Metropole | Unknown | ERS6494827 |
| 201811415 | 2018 | France | Metropole | Unknown | ERS6494828 |
| 201811447 | 2018 | France | Metropole | Unknown | ERS6492921 |
| 201811448 | 2018 | France | Metropole | Morocco | ERS6494829 |
| 201811450 | 2018 | France | Metropole | Unknown | ERS6494830 |
| 201811452 | 2018 | France | Metropole | Madagascar | ERS6494831 |
| 201900021 | 2018 | France | Metropole | None | ERS6494832 |
| 201900028 | 2018 | France | Metropole | Unknown | ERS6494833 |
| 201900037 | 2018 | France | Metropole | Portugal | ERS6494834 |
| 201900041 | 2018 | France | Metropole | Unknown | ERS6494835 |
| 201900071 | 2018 | France | Metropole | Mexico | ERS6494837 |
| 201900089 | 2018 | France | Metropole | None | ERS6494838 |
| 201900276 | 2018 | France | Metropole | Unknown | ERS6494841 |
| 201900342 | 2018 | France | French Guiana | Unknown | ERS6494845 |
| 201900346 | 2018 | France | French Guiana | Unknown | ERS6494846 |
| 201900347 | 2018 | France | French Guiana | Unknown | ERS6494847 |
| 201900350 | 2018 | France | French Guiana | Unknown | ERS6494848 |
| 201900594 | 2018 | France | French Guiana | Unknown | ERS6494859 |
| 201900800 | 2018 | France | French Guiana | Unknown | ERS6494874 |
| 201900813 | 2018 | France | Metropole | Unknown | ERS6492922 |
| 201900814 | 2018 | France | Metropole | Unknown | ERS6492923 |
| 201900815 | 2018 | France | Metropole | Unknown | ERS6492924 |
| 201901184 | 2018 | France | French Guiana | Unknown | ERS6494905 |
| 201902073 | 2018 | France | Metropole | Unknown | ERS6494985 |
| 201902077 | 2018 | France | Metropole | Cape Verde | ERS6494989 |
| 201908168 | 2018 | France | Mayotte | Unknown | ERS6495510 |
| 201908175 | 2018 | France | Mayotte | Unknown | ERS6495513 |
| 201908182 | 2018 | France | Mayotte | Unknown | ERS6495514 |
| 201908188 | 2018 | France | Mayotte | Unknown | ERS6495515 |
| 201908214 | 2018 | France | Mayotte | Unknown | ERS6495520 |
| 201900053 | 2019 | France | Metropole | Unknown | ERS6575939 |
| 201900067 | 2019 | France | Metropole | Unknown | ERS6494836 |
| 201900269 | 2019 | France | Metropole | Cameroon | ERS6494839 |
| 201900275 | 2019 | France | Metropole | Unknown | ERS6494840 |
| 201900295 | 2019 | France | Metropole | Unknown | ERS6494842 |
| 201900310 | 2019 | France | Metropole | India | ERS6494843 |
| 201900319 | 2019 | France | Metropole | None | ERS6494844 |
| 201900388 | 2019 | France | Metropole | Unknown | ERS6494849 |
| 201900396 | 2019 | France | Metropole | Unknown | ERS6494850 |
| 201900402 | 2019 | France | Metropole | Unknown | ERS6494851 |
| 201900413 | 2019 | France | Metropole | Thailand | ERS6494852 |
| 201900431 | 2019 | France | Metropole | Unknown | ERS6494853 |
| 201900455 | 2019 | France | Metropole | None | ERS6494854 |
| 201900475 | 2019 | France | Metropole | None | ERS6494855 |
| 201900492 | 2019 | France | Metropole | Unknown | ERS6494856 |
| 201900530 | 2019 | France | Metropole | Unknown | ERS6494857 |
| 201900565 | 2019 | France | Metropole | Unknown | ERS6494858 |
| 201900607 | 2019 | France | Metropole | None | ERS6494860 |
| 201900656 | 2019 | France | Metropole | Sri Lanka | ERS6494861 |
| 201900682 | 2019 | France | Metropole | Indonesia | ERS6494862 |
| 201900710 | 2019 | France | French Guiana | Unknown | ERS6494863 |
| 201900711 | 2019 | France | French Guiana | Unknown | ERS6494864 |
| 201900716 | 2019 | France | Metropole | None | ERS6494865 |
| 201900724 | 2019 | France | Metropole | Colombia | ERS6494866 |
| 201900733 | 2019 | France | Metropole | Africa | ERS6494867 |
| 201900740 | 2019 | France | Metropole | Unknown | ERS6494868 |
| 201900751 | 2019 | France | Metropole | None | ERS6494869 |
| 201900761 | 2019 | France | Metropole | Senegal | ERS6494870 |
| 201900766 | 2019 | France | Metropole | Unknown | ERS6494871 |
| 201900780 | 2019 | France | Metropole | None | ERS6494872 |
| 201900783 | 2019 | France | Metropole | Unknown | ERS6494873 |
| 201900829 | 2019 | France | French Guiana | Unknown | ERS6494875 |
| 201900850 | 2019 | France | Metropole | Unknown | ERS6494876 |
| 201900863 | 2019 | France | Metropole | Unknown | ERS6494877 |
| 201900891 | 2019 | France | Metropole | None | ERS6494878 |
| 201900902 | 2019 | France | Metropole | Unknown | ERS6494879 |
| 201900918 | 2019 | France | French Guiana | Unknown | ERS6494880 |
| 201900932 | 2019 | France | Metropole | None | ERS6494881 |
| 201900936 | 2019 | France | Metropole | Unknown | ERS6494882 |
| 201900946 | 2019 | France | Metropole | Philippines | ERS6494883 |
| 201900978 | 2019 | France | Metropole | None | ERS6494884 |
| 201900994 | 2019 | France | Metropole | Unknown | ERS6494885 |
| 201900999 | 2019 | France | Metropole | Kenya | ERS6494886 |
| 201901000 | 2019 | France | Metropole | Dominican Republic | ERS6494887 |
| 201901002 | 2019 | France | Metropole | Unknown | ERS6494888 |
| 201901005 | 2019 | France | French Guiana | Unknown | ERS6494889 |
| 201901052 | 2019 | France | Metropole | Unknown | ERS6494890 |
| 201901053 | 2019 | France | Metropole | Brazil | ERS6494891 |
| 201901058 | 2019 | France | Metropole | Unknown | ERS6494892 |
| 201901059 | 2019 | France | Metropole | Unknown | ERS6494893 |
| 201901075 | 2019 | France | Metropole | Unknown | ERS6494894 |
| 201901083 | 2019 | France | Metropole | Unknown | ERS6494895 |
| 201901087 | 2019 | France | Metropole | Unknown | ERS6494896 |
| 201901088 | 2019 | France | Metropole | Unknown | ERS6494897 |
| 201901106 | 2019 | France | Metropole | Unknown | ERS6494898 |
| 201901119 | 2019 | France | Metropole | Unknown | ERS6494899 |
| 201901175 | 2019 | France | French Guiana | Unknown | ERS6494900 |
| 201901176 | 2019 | France | French Guiana | Unknown | ERS6494901 |
| 201901177 | 2019 | France | French Guiana | Unknown | ERS6494902 |
| 201901178 | 2019 | France | French Guiana | Unknown | ERS6494903 |
| 201901179 | 2019 | France | French Guiana | Unknown | ERS6494904 |
| 201901197 | 2019 | France | Metropole | Unknown | ERS6494907 |
| 201901211 | 2019 | France | Metropole | Unknown | ERS6494908 |
| 201901224 | 2019 | France | Metropole | Unknown | ERS6494909 |

| | | | | | |
|---|---|---|---|---|---|
| 201901249 | 2019 | France | Metropole | Cameroon | ERS6494910 |
| 201901275 | 2019 | France | Metropole | None | ERS6494911 |
| 201901285 | 2019 | France | Metropole | None | ERS6494912 |
| 201901289 | 2019 | France | Metropole | None | ERS6494913 |
| 201901334 | 2019 | France | Metropole | Morocco | ERS6494914 |
| 201901359 | 2019 | France | Metropole | None | ERS6494916 |
| 201901374 | 2019 | France | Metropole | None | ERS6494917 |
| 201901404 | 2019 | France | Metropole | India | ERS6494918 |
| 201901405 | 2019 | France | Metropole | None | ERS6494919 |
| 201901476 | 2019 | France | Metropole | Unknown | ERS6494920 |
| 201901489 | 2019 | France | French Guiana | Unknown | ERS6494921 |
| 201901491 | 2019 | France | French Guiana | Unknown | ERS6494922 |
| 201901498 | 2019 | France | Metropole | Unknown | ERS6494923 |
| 201901505 | 2019 | France | Metropole | Unknown | ERS6494924 |
| 201901525 | 2019 | France | Metropole | Thailand | ERS6494925 |
| 201901532 | 2019 | France | Metropole | Ethiopia | ERS6494926 |
| 201901543 | 2019 | France | Metropole | Unknown | ERS6494927 |
| 201901577 | 2019 | France | Metropole | Unknown | ERS6494928 |
| 201901635 | 2019 | France | Metropole | Unknown | ERS6494929 |
| 201901668 | 2019 | France | Metropole | Mexico | ERS6494930 |
| 201901669 | 2019 | France | Metropole | Mexico | ERS6494931 |
| 201901674 | 2019 | France | Metropole | Unknown | ERS6494932 |
| 201901682 | 2019 | France | Metropole | Unknown | ERS6494933 |
| 201901703 | 2019 | France | Metropole | None | ERS6494934 |
| 201901719 | 2019 | France | Metropole | Morocco | ERS6494935 |
| 201901762 | 2019 | France | Metropole | Unknown | ERS6494936 |
| 201901791 | 2019 | France | French Guiana | Unknown | ERS6494937 |
| 201901821 | 2019 | France | Metropole | Morocco | ERS6494938 |
| 201901836 | 2019 | France | Metropole | Peru | ERS6494939 |
| 201901898 | 2019 | France | Metropole | Unknown | ERS6494940 |
| 201901918 | 2019 | France | Metropole | None | ERS6494941 |
| 201901923 | 2019 | France | Metropole | Unknown | ERS6494942 |
| 201901963 | 2019 | France | Metropole | Unknown | ERS6494975 |
| 201901966 | 2019 | France | Metropole | Morocco | ERS6494976 |
| 201902035 | 2019 | France | Metropole | Colombia | ERS6494977 |
| 201902038 | 2019 | France | Metropole | Unknown | ERS6494978 |
| 201902048 | 2019 | France | Metropole | Unknown | ERS6494979 |
| 201902055 | 2019 | France | Metropole | Togo | ERS6494980 |
| 201902066 | 2019 | France | Metropole | Unknown | ERS6494981 |
| 201902067 | 2019 | France | Metropole | China | ERS6494982 |
| 201902068 | 2019 | France | Metropole | India | ERS6494983 |
| 201902070 | 2019 | France | Metropole | Egypt | ERS6494984 |
| 201902074 | 2019 | France | Metropole | Unknown | ERS6494986 |
| 201902075 | 2019 | France | Metropole | Unknown | ERS6494987 |
| 201902076 | 2019 | France | Metropole | Egypt | ERS6494988 |
| 201902078 | 2019 | France | Metropole | Unknown | ERS6494990 |
| 201902079 | 2019 | France | Metropole | Senegal | ERS6494991 |
| 201902080 | 2019 | France | Metropole | Egypt | ERS6494992 |
| 201902081 | 2019 | France | Metropole | Mali | ERS6494993 |
| 201902082 | 2019 | France | Metropole | Unknown | ERS6494994 |
| 201902083 | 2019 | France | Metropole | Philippines | ERS6494995 |
| 201902123 | 2019 | France | Metropole | None | ERS6494996 |
| 201902127 | 2019 | France | Metropole | Unknown | ERS6494997 |
| 201902128 | 2019 | France | Metropole | Unknown | ERS6494998 |
| 201902191 | 2019 | France | Metropole | Unknown | ERS6494999 |
| 201902199 | 2019 | France | Metropole | None | ERS6495000 |
| 201902270 | 2019 | France | Metropole | Unknown | ERS6495001 |
| 201902277 | 2019 | France | Metropole | South Africa | ERS6495002 |
| 201902286 | 2019 | France | Metropole | Cambodia | ERS6495003 |
| 201902303 | 2019 | France | Metropole | Cuba | ERS6495004 |
| 201902337 | 2019 | France | Metropole | Unknown | ERS6495005 |
| 201902375 | 2019 | France | Metropole | Central African Republic | ERS6495006 |
| 201902384 | 2019 | France | Metropole | Unknown | ERS6495007 |
| 201902428 | 2019 | France | Metropole | Morocco | ERS6495008 |
| 201902443 | 2019 | France | Metropole | Morocco | ERS6495009 |
| 201902465 | 2019 | France | Metropole | Unknown | ERS6495010 |
| 201902473 | 2019 | France | Metropole | Sri Lanka | ERS6495011 |
| 201902475 | 2019 | France | Metropole | South Africa | ERS6495012 |
| 201902481 | 2019 | France | Metropole | None | ERS6495013 |
| 201902486 | 2019 | France | Metropole | Dominican Republic | ERS6495014 |
| 201902501 | 2019 | France | Metropole | None | ERS6495015 |
| 201902508 | 2019 | France | Metropole | None | ERS6495016 |
| 201902523 | 2019 | France | Metropole | Morocco | ERS6495017 |
| 201902545 | 2019 | France | Metropole | None | ERS6495018 |
| 201902579 | 2019 | France | Metropole | Oman | ERS6495019 |
| 201902580 | 2019 | France | Metropole | Oman | ERS6575940 |
| 201902607 | 2019 | France | Metropole | Unknown | ERS6495020 |
| 201902621 | 2019 | France | Metropole | Unknown | ERS6495021 |
| 201902629 | 2019 | France | Metropole | El Savador | ERS6575941 |
| 201902630 | 2019 | France | Metropole | None | ERS6495022 |
| 201902637 | 2019 | France | Metropole | None | ERS6495023 |
| 201902691 | 2019 | France | Metropole | Unknown | ERS6495024 |
| 201902700 | 2019 | France | French Guiana | Unknown | ERS6495025 |
| 201902701 | 2019 | France | French Guiana | Unknown | ERS6495026 |
| 201902702 | 2019 | France | French Guiana | Unknown | ERS6495027 |
| 201902703 | 2019 | France | French Guiana | Unknown | ERS6495028 |
| 201902704 | 2019 | France | French Guiana | Unknown | ERS6495029 |
| 201902705 | 2019 | France | French Guiana | Unknown | ERS6495030 |
| 201902706 | 2019 | France | French Guiana | Unknown | ERS6495031 |
| 201902751 | 2019 | France | Metropole | Unknown | ERS6495032 |
| 201902877 | 2019 | France | Metropole | Unknown | ERS6495034 |
| 201902911 | 2019 | France | Metropole | Thailand | ERS6495035 |
| 201902948 | 2019 | France | Metropole | Unknown | ERS6495036 |
| 201902951 | 2019 | France | Metropole | Dominican Republic | ERS6495037 |
| 201903063 | 2019 | France | Metropole | Unknown | ERS6495038 |
| 201903095 | 2019 | France | Metropole | Singapore | ERS6495039 |
| 201903195 | 2019 | France | Metropole | Unknown | ERS6495040 |
| 201903202 | 2019 | France | Metropole | Unknown | ERS6495041 |
| 201903215 | 2019 | France | Metropole | Unknown | ERS6495042 |
| 201903220 | 2019 | France | Metropole | Unknown | ERS6495043 |
| 201903232 | 2019 | France | Metropole | Unknown | ERS6495044 |
| 201903286 | 2019 | France | Metropole | Unknown | ERS6495045 |
| 201903300 | 2019 | France | Metropole | Mayotte | ERS6495046 |

| | | | | | |
|---|---|---|---|---|---|
| 201903304 | 2019 | France | Metropole | Mayotte | ERS6495047 |
| 201903340 | 2019 | France | Metropole | Jordan | ERS6495048 |
| 201903362 | 2019 | France | Metropole | Mexico | ERS6495049 |
| 201903363 | 2019 | France | Metropole | Mexico | ERS6495050 |
| 201903385 | 2019 | France | Metropole | Unknown | ERS6495051 |
| 201903418 | 2019 | France | Metropole | None | ERS6495052 |
| 201903427 | 2019 | France | Metropole | Unknown | ERS6495053 |
| 201903458 | 2019 | France | Metropole | Morocco | ERS6495054 |
| 201903493 | 2019 | France | Metropole | None | ERS6495055 |
| 201903517 | 2019 | France | Metropole | Spain | ERS6495056 |
| 201903518 | 2019 | France | Metropole | Philippines | ERS6495057 |
| 201903554 | 2019 | France | Metropole | None | ERS6495058 |
| 201903555 | 2019 | France | Metropole | None | ERS6495059 |
| 201903556 | 2019 | France | Metropole | Morocco | ERS6495060 |
| 201903690 | 2019 | France | Metropole | Cambodia | ERS6495061 |
| 201903747 | 2019 | France | Metropole | Unknown | ERS6495062 |
| 201903782 | 2019 | France | Metropole | Unknown | ERS6495063 |
| 201903803 | 2019 | France | Metropole | Unknown | ERS6495064 |
| 201903879 | 2019 | France | Metropole | Unknown | ERS6495065 |
| 201903902 | 2019 | France | Metropole | Unknown | ERS6495066 |
| 201903956 | 2019 | France | Metropole | Cambodia | ERS6495067 |
| 201903976 | 2019 | France | Metropole | India | ERS6495068 |
| 201904005 | 2019 | France | Metropole | Pakistan | ERS6495069 |
| 201904057 | 2019 | France | Metropole | None | ERS6495070 |
| 201904113 | 2019 | France | Metropole | Unknown | ERS6492925 |
| 201904126 | 2019 | France | Metropole | None | ERS6495071 |
| 201904370 | 2019 | France | Metropole | Unknown | ERS6495072 |
| 201904378 | 2019 | France | Metropole | Senegal | ERS6495073 |
| 201904438 | 2019 | France | Reunion | Madagascar | ERS6495074 |
| 201904455 | 2019 | France | Metropole | Unknown | ERS6495075 |
| 201904458 | 2019 | France | Metropole | South Africa | ERS6495076 |
| 201904459 | 2019 | France | Metropole | None | ERS6495077 |
| 201904506 | 2019 | France | Metropole | None | ERS6495078 |
| 201904519 | 2019 | France | Metropole | None | ERS6495079 |
| 201904525 | 2019 | France | Metropole | Indonesia | ERS6495080 |
| 201904534 | 2019 | France | Metropole | Unknown | ERS6495081 |
| 201904637 | 2019 | France | Metropole | Tanzania | ERS6495082 |
| 201904686 | 2019 | France | Metropole | Unknown | ERS6495083 |
| 201904769 | 2019 | France | Metropole | Guinea | ERS6495084 |
| 201904828 | 2019 | France | Metropole | Nepal | ERS6492926 |
| 201904837 | 2019 | France | Metropole | Unknown | ERS6492927 |
| 201904869 | 2019 | France | Metropole | None | ERS6495085 |
| 201904893 | 2019 | France | Metropole | India | ERS6495086 |
| 201904910 | 2019 | France | Metropole | Indonesia | ERS6495087 |
| 201904911 | 2019 | France | Metropole | Unknown | ERS6495088 |
| 201904961 | 2019 | France | Metropole | Unknown | ERS6495089 |
| 201904971 | 2019 | France | Metropole | India | ERS6495090 |
| 201904981 | 2019 | France | Metropole | None | ERS6495091 |
| 201905024 | 2019 | France | Metropole | None | ERS6495093 |
| 201905339 | 2019 | France | Metropole | None | ERS6495097 |
| 201905343 | 2019 | France | Metropole | None | ERS6495098 |
| 201905344 | 2019 | France | Metropole | Unknown | ERS6495099 |
| 201905345 | 2019 | France | Metropole | Unknown | ERS6495100 |
| 201905346 | 2019 | France | Metropole | Unknown | ERS6495101 |
| 201905347 | 2019 | France | Metropole | Unknown | ERS6495102 |
| 201905353 | 2019 | France | Metropole | Unknown | ERS6495103 |
| 201905364 | 2019 | France | Metropole | Egypt | ERS6495104 |
| 201905378 | 2019 | France | Metropole | None | ERS6495105 |
| 201905420 | 2019 | France | Metropole | Unknown | ERS6495106 |
| 201905421 | 2019 | France | Metropole | Indonesia | ERS6495107 |
| 201905484 | 2019 | France | Metropole | Unknown | ERS6495108 |
| 201905488 | 2019 | France | Metropole | Unknown | ERS6495109 |
| 201905544 | 2019 | France | Metropole | Unknown | ERS6495110 |
| 201905545 | 2019 | France | Metropole | Unknown | ERS6495111 |
| 201905559 | 2019 | France | Metropole | Unknown | ERS6495112 |
| 201905594 | 2019 | France | Metropole | Unknown | ERS6495113 |
| 201905595 | 2019 | France | Metropole | Unknown | ERS6495114 |
| 201905610 | 2019 | France | Metropole | Unknown | ERS6495115 |
| 201905690 | 2019 | France | Metropole | None | ERS6495116 |
| 201905782 | 2019 | France | Metropole | Unknown | ERS6495117 |
| 201905824 | 2019 | France | Metropole | Unknown | ERS6495118 |
| 201905825 | 2019 | France | Metropole | Mali | ERS6495119 |
| 201905832 | 2019 | France | Metropole | Tunisia | ERS6495120 |
| 201905843 | 2019 | France | French Guiana | Unknown | ERS6495121 |
| 201905853 | 2019 | France | Metropole | Unknown | ERS6495122 |
| 201905857 | 2019 | France | Metropole | Unknown | ERS6495123 |
| 201905858 | 2019 | France | Metropole | Unknown | ERS6495124 |
| 201905875 | 2019 | France | Metropole | Unknown | ERS6495125 |
| 201905885 | 2019 | France | Metropole | Senegal | ERS6495126 |
| 201905900 | 2019 | France | Metropole | India | ERS6495127 |
| 201905901 | 2019 | France | Metropole | Unknown | ERS6495128 |
| 201905955 | 2019 | France | Metropole | Unknown | ERS6495129 |
| 201905959 | 2019 | France | French Guiana | Unknown | ERS6495130 |
| 201905964 | 2019 | France | Metropole | India | ERS6495131 |
| 201905967 | 2019 | France | Metropole | None | ERS6495132 |
| 201906055 | 2019 | France | Metropole | Egypt | ERS6495133 |
| 201906060 | 2019 | France | Metropole | Unknown | ERS6495134 |
| 201906092 | 2019 | France | Metropole | Unknown | ERS6495135 |
| 201906103 | 2019 | France | Metropole | Unknown | ERS6495136 |
| 201906104 | 2019 | France | Metropole | Unknown | ERS6495137 |
| 201906105 | 2019 | France | Metropole | Senegal | ERS6495138 |
| 201906114 | 2019 | France | Metropole | Unknown | ERS6495139 |
| 201906159 | 2019 | France | Metropole | None | ERS6495140 |
| 201906166 | 2019 | France | Metropole | Unknown | ERS6495141 |
| 201906173 | 2019 | France | Metropole | Colombia | ERS6495142 |
| 201906188 | 2019 | France | Metropole | Unknown | ERS6495143 |
| 201906192 | 2019 | France | Metropole | Colombia | ERS6495144 |
| 201906202 | 2019 | France | Metropole | India | ERS6495145 |
| 201906203 | 2019 | France | Metropole | None | ERS6495146 |
| 201906533 | 2019 | France | Metropole | None | ERS6495147 |
| 201906557 | 2019 | France | Metropole | None | ERS6495148 |
| 201906645 | 2019 | France | Metropole | Unknown | ERS6495149 |
| 201906684 | 2019 | France | Metropole | Unknown | ERS6495150 |

| | | | | | |
|---|---|---|---|---|---|
| 201906714 | 2019 | France | Metropole | Unknown | ERS6495151 |
| 201906746 | 2019 | France | Metropole | Unknown | ERS6495152 |
| 201906751 | 2019 | France | Metropole | Unknown | ERS6495153 |
| 201906763 | 2019 | France | Metropole | Morocco | ERS6495154 |
| 201906835 | 2019 | France | Metropole | Morocco | ERS6495155 |
| 201906853 | 2019 | France | Metropole | Unknown | ERS6495156 |
| 201906932 | 2019 | France | Metropole | Unknown | ERS6495157 |
| 201906956 | 2019 | France | Metropole | None | ERS6495158 |
| 201906984 | 2019 | France | Metropole | Senegal | ERS6495159 |
| 201906985 | 2019 | France | Metropole | Unknown | ERS6495160 |
| 201907010 | 2019 | France | Metropole | Unknown | ERS6495161 |
| 201907035 | 2019 | France | Metropole | Unknown | ERS6495162 |
| 201907038 | 2019 | France | Metropole | Unknown | ERS6495163 |
| 201907048 | 2019 | France | Metropole | Philippines | ERS6495164 |
| 201907083 | 2019 | France | Metropole | Unknown | ERS6495165 |
| 201907091 | 2019 | France | Metropole | Morocco | ERS6495166 |
| 201907092 | 2019 | France | Metropole | Morocco | ERS6495167 |
| 201907108 | 2019 | France | Metropole | Turkey | ERS6495168 |
| 201907109 | 2019 | France | Metropole | India | ERS6495169 |
| 201907110 | 2019 | France | Metropole | Egypt | ERS6495170 |
| 201907123 | 2019 | France | Metropole | Mali | ERS6495171 |
| 201907131 | 2019 | France | Metropole | Algeria | ERS6495172 |
| 201907158 | 2019 | France | Metropole | Unknown | ERS6495173 |
| 201907172 | 2019 | France | Metropole | Unknown | ERS6495174 |
| 201907173 | 2019 | France | Metropole | Unknown | ERS6575942 |
| 201907204 | 2019 | France | Metropole | Unknown | ERS6495448 |
| 201907206 | 2019 | France | Metropole | Unknown | ERS6495449 |
| 201907208 | 2019 | France | Metropole | Morocco | ERS6495450 |
| 201907217 | 2019 | France | Metropole | Peru | ERS6495451 |
| 201907329 | 2019 | France | Metropole | None | ERS6495452 |
| 201907347 | 2019 | France | Metropole | Nepal | ERS6495453 |
| 201907359 | 2019 | France | Metropole | None | ERS6495454 |
| 201907369 | 2019 | France | Metropole | Unknown | ERS6495455 |
| 201907375 | 2019 | France | Metropole | Senegal | ERS6495456 |
| 201907376 | 2019 | France | Metropole | Unknown | ERS6495457 |
| 201907377 | 2019 | France | Metropole | Unknown | ERS6495458 |
| 201907378 | 2019 | France | Metropole | Spain | ERS6495459 |
| 201907379 | 2019 | France | Metropole | Unknown | ERS6495460 |
| 201907401 | 2019 | France | Metropole | Unknown | ERS6495461 |
| 201907414 | 2019 | France | Metropole | Montenegro | ERS6495462 |
| 201907494 | 2019 | France | Metropole | None | ERS6495463 |
| 201907510 | 2019 | France | Metropole | Morocco | ERS6495464 |
| 201907529 | 2019 | France | Metropole | Unknown | ERS6495465 |
| 201907538 | 2019 | France | Metropole | Tunisia | ERS6495466 |
| 201907540 | 2019 | France | Metropole | Unknown | ERS6495467 |
| 201907551 | 2019 | France | Metropole | Morocco | ERS6495468 |
| 201907576 | 2019 | France | Metropole | Unknown | ERS6495469 |
| 201907593 | 2019 | France | Metropole | Ecuador | ERS6495470 |
| 201907612 | 2019 | France | Metropole | Unknown | ERS6495471 |
| 201907632 | 2019 | France | Metropole | Egypt | ERS6495472 |
| 201907664 | 2019 | France | Metropole | None | ERS6495473 |
| 201907665 | 2019 | France | Metropole | Israel | ERS6492928 |
| 201907681 | 2019 | France | Metropole | Unknown | ERS6495474 |
| 201907700 | 2019 | France | Metropole | Unknown | ERS6495475 |
| 201907701 | 2019 | France | Metropole | Unknown | ERS6495476 |
| 201907720 | 2019 | France | Metropole | None | ERS6495477 |
| 201907729 | 2019 | France | Metropole | None | ERS6495478 |
| 201907764 | 2019 | France | Metropole | None | ERS6495479 |
| 201907765 | 2019 | France | Metropole | None | ERS6575943 |
| 201907784 | 2019 | France | Metropole | Morocco | ERS6495480 |
| 201907801 | 2019 | France | Metropole | Greece | ERS6495481 |
| 201907834 | 2019 | France | Metropole | Unknown | ERS6495482 |
| 201907848 | 2019 | France | Metropole | None | ERS6495483 |
| 201907849 | 2019 | France | Metropole | Unknown | ERS6495484 |
| 201907850 | 2019 | France | Metropole | Unknown | ERS6495485 |
| 201907851 | 2019 | France | Metropole | Unknown | ERS6575944 |
| 201907857 | 2019 | France | Metropole | Unknown | ERS6495486 |
| 201907860 | 2019 | France | Metropole | Unknown | ERS6495487 |
| 201907866 | 2019 | France | Metropole | Unknown | ERS6495488 |
| 201907869 | 2019 | France | Metropole | Morocco | ERS6495489 |
| 201907882 | 2019 | France | Metropole | Unknown | ERS6495490 |
| 201907887 | 2019 | France | Metropole | Algeria | ERS6495491 |
| 201907888 | 2019 | France | Metropole | Unknown | ERS6495492 |
| 201907951 | 2019 | France | Metropole | None | ERS6495493 |
| 201907964 | 2019 | France | Metropole | Unknown | ERS6495494 |
| 201907985 | 2019 | France | Metropole | Unknown | ERS6495495 |
| 201907987 | 2019 | France | Metropole | None | ERS6495496 |
| 201908000 | 2019 | France | Metropole | Italy | ERS6495497 |
| 201908011 | 2019 | France | Metropole | Georgia | ERS6495498 |
| 201908022 | 2019 | France | Metropole | Morocco | ERS6495499 |
| 201908023 | 2019 | France | Metropole | Unknown | ERS6495500 |
| 201908031 | 2019 | France | Metropole | Unknown | ERS6495501 |
| 201908032 | 2019 | France | Metropole | Unknown | ERS6495502 |
| 201908033 | 2019 | France | Metropole | Unknown | ERS6495503 |
| 201908051 | 2019 | France | Metropole | Morocco | ERS6495504 |
| 201908102 | 2019 | France | Metropole | Unknown | ERS6495505 |
| 201908104 | 2019 | France | Metropole | None | ERS6495506 |
| 201908113 | 2019 | France | Metropole | South Africa | ERS6495507 |
| 201908114 | 2019 | France | Metropole | Morocco | ERS6495508 |
| 201908157 | 2019 | France | Metropole | Unknown | ERS6495509 |
| 201908171 | 2019 | France | Mayotte | Unknown | ERS6495511 |
| 201908174 | 2019 | France | Mayotte | Unknown | ERS6495512 |
| 201908195 | 2019 | France | Mayotte | Unknown | ERS6495516 |
| 201908196 | 2019 | France | Mayotte | Unknown | ERS6495517 |
| 201908208 | 2019 | France | Mayotte | Unknown | ERS6495518 |
| 201908210 | 2019 | France | Mayotte | Unknown | ERS6495519 |
| 201908231 | 2019 | France | Metropole | None | ERS6495521 |
| 201908234 | 2019 | France | Metropole | None | ERS12446240 |
| 201908244 | 2019 | France | Metropole | Morocco | ERS6495522 |
| 201908245 | 2019 | France | Metropole | Morocco | ERS6576213 |
| 201908252 | 2019 | France | Metropole | Unknown | ERS6495523 |
| 201908272 | 2019 | France | Metropole | Unknown | ERS6495524 |
| 201908280 | 2019 | France | Metropole | Morocco | ERS6495525 |

| | | | | | |
|---|---|---|---|---|---|
| 201908305 | 2019 | France | Metropole | Unknown | ERS6495526 |
| 201908320 | 2019 | France | Metropole | Unknown | ERS6495527 |
| 201908330 | 2019 | France | Metropole | None | ERS6495528 |
| 201908347 | 2019 | France | Metropole | Unknown | ERS6495529 |
| 201908357 | 2019 | France | Metropole | Unknown | ERS6495530 |
| 201908364 | 2019 | France | Metropole | Morocco | ERS6495531 |
| 201908372 | 2019 | France | Metropole | Turkey | ERS6495532 |
| 201908373 | 2019 | France | Metropole | Unknown | ERS6495533 |
| 201908400 | 2019 | France | Metropole | Tanzania | ERS6495534 |
| 201908412 | 2019 | France | Metropole | Unknown | ERS6495535 |
| 201908413 | 2019 | France | Metropole | Colombia | ERS6495536 |
| 201908415 | 2019 | France | Metropole | India | ERS6495537 |
| 201908418 | 2019 | France | Metropole | Côte d'Ivoire | ERS6495538 |
| 201908442 | 2019 | France | Metropole | Unknown | ERS6495539 |
| 201908490 | 2019 | France | Metropole | Unknown | ERS6495540 |
| 201908550 | 2019 | France | Metropole | Madagascar | ERS6495541 |
| 201908554 | 2019 | France | Metropole | Unknown | ERS6495542 |
| 201908573 | 2019 | France | Metropole | Morocco | ERS6495543 |
| 201908581 | 2019 | France | Metropole | Malaysia_Singapore | ERS6495544 |
| 201908585 | 2019 | France | Metropole | Unknown | ERS6495545 |
| 201908588 | 2019 | France | Metropole | Unknown | ERS6495547 |
| 201908589 | 2019 | France | Metropole | Unknown | ERS6495548 |
| 201908591 | 2019 | France | Metropole | Unknown | ERS6495549 |
| 201908645 | 2019 | France | Metropole | Greece | ERS6495550 |
| 201908671 | 2019 | France | Metropole | Morocco | ERS6495551 |
| 201908672 | 2019 | France | Metropole | Tanzania | ERS6495552 |
| 201908697 | 2019 | France | Metropole | Unknown | ERS6495553 |
| 201908698 | 2019 | France | Metropole | Unknown | ERS6495554 |
| 201908718 | 2019 | France | Metropole | Algeria | ERS6495555 |
| 201908737 | 2019 | France | Metropole | Unknown | ERS6495557 |
| 201908738 | 2019 | France | French Guiana | Unknown | ERS6495558 |
| 201908750 | 2019 | France | Metropole | Egypt | ERS6495559 |
| 201908751 | 2019 | France | Metropole | Serbia | ERS6495560 |
| 201908778 | 2019 | France | Metropole | None | ERS6495562 |
| 201908783 | 2019 | France | Metropole | Unknown | ERS6495563 |
| 201908799 | 2019 | France | French Guiana | Unknown | ERS6495564 |
| 201908821 | 2019 | France | Metropole | None | ERS6495565 |
| 201908832 | 2019 | France | Metropole | Morocco | ERS6495566 |
| 201908860 | 2019 | France | Metropole | Egypt | ERS6495567 |
| 201908899 | 2019 | France | Metropole | None | ERS6495568 |
| 201908922 | 2019 | France | Metropole | Unknown | ERS6495569 |
| 201908945 | 2019 | France | Metropole | Unknown | ERS6495571 |
| 201908973 | 2019 | France | Metropole | Unknown | ERS6495572 |
| 201908995 | 2019 | France | Metropole | Turkey | ERS6495574 |
| 201909029 | 2019 | France | Metropole | Unknown | ERS6495575 |
| 201909079 | 2019 | France | Metropole | Unknown | ERS6495576 |
| 201909081 | 2019 | France | Metropole | Unknown | ERS6495577 |
| 201909101 | 2019 | France | Metropole | None | ERS6495578 |
| 201909147 | 2019 | France | Metropole | Unknown | ERS6495579 |
| 201909153 | 2019 | France | Metropole | None | ERS6495580 |
| 201909183 | 2019 | France | Metropole | Morocco | ERS6495581 |
| 201909193 | 2019 | France | Metropole | Morocco | ERS6495582 |
| 201909194 | 2019 | France | Metropole | Algeria | ERS6495583 |
| 201909234 | 2019 | France | Metropole | Spain | ERS6495584 |
| 201909242 | 2019 | France | Metropole | Unknown | ERS6495585 |
| 201909276 | 2019 | France | Mayotte | Unknown | ERS6495588 |
| 201909284 | 2019 | France | Metropole | Unknown | ERS6495589 |
| 201909285 | 2019 | France | Metropole | Morocco | ERS6495590 |
| 201909289 | 2019 | France | Metropole | Spain | ERS6495591 |
| 201909290 | 2019 | France | Metropole | Unknown | ERS6495592 |
| 201909292 | 2019 | France | Metropole | Unknown | ERS6495593 |
| 201909346 | 2019 | France | Metropole | Egypt | ERS6495594 |
| 201909362 | 2019 | France | Metropole | Unknown | ERS6495595 |
| 201909377 | 2019 | France | Metropole | Unknown | ERS6495596 |
| 201909414 | 2019 | France | Metropole | None | ERS6495597 |
| 201909467 | 2019 | France | Metropole | Morocco | ERS6495598 |
| 201909480 | 2019 | France | Metropole | None | ERS6495599 |
| 201909481 | 2019 | France | Metropole | Unknown | ERS6495600 |
| 201909505 | 2019 | France | Metropole | Unknown | ERS6495601 |
| 201909520 | 2019 | France | Metropole | Unknown | ERS6495602 |
| 201909527 | 2019 | France | Metropole | Unknown | ERS6495603 |
| 201909538 | 2019 | France | Metropole | Mexico | ERS6495604 |
| 201909544 | 2019 | France | Metropole | Unknown | ERS6495605 |
| 201909571 | 2019 | France | Metropole | None | ERS6495606 |
| 201909591 | 2019 | France | Metropole | Unknown | ERS6495607 |
| 201909651 | 2019 | France | Metropole | Algeria | ERS6495608 |
| 201909673 | 2019 | France | Metropole | None | ERS6495609 |
| 201909725 | 2019 | France | Metropole | Unknown | ERS6495610 |
| 201909776 | 2019 | France | Metropole | Moldovia | ERS6495611 |
| 201909798 | 2019 | France | Metropole | Unknown | ERS6495612 |
| 201909824 | 2019 | France | Metropole | Unknown | ERS6495613 |
| 201909825 | 2019 | France | Metropole | Unknown | ERS6495614 |
| 201909826 | 2019 | France | Metropole | Unknown | ERS6495615 |
| 201909827 | 2019 | France | Metropole | Unknown | ERS6495616 |
| 201909832 | 2019 | France | Metropole | Unknown | ERS6495617 |
| 201909849 | 2019 | France | Metropole | None | ERS6495618 |
| 201909876 | 2019 | France | Metropole | None | ERS6492929 |
| 201909910 | 2019 | France | Metropole | Unknown | ERS6495619 |
| 201909911 | 2019 | France | Metropole | Unknown | ERS6492930 |
| 201909917 | 2019 | France | Metropole | Sri Lanka | ERS6495620 |
| 201909920 | 2019 | France | Metropole | Morocco | ERS6495621 |
| 201909936 | 2019 | France | Metropole | None | ERS6492931 |
| 201909937 | 2019 | France | Metropole | None | ERS6492932 |
| 201909978 | 2019 | France | Metropole | None | ERS6495622 |
| 201909982 | 2019 | France | Metropole | Morocco | ERS6495623 |
| 201909983 | 2019 | France | Metropole | Unknown | ERS6495624 |
| 201910012 | 2019 | France | French Guiana | Unknown | ERS6495625 |
| 201910025 | 2019 | France | Metropole | None | ERS6495626 |
| 201910033 | 2019 | France | Metropole | Unknown | ERS6495627 |
| 201910034 | 2019 | France | Metropole | Egypt | ERS6495628 |
| 201910039 | 2019 | France | Metropole | None | ERS6495629 |
| 201910042 | 2019 | France | Metropole | Unknown | ERS6495631 |
| 201910055 | 2019 | France | Metropole | Algeria | ERS6495632 |

| | | | | | |
|---|---|---|---|---|---|
| 201910061 | 2019 | France | Metropole | Unknown | ERS6495633 |
| 201910062 | 2019 | France | Metropole | Unknown | ERS6495634 |
| 201910090 | 2019 | France | Metropole | Cape Verde | ERS6495635 |
| 201910104 | 2019 | France | Metropole | Unknown | ERS6495636 |
| 201910112 | 2019 | France | Metropole | Unknown | ERS6495637 |
| 201910113 | 2019 | France | Metropole | Colombia | ERS6495638 |
| 201910122 | 2019 | France | Metropole | Spain | ERS12446241 |
| 201910147 | 2019 | France | Metropole | Unknown | ERS6495639 |
| 201910150 | 2019 | France | Reunion | Madagascar | ERS6495640 |
| 201910179 | 2019 | France | Metropole | Morocco | ERS6495641 |
| 201910208 | 2019 | France | Metropole | Unknown | ERS6495642 |
| 201910218 | 2019 | France | Metropole | India | ERS6495643 |
| 201910219 | 2019 | France | Metropole | Unknown | ERS6495644 |
| 201910220 | 2019 | France | Metropole | None | ERS6495645 |
| 201910269 | 2019 | France | Metropole | Senegal | ERS6495646 |
| 201910295 | 2019 | France | Metropole | None | ERS6492933 |
| 201910300 | 2019 | France | Metropole | Unknown | ERS6492934 |
| 201910304 | 2019 | France | Metropole | Unknown | ERS6574849 |
| 201910325 | 2019 | France | Metropole | Unknown | ERS6574850 |
| 201910328 | 2019 | France | Metropole | None | ERS6574851 |
| 201910348 | 2019 | France | Metropole | Unknown | ERS6574852 |
| 201910350 | 2019 | France | Metropole | Unknown | ERS6574853 |
| 201910391 | 2019 | France | Metropole | Algeria | ERS6574854 |
| 201910405 | 2019 | France | Metropole | Central African Republic | ERS6574855 |
| 201910409 | 2019 | France | Reunion | Unknown | ERS6574856 |
| 201910436 | 2019 | France | Metropole | Unknown | ERS6492935 |
| 201910438 | 2019 | France | Metropole | Unknown | ERS6574857 |
| 201910445 | 2019 | France | Metropole | None | ERS6574858 |
| 201910446 | 2019 | France | Metropole | None | ERS6574859 |
| 201910447 | 2019 | France | Metropole | None | ERS6574860 |
| 201910460 | 2019 | France | Metropole | Unknown | ERS6574861 |
| 201910465 | 2019 | France | Metropole | Senegal | ERS6574862 |
| 201910474 | 2019 | France | Metropole | Senegal | ERS6574863 |
| 201910492 | 2019 | France | Metropole | Unknown | ERS6574864 |
| 201910509 | 2019 | France | Metropole | None | ERS6574865 |
| 201910532 | 2019 | France | Metropole | Unknown | ERS6574866 |
| 201910541 | 2019 | France | Metropole | Unknown | ERS6574867 |
| 201910546 | 2019 | France | Metropole | Unknown | ERS6492936 |
| 201910551 | 2019 | France | Metropole | None | ERS6574868 |
| 201910554 | 2019 | France | Metropole | Senegal | ERS6574869 |
| 201910556 | 2019 | France | Metropole | Madagascar | ERS6574870 |
| 201910566 | 2019 | France | Metropole | Unknown | ERS6574871 |
| 201910648 | 2019 | France | Metropole | Algeria | ERS6574872 |
| 201910650 | 2019 | France | Metropole | Morocco | ERS6574873 |
| 201910674 | 2019 | France | Reunion | Madagascar | ERS6575035 |
| 201910698 | 2019 | France | Metropole | Unknown | ERS6575036 |
| 201910715 | 2019 | France | Metropole | Togo | ERS6574874 |
| 201910726 | 2019 | France | Metropole | Egypt | ERS6575037 |
| 201910729 | 2019 | France | Metropole | Greece | ERS6575038 |
| 201910769 | 2019 | France | Metropole | Unknown | ERS6575039 |
| 201910789 | 2019 | France | Metropole | Unknown | ERS6575041 |
| 201910790 | 2019 | France | Metropole | Unknown | ERS6492937 |
| 201910791 | 2019 | France | Metropole | Unknown | ERS6492938 |
| 201910792 | 2019 | France | Metropole | Unknown | ERS6575042 |
| 201910793 | 2019 | France | Metropole | Unknown | ERS6575043 |
| 201910794 | 2019 | France | Metropole | Unknown | ERS6575044 |
| 201910803 | 2019 | France | Metropole | None | ERS6575045 |
| 201910807 | 2019 | France | Metropole | Germany | ERS6575046 |
| 201910819 | 2019 | France | Metropole | Unknown | ERS6575047 |
| 201910824 | 2019 | France | Metropole | Unknown | ERS6575048 |
| 201910825 | 2019 | France | Metropole | Unknown | ERS6575049 |
| 201910826 | 2019 | France | Metropole | Unknown | ERS6575050 |
| 201910852 | 2019 | France | Metropole | Unknown | ERS6575051 |
| 201910868 | 2019 | France | Metropole | Iraq | ERS6575052 |
| 201910873 | 2019 | France | Metropole | Thailand | ERS6575053 |
| 201910886 | 2019 | France | Metropole | None | ERS6575054 |
| 201910890 | 2019 | France | Metropole | Unknown | ERS6575055 |
| 201910893 | 2019 | France | Metropole | None | ERS6575056 |
| 201910905 | 2019 | France | Metropole | None | ERS6575057 |
| 201910916 | 2019 | France | Metropole | Unknown | ERS6575058 |
| 201910919 | 2019 | France | Metropole | Unknown | ERS6575059 |
| 201910920 | 2019 | France | Metropole | Mexico | ERS6575060 |
| 201910921 | 2019 | France | Metropole | Unknown | ERS6575061 |
| 201910923 | 2019 | France | Metropole | Unknown | ERS6575062 |
| 201910938 | 2019 | France | Metropole | Unknown | ERS6575063 |
| 201910939 | 2019 | France | Metropole | Unknown | ERS6575064 |
| 201910940 | 2019 | France | Metropole | Unknown | ERS6575065 |
| 201910947 | 2019 | France | Metropole | Unknown | ERS6575066 |
| 201910969 | 2019 | France | Metropole | Indonesia | ERS6575067 |
| 201910971 | 2019 | France | Metropole | Unknown | ERS6575068 |
| 201910977 | 2019 | France | Metropole | Niger | ERS6575069 |
| 201910995 | 2019 | France | Metropole | Reunion | ERS6575070 |
| 201911002 | 2019 | France | Metropole | Unknown | ERS6575071 |
| 201911006 | 2019 | France | Metropole | Unknown | ERS6575072 |
| 201911037 | 2019 | France | Metropole | Unknown | ERS6575073 |
| 201911039 | 2019 | France | Metropole | Unknown | ERS6575074 |
| 201911040 | 2019 | France | Metropole | Unknown | ERS6575075 |
| 201911042 | 2019 | France | Metropole | Unknown | ERS6575076 |
| 201911045 | 2019 | France | Metropole | Unknown | ERS6575077 |
| 201911077 | 2019 | France | Metropole | South Africa | ERS6575078 |
| 201911079 | 2019 | France | French Guiana | Unknown | ERS6575079 |
| 201911084 | 2019 | France | Metropole | Unknown | ERS6575080 |
| 201911085 | 2019 | France | Metropole | Unknown | ERS6575081 |
| 201911086 | 2019 | France | Metropole | Unknown | ERS6575082 |
| 201911087 | 2019 | France | Metropole | Unknown | ERS6575083 |
| 201911094 | 2019 | France | Metropole | Unknown | ERS6575084 |
| 201911165 | 2019 | France | Metropole | Unknown | ERS6575085 |
| 201911173 | 2019 | France | Reunion | None | ERS6575086 |
| 201911174 | 2019 | France | Reunion | None | ERS6575087 |
| 201911179 | 2019 | France | Metropole | Unknown | ERS6575088 |
| 201911215 | 2019 | France | Metropole | Egypt | ERS6575089 |
| 201911216 | 2019 | France | Metropole | None | ERS6575090 |
| 201911239 | 2019 | France | Mayotte | Unknown | ERS6575091 |

| | | | | | |
|---|---|---|---|---|---|
| 201911245 | 2019 | France | Mayotte | Unknown | ERS6575092 |
| 201911259 | 2019 | France | Metropole | Tanzania | ERS6575093 |
| 201911260 | 2019 | France | Metropole | Unknown | ERS6575094 |
| 201911261 | 2019 | France | Metropole | Unknown | ERS6575095 |
| 201911263 | 2019 | France | Metropole | Unknown | ERS6575096 |
| 201911294 | 2019 | France | Metropole | Unknown | ERS6575097 |
| 201911296 | 2019 | France | Metropole | Unknown | ERS6575098 |
| 201911310 | 2019 | France | Metropole | Unknown | ERS6575099 |
| 201911317 | 2019 | France | Metropole | Unknown | ERS6575100 |
| 201911323 | 2019 | France | Metropole | Unknown | ERS6575101 |
| 201911328 | 2019 | France | Metropole | Unknown | ERS6575102 |
| 201911333 | 2019 | France | Metropole | Unknown | ERS6575103 |
| 201911334 | 2019 | France | Metropole | Unknown | ERS6575104 |
| 201911336 | 2019 | France | Metropole | Unknown | ERS6575105 |
| 201911340 | 2019 | France | Metropole | Unknown | ERS6575106 |
| 201911341 | 2019 | France | Metropole | Unknown | ERS6575107 |
| 201911342 | 2019 | France | Metropole | Unknown | ERS6575108 |
| 201911345 | 2019 | France | Metropole | Unknown | ERS6575109 |
| 201911346 | 2019 | France | Metropole | Unknown | ERS6575110 |
| 201911347 | 2019 | France | Metropole | Unknown | ERS6575111 |
| 201911348 | 2019 | France | Metropole | Unknown | ERS6575112 |
| 201911350 | 2019 | France | Metropole | Unknown | ERS6575113 |
| 201911351 | 2019 | France | Metropole | Unknown | ERS6575114 |
| 201911352 | 2019 | France | Metropole | Unknown | ERS6575115 |
| 201911378 | 2019 | France | Metropole | Unknown | ERS6575116 |
| 201911384 | 2019 | France | Metropole | None | ERS6575117 |
| 201911396 | 2019 | France | Metropole | Unknown | ERS6575118 |
| 201911404 | 2019 | France | Metropole | Unknown | ERS6575119 |
| 201911408 | 2019 | France | Metropole | Tanzania | ERS6575120 |
| 201911418 | 2019 | France | Metropole | Guinea | ERS6492939 |
| 201911426 | 2019 | France | Metropole | Unknown | ERS6492940 |
| 201911428 | 2019 | France | Metropole | Unknown | ERS6492941 |
| 201911433 | 2019 | France | Metropole | None | ERS6492942 |
| 201911445 | 2019 | France | Metropole | Unknown | ERS6492943 |
| 201911446 | 2019 | France | Metropole | None | ERS6492944 |
| 201911449 | 2019 | France | Metropole | Unknown | ERS6492945 |
| 201911450 | 2019 | France | Metropole | Unknown | ERS6492946 |
| 201911457 | 2019 | France | Metropole | Unknown | ERS6492947 |
| 201911458 | 2019 | France | Metropole | Unknown | ERS6492948 |
| 201911459 | 2019 | France | Metropole | Unknown | ERS6492949 |
| 201911461 | 2019 | France | Metropole | Unknown | ERS6492950 |
| 201911465 | 2019 | France | Reunion | Madagascar | ERS6492951 |
| 201911468 | 2019 | France | Metropole | Unknown | ERS6492952 |
| 201911495 | 2019 | France | Metropole | Unknown | ERS6492953 |
| 201911497 | 2019 | France | Metropole | Unknown | ERS6492954 |
| 201911505 | 2019 | France | Metropole | Unknown | ERS6492955 |
| 201911509 | 2019 | France | Metropole | Unknown | ERS6492956 |
| 201911510 | 2019 | France | Metropole | Unknown | ERS6492957 |
| 201911511 | 2019 | France | Metropole | Unknown | ERS6495811 |
| 201911512 | 2019 | France | Metropole | Unknown | ERS6492958 |
| 201911513 | 2019 | France | Metropole | None | ERS6492959 |
| 201911514 | 2019 | France | Metropole | Unknown | ERS6492960 |
| 201911518 | 2019 | France | Metropole | Unknown | ERS6492961 |
| 201911540 | 2019 | France | Metropole | Unknown | ERS6492962 |
| 201911551 | 2019 | France | Metropole | Unknown | ERS6492963 |
| 201911556 | 2019 | France | Metropole | None | ERS6495812 |
| 201911568 | 2019 | France | Metropole | Unknown | ERS6495813 |
| 201911576 | 2019 | France | Metropole | None | ERS6495814 |
| 201911610 | 2019 | France | Metropole | Unknown | ERS6495815 |
| 201911625 | 2019 | France | Metropole | None | ERS6495816 |
| 201911626 | 2019 | France | Metropole | None | ERS6495817 |
| 201911627 | 2019 | France | Metropole | None | ERS6495818 |
| 201911638 | 2019 | France | Metropole | None | ERS6495647 |
| 201911641 | 2019 | France | Metropole | None | ERS6495819 |
| 201911645 | 2019 | France | Metropole | Unknown | ERS6495820 |
| 201911650 | 2019 | France | Metropole | None | ERS6495821 |
| 202000038 | 2019 | France | Metropole | None | ERS6495822 |
| 202000059 | 2019 | France | Reunion | None | ERS6495823 |
| 202000151 | 2019 | France | French Guiana | Unknown | ERS6495825 |
| 202000152 | 2019 | France | French Guiana | Unknown | ERS6495826 |
| 202000158 | 2019 | France | Metropole | Unknown | ERS6495827 |
| 202000159 | 2019 | France | Metropole | Madagascar | ERS6495828 |
| 202000163 | 2019 | France | Reunion | Unknown | ERS6495829 |
| 202000221 | 2019 | France | Metropole | Unknown | ERS6495833 |
| 202000223 | 2019 | France | Metropole | Unknown | ERS6495834 |
| 202000258 | 2019 | France | Reunion | Unknown | ERS6495835 |
| 202000123 | 2020 | France | Metropole | Egypt | ERS6495824 |
| 202000169 | 2020 | France | Metropole | Unknown | ERS6495830 |
| 202000170 | 2020 | France | Metropole | Unknown | ERS6495831 |
| 202000197 | 2020 | France | Metropole | Unknown | ERS6495832 |
| 202000267 | 2020 | France | Metropole | None | ERS6495836 |
| 202000274 | 2020 | France | Metropole | None | ERS6495837 |
| 202000288 | 2020 | France | Metropole | Unknown | ERS6495838 |
| 202000293 | 2020 | France | Metropole | Madagascar | ERS6495839 |
| 202000316 | 2020 | France | Metropole | Unknown | ERS6495840 |
| 202000322 | 2020 | France | Metropole | Unknown | ERS6495841 |
| 202000351 | 2020 | France | Metropole | Unknown | ERS6495842 |
| 202000371 | 2020 | France | Metropole | Unknown | ERS6495843 |
| 202000378 | 2020 | France | Metropole | Cuba | ERS6495844 |
| 202000382 | 2020 | France | Metropole | Tunisia | ERS6495845 |
| 202000386 | 2020 | France | Metropole | Unknown | ERS6495846 |
| 202000436 | 2020 | France | Metropole | Unknown | ERS6575377 |
| 202000460 | 2020 | France | Reunion | Madagascar | ERS6575378 |
| 202000461 | 2020 | France | Reunion | Madagascar | ERS6575379 |
| 202000462 | 2020 | France | Reunion | Madagascar | ERS6575380 |
| 202000464 | 2020 | France | Metropole | Unknown | ERS6575381 |
| 202000471 | 2020 | France | Metropole | None | ERS6575382 |
| 202000473 | 2020 | France | Metropole | Unknown | ERS6575383 |
| 202000511 | 2020 | France | Metropole | None | ERS6575384 |
| 202000529 | 2020 | France | Metropole | None | ERS6575385 |
| 202000532 | 2020 | France | Metropole | Unknown | ERS6575386 |
| 202000533 | 2020 | France | Metropole | Unknown | ERS6575387 |
| 202000561 | 2020 | France | Metropole | Unknown | ERS6575388 |

| | | | | | |
|---|---|---|---|---|---|
| 202000562 | 2020 | France | Metropole | Unknown | ERS6575389 |
| 202000571 | 2020 | France | Metropole | Unknown | ERS6575390 |
| 202000579 | 2020 | France | Metropole | Unknown | ERS6575391 |
| 202000604 | 2020 | France | Reunion | Madagascar | ERS6575945 |
| 202000605 | 2020 | France | Reunion | Madagascar | ERS6575946 |
| 202000617 | 2020 | France | Metropole | None | ERS6575947 |
| 202000618 | 2020 | France | Metropole | None | ERS6575948 |
| 202000619 | 2020 | France | Metropole | None | ERS6575949 |
| 202000642 | 2020 | France | Metropole | Mexico | ERS6575950 |
| 202000650 | 2020 | France | Metropole | Unknown | ERS6575951 |
| 202000651 | 2020 | France | Metropole | Unknown | ERS6575952 |
| 202000652 | 2020 | France | Metropole | Unknown | ERS6575953 |
| 202000654 | 2020 | France | Metropole | Unknown | ERS6575954 |
| 202000676 | 2020 | France | Metropole | Unknown | ERS6575955 |
| 202000715 | 2020 | France | Metropole | Unknown | ERS6575956 |
| 202000731 | 2020 | France | Metropole | Argentina | ERS6575957 |
| 202000757 | 2020 | France | Metropole | Unknown | ERS6575958 |
| 202000767 | 2020 | France | Metropole | Cameroon | ERS6575959 |
| 202000772 | 2020 | France | Metropole | India | ERS6575960 |
| 202000803 | 2020 | France | Metropole | Unknown | ERS6575961 |
| 202000821 | 2020 | France | Metropole | None | ERS6575962 |
| 202000823 | 2020 | France | Reunion | Unknown | ERS6575963 |
| 202000828 | 2020 | France | Metropole | Unknown | ERS6575964 |
| 202000882 | 2020 | France | Metropole | Unknown | ERS6575965 |
| 202000891 | 2020 | France | Metropole | Unknown | ERS6575966 |
| 202000895 | 2020 | France | Reunion | Madagascar | ERS6575967 |
| 202000912 | 2020 | France | Metropole | Mexico | ERS6575968 |
| 202000934 | 2020 | France | French Guiana | Unknown | ERS6575969 |
| 202000943 | 2020 | France | French Guiana | Unknown | ERS6575970 |
| 202000944 | 2020 | France | French Guiana | Unknown | ERS6575971 |
| 202000957 | 2020 | France | Metropole | None | ERS6575972 |
| 202000958 | 2020 | France | Metropole | Unknown | ERS6575973 |
| 202000959 | 2020 | France | Metropole | Vietnam | ERS6575974 |
| 202000971 | 2020 | France | Metropole | Unknown | ERS6575975 |
| 202000995 | 2020 | France | Metropole | Unknown | ERS6575976 |
| 202001001 | 2020 | France | Metropole | Unknown | ERS6575977 |
| 202001025 | 2020 | France | Metropole | Unknown | ERS6575978 |
| 202001047 | 2020 | France | Metropole | Guinea | ERS6575979 |
| 202001056 | 2020 | France | Metropole | Unknown | ERS6575980 |
| 202001057 | 2020 | France | Metropole | Unknown | ERS6576214 |
| 202001062 | 2020 | France | Metropole | Unknown | ERS6576215 |
| 202001063 | 2020 | France | Metropole | Unknown | ERS6576216 |
| 202001076 | 2020 | France | Metropole | Unknown | ERS6575981 |
| 202001077 | 2020 | France | Metropole | Unknown | ERS6575982 |
| 202001092 | 2020 | France | Metropole | Unknown | ERS6575983 |
| 202001104 | 2020 | France | Metropole | Unknown | ERS6575984 |
| 202001111 | 2020 | France | Metropole | None | ERS6575985 |
| 202001113 | 2020 | France | Metropole | Unknown | ERS12446242 |
| 202001114 | 2020 | France | Metropole | Unknown | ERS6575986 |
| 202001125 | 2020 | France | Metropole | Madagascar | ERS6575987 |
| 202001148 | 2020 | France | Metropole | Tanzania | ERS12446243 |
| 202001172 | 2020 | France | Metropole | New Caledonia | ERS6575988 |
| 202001191 | 2020 | France | Metropole | Unknown | ERS12446244 |
| 202001254 | 2020 | France | Metropole | Unknown | ERS6575989 |
| 202001265 | 2020 | France | Mayotte | Unknown | ERS6576217 |
| 202001321 | 2020 | France | Metropole | Unknown | ERS6575990 |
| 202001332 | 2020 | France | Metropole | Unknown | ERS6575991 |
| 202001380 | 2020 | France | Metropole | Unknown | ERS6575992 |
| 202001387 | 2020 | France | Reunion | Unknown | ERS6575993 |
| 202001405 | 2020 | France | Metropole | Unknown | ERS6575994 |
| 202001406 | 2020 | France | Metropole | Unknown | ERS6575995 |
| 202001421 | 2020 | France | Metropole | None | ERS6575996 |
| 202001422 | 2020 | France | Metropole | Unknown | ERS6575997 |
| 202001430 | 2020 | France | Metropole | None | ERS6576218 |
| 202001457 | 2020 | France | French Guiana | Unknown | ERS6575998 |
| 202001499 | 2020 | France | French Guiana | Unknown | ERS6575999 |
| 202001520 | 2020 | France | Metropole | Unknown | ERS6576000 |
| 202001551 | 2020 | France | Metropole | Benin | ERS6576001 |
| 202001554 | 2020 | France | Metropole | Benin | ERS6576002 |
| 202001572 | 2020 | France | Metropole | None | ERS6576003 |
| 202001573 | 2020 | France | Metropole | None | ERS6576004 |
| 202001574 | 2020 | France | Metropole | Unknown | ERS6576005 |
| 202001611 | 2020 | France | Metropole | Unknown | ERS6576006 |
| 202001623 | 2020 | France | Metropole | Unknown | ERS6576007 |
| 202001631 | 2020 | France | Metropole | Unknown | ERS6576008 |
| 202001638 | 2020 | France | Metropole | Unknown | ERS6576009 |
| 202001713 | 2020 | France | Metropole | Unknown | ERS6576010 |
| 202001772 | 2020 | France | Metropole | Mauritius | ERS6576219 |
| 202001781 | 2020 | France | Metropole | Cambodia | ERS6576011 |
| 202001783 | 2020 | France | Metropole | None | ERS6576220 |
| 202001820 | 2020 | France | Metropole | Cambodia | ERS6576221 |
| 202001835 | 2020 | France | French Guiana | Unknown | ERS6576222 |
| 202001911 | 2020 | France | Metropole | Unknown | ERS6576223 |
| 202001914 | 2020 | France | Metropole | None | ERS6576224 |
| 202001961 | 2020 | France | Metropole | Indonesia | ERS6576225 |
| 202001966 | 2020 | France | Metropole | Unknown | ERS6576226 |
| 202001995 | 2020 | France | Metropole | Unknown | ERS6576227 |
| 202002009 | 2020 | France | French Guiana | Unknown | ERS6576228 |
| 202002061 | 2020 | France | Metropole | Unknown | ERS6576229 |
| 202002063 | 2020 | France | Metropole | None | ERS6576230 |
| 202002129 | 2020 | France | French Guiana | Unknown | ERS6576231 |
| 202002140 | 2020 | France | Metropole | Unknown | ERS6576232 |
| 202002142 | 2020 | France | Metropole | Unknown | ERS6576233 |
| 202002145 | 2020 | France | Metropole | Unknown | ERS6576234 |
| 202002169 | 2020 | France | Metropole | Unknown | ERS6576235 |
| 202002188 | 2020 | France | Metropole | Unknown | ERS6576236 |
| 202002213 | 2020 | France | French Guiana | Unknown | ERS6576237 |
| 202002283 | 2020 | France | Metropole | Unknown | ERS6576238 |
| 202002357 | 2020 | France | French Guiana | Unknown | ERS6576239 |
| 202002358 | 2020 | France | French Guiana | Unknown | ERS6576240 |
| 202002377 | 2020 | France | Mayotte | None | ERS6576241 |
| 202002378 | 2020 | France | Mayotte | None | ERS6576242 |
| 202002382 | 2020 | France | Mayotte | None | ERS6576243 |

| | | | | | |
|---|---|---|---|---|---|
| 202002451 | 2020 | France | Metropole | Unknown | ERS6576244 |
| 202002597 | 2020 | France | Metropole | Unknown | ERS6576245 |
| 202002692 | 2020 | France | Metropole | Unknown | ERS6576246 |
| 202002717 | 2020 | France | Metropole | None | ERS6576247 |
| 202002865 | 2020 | France | French Guiana | Unknown | ERS6576248 |
| 202002975 | 2020 | France | Metropole | Unknown | ERS6576249 |
| 202003021 | 2020 | France | Metropole | Unknown | ERS6576250 |
| 202003031 | 2020 | France | Metropole | Unknown | ERS6576251 |
| 202003044 | 2020 | France | Metropole | Unknown | ERS6576252 |
| 202003088 | 2020 | France | Metropole | Unknown | ERS6576253 |
| 202003345 | 2020 | France | Metropole | Unknown | ERS6576254 |
| 202003459 | 2020 | France | Metropole | Unknown | ERS6576255 |
| 202003478 | 2020 | France | Metropole | None | ERS6576256 |
| 202003568 | 2020 | France | Mayotte | Unknown | ERS6576257 |
| 202003570 | 2020 | France | Mayotte | Unknown | ERS6576258 |
| 202003573 | 2020 | France | Mayotte | Unknown | ERS6576259 |
| 202003618 | 2020 | France | Metropole | Unknown | ERS6576260 |
| 202003644 | 2020 | France | Metropole | Unknown | ERS6576261 |
| 202003679 | 2020 | France | Metropole | Unknown | ERS6576262 |
| 202003706 | 2020 | France | Metropole | Unknown | ERS6576263 |
| 202003879 | 2020 | France | Metropole | Unknown | ERS6576264 |
| 202003882 | 2020 | France | French Guiana | French Guiana | ERS6576265 |
| 202003947 | 2020 | France | Metropole | None | ERS6576266 |
| 202003965 | 2020 | France | Metropole | None | ERS6576267 |
| 202003972 | 2020 | France | Metropole | None | ERS6576268 |
| 202003974 | 2020 | France | Metropole | Unknown | ERS6576269 |
| 202004050 | 2020 | France | Metropole | None | ERS6576270 |
| 202004125 | 2020 | France | Metropole | Unknown | ERS6576271 |
| 202004139 | 2020 | France | Metropole | Unknown | ERS6576272 |
| 202004184 | 2020 | France | Metropole | None | ERS6576273 |
| 202004237 | 2020 | France | Metropole | Unknown | ERS6576274 |
| 202004408 | 2020 | France | Metropole | None | ERS6576275 |
| 202004430 | 2020 | France | Metropole | Unknown | ERS6576276 |
| 202004441 | 2020 | France | Metropole | None | ERS6576277 |
| 202004492 | 2020 | France | Metropole | None | ERS6576278 |
| 202004493 | 2020 | France | Metropole | Unknown | ERS6576279 |
| 202004550 | 2020 | France | Metropole | Unknown | ERS12446245 |
| 202004649 | 2020 | France | Metropole | None | ERS6576280 |
| 202004795 | 2020 | France | Metropole | Unknown | ERS6576281 |
| 202004830 | 2020 | France | Reunion | Unknown | ERS6576282 |
| 202004834 | 2020 | France | Metropole | None | ERS6576283 |
| 202004884 | 2020 | France | Metropole | Unknown | ERS6576284 |
| 202004899 | 2020 | France | Metropole | Unknown | ERS6576286 |
| 202005096 | 2020 | France | Metropole | Senegal | ERS12446246 |
| 202005154 | 2020 | France | Reunion | None | ERS6576287 |
| 202005237 | 2020 | France | Metropole | None | ERS6576288 |
| 202005269 | 2020 | France | Metropole | Niger | ERS6576289 |
| 202005291 | 2020 | France | Metropole | Unknown | ERS6576290 |
| 202005297 | 2020 | France | Metropole | Unknown | ERS6576291 |
| 202005338 | 2020 | France | Metropole | Unknown | ERS6576292 |
| 202005339 | 2020 | France | Metropole | Unknown | ERS6576293 |
| 202005394 | 2020 | France | Metropole | None | ERS6576294 |
| 202005399 | 2020 | France | Metropole | Unknown | ERS6576295 |
| 202005406 | 2020 | France | Metropole | Unknown | ERS6576296 |
| 202005425 | 2020 | France | Metropole | Togo | ERS6576297 |
| 202005655 | 2020 | France | Metropole | Tunisia | ERS6576298 |
| 202005808 | 2020 | France | Metropole | Unknown | ERS6576299 |
| 202005851 | 2020 | France | Metropole | None | ERS6576300 |
| 202005861 | 2020 | France | Metropole | Unknown | ERS6576301 |
| 202005871 | 2020 | France | Metropole | None | ERS6576302 |
| 202005884 | 2020 | France | Metropole | Unknown | ERS6576303 |
| 202006001 | 2020 | France | Metropole | Unknown | ERS6576304 |
| 202006038 | 2020 | France | Metropole | Unknown | ERS6576305 |
| 202006235 | 2020 | France | Metropole | Unknown | ERS6576306 |
| 202006308 | 2020 | France | Metropole | Unknown | ERS6576307 |
| 202006311 | 2020 | France | Metropole | Unknown | ERS6576308 |
| 202006312 | 2020 | France | Metropole | Unknown | ERS6576309 |
| 202006637 | 2020 | France | Metropole | None | ERS6576310 |
| 202006729 | 2020 | France | Metropole | None | ERS6576311 |
| 202006730 | 2020 | France | Metropole | None | ERS6576312 |
| 202006740 | 2020 | France | Metropole | Unknown | ERS6576313 |
| 202006782 | 2020 | France | Metropole | Unknown | ERS6576314 |
| 202007026 | 2020 | France | Metropole | Morocco | ERS6576315 |
| 202007046 | 2020 | France | Metropole | Unknown | ERS6576316 |
| 202007051 | 2020 | France | Metropole | Unknown | ERS6576317 |
| 202007085 | 2020 | France | Metropole | Egypt | ERS6576318 |
| 202007105 | 2020 | France | Metropole | Unknown | ERS6576319 |
| 202007128 | 2020 | France | Metropole | Unknown | ERS6576320 |
| 202007186 | 2020 | France | Metropole | Unknown | ERS6576321 |
| 202007345 | 2020 | France | French Guiana | Unknown | ERS6576322 |
| 202007406 | 2020 | France | Metropole | Unknown | ERS6576323 |
| 202007417 | 2020 | France | Metropole | None | ERS6576324 |
| 202007434 | 2020 | France | Metropole | Unknown | ERS6576325 |
| 202007486 | 2020 | France | Metropole | None | ERS6576326 |
| 202007497 | 2020 | France | Metropole | Unknown | ERS6576327 |
| 202007629 | 2020 | France | Metropole | None | ERS6576328 |
| 202007640 | 2020 | France | Metropole | Unknown | ERS6576329 |
| 202007708 | 2020 | France | Metropole | Unknown | ERS6576330 |
| 202007774 | 2020 | France | Mayotte | Unknown | ERS6576331 |
| 202007845 | 2020 | France | Metropole | Unknown | ERS6576332 |
| 202007856 | 2020 | France | Metropole | None | ERS6576333 |
| 202007858 | 2020 | France | Metropole | Unknown | ERS6576334 |
| 202007876 | 2020 | France | Metropole | Unknown | ERS6576335 |
| 202007877 | 2020 | France | Metropole | Congo | ERS6576336 |
| 202007896 | 2020 | France | Metropole | Unknown | ERS6576337 |
| 202007960 | 2020 | France | Metropole | Unknown | ERS6576338 |
| 202007978 | 2020 | France | Metropole | Unknown | ERS6576339 |
| 202007982 | 2020 | France | Metropole | Unknown | ERS6576340 |
| 202008000 | 2020 | France | Metropole | None | ERS6576341 |
| 202008072 | 2020 | France | Metropole | Unknown | ERS6576342 |
| 202008077 | 2020 | France | Metropole | None | ERS6576343 |
| 202008091 | 2020 | France | Metropole | Unknown | ERS6576344 |
| 202008110 | 2020 | France | Metropole | Unknown | ERS6576345 |

| | | | | | |
|---|---|---|---|---|---|
| 202008118 | 2020 | France | Metropole | Unknown | ERS12446247 |
| 202008158 | 2020 | France | Metropole | None | ERS6576346 |
| 202008171 | 2020 | France | Metropole | Unknown | ERS6576347 |
| 202008214 | 2020 | France | Metropole | Unknown | ERS6576348 |
| 202008242 | 2020 | France | Metropole | Unknown | ERS6576349 |
| 202008246 | 2020 | France | Metropole | Unknown | ERS6576350 |
| 202008247 | 2020 | France | Metropole | Egypt | ERS6576351 |
| 202008296 | 2020 | France | French Guiana | Unknown | ERS6576352 |
| 202008337 | 2020 | France | Metropole | Unknown | ERS6576353 |
| 202008344 | 2020 | France | Metropole | Reunion | ERS6576354 |
| 202008346 | 2020 | France | Metropole | Unknown | ERS6576355 |
| 202008392 | 2020 | France | Metropole | Unknown | ERS6576357 |
| 202008421 | 2020 | France | Metropole | Unknown | ERS6576358 |
| 202008509 | 2020 | France | Metropole | Unknown | ERS6576359 |
| 202008533 | 2020 | France | Metropole | Unknown | ERS6576360 |
| 202008535 | 2020 | France | Metropole | Unknown | ERS6576361 |
| 202008564 | 2020 | France | Metropole | Unknown | ERS6576362 |
| 202008583 | 2020 | France | Metropole | Spain | ERS6576363 |
| 202008602 | 2020 | France | Metropole | Unknown | ERS6576364 |
| 202008603 | 2020 | France | Metropole | Unknown | ERS6576365 |
| 202008691 | 2020 | France | Metropole | None | ERS6578763 |
| 202008707 | 2020 | France | Metropole | Unknown | ERS6578764 |
| 202008716 | 2020 | France | Metropole | Central African Republic | ERS6578765 |
| 202100038 | 2020 | France | Metropole | Unknown | ERS12156937 |
| 202100124 | 2020 | France | Metropole | Unknown | ERS12156938 |
| 202100202 | 2020 | France | Mayotte | Unknown | ERS12156941 |
| 202100207 | 2020 | France | Mayotte | Unknown | ERS12156942 |
| 202100208 | 2020 | France | Mayotte | Unknown | ERS12156943 |
| 202100162 | 2021 | France | Metropole | Tanzania | ERS12156939 |
| 202100168 | 2021 | France | Metropole | Unknown | ERS12156940 |
| 202100233 | 2021 | France | Metropole | Mayotte | ERS12156944 |
| 202100239 | 2021 | France | Metropole | Senegal | ERS12156945 |
| 202100259 | 2021 | France | Metropole | Tanzania | ERS12156946 |
| 202100286 | 2021 | France | Metropole | Tanzania | ERS12156947 |
| 202100362 | 2021 | France | Metropole | Unknown | ERS12156948 |
| 202100373 | 2021 | France | Metropole | Unknown | ERS12156949 |
| 202100374 | 2021 | France | French Guiana | None | ERS12156950 |
| 202100396 | 2021 | France | Metropole | Unknown | ERS12156951 |
| 202100420 | 2021 | France | Metropole | Unknown | ERS12156952 |
| 202100452 | 2021 | France | Metropole | Unknown | ERS12156953 |
| 202100474 | 2021 | France | Metropole | None | ERS12156954 |
| 202100476 | 2021 | France | Metropole | Pakistan | ERS12156955 |
| 202100478 | 2021 | France | Metropole | Unknown | ERS12156956 |
| 202100480 | 2021 | France | Metropole | Unknown | ERS12156957 |
| 202100488 | 2021 | France | French Guiana | Unknown | ERS12156958 |
| 202100504 | 2021 | France | Metropole | Unknown | ERS12156959 |
| 202100558 | 2021 | France | Metropole | Unknown | ERS12156960 |
| 202100655 | 2021 | France | Metropole | Unknown | ERS12156961 |
| 202100656 | 2021 | France | Metropole | Unknown | ERS12156962 |
| 202100739 | 2021 | France | Metropole | India | ERS12156963 |
| 202100759 | 2021 | France | Metropole | Unknown | ERS12156964 |
| 202100791 | 2021 | France | Metropole | Unknown | ERS12156965 |
| 202100906 | 2021 | France | Metropole | None | ERS12156967 |
| 202100932 | 2021 | France | Metropole | Unknown | ERS12156968 |
| 202100945 | 2021 | France | Metropole | None | ERS12156969 |
| 202100975 | 2021 | France | Metropole | Unknown | ERS12156970 |
| 202100983 | 2021 | France | Metropole | Africa | ERS12156971 |
| 202101010 | 2021 | France | Metropole | Unknown | ERS12156972 |
| 202101155 | 2021 | France | Metropole | Unknown | ERS12156973 |
| 202101207 | 2021 | France | Metropole | Unknown | ERS12156974 |
| 202101222 | 2021 | France | Metropole | Unknown | ERS12156975 |
| 202101234 | 2021 | France | Metropole | Africa | ERS12156976 |
| 202101241 | 2021 | France | Metropole | None | ERS12156977 |
| 202101338 | 2021 | France | Metropole | Tanzania | ERS12156978 |
| 202101379 | 2021 | France | Metropole | Unknown | ERS12156979 |
| 202101412 | 2021 | France | Metropole | Unknown | ERS12156980 |
| 202101428 | 2021 | France | Metropole | Unknown | ERS12156981 |
| 202101449 | 2021 | France | Metropole | Africa | ERS12156982 |
| 202101453 | 2021 | France | French Guiana | Unknown | ERS12156983 |
| 202101454 | 2021 | France | French Guiana | Unknown | ERS12156984 |
| 202101455 | 2021 | France | French Guiana | Unknown | ERS12156985 |
| 202101477 | 2021 | France | Metropole | Rwanda | ERS12156986 |
| 202101541 | 2021 | France | Metropole | None | ERS12156987 |
| 202101576 | 2021 | France | Mayotte | Unknown | ERS12156988 |
| 202101579 | 2021 | France | Metropole | Unknown | ERS12156989 |
| 202101582 | 2021 | France | Mayotte | Unknown | ERS12156990 |
| 202101584 | 2021 | France | Mayotte | Unknown | ERS12156991 |
| 202101585 | 2021 | France | Mayotte | Unknown | ERS12156992 |
| 202101587 | 2021 | France | Mayotte | Unknown | ERS12156993 |
| 202101590 | 2021 | France | Mayotte | Unknown | ERS12156994 |
| 202101599 | 2021 | France | Metropole | None | ERS12156995 |
| 202101624 | 2021 | France | Metropole | Unknown | ERS12156996 |
| 202101651 | 2021 | France | Metropole | None | ERS12156997 |
| 202101673 | 2021 | France | Metropole | Unknown | ERS12156998 |
| 202101706 | 2021 | France | Metropole | Unknown | ERS12156999 |
| 202101718 | 2021 | France | Metropole | Unknown | ERS12157000 |
| 202101727 | 2021 | France | Metropole | None | ERS12157001 |
| 202101759 | 2021 | France | Metropole | Unknown | ERS12157002 |
| 202101760 | 2021 | France | Metropole | Unknown | ERS12157003 |
| 202101779 | 2021 | France | Metropole | Unknown | ERS12157004 |
| 202101806 | 2021 | France | Metropole | Unknown | ERS12157005 |
| 202101814 | 2021 | France | French Guiana | Unknown | ERS12157006 |
| 202101879 | 2021 | France | Metropole | None | ERS12157007 |
| 202101885 | 2021 | France | Metropole | Unknown | ERS12157008 |
| 202101895 | 2021 | France | Metropole | Unknown | ERS12157009 |
| 202101918 | 2021 | France | Metropole | Unknown | ERS12157010 |
| 202101930 | 2021 | France | Metropole | Unknown | ERS12157011 |
| 202101941 | 2021 | France | Metropole | Unknown | ERS12157012 |
| 202102013 | 2021 | France | Metropole | Unknown | ERS12157013 |
| 202102119 | 2021 | France | Metropole | Unknown | ERS12157014 |
| 202102131 | 2021 | France | Metropole | Mozambique | ERS12157015 |
| 202102160 | 2021 | France | Metropole | Mexico | ERS12157016 |
| 202102200 | 2021 | France | Metropole | Unknown | ERS12446248 |

| | | | | | |
|---|---|---|---|---|---|
| 202102275 | 2021 | France | Metropole | Unknown | ERS12157017 |
| 202102283 | 2021 | France | Metropole | None | ERS12157018 |
| 202102392 | 2021 | France | French Guiana | Unknown | ERS12157019 |
| 202102436 | 2021 | France | Metropole | Unknown | ERS12157020 |
| 202102474 | 2021 | France | Metropole | Unknown | ERS12157021 |
| 202102699 | 2021 | France | Metropole | Iraq | ERS12157022 |
| 202102707 | 2021 | France | Metropole | Unknown | ERS12157023 |
| 202102709 | 2021 | France | French Guiana | Unknown | ERS12157024 |
| 202102715 | 2021 | France | Metropole | Unknown | ERS12157025 |
| 202102776 | 2021 | France | Metropole | None | ERS12157026 |
| 202102820 | 2021 | France | Metropole | None | ERS12157027 |
| 202102823 | 2021 | France | Metropole | Unknown | ERS12157028 |
| 202102843 | 2021 | France | Metropole | None | ERS12157029 |
| 202102858 | 2021 | France | Metropole | Unknown | ERS12157030 |
| 202102980 | 2021 | France | Metropole | None | ERS12157031 |
| 202102993 | 2021 | France | Metropole | Unknown | ERS12157032 |
| 202103005 | 2021 | France | Metropole | None | ERS12157033 |
| 202103026 | 2021 | France | Metropole | None | ERS12157034 |
| 202103028 | 2021 | France | Metropole | Unknown | ERS12157035 |
| 202103103 | 2021 | France | Metropole | Unknown | ERS12157036 |
| 202103128 | 2021 | France | Metropole | Senegal | ERS12157037 |
| 202103134 | 2021 | France | Metropole | None | ERS12157038 |
| 202103154 | 2021 | France | Reunion | Unknown | ERS12157039 |
| 202103166 | 2021 | France | Metropole | None | ERS12157040 |
| 202103226 | 2021 | France | Metropole | None | ERS12157041 |
| 202103236 | 2021 | France | French Guiana | Unknown | ERS12157042 |
| 202103241 | 2021 | France | Metropole | None | ERS12157043 |
| 202103248 | 2021 | France | Metropole | None | ERS12157044 |
| 202103264 | 2021 | France | Metropole | Unknown | ERS12157045 |
| 202103266 | 2021 | France | Metropole | Benin | ERS12157046 |
| 202103281 | 2021 | France | Metropole | Africa | ERS12157047 |
| 202103290 | 2021 | France | Metropole | None | ERS12157048 |
| 202103328 | 2021 | France | Mayotte | Unknown | ERS12157049 |
| 202103329 | 2021 | France | Mayotte | Unknown | ERS12157050 |
| 202103342 | 2021 | France | Metropole | None | ERS12157051 |
| 202103369 | 2021 | France | Metropole | Unknown | ERS12157052 |
| 202103419 | 2021 | France | Metropole | None | ERS12157053 |
| 202103422 | 2021 | France | Metropole | Indonesia | ERS12157054 |
| 202103424 | 2021 | France | Metropole | Cameroon | ERS12157055 |
| 202103519 | 2021 | France | Metropole | Unknown | ERS12157056 |
| 202103563 | 2021 | France | Metropole | Unknown | ERS12157057 |
| 202103567 | 2021 | France | Metropole | Unknown | ERS12157058 |
| 202103655 | 2021 | France | Metropole | Unknown | ERS12157059 |
| 202103754 | 2021 | France | Metropole | Unknown | ERS12157060 |
| 202103844 | 2021 | France | Metropole | Unknown | ERS12157061 |
| 202103880 | 2021 | France | Metropole | None | ERS12157062 |
| 202103899 | 2021 | France | Metropole | Senegal | ERS12157063 |
| 202103930 | 2021 | France | Metropole | None | ERS12157064 |
| 202104044 | 2021 | France | Metropole | Egypt | ERS12157065 |
| 202104060 | 2021 | France | Metropole | Unknown | ERS12157066 |
| 202104078 | 2021 | France | Metropole | None | ERS12157067 |
| 202104261 | 2021 | France | Metropole | None | ERS12157068 |
| 202104281 | 2021 | France | Metropole | None | ERS12157069 |
| 202104322 | 2021 | France | Metropole | Unknown | ERS12157070 |
| 202104371 | 2021 | France | Metropole | Unknown | ERS12157071 |
| 202104519 | 2021 | France | Metropole | None | ERS12157072 |
| 202104674 | 2021 | France | Metropole | Unknown | ERS12157073 |
| 202104685 | 2021 | France | Metropole | Unknown | ERS12157074 |
| 202104717 | 2021 | France | Metropole | Unknown | ERS12157075 |
| 202104831 | 2021 | France | Metropole | None | ERS12157076 |
| 202104930 | 2021 | France | Metropole | Unknown | ERS12157077 |
| 202104991 | 2021 | France | Metropole | Unknown | ERS12157078 |
| 202105132 | 2021 | France | Metropole | Unknown | ERS12157079 |
| 202105166 | 2021 | France | Metropole | Unknown | ERS12157080 |
| 202105200 | 2021 | France | Metropole | Unknown | ERS12157081 |
| 202105282 | 2021 | France | Metropole | Cameroon | ERS12157082 |
| 202105284 | 2021 | France | Metropole | None | ERS12157083 |
| 202105285 | 2021 | France | Metropole | None | ERS12157084 |
| 202105293 | 2021 | France | Metropole | Unknown | ERS12157085 |
| 202105294 | 2021 | France | Metropole | Unknown | ERS12157086 |
| 202105373 | 2021 | France | Metropole | Unknown | ERS12157087 |
| 202105403 | 2021 | France | Metropole | Morocco | ERS12157088 |
| 202105408 | 2021 | France | Metropole | Unknown | ERS12157089 |
| 202105424 | 2021 | France | Metropole | Unknown | ERS12157090 |
| 202105426 | 2021 | France | Metropole | Unknown | ERS12157091 |
| 202105440 | 2021 | France | Metropole | Unknown | ERS12157092 |
| 202105446 | 2021 | France | Metropole | Unknown | ERS12157093 |
| 202105523 | 2021 | France | Metropole | None | ERS12157094 |
| 202105574 | 2021 | France | Metropole | None | ERS12157095 |
| 202105576 | 2021 | France | Metropole | Morocco | ERS12157096 |
| 202105608 | 2021 | France | Metropole | Côte d'Ivoire | ERS12157097 |
| 202105725 | 2021 | France | Metropole | Unknown | ERS12157098 |
| 202105756 | 2021 | France | Metropole | Unknown | ERS12157099 |
| 202105763 | 2021 | France | Metropole | Senegal | ERS12157100 |
| 202105800 | 2021 | France | Metropole | Unknown | ERS12157101 |
| 202105814 | 2021 | France | Metropole | None | ERS12157102 |
| 202105817 | 2021 | France | Metropole | Unknown | ERS12157103 |
| 202105818 | 2021 | France | Metropole | Unknown | ERS12157104 |
| 202105866 | 2021 | France | Metropole | Morocco | ERS12157105 |
| 202105873 | 2021 | France | Metropole | Tanzania | ERS12157106 |
| 202105882 | 2021 | France | Metropole | Unknown | ERS12157107 |
| 202105919 | 2021 | France | Metropole | Egypt | ERS12157108 |
| 202105920 | 2021 | France | Metropole | Morocco | ERS12157109 |
| 202105953 | 2021 | France | Metropole | Unknown | ERS12157110 |
| 202105986 | 2021 | France | Metropole | None | ERS12157111 |
| 202106000 | 2021 | France | Metropole | Unknown | ERS12157112 |
| 202106014 | 2021 | France | Metropole | Morocco | ERS12157113 |
| 202106029 | 2021 | France | Metropole | Unknown | ERS12157114 |
| 202106050 | 2021 | France | Metropole | Kenya | ERS12157115 |
| 202106151 | 2021 | France | Mayotte | Unknown | ERS12157116 |
| 202106153 | 2021 | France | Mayotte | Unknown | ERS12157117 |
| 202106154 | 2021 | France | Mayotte | Unknown | ERS12157118 |
| 202106157 | 2021 | France | Mayotte | Unknown | ERS12157119 |

| | | | | | |
|---|---|---|---|---|---|
| 202106161 | 2021 | France | Mayotte | Unknown | ERS12157120 |
| 202106219 | 2021 | France | Metropole | None | ERS12157121 |
| 202106247 | 2021 | France | Metropole | Unknown | ERS12157122 |
| 202106248 | 2021 | France | Metropole | None | ERS12157123 |
| 202106255 | 2021 | France | Metropole | Unknown | ERS12157124 |
| 202106287 | 2021 | France | Metropole | Gambia | ERS12157125 |
| 202106289 | 2021 | France | Metropole | Unknown | ERS12157126 |
| 202106316 | 2021 | France | Metropole | Unknown | ERS12157127 |
| 202106338 | 2021 | France | Metropole | Unknown | ERS12157128 |
| 202106341 | 2021 | France | Metropole | Unknown | ERS12157129 |
| 202106367 | 2021 | France | Metropole | Lebanon | ERS12157130 |
| 202106388 | 2021 | France | Metropole | Morocco | ERS12157131 |
| 202106433 | 2021 | France | Metropole | Unknown | ERS12157132 |
| 202106455 | 2021 | France | Metropole | None | ERS12157133 |
| 202106464 | 2021 | France | Metropole | Unknown | ERS12157134 |
| 202106466 | 2021 | France | Metropole | Morocco | ERS12157135 |
| 202106503 | 2021 | France | Metropole | None | ERS12157136 |
| 202106524 | 2021 | France | Metropole | Unknown | ERS12157137 |
| 202106544 | 2021 | France | Metropole | Morocco | ERS12157138 |
| 202106573 | 2021 | France | Metropole | Unknown | ERS12157139 |
| 202106613 | 2021 | France | Reunion | None | ERS12157140 |
| 202106615 | 2021 | France | Metropole | None | ERS12157141 |
| 202106629 | 2021 | France | Metropole | Unknown | ERS12157142 |
| 202106648 | 2021 | France | Metropole | Morocco | ERS12157143 |
| 202106652 | 2021 | France | Metropole | Unknown | ERS12157144 |
| 202106664 | 2021 | France | Metropole | None | ERS12157145 |
| 202106665 | 2021 | France | Metropole | Egypt | ERS12157146 |
| 202106668 | 2021 | France | Metropole | Côte d'Ivoire | ERS12157147 |
| 202106716 | 2021 | France | Metropole | Spain | ERS12157148 |
| 202106740 | 2021 | France | Metropole | Unknown | ERS12157149 |
| 202106756 | 2021 | France | Metropole | Mali | ERS12157150 |
| 202106764 | 2021 | France | Metropole | None | ERS12157151 |
| 202106790 | 2021 | France | Metropole | Mauritania | ERS12157152 |
| 202106829 | 2021 | France | Metropole | Unknown | ERS12157153 |
| 202106837 | 2021 | France | Metropole | Unknown | ERS12157154 |
| 202106851 | 2021 | France | Metropole | Unknown | ERS12157155 |
| 202106852 | 2021 | France | Reunion | Unknown | ERS12157156 |
| 202106938 | 2021 | France | Metropole | Senegal | ERS12157157 |
| 202106939 | 2021 | France | Metropole | Unknown | ERS12157158 |
| 202106953 | 2021 | France | Metropole | Egypt | ERS12157159 |
| 202106954 | 2021 | France | Metropole | Morocco | ERS12157160 |
| 202106956 | 2021 | France | Metropole | Unknown | ERS12157161 |
| 202106959 | 2021 | France | Metropole | Morocco | ERS12157162 |
| 202106961 | 2021 | France | Metropole | Unknown | ERS12157163 |
| 202106985 | 2021 | France | Metropole | None | ERS12157164 |
| 202107022 | 2021 | France | Metropole | None | ERS12157165 |
| 202107027 | 2021 | France | Metropole | Unknown | ERS12157166 |
| 202107051 | 2021 | France | Metropole | None | ERS12157167 |
| 202107052 | 2021 | France | Metropole | None | ERS12157168 |
| 202107111 | 2021 | France | Metropole | Unknown | ERS12157169 |
| 202107115 | 2021 | France | Metropole | None | ERS12157170 |
| 202107119 | 2021 | France | Metropole | Unknown | ERS12157171 |
| 202107135 | 2021 | France | Metropole | Unknown | ERS12157172 |
| 202107156 | 2021 | France | Metropole | Unknown | ERS12157173 |
| 202107188 | 2021 | France | Metropole | Unknown | ERS12157174 |
| 202107227 | 2021 | France | Metropole | Unknown | ERS12157175 |
| 202107257 | 2021 | France | Metropole | Unknown | ERS12157176 |
| 202107258 | 2021 | France | Metropole | Unknown | ERS12157177 |
| 202107260 | 2021 | France | Metropole | Unknown | ERS12157178 |
| 202107323 | 2021 | France | Metropole | Unknown | ERS12157179 |
| 202107375 | 2021 | France | Metropole | Unknown | ERS12157180 |
| 202107377 | 2021 | France | Metropole | Unknown | ERS12157181 |
| 202107395 | 2021 | France | Metropole | Unknown | ERS12157182 |
| 202107415 | 2021 | France | Metropole | Unknown | ERS12157183 |
| 202107446 | 2021 | France | Metropole | None | ERS12157184 |
| 202107464 | 2021 | France | Metropole | Unknown | ERS12157185 |
| 202107466 | 2021 | France | Metropole | None | ERS12157186 |
| 202107510 | 2021 | France | Metropole | None | ERS12157187 |
| 202107511 | 2021 | France | Metropole | Unknown | ERS12157188 |
| 202107521 | 2021 | France | Metropole | Unknown | ERS12157189 |
| 202107554 | 2021 | France | Metropole | Morocco | ERS12157190 |
| 202107563 | 2021 | France | Metropole | None | ERS12157191 |
| 202107582 | 2021 | France | Metropole | Unknown | ERS12157192 |
| 202107585 | 2021 | France | Metropole | Unknown | ERS12157193 |
| 202107606 | 2021 | France | Metropole | Morocco | ERS12157194 |
| 202107658 | 2021 | France | Metropole | Unknown | ERS12157195 |
| 202107660 | 2021 | France | Metropole | Unknown | ERS12157196 |
| 202107666 | 2021 | France | Metropole | None | ERS12157197 |
| 202107677 | 2021 | France | Metropole | None | ERS12157198 |
| 202107704 | 2021 | France | Metropole | Morocco | ERS12157199 |
| 202107705 | 2021 | France | Metropole | Unknown | ERS12157200 |
| 202107712 | 2021 | France | Metropole | Unknown | ERS12157201 |
| 202107713 | 2021 | France | Metropole | Unknown | ERS12157202 |
| 202107714 | 2021 | France | Metropole | Unknown | ERS12157203 |
| 202107720 | 2021 | France | Metropole | Morocco | ERS12157204 |
| 202107776 | 2021 | France | Metropole | Unknown | ERS12157205 |
| 202107780 | 2021 | France | Metropole | Unknown | ERS12157206 |
| 202107815 | 2021 | France | Metropole | Morocco | ERS12157207 |
| 202107816 | 2021 | France | Metropole | Morocco | ERS12157208 |
| 202107818 | 2021 | France | Metropole | Unknown | ERS12157209 |
| 202107828 | 2021 | France | Metropole | Unknown | ERS12157210 |
| 202107834 | 2021 | France | Metropole | Unknown | ERS12157211 |
| 202107845 | 2021 | France | Metropole | Unknown | ERS12157212 |
| 202107853 | 2021 | France | Metropole | Unknown | ERS12157213 |
| 202107855 | 2021 | France | Metropole | Unknown | ERS12157214 |
| 202107858 | 2021 | France | Metropole | Unknown | ERS12157215 |
| 202107873 | 2021 | France | Metropole | None | ERS12157216 |
| 202107902 | 2021 | France | Metropole | Unknown | ERS12157217 |
| 202107916 | 2021 | France | Metropole | None | ERS12157218 |
| 202107972 | 2021 | France | Metropole | Unknown | ERS12157219 |
| 202107994 | 2021 | France | Metropole | None | ERS12157220 |
| 202107995 | 2021 | France | Metropole | Unknown | ERS12157221 |
| 202107996 | 2021 | France | Metropole | Unknown | ERS12157222 |

| | | | | | |
|---|---|---|---|---|---|
| 202107997 | 2021 | France | Metropole | Unknown | ERS12157223 |
| 202107998 | 2021 | France | Metropole | Unknown | ERS12157224 |
| 202108011 | 2021 | France | Metropole | Unknown | ERS12157225 |
| 202108032 | 2021 | France | Metropole | None | ERS12157226 |
| 202108051 | 2021 | France | Metropole | Unknown | ERS12157227 |
| 202108086 | 2021 | France | Metropole | Unknown | ERS12446249 |
| 202108119 | 2021 | France | Metropole | None | ERS12446250 |
| 202108174 | 2021 | France | Metropole | Unknown | ERS12446251 |
| 202108214 | 2021 | France | Metropole | Unknown | ERS12446252 |
| 202108221 | 2021 | France | Metropole | Unknown | ERS12446253 |
| 202108294 | 2021 | France | Metropole | Unknown | ERS12446254 |
| 202108300 | 2021 | France | Metropole | Unknown | ERS12446255 |
| 202108312 | 2021 | France | Metropole | Unknown | ERS12446256 |
| 202108318 | 2021 | France | Metropole | Unknown | ERS12446257 |
| 202108321 | 2021 | France | Metropole | Unknown | ERS12446258 |
| 202108326 | 2021 | France | Metropole | None | ERS12446259 |
| 202108340 | 2021 | France | Metropole | Unknown | ERS12446260 |
| 202108363 | 2021 | France | Metropole | Unknown | ERS12446261 |
| 202108364 | 2021 | France | Metropole | Unknown | ERS12446262 |
| 202108395 | 2021 | France | Metropole | Unknown | ERS12446263 |
| 202108407 | 2021 | France | Metropole | Morocco | ERS12446264 |
| 202108410 | 2021 | France | Metropole | Unknown | ERS12446265 |
| 202108451 | 2021 | France | Metropole | Unknown | ERS12446266 |
| 202108481 | 2021 | France | Metropole | None | ERS12446267 |
| 202108511 | 2021 | France | Metropole | Senegal | ERS12446268 |
| 202108535 | 2021 | France | Metropole | None | ERS12446269 |
| 202108558 | 2021 | France | Metropole | Unknown | ERS12446270 |
| 202108630 | 2021 | France | Metropole | Unknown | ERS12446271 |
| 202108658 | 2021 | France | Metropole | Morocco | ERS12446272 |
| 202108659 | 2021 | France | Metropole | Unknown | ERS12446273 |
| 202108670 | 2021 | France | Metropole | Unknown | ERS12446274 |
| 202108687 | 2021 | France | Metropole | Unknown | ERS12446275 |
| 202108690 | 2021 | France | Metropole | Unknown | ERS12446276 |
| 202108691 | 2021 | France | Metropole | Unknown | ERS12446277 |
| 202108692 | 2021 | France | Metropole | None | ERS12446278 |
| 202108693 | 2021 | France | Metropole | None | ERS12446279 |
| 202108694 | 2021 | France | Metropole | Unknown | ERS12446280 |
| 202108695 | 2021 | France | Metropole | Unknown | ERS12446281 |
| 202108696 | 2021 | France | Metropole | Unknown | ERS12446282 |
| 202108697 | 2021 | France | Metropole | Morocco | ERS12446283 |
| 202108698 | 2021 | France | Metropole | None | ERS12446284 |
| 202108699 | 2021 | France | Metropole | Unknown | ERS12446285 |
| 202108700 | 2021 | France | Metropole | None | ERS12446286 |
| 202108701 | 2021 | France | Metropole | Unknown | ERS12446287 |
| 202108702 | 2021 | France | Metropole | Unknown | ERS12446288 |
| 202108795 | 2021 | France | Metropole | Unknown | ERS12446289 |
| 202108801 | 2021 | France | Metropole | Unknown | ERS12446290 |
| 202108853 | 2021 | France | Metropole | Morocco | ERS12446291 |
| 202108894 | 2021 | France | Metropole | Unknown | ERS12446292 |
| 202108895 | 2021 | France | Metropole | Unknown | ERS12446293 |
| 202108919 | 2021 | France | Metropole | None | ERS12446294 |
| 202108928 | 2021 | France | Metropole | Unknown | ERS12446295 |
| 202108943 | 2021 | France | Metropole | Unknown | ERS12446296 |
| 202108950 | 2021 | France | Metropole | Mayotte | ERS12446297 |
| 202108953 | 2021 | France | Metropole | Unknown | ERS12446298 |
| 202109025 | 2021 | France | Metropole | Lebanon | ERS12446299 |
| 202109058 | 2021 | France | Metropole | Unknown | ERS12446300 |
| 202109152 | 2021 | France | Metropole | None | ERS12446301 |
| 202109163 | 2021 | France | Metropole | Unknown | ERS12446302 |
| 202109182 | 2021 | France | Metropole | Unknown | ERS12446303 |
| 202109203 | 2021 | France | Metropole | Unknown | ERS12446304 |
| 202109232 | 2021 | France | Metropole | Unknown | ERS12446305 |
| 202109345 | 2021 | France | Metropole | None | ERS12446306 |
| 202109371 | 2021 | France | Metropole | Unknown | ERS12446307 |
| 202109380 | 2021 | France | Metropole | Morocco | ERS12446308 |
| 202109410 | 2021 | France | Metropole | Unknown | ERS12446309 |
| 202109419 | 2021 | France | Metropole | Unknown | ERS12446310 |
| 202109427 | 2021 | France | Metropole | Morocco | ERS12446311 |
| 202109470 | 2021 | France | Metropole | Egypt | ERS12446312 |
| 202109489 | 2021 | France | Metropole | Morocco | ERS12446313 |
| 202109504 | 2021 | France | Metropole | Egypt | ERS12446314 |
| 202109540 | 2021 | France | Metropole | Unknown | ERS12446315 |
| 202109570 | 2021 | France | Metropole | Unknown | ERS12446316 |
| 202109583 | 2021 | France | Metropole | Morocco | ERS12446317 |
| 202109588 | 2021 | France | Metropole | Unknown | ERS12446318 |
| 202109651 | 2021 | France | Metropole | None | ERS12446319 |
| 202109656 | 2021 | France | Metropole | Unknown | ERS12446320 |
| 202109793 | 2021 | France | Metropole | Morocco | ERS12446321 |
| 202109813 | 2021 | France | Metropole | None | ERS12446322 |
| 202109848 | 2021 | France | Metropole | Unknown | ERS12446323 |
| 202109849 | 2021 | France | Metropole | Unknown | ERS12446324 |
| 202109862 | 2021 | France | Metropole | Senegal | ERS12446325 |
| 202109869 | 2021 | France | Metropole | Unknown | ERS12446326 |
| 202109870 | 2021 | France | Metropole | Unknown | ERS12446327 |
| 202109898 | 2021 | France | Metropole | Unknown | ERS12446328 |
| 202109904 | 2021 | France | Metropole | Egypt | ERS12446329 |
| 202109909 | 2021 | France | Metropole | Unknown | ERS12446330 |
| 202109925 | 2021 | France | Metropole | Tanzania | ERS12446331 |
| 202109932 | 2021 | France | Metropole | Unknown | ERS12446332 |
| 202109943 | 2021 | France | Metropole | Unknown | ERS12446333 |
| 202109944 | 2021 | France | Metropole | Morocco | ERS12446334 |
| 202109960 | 2021 | France | Metropole | Unknown | ERS12446335 |
| 202109980 | 2021 | France | Metropole | Unknown | ERS12446336 |
| 202109992 | 2021 | France | Metropole | None | ERS12446337 |
| 202109997 | 2021 | France | Metropole | None | ERS12446338 |
| 202110006 | 2021 | France | Metropole | Unknown | ERS12446339 |
| 202110015 | 2021 | France | Metropole | Morocco | ERS12446340 |
| 202110034 | 2021 | France | Metropole | Unknown | ERS12446341 |
| 202110049 | 2021 | France | Metropole | Morocco | ERS12446342 |
| 202110090 | 2021 | France | Metropole | Morocco | ERS12446343 |
| 202110107 | 2021 | France | Metropole | Algeria | ERS12446344 |
| 202110115 | 2021 | France | Metropole | Unknown | ERS12446345 |
| 202110142 | 2021 | France | Metropole | Unknown | ERS12446346 |

| | | | | | |
|---|---|---|---|---|---|
| 202110146 | 2021 | France | Metropole | Unknown | ERS12446347 |
| 202110148 | 2021 | France | Metropole | Morocco | ERS12446348 |
| 202110149 | 2021 | France | Metropole | Morocco | ERS12446349 |
| 202110175 | 2021 | France | Metropole | Unknown | ERS12446350 |
| 202110231 | 2021 | France | Mayotte | Unknown | ERS12446351 |
| 202110232 | 2021 | France | Mayotte | Unknown | ERS12446352 |
| 202110233 | 2021 | France | Mayotte | Unknown | ERS12446353 |
| 202110234 | 2021 | France | Mayotte | Unknown | ERS12446354 |
| 202110237 | 2021 | France | Mayotte | Unknown | ERS12446355 |
| 202110251 | 2021 | France | Metropole | Unknown | ERS12446356 |
| 202110297 | 2021 | France | Metropole | Senegal | ERS12446357 |
| 202110341 | 2021 | France | Metropole | Unknown | ERS12446358 |
| 202110347 | 2021 | France | Metropole | Unknown | ERS12446359 |
| 202110403 | 2021 | France | Metropole | Unknown | ERS12446360 |
| 202110415 | 2021 | France | Metropole | None | ERS12446361 |
| 202110472 | 2021 | France | Metropole | None | ERS12446362 |
| 202110475 | 2021 | France | Metropole | Unknown | ERS12446363 |
| 202110554 | 2021 | France | Metropole | Unknown | ERS12446364 |
| 202110584 | 2021 | France | Metropole | Unknown | ERS12446365 |
| 202110605 | 2021 | France | Metropole | Egypt | ERS12446366 |
| 202110616 | 2021 | France | Metropole | Unknown | ERS12446367 |
| 202110617 | 2021 | France | Metropole | Unknown | ERS12446368 |
| 202110618 | 2021 | France | Metropole | Unknown | ERS12446369 |
| 202110641 | 2021 | France | Metropole | Unknown | ERS12446370 |
| 202110650 | 2021 | France | Metropole | None | ERS12446371 |
| 202110656 | 2021 | France | Metropole | Unknown | ERS12446372 |
| 202110674 | 2021 | France | Metropole | Unknown | ERS12446373 |
| 202110699 | 2021 | France | Metropole | None | ERS12446374 |
| 202110735 | 2021 | France | Metropole | Unknown | ERS12446375 |
| 202110745 | 2021 | France | Metropole | Unknown | ERS12446376 |
| 202110841 | 2021 | France | Metropole | Spain | ERS12446377 |
| 202110865 | 2021 | France | Metropole | Unknown | ERS12446378 |
| 202110869 | 2021 | France | Metropole | Unknown | ERS12446379 |
| 202110872 | 2021 | France | Metropole | Unknown | ERS12446380 |
| 202110877 | 2021 | France | Metropole | Unknown | ERS12446381 |
| 202110883 | 2021 | France | Metropole | Unknown | ERS12446382 |
| 202110898 | 2021 | France | Metropole | Unknown | ERS12446383 |
| 202110909 | 2021 | France | Metropole | Unknown | ERS12446384 |
| 202110959 | 2021 | France | Metropole | Unknown | ERS12446385 |
| 202110960 | 2021 | France | Metropole | Unknown | ERS12446386 |
| 202110961 | 2021 | France | Metropole | Unknown | ERS12446387 |
| 202110978 | 2021 | France | Metropole | Africa | ERS12446388 |
| 202110980 | 2021 | France | Metropole | Africa | ERS12446389 |
| 202110981 | 2021 | France | Metropole | Africa | ERS12446390 |
| 202110987 | 2021 | France | Metropole | Egypt | ERS12446391 |
| 202111030 | 2021 | France | Metropole | Unknown | ERS12446392 |
| 202111080 | 2021 | France | Metropole | Unknown | ERS12446393 |
| 202111082 | 2021 | France | Metropole | Unknown | ERS12446394 |
| 202111134 | 2021 | France | Metropole | Unknown | ERS12446395 |
| 202111144 | 2021 | France | Metropole | Unknown | ERS12446396 |
| 202111148 | 2021 | France | Metropole | None | ERS12446397 |
| 202111160 | 2021 | France | Metropole | Unknown | ERS12446398 |
| 202111184 | 2021 | France | Metropole | None | ERS12446399 |
| 202111192 | 2021 | France | Metropole | Djibouti | ERS12446400 |
| 202111254 | 2021 | France | Metropole | Mozambique | ERS12446401 |
| 202111298 | 2021 | France | Metropole | Nigeria | ERS12446402 |
| 202200007 | 2021 | France | Metropole | None | ERS12446403 |
| 202200109 | 2021 | France | Metropole | Unknown | ERS12446404 |
| 202200139 | 2021 | France | Metropole | Egypt | ERS12446405 |
| 202200143 | 2021 | France | Metropole | Unknown | ERS12446406 |
